# Virtual aquatic ecology: stepped simulations of gross production within constraints explain biomass changes

**DOI:** 10.64898/2025.12.01.691727

**Authors:** A.J.R. Cotter

**Affiliations:** Independent researcher, Fishworld Science, Broadford, Victoria, Australia 3658

**Keywords:** aquatic ecosystems, multi-species modeling, linear programming, generalized Lotka-Volterra model, gross production, ecological risk assessment, nutrient recycling, biomass recycling, fisheries, seasonal succession, aquatic ecosystems, multi-species modeling, linear programming, generalized Lotka-Volterra model, gross production, ecological risk assessment, nutrient recycling, biomass recycling, fisheries, seasonal succession

## Abstract

A simulator, ‘ECOLPS’ in R, is developed and trialed for ecological studies of closed aquatic ecosystems. Its constraint-based approach contrasts with function-based models widely applied in ecology. Total gross production (ΣGP) by ‘wild components’ (= species/life stages, grouped by ecological roles) is maximized within constraints over short time steps using linear programming, thereby enabling simulations of opportunistic growth and harvesting by competing components. Constraints use integrated terms from an 𝓃-component generalization of the Lotka-Volterra (LV) predator-prey model, and from mass-based models for fisheries, non-living organics, nutrients and essential habitats. Seasonality uses programmed temperature and light indices. Trial simulations, total 12, sought first to confirm conformance with LV theory. Further trials of gradually increasing ecosystem complexity were designed to simulate wellknown ecological events. They found biomass oscillations depending on starting biomasses and seasons, predator satiation, competitive exclusion, predator diets dependent on available prey, instability of unconstrained 3-level food chains, limitation of GP by essential habitats and nutrients, recycling of nutrient and biomass, trophic cascades and wasp-waist systems caused by fishing, and seasonal succession. Step-wise studies of simulated series explained results. A ‘constrained-GP’ hypothesis is proposed. Priorities for further developments are suggested. ECOLPS simulations could support cause-and-effect investigations, field work, and aquatic ecological risk assessments.

---

Modeling of aquatic ecosystems is widely practiced with the general aim of improving understanding of living processes and their vulnerabilities, most notably, to fishing and projected climate change (1–3). Despite the diversity (4, 5) of available models, many are ‘function-based’, relying on mathematical functions to represent relationships among living components and environmental variables. The dubious implication is that the often wide-ranging fluctuations that can occur in wild populations (6–8) are merely mechanical responses to changing circumstances. Solving function-based models is typically complex, possibly requiring simplifications, iterative numerical methods, weakly justified statistical distributions, a long-term ecological equilibrium, and an assumed lack of chaotic variation which can arise even from simple functions (9). The patchy use of ecosystem-model projections for decision making (2, 10) may partly stem from these, and other (11) significant difficulties.

The importance of constraints in ecology has long been acknowledged (12, 13) though seldom used by modelers, judging from the cited reviews. Constraints, when active, are, by definition, intrusive and restrictive for living organisms so responses are automatic, not just implicit as when a living response is calculated mathematically from other variables. Furthermore, constraints can be linear functions of a rate or quantity, making them simple to evaluate. These merits suggest simulation of an ecosystem with a ‘constraint-based’ approach involving optimization of an objective function indicative of healthy functioning, subject only to modeled constraints bounding a feasible region for all living components. Among optimization techniques (14, 15), linear programming (LP) stands out for efficiency with many variables and constraints. Hitherto, LP has seen little use in ecology since early applications for policy optimization (16).

This paper develops and tests an algorithm to simulate gross production, relative organic biomasses, harvesting, etc. in simple, conceptual aquatic-ecosystems populated only by autotrophs and heterotrophs. The ecosystem is closed to fluxes of living organisms and materials and has only one ‘water’ and one ‘non-living organics’ component; these restrictions avoid the need for mass-transfer functions. The LP step is carried out over short, unit time intervals Δ with calculated results enabling updated constraint limits for the next step, and so on as required. Stepwise simulations allow output time series to be highly flexible despite the linearity requirements of LP, as well as facilitating analyses of cause and effect. Software in R (17) called ‘Ecological linear programming system’ or ECOLPS, was developed in parallel with the algorithm. Since both were thought novel, their capabilities and ecological realism were assessed initially by challenging ECOLPS to simulate a Lotka-Volterra 2-component system in accordance with standard theory, and to carry out other simple tests as expected. Further trial simulations, 12 in all with gradually increasing complexity, were designed to reveal ecological events known to occur in aquatic ecosystems. ECOLPS processed each trial in seconds. A ‘constrained-GP’ hypothesis is put forward to summarize understanding gained. Possible uses for, and development of constraint-based simulations are discussed.

## Theory

### Outline, notation and definitions

Theory begins by defining gross production (GP) and its dispersal routes, justifying measurement of ecosystem health with total GP, ‘ΣGP’, and describing the LP problem. The Lotka-Volterra (LV) predatorprey model is summarized (i) for comparisons with trial simulations and (ii) as the basis for a generalization, the ‘GLV’, to biomasses of **𝓃**_W_ living components and **𝓃**_F_ fisheries. (A previous generalization (18, App. 3) was too restrictive.) The GLV supports feeding, fishing, and growth rate constraints. Other mass models needed for constraints deal with NLO receiving all dropped biomass; essential nutrients in water, ‘H2O’, NLO and living tissues; and essential habitats. Reflecting the broad ambit of the GLV, the term *food* is used to include grazed plants, heterotrophic prey, components targeted by fisheries, and NLO eaten by heterotrophic *scavengers. Consumers* include heterotrophs and fisheries. Three types of constraint are distinguished:

- Rate constraints on physiological, physical or chemical processes contributing to GP.
- Resource constraints caused by resources used up by GP, such as habitats and nutrients.
- Food constraints caused by total consumption of a food component harvested by one or more consumers. Food constraints link GP of heterotrophs with harvesting of their foods. Updating of constraints after each Δ_*t*_ using slack variables is described. Simulations may include seasonal variations. These are based on 0-to-1 temperature *T*_*t*_ and light *I*_*t*_ indices supplied over time in a supplementary ‘Environ.file’ read by ECOLPS after each LP step.

The extensive notation needed is obtained with different fonts and cases (l.c. or u.c.). A *component*, a term from ERAEF (19, 20), is a relevant living or non-living part of an ecosystem. A living component, or *wild w*, consists of all the individuals belonging to a set of one or more living species and/or life stages defined by shared methods of production, needs for resources, vulnerabilities to harvesters and, possibly, other features depending on information available. *w* is a member of the set of all wilds, denoted *w* ∈ W. W has overlapping subsets depending on trophic roles. So *w* may also be a group of autotrophs *a* ∈ A, heterotrophs *h* ∈ H, foods *m* ∈ M and/or of consumers *g* ∈ G = H ∪ F. For reference, Table 1 lists living and non-living components as sets (upright, u.c.), their members (italic, l.c.), interrelationships, and required numbers (**𝓃**) of members for viable simulations. Table 2 lists associated time-dependent, non-negative (italic, u.c.) and cumulative (calligraphic, u.c.) variables with definitions and uses. Table 3 summarizes model parameters (calligraphic, l.c.), all non-negative. Table 4 lists special symbols and abbreviations used.

**Table 1.**
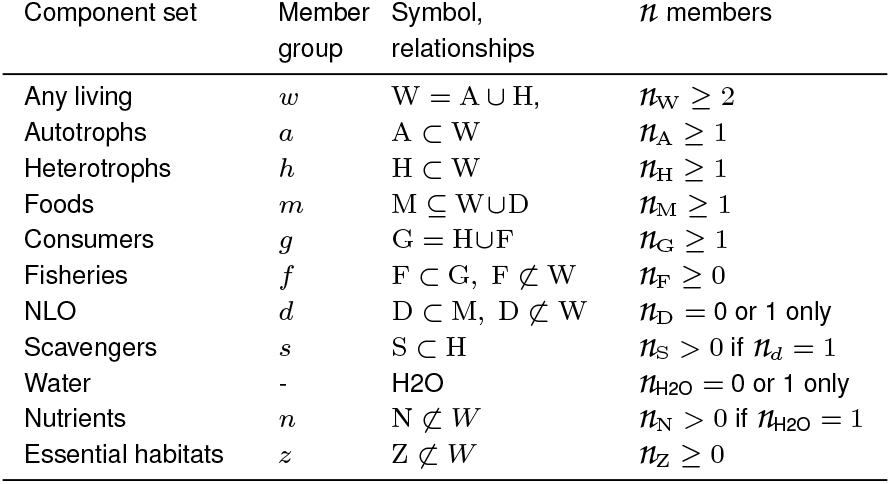
Components for LP simulations.

**Table 2.** Time-dependent, non-negative variables for LP simulations.

| Name | Symbol | Definition | Components described |
| --- | --- | --- | --- |
| Biomass | $B$ | organic weight | $w \in W$ or $d \in D$ |
| GP, fishery landings | $\mathcal{G}$ | GP or landings over $\Delta$ | $w \in W$ or $f \in F$ |
| Harvest | $\mathcal{H}$ | harvest over $\Delta$ | $m \in M$ and $g \in G$ |
| Self losses | $\mathcal{S}$ | self losses over $\Delta$ | $w \in W$ or $d \in D$ |
| Losses | $\mathcal{L}$ | self loss & harvest over $\Delta$ | $w \in W$ or $d \in D$ |
| Habitat size, area | $A$ | habitat available | $z \in Z$ |
| Mass | $Q$ | available weight | $n \in N$ |
| Effort | $E$ | fishing effort over $\Delta$ | $f \in F$ |
| Quota | $Y$ | allowed landings | $m \in M$ and $f \in F$ |
| Catch | $\mathcal{C}$ | $(\mathcal{G} + \mathcal{D})_f$ over $\Delta$ | $m \in M$ and $f \in F$ |
| Discards | $\mathcal{D}$ | discards over $\Delta$ | $m \in M$ and $f \in F$ |
| Temperature | $T$ | temperature index $\leq 1$ | H2O |
| Irradiance, light | $I$ | irradiance index $\leq 1$ | H2O |

**Table 3.** Non-negative parameters for LP simulations.

| Name | Symbol | Associated components | Units |
| --- | --- | --- | --- |
| Self loss rate | $s < 2$ | $W \cup D$ | $\Delta^{-1}$ |
| Self growth rate | $r$ | $A$ | $\Delta^{-1}$ |
| Harvest rate | $\dot{h}$ | $M$ and $G$ | $B_h^{-1} \Delta^{-1}$ or $E_f^{-1} \Delta^{-1}$ |
| Assimilation efficiency | $a < 1$ | $M$ and $H$ | 1 |
| Retention efficiency | $a < 1$ | $M$ and $F$ | 1 |
| Conversion rate | $g = a\dot{h}$ | $M$ and $G$ | $B_h^{-1} \Delta^{-1}$ or $E_f^{-1} \Delta^{-1}$ |
| Satiated growth rate | $g^{\max}$ | $H$ | $\Delta^{-1}$ |
| Satiated harvest rate | $\dot{h}^{\max}$ | $H$ | $\Delta^{-1}$ |
| Particulate self loss | $p \leq 1$ | $W$; ($p_d = 0$) | 1 |
| Concentration (mass) | $c$ | $N$ in $W$ or $D$ | $Q_{n,w} B_w^{-1}$ or $Q_{n,d} B_d^{-1}$ |
| Occupation rate | $v$ | $W$ and $Z$ | $A_z B_w^{-1}$ |

**Table 4.**
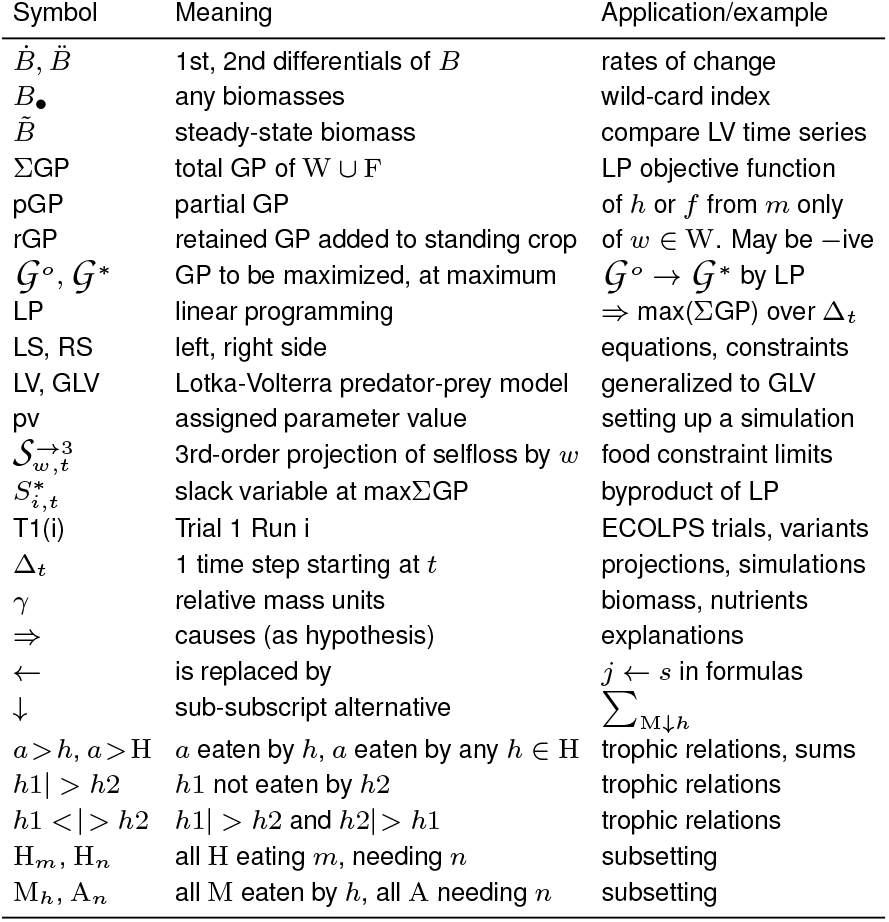
Special symbols and abbreviations.

Subscripting of terms may be omitted when obvious. For example, harvesting rate of *m* by *h*, **𝒽**_*m,h*_*B*_*m,t*_*B*_*h,t*_, becomes **𝒽***B*_*m*_*B*_*h*_. Foods *always* precede consumers in such terms and subscripts. Summation ranges, for example over heterotrophs *h* ∈ H, are ∑ _H_ or, over all foods of *h*, are ∑ _M>h_ where *>* between two component sets signifies ‘harvested by’. Matrices of trophic terms have foods in rows and consumers in columns, unless transposed, ^*T*^ . Note that ‘foods’ are defined as harvested *and* assimilated; totally unassimilated harvesting is briefly discussed in Annex S3. Vectors are bold, l.c.; for example **g** = [**𝒢**_1_, **𝒢**_2_, … ]^*T*^ where **𝒢** is notation for GP. Numerical *projections* (21), shown as superscript ^→*x*^, refer to model-dependent calculation of conditions at *t* + Δ from those at *t* using an *x*-order Taylor series, see Annex S1; for example a 3^rd^-order projection of GP of *w* over Δ_*t*_ starting from 0 is 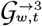.

GP or **𝒢**_*w,t*_ is defined as the notional total quantity of organic biomass formed over Δ_*t*_ by anabolic metabolism of *w*, either autotrophically by photosynthesis or heterotrophically by harvesting and conversion of organic matter (‘secondary production’). **𝒢**_*w,t*_ is distributed continuously via three GP pathways, Fig.1:-

i. Self-lost GP:- (a) *Metabolized* for respiration or release to water as excretions (including secretions), or reproductive materials deemed to belong to another component = *w*; or (b) *Particulate* including molted exoskeletons, body parts, and corpses not resulting from harvesting. The *particulate self-loss* proportion, 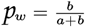.
ii. Harvested GP:- (a) *Assimilated* by heterotrophs eating *w* to fuel their own GPs; or (b) *Dropped* including feces holding indigestible parts of *w*, and body parts of *w* regurgitated or discarded by the harvester. The *assimilation* (also ‘ecological’ (18)) *efficiency*, Table 3, is 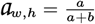.
iii. Retained GP (rGP):- GP added to standing crop *B*_*w,t*_ pending later self-loss or harvesting. rGP may be negative.

*Partial* GP or pGP, notated as **𝒢**_*m,h,t*_, refers to GP of *h* specifically from food *m* over Δ . So ∑_M*>h*_ **𝒢**_*m,h*_ = **𝒢** _*h*_ . For partial harvests, ∑_M*>h*_ **ℋ**_*m,h*_ = **ℋ**_*h*_; likewise ∑_*m>*H_ **ℋ**_*m,h*_ = **ℋ**_*m*._

For brevity, harvesting rate of *m* by *h*, written in full as **𝒽**_*m,h*_*B*_*m,t*_*B*_*h,t*_, becomes **𝒽***B*_*m*_*B*_*h*_. Foods always precede consumers in such terms and subscripts. Summation ranges, for example over heterotrophs *h* ∈ H, are ∑_H_. Matrices of trophic terms have foods in rows and consumers in columns, unless transposed, ^*T*^ . So, positive values in the *w*’th row of harvest-coefficient matrix [**𝒽**] identify the consumers *g* ∈G_*w*_ eating *w*. Positive values in the *g*’th column of assimilation-coefficient matrix [**𝒶**] identify the foods *m* ∈ M_*g*_ eaten by *g*. Note that ‘foods’ are defined as harvested *and* assimilated; unassimilated harvesting is not discussed. Vectors are bold, l.c.; for example **g** = [**𝒢**_1_, **𝒢**_2_, … ]^*T*^ . Numerical *projections* (21), shown as superscript ^→*x*^, refer to model-dependent calculation of conditions at *t* + Δ from those at *t* using an *x*-order Taylor series, see Annex S1; for example a 3^rd^-order projection of GP of *w* over Δ_*t*_ is 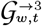.

### Simulating an aquatic ecosystem by maximizing ΣGP within constraints

E.Odum wrote ‘Nature maximizes for gross production’ (22, p. 48). Lotka wrote similarly of energy flux through an ecosystem (23). In support of these views, GP provides all the energy and biomass required by an organism for activity, harvesting, growth, reproduction, and self repair, implying that every wild must strive to maximize GP for best selective advantage within any constraints acting on it.

Simulation of unfished aquatic ecosystems by maximization of ΣGP subject to constraints on GPs of individual wilds, accords with this idea. Furthermore, ΣGP incorporates the productive outputs of heterotrophs and their foods because, by the definition of GP above, when any *h* consumes any *m*, the harvested *B*_*m*_ contributes to **𝒢**_*h*_ but is not subtracted from **𝒢**_*m*_. So **𝒢**_*m*_ + **𝒢**_*h*_, increases monotonically over time even though *B*_*m*_ + *B*_*h*_ may increase or decrease.

Fisheries *f* ∈ F can be incorporated as consumers *g* ∈ G along-side heterotrophs in the ΣGP objective function by choosing a biomass variable that fishers strive to maximize. Marketable landings **𝒢**_*f*_ is proposed because of their motivating financial value. Catches **ℋ**_*f*_, on the other hand, are better for measuring harvests because they include discarded biomass (24, 25) such as unsaleable species, size groups and offal.

### Linear programming

The LP task is to maximize ΣGP over each Δ_*t*_:-

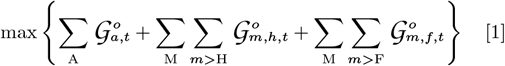

subject to all 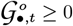, and to **𝓃**_**i**_ other linear constraints,

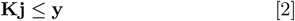

Where

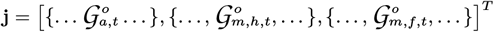

is the column vector of the **𝓃** _j_ = **𝓃**_A_ + ∑_M_ **𝓃**_m< G_ quantities on the RS of Eq. (1), **K** is an **𝓃**_**i**_ *×***𝓃**_**j**_ matrix of coefficients, and **y** is an **𝓃**_**i**_ vector of limiting values. Signs of coefficients in **K** are adjusted if necessary so that Relation (2) is always ≤; this simplifies LP computations (15). All 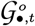 in Eq. 1 receive unit weighting so that they are all assigned equal ecological significance, given that their different abilities to harvest and grow are accounted for in constraint formulas.

Slack variables, 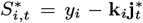 for *i* = 1, …, **𝓃**_**i**_ measure the unutilized capacity under the *i*th constraint at max ΣGP, denoted ^∗^. 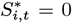 identifies an ‘active’ constraint, one of the intersecting group of **𝓃**_active_ ≥ **𝓃**_**j**_ constraints binding the maximum; (**𝓃**_active −_ **𝓃**_**j**_) redundant constraints may occur at the LP solution (14). Slack variables from food constraints are used to update biomasses after each Δ_*t*_; see Eq. (23).

### Mass models

#### The Lotka-Volterra predator-prey model

The LV model for numbers of one predator and one prey (26, 27) is here applied to organic biomasses, *B*. They better measure the nutritive value of food and are consistent with GP as maximized in Eq. (1). Additionally, during reproduction, *B*_*w*_ gradually increases as gametes or embryos mature internally, then are released without further immediate organic gains, whereas abundances of many aquatic species spike sharply upwards when large numbers of very small fertilized embryos are released to water (28). Thus biomass also offers better continuity of variation and avoids dependence on imprecise stock-recruitment models (29). Retained GP (rGP) rates at *t* for autotroph *a* and heterotroph *h, a>h*, are

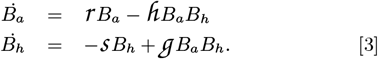

Coefficients are listed in Table 3. Conversion is proportional to harvest:

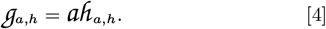

Eq. (3) may be set to zero and solved to find:

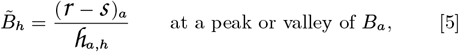

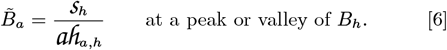

These are also *steady-state* (30), or ‘equilibrium’ (31) values reached if oscillations are allowed time to die away. Additionally, oscillatory frequency *ϕ* of an LV system is ‘governed’ (30, p. 220) by

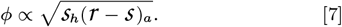

Three uses for the 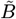and *ϕ* are:

1. Simulated LV steady-state values may be verified with formulas for 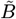 and *ϕ*.
2. Values of parameters may be chosen to achieve relative steady states in an LV simulation. For example, set

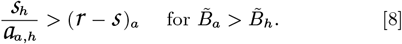
3. Since time cancels when Eq. (5) or (6) is solved for units from Table 3, but Eq. (7) has units Δ^−1^, simulated cycles may be slowed or hastened without altering steady-state biomasses by multiplying all time-dependent parameters by a constant. For example, halving the values, and thus doubling the time unit, of **𝓈, 𝓇**, **𝒽** should also halve the frequency – or double the wavelength – of LV oscillations.

#### A generalized Lotka-Volterra model (GLV)

The LV model can be generalized to food webs and fisheries by considering that wilds *w* ∈ W in an unfished ecosystem may each be a food for *h* ∈ H_*w*_ and, if themselves heterotrophic, a consumer eating foods *m* ∈ M_*w*_ . Rewriting Eq. (3) for these two roles, *i* and *j*, of *w*:-

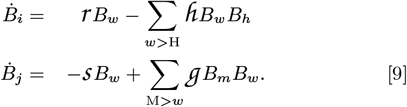

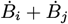 gives the rGP rate for *w*:-

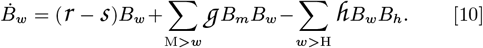

Self-growth coefficient, **𝓇**_*w*_, is the maximum rate of photosynthesis per unit of *B*_*w*_ if *w* ∈ A, or 0 otherwise; **𝒽** _*w,h*_ is the maximum rate of harvesting of *w* per unit of *B*_*h*_; and **ℊ**_*m,w*_ is the maximum rate of conversion of *m* per unit of *B*_*w*_ . **𝓈**_*w*_ is the rate of self-loss taken as a maximum during starvation assuming that extreme conditions such as anoxia or toxic pollution are not present. **𝓈**_*w*_ is assumed constant whether *w* is feeding or not. See Table 3 for ranges and units of parameters. The proposed GLV model consists of **𝓃**_W_ Eq. (10). These might be solvable simultaneously for *B* given parameter values (pvs) but results may not then be ecologically feasible, for example if some *B* are negative. The three RS terms of Eq. (10) are the *self-change*(=*self-gain*−*self-loss*), *conversion* (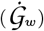), and *harvesting* (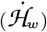 ) rates. Taking self and harvesting losses to the LS to join rGP, and matching the three LS terms with the three GP pathways defined earlier, maximum GP rates are seen to be

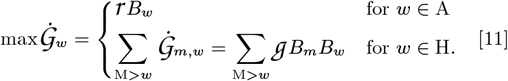

Multiplying Eq. (4) by *B*_*m*_*B*_*h*_ shows that the harvesting rate corresponding to a pGP rate of heterotroph *h* is

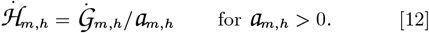

Subtracting fishery catch rates 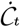 based on fishing effort *E*_*f,t*_ from Eq. (10) gives

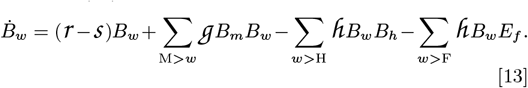

*E* can be measured in various ways (32), for example as trawls or hooks in the water, or length of fixed nets set, here assumed constant over each Δ_*t*_. Conversion of catch to landing rates uses *retention efficiency*, **𝒶**_*m,f*_, analogous to **𝒶**_*m,h*_ applied to wild harvests:-

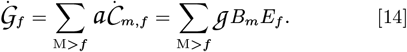

#### Non-living organic biomass model

Four *NLO pathways* affect simulated NLO mass, *B*_*d*_, Fig. 2:-

i. Gains of particulates self lost by W, their GP pathway (i)b in Fig. 1.

**Fig. 1.**
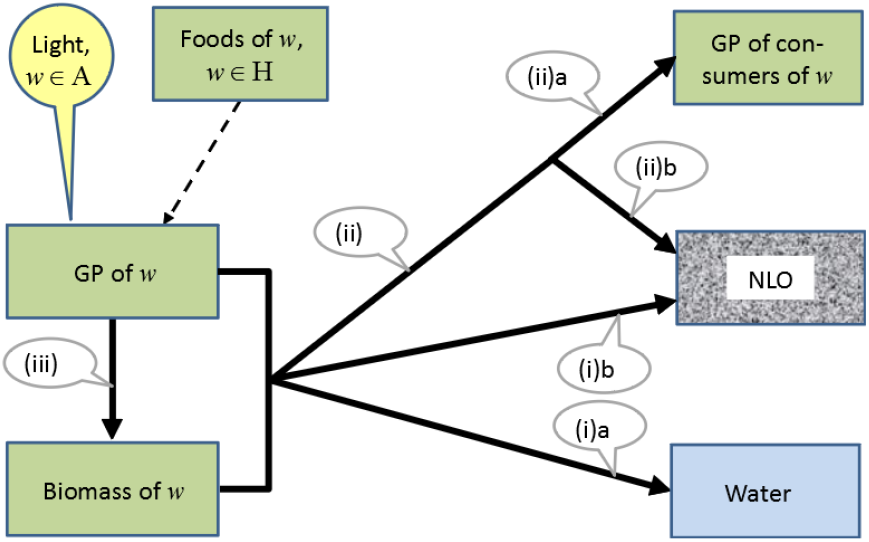
GP pathways identified for simulating wild component *w* in a closed ecosystem, numbered as in the text. Broken arrows=GP pathways of other wild components; NLO=non-living organics; GP=gross production; A=autotrophs; H=heterotrophs.
ii. Gains of particulates dropped or discarded during harvesting of W. See GP pathway (ii)b for W in Fig. 1.
iii. Self losses by biological or chemical decomposition of NLO within *d*. Products transfer to water as solutes only, so **𝒫**_*d*_ = 0.
iv. Losses harvested and assimilated by scavengers S eating NLO. Any dropped NLO returns to *d* and can be ignored; so **𝒶**_*d,s*_ = 1 and **𝒽**_*d,s*_ = **ℊ**_*d,s*_. A GLV rate equation for these pathways is

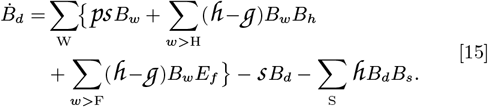

RS terms, in order, model rates for NLO pathways (i), (ii) for H and F, (iii), and (iv).

#### Essential nutrient model

Nutrients *n* ∈ N are essential for autotrophs A_*n*_ ⊆ A, and for heterotrophs H_*n*_ ⊆ H requiring *n* obtained by eating foods A_*n*_ (33) or H_*n*_ also needing, and therefore storing, **𝓃**. Let initial masses of each *n* in water be *Q*_*n*,H2O,*t*=0_, those in NLO be *Q*_*n,d,t*=0_, and let concentrations of *n* in organic tissues of wilds be **𝒸**_*n,w*_ measured as notional averages over all biomass of individuals belonging to *w*. The **𝒸**_*n,w*_ are assumed constant as a result of homeostatic processes; this is a simplification of evidence (34, 35),(36)[§1.5.3 & Box 4.1]. Quantities of *n* in *w* are then **𝒸**_*n,w*_ *B*_*w,t*_. Five *nutrient pathways*, Fig. 3, direct simulated flows of *n*:-

**Fig. 2.**
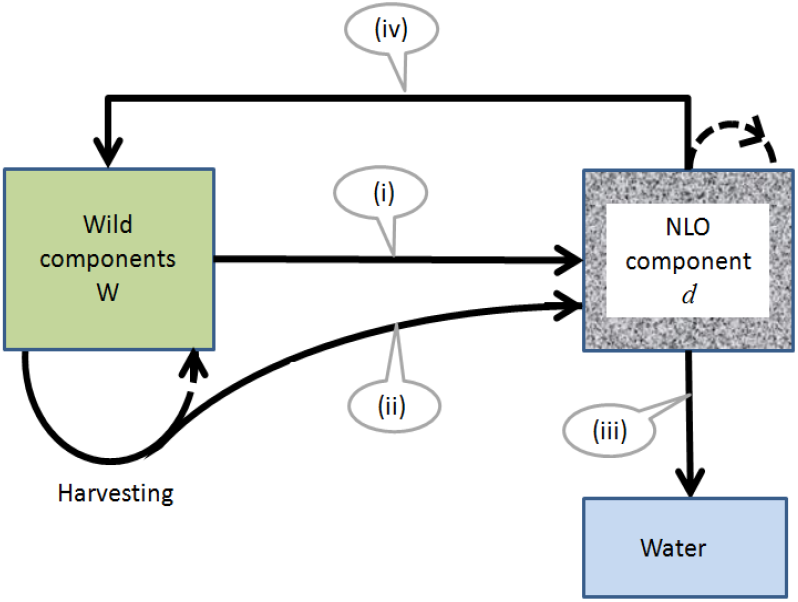
Non-living organic (NLO) pathways identified for simulating a closed ecosystem, numbered as in the text. Broken arrows=harvest returned to source.

**Fig. 3.**
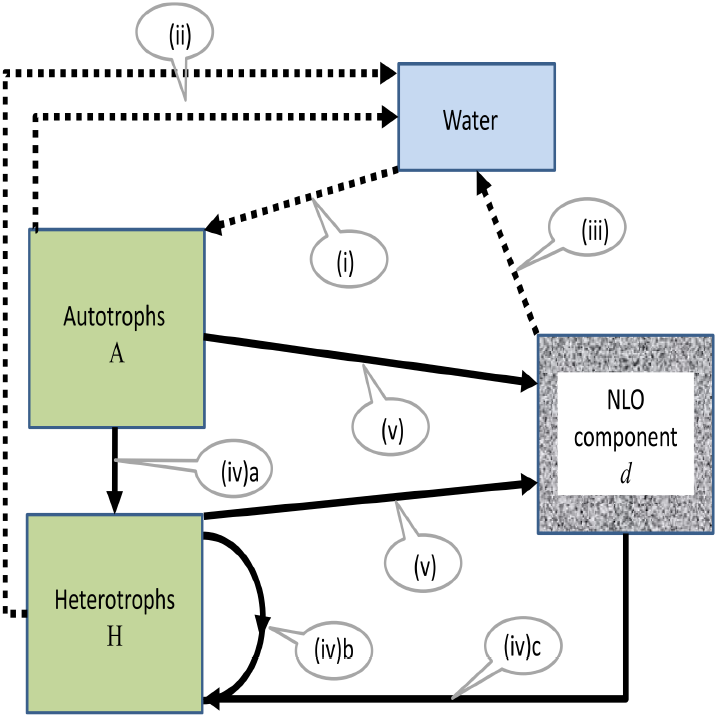
Nutrient pathways identified for simulating components in a closed ecosystem, numbered as in the text. Broken arrows=transport via water; solid arrows=transport via biomass or organic matter. NLO=non-living organic matter.

i) Uptake of *n* from water by A_*n*_ for photosynthesis.

ii) Transfers of dissolved *n* from wilds to water as excretions (including any secretions).

iii) Transfers of dissolved *n* to water self-lost by *d*, NLO pathway (iii) in Fig. 2.

iv) Assimilation of *n* by H_*n*_ from (a) autotrophic, (b) heterotrophic, and (c) NLO foods via GP pathway (ii)a in Fig. 1 and NLO pathway (iv) in Fig. 2.

v) Transfers to *d* of *n* bound to particulate self and harvested losses of W via their GP pathways (i)b and (ii)b in Fig. 1, and NLO pathways (i) and (ii) in Fig. 2.

For autotrophs, let the net rate of transfer of *n* mass to water at *t* via nutrient paths (i) and (ii) be

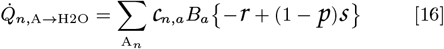

where 1 − **𝒫**_*a*_ is the dissolved proportion of **𝓈**_*a*_. For heterotrophs, let rate of loss of *n* to water via nutrient path (ii) depend on surplus *n* assimilated from foods, nutrient paths (iv). From Eq. (11) with *w* ← *h*, the rate of supply of *n* to *h* from assimilated food *m* is **𝒸**_*n,m*_**ℊ** *B*_*m*_*B*_*h*_, and the rate of need for additional *n* by growing *h* is **𝒸**_*n,h*_**ℊ** *B*_*m*_*B*_*h*_. If **𝒸**_*n,h*_ is to remain constant by excretion, the total rate of transfer of *n* to water by all *h ∈* H is

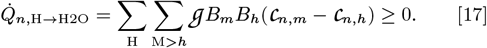

Let the rate of gain of *n* mass in water at *t* caused by efflux of dissolved *n* from NLO *d*, nutrient path (iii), be

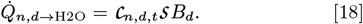

**𝒸**_*n,d,t*_ varies with *t*. 1 − **𝒫**_*d*_ = 1 is omitted; see NLO path (iii). Lastly, the net rate of gain at *t* of *n* mass by NLO from wilds via nutrient paths (iv)c and (v) is, from Eq. (15),

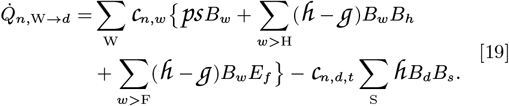

In a *closed*, unfished ecosystem, the total *n* mass *Q*_*n*Σ_ in a simulation that includes recycling of *n* via water, NLO and scavengers S should be constant:

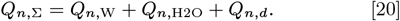

Eq. (16)-(19) all describe rates of lossless transfers of *n* between two RS terms of Eq. (20), and therefore not affecting *Q*_*n*,Σ_. However, in trials including *n* recycling (below), constant *Q*_*n*,Σ_ could not be reliably achieved without trial-and-error adjustment of pvs. Possibly, biases inherent in projected self losses used in several constraint-limit formulas, see Table 5 below, may underlie these unwanted trends in *Q*_*n*,Σ_.

**Table 5.** Simulating aquatic ecosystems:- Proposed linear constraints on GP over Δ_*t*_.

| Name [code] | Const-<br>trained | Processes<br>affected | Left side | Limit | N cons-<br>traints | Slack measures:- |
| --- | --- | --- | --- | --- | --- | --- |
| Autotrophic GP<br>[GP.a] | $a \in A$ | photo-<br>synthesis | $\mathcal{G}_{a,t}^o$ | $\leq \mathcal{G}_{a,t}^{\rightarrow 3}$ | $n_A$ | lost GP<br>of $a$ |
| Consumer pGP<br>[m.pGP.g] | $g \in G$ | growth from<br>$m_g$ | $\mathcal{G}_{m,g,t}^o$ | $\leq \mathcal{G}_{m,g,t}^{\rightarrow 3}$ | $\sum_G n_{M_g}$ | lost pGP of<br>$g$ from $m_g$ |
| Living food<br>[m.FOOD.G] | $g \in G_{M \cap W}$ | harvest of<br>$m \in M$ | $\sum_{m > G} \frac{\mathcal{G}_{m,g,t}^o}{a_{m,g}} - \mathcal{G}_{m,t}^o$ | $\leq B_{m,t} - \mathcal{S}_{m,t}^{\rightarrow 3}$ | $n_M$ | $B_{m,t+\Delta}$ for<br>$m \in M$ |
| Dummy food<br>[w.FOOD] | None | None | $-\mathcal{G}_{w,t}^o$ | $\leq B_{w,t} - \mathcal{S}_{w,t}^{\rightarrow 3}$ | $n_{W \notin M}$ | $B_{w,t+\Delta}$ for<br>$w \notin M$ |
| Satiated GP<br>[M.SATE.h] | $h \in H$ | growth<br>when $M_h$<br>in<br>excess | $\sum_{M > h} \mathcal{G}_{m,h,t}^o$ | $\leq \mathcal{G}_h^{\max} B_{h,t}$ | $\leq n_H$ | lost GP of $h$ from<br>$M_h$ |
| Heterotrophic har-<br>vest [M.HARV.h] | $h \in H$ | harvest<br>when $M_h$<br>in<br>excess | $\sum_{M > h} \frac{\mathcal{G}_{m,h,t}^o}{a_{m,h}}$ | $\leq \mathcal{H}_h^{\max} B_{h,t}$ | $\leq n_H$ | lost harvest of<br>$M_h$ by $h$ |
| Habitat [z.HAB.W] | $w \in W_z$ | rGP of<br>$w$ need-<br>ing $z$ | $\sum_{W_z} v_{z,w} \left\{ \mathcal{G}_{w,t}^o - \sum_{w > G} \frac{\mathcal{G}_{w,g,t}^o}{a_{w,g}} \right\}$ | $\leq A_{z,t} + \sum_{W_z} v_{z,w} \mathcal{S}_{w,t}^{\rightarrow 3}$ | $n_Z$ | $A_{z,t+\Delta}$ |
| Autotrophic nutri-<br>ent [n.NUTR.A] | $A_n$ | GP of $a$<br>needing<br>$n \in N$ | $\sum_{A_n} c_{n,a} \mathcal{G}_{a,t}^o - \sum_H \sum_{M > h} \mathcal{G}_{m,h,t}^o (c_{n,m} - c_{n,h})$ | $\leq Q_{n,H_2O,t} + \sum_{i \in A \cup d} c_{n,i} (1-p) \mathcal{S}_{i,t}^{\rightarrow 3}$ | $n_N$ | $Q_{n,H_2O,t+\Delta}$ |
| Excreted nutrient<br>[n.EXCR.h] | $h \in H_{n \in N}$ | growth of H<br>needing $n$ | $-\sum_{M > h} \mathcal{G}_{m,h,t}^o (c_{n,m} - c_{n,h})$ | $\leq 0$ | $\sum_N n_{H_n}$ | excretion of $n$ by<br>$h$ |
| Non-living food<br>[d.FOOD.S] | $s \in S_d$ | harvest of<br>$d \in D$ | $\sum_{d > S} \mathcal{G}_{d,s,t}^o - \sum_M \sum_{m > G} (a_{m,g}^{-1} - 1) \mathcal{G}_{m,g,t}^o$ | $\leq B_{d,t} - \mathcal{S}_{d,t}^{\rightarrow 3} + \sum_W p_w \mathcal{S}_{w,t}^{\rightarrow 3}$ | $n_d$ | $B_{d,t+\Delta}$ |

#### Essential habitats

One or more wilds *w* ∈ W_*z*_ may compete to live in essential habitat *z* ∈ Z whose total ‘size’ *A*_*z*_ is fixed. *z* might be a nursery area or feeding ground, and *A*_*z*_ might be an area of sea floor or a limited number of habitable crevices present. Then *z* is a potentially constraining resource and *w* ∈ W_*z*_ may live nowhere other than *z* if *z* is truly *essential*. Any *w* ∉W taking space in *z* when it need not is ignored here. Let the occupation coefficient **𝓋**_*z,w*_ be the space in *z* needed per unit biomass of *w* ∈ W_*z*_ . Then the space left in *z* for filling by rGP of *w* at *t* is:

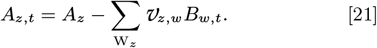

### Constraints

#### Preliminaries

Constraints may be active directly on GP, or indirectly via rGP. Proposed linear constraints over Δ_*t*_ are listed in Table 5. Constraints are named, for example ‘m.FOOD.G’ where ‘m’, in this case an individual component, is the constraining cause if clear, otherwise omitted. ‘FOOD’ (u.c.) is the type of constraint, and ‘G’, in this case is a constrained set of components. Pvs and fishing effort *E*_*f,t*_ are constant within Δ_*t*_ but may be varied prior to Δ_*t*+1_; see section below.

The LS of each constraint is a linear function of **𝒢** ^*o*^ terms in the objective function, Eq. (1). The **𝒢**^∗^, that is, the **𝒢** ^*o*^ after maximization by LP, serve to numerically integrate the max differentials in Eq. (11) over Δ_*t*_ up to the amount permitted by active constraints. Assimilated harvests, if relevant to the constraint, appear on the LS as **𝒢** ^*o*^*/***𝒶** if **𝒶** *>* 0; this term integrates the differentials in Eq. (12).

The RS of constraints are upper limits. For rate constraints, GP.a and m.pGP.g, the RS is a projected integral of the appropriate maximum GP rate over Δ_*t*_ approximated with a 3^rd^-order Taylor series for integrals; see Annex S1. This numerical method (37) is compatible with the autonomous structure (31) of GLV equations. Starting from **𝒢**_*w,t*_ = 0,

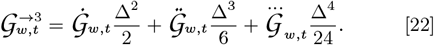

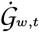 is from Eq. (11) with *w* ← *a* or *h*, and 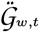 and 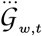 are from Annex S1 Eqs. (S7) and (S9). Since GP.a and m.pGP.g set maximum GP rates, they are only active when all more restrictive constraints are inactive, implying that Eq. (22) is safely valid when it is needed. Negative 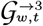 can occur because of poor approximation by Eq. (22), for example when the true value dips sharply to almost zero but the approximation overshoots. In this study, all negative values were replaced with zeros, or were avoided by adjusting selected pvs.

Self losses integrated over Δ_*t*_, **𝒮**_*w,t*_, are required for the RS of FOOD, HAB and NUTR constraints, Table 5. 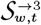 can be calculated from . (22) with **𝒢** ← **𝒮** . Referring to Annex S1, 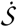, 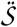 and 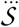 are in the wild diagonals of 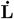,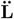 and 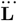 respectively. Self losses are unconstrained. However, if biomasses are increasing from *B*_*t*_ over Δ_*t*_, 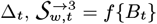 may under-estimate **𝒮**_*w,t*_ and vice-versa, with consequent effects on the RS limits. These are called ‘self-loss errors’ below.

#### Constraint formulas

These notes supplement Table 5. GP.a, m.pGP.g, m.FOOD.G, and FOOD.g are *minimal* constraints because all are required for every simulation. The **K**_*•,•*_ below, are elements of coefficient matrix **K** in Eq. (2). The first subscript identifies the row *within the subset* holding the constraint type, and the second identifies the column. Unspecified elements are zero.

##### Autotrophic GP (GP.a)

The maximum GP of *a* is from Eq. (11), (22). **K**_*a,a*_ = 1 for *a* ∈ A.

##### Consumer pGP (m.pGP.g)

The maximum pGP of *m>g* is from Eq. (11), (22). **K**_*m,m>g*_ = 1 for all *m* M and *g* G.

##### Living-food (m.FOOD.G)

LS: Harvests of *m* from Eq. (12), integrated for each consumer as 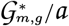, are replenished by GP of *m* itself. RS: The *B*_*m,t*_ are reduced by projected self losses. See Eq. (10) or (13) with *m* ← *m*^*′*^, *w* ← *m*. 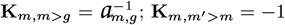 for all *m ∈* M, all *m ∈* M_*m*_ and all *g* ∈ G.

##### Dummy-food (w.FOOD)

‘Dummy’ food constraints are applied to unharvested foods, for example unfished top predators, to supply slack variables for Eq. (23) below. **K**_*w,m>w*_ = − 1 for all *w* ∉M, *m* ∈ M_*w*_ .

##### Satiated GP (M.SATE.h)

Satiated GP of *h* summed over all of its foods *m* ∈ M is limited by a maximum assumed proportional to *B*_*h,t*_ with coefficient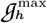. cf. **ℊ**_*m,h*_, Table 3. M.SATE.h is motivated by Holling type I or II responses of predators (10, 38). **K**_*h,m>h*_ = 1 for 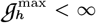,*h* ∈ H, and all *m* ∈ M_*h*_.

##### Heterotrophic harvest (M.HARV.h)

Harvesting in the presence of excessive food may be limited similarly with 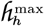 given independent growth and harvesting rates. 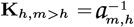 for 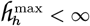,*h* ∈ H, and all *m* ∈ M_*h*_ .

##### Essential habitat

**(z.HAB.W)** is required if habitat *z* Z of finite size *A*_*z*_ has any **𝓋**_*z,w*_ *>* 0. Occupancy by rGP of *w* W_*z*_ during Δ_*t*_ must be to size currently available, *A*_*z,t*_, Eq. (21). LS: Use of *z* by GP of *w* ∈ W_*z*_ is corrected for size released by harvests of *w*. RS: *A*_*z,t*_ is increased for size released by self losses of *w*. For **K** with row *z* holding z.HAB.W: 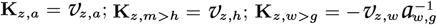

##### Autotrophic nutrient

**(n.NUTR.A)** is required if any **𝒸** _*n,a*_ >0. Summed photosynthetic removal of *n* from water by *a* must be ≤ the amount of *n* available over Δ_*t*_. LS Term 1: Removal of *n* by all A_*n*_ is summed; see RS term 1 of Eq. (16). LS Term 2: *n* excreted into the water by *h* ∈ H is subtracted; see Eq. (17). RS Term 2: Dissolved self-losses from all *a* ∈ A and *d* ∈ D are added; see Eq. (16), (18). For **K** with row *n* holding n.NUTR.A: **K**_*n,a*_ = **𝒸**_*n,a*_; **K**_*n,m>h*_ = − **𝒸**_*n,m*_ + **𝒸**_*n,h*_.

##### Excreted nutrient

**(n.EXCR.h)** is required for any *h* with homeostatic **𝒸**_*n,h*_ *>* 0. Relationship (17) for individual *h* is here negative. For **K** with row *i* (for each {*n, h*} pair) holding n.EXCR.h: **K**_*i,m >h*_ = **𝒸**_*n,h*_ − **𝒸**_*n,m*_ .

##### Non-living-food

**(d.FOOD.S)** is required if NLO component *d* and scavengers *s* ∈ S are present with **ℊ**_*d,s*_ *>* 0. Harvests of *d* by S must be ≤ NLO mass available over Δ_*t*_. d.FOOD.S uses integrated rate terms from Eq. (15). LS: Term 1: Harvests of *d* by S are summed; note that harvest **ℋ**_*d,s*_ = **𝒢**_*d,s*_ because the GP-to-harvest factor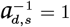 . Term 2: Replenishing gains of NLO from all dropped harvests of wild foods, calculated as (harvest - GP)= (**𝒶**^−1^ 1)**𝒢** ^*o*^, are subtracted. RS: Term 2: *d*’s dissolved self losses to water are subtracted from *B*_*d,t*_. Term 3: All dropped particulate wild self losses are added. For **K** with *d*.FOOD.S coefficients in row *i*: **K**_*i,d>s*_ = 1, and 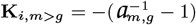 for all *m ∈* M and *g ∈* G_*m*_.

##### Updating biomasses

Having found maximum ΣGP over Δ_*t*_ by LP, wild biomasses needed to re-calculate constraint limits and run the LP projection over Δ_*t*+Δ_ are obtained from the slack variables of FOOD constraints:

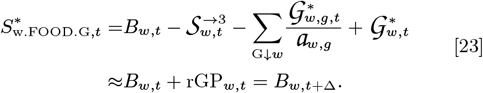

This is an approximation because of the possible self-loss error. Mass of NLO, essential *n* in water, and habitat *A*_*z*_ available at Δ_*t*+Δ_ are found from slack variables 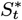see Table 5, last column.

The concentration of *n* in NLO *d* at *t* + Δ, **𝒸**_*n,d,t*+Δ_ is needed for the RS of n.NUTR.A, Table 5. The mass of *n* in NLO at *t* + Δ is obtained from Eq. (18), (19) integrated over Δ_*t*_ either with third-order projections or, where possible, **𝒢**^∗^ from LP for better accuracy:-

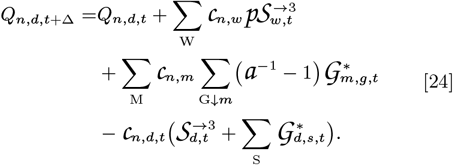

The new concentration of **𝓃** in *d* is

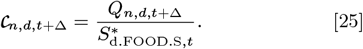

##### Timed adjustments of simulations

Supply of a days *×* updates ‘Envir.file’ causes ECOLPS to search there after each Δ_*t*_ for changed environmental conditions applicable over Δ_*t*+1_; see Annex S3. For example, autotrophic self-growth rate can be modeled as a seasonal function of light index 0 ≤ *I*_*t*_ ≤ 1 (darkest to brightest) with values in rows of Envir.file varying from dim in winter to bright in summer. A temperature index 0 ≤ *l*_*T*_ ≤ *T*_*t*_ ≤ *h*_*T*_ ≤ 1 (coolest to warmest) may also be included to prevent light-dependent self growth in unfavorable temperatures. A light-saturation curve is available (36, 39) but a simpler proportional system was found easier to apply. With parameter *α*_*I*_ *>* 0,

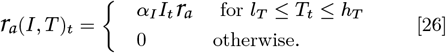

Similarly, whole columns in [**𝒽**] may be adjusted uniformly with temperature using *α*_*T*_ *T*_*t*_**𝒽**_*•,h*_.

### Simulations

The ECOLPS simulator, described in Annex S3, is available with all Trial.txt files from [URL to be determined]. Readers may use these to re-run, extend or vary trials. LV parameters were set arbitrarily and relatively, sometimes using methods in Annex S2. Annex S4 archives printed pvs and Controls.txt settings for every trial. 1000Δs are displayed for most non-seasonal trials, 8 years of 12Δs for seasonal trials with 1Δ = 1 ‘month’. Additional unillustrated runs were made to check stability. Abbreviations: ‘T1(ii)’ = Trial 1 Run (ii); ‘⇒’ means ‘causes, as a hypothesis,’.

#### Trial 1. The minimal model with alga *a*, **grazer** *h, a > h*

**Aim:** To simulate basic LV theory. **Method:** Pvs are in Table 6; Fig. 4 shows the configuration of constraints, objective function and feasible region at *t* = 0. **ℊ** ^max^ was set high so that a.SATE.h was never active in T1. **Runs:** (i) Starting biomasses arbitrarily displaced from steady-state values. (ii) Starting biomasses set exactly at steady-states. (iii) Run(i) repeated with values of time-dependent parameters **𝓈, 𝓇** and **𝒽** halved. (iv) Run(i) repeated with **𝓇**_*a*_ set at 7 different values.

**Table 6.**
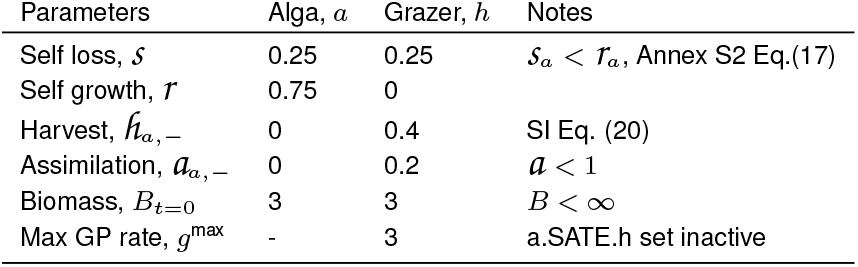
LV trials 1 to 7. Values set for parameters; adjustments in text.

**Fig. 4.**
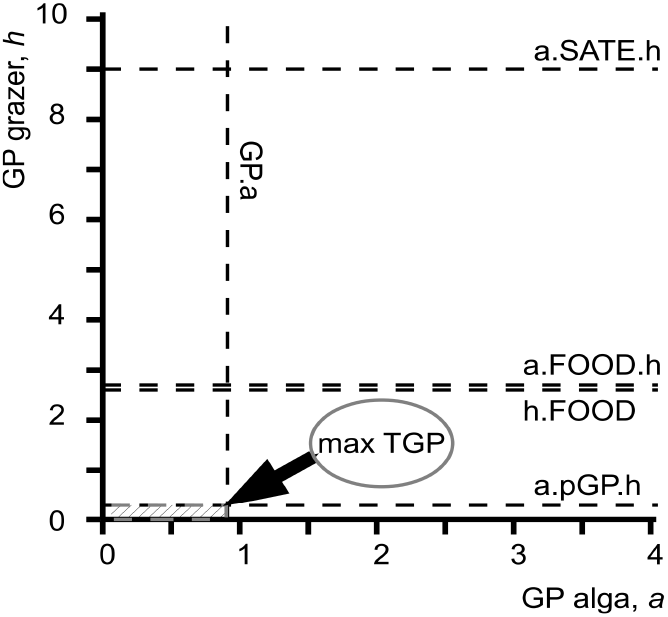
Trial 1, run 1. First LP projection:- configuration of constraint limits, objective function (diagonal hatching over the feasible region), and max ΣGP=‘max TGP’.

##### Results. Run(i)

Fig. 5(top). Only GP.a and a.pGP.h were active throughout the simulation. Biomass time series showed oscillations damped towards steady states, 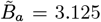and 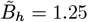, in conformity with Eq. (5), (6). Every oscillation traversed the 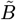 values at peaks and valleys of the other biomass variable, as expected, from near the start of the series, see Table S3, indicating stability of the system throughout. Wavelengths for both series were 36Δs. Damped cycles contrasted with irregular but undamped cycles reported for lynx (40) and their prey, snowshoe hare (41). **Analysis**: Damping can be explained as follows. Initial biomasses, Table 6, were arbitrarily displaced from steady states: 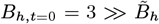, and 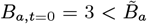. High, displaced *B*_*h,t*=0_ ⇒ fast initial harvesting of *B*_*a*_ ⇒ an extremely low first valley of *B*_*a*_ ⇒ *B*_*h*_ declined to an extremely low level by starvation ⇒ fast recovery of *B*_*a*_ while *B*_*h*_ was low fast increase of *B*_*h*_ in response ⇒ *B*_*a*_ peaked when self and harvesting losses exceeded self growth, then declined rapidly because of high *B*_*h*_ ⇒ *B*_*a*_ returned to its starting value, *B*_*a,t*=0_ = 3, when *B*_*h*_, although at a peak, was slightly less than its starting value, *B*_*h,t*=0_ = 3 ⇒ the second dip of *B*_*a*_ was not as low as the first because of lower grazing, and the second dip of *B*_*h*_ was slightly higher than the first because of higher food presence per unit weight of *h*, and so on.

**Fig. 5.**
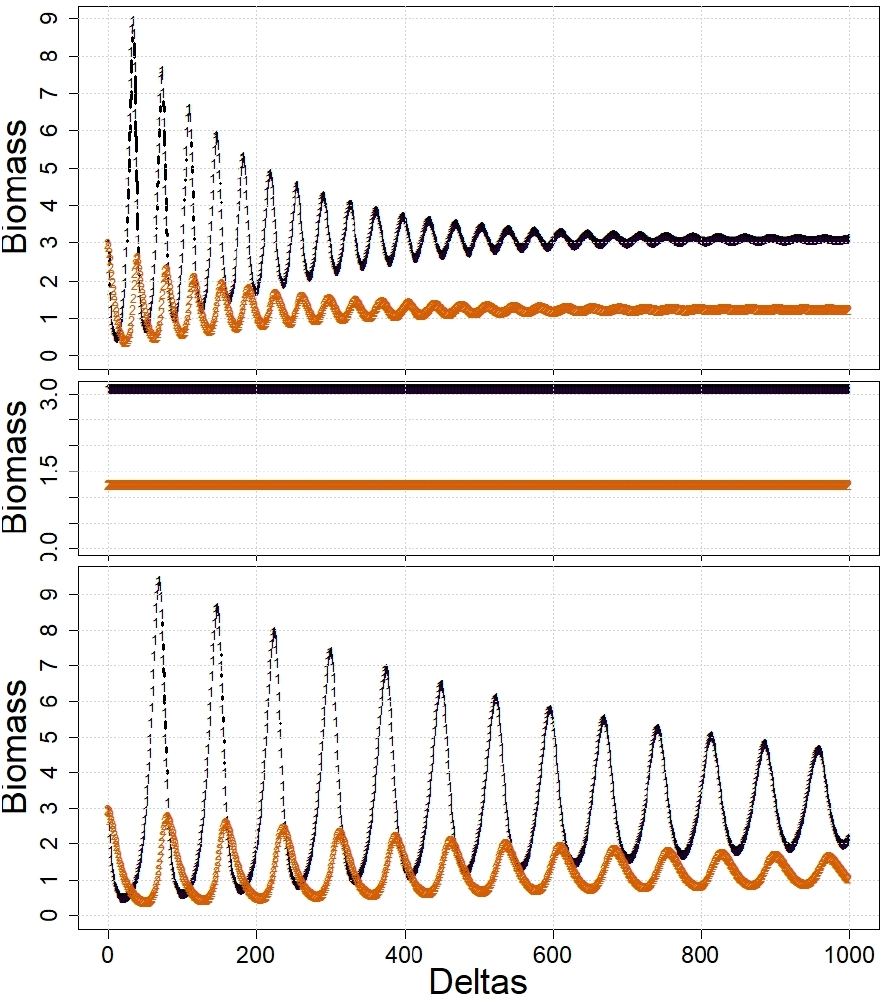
Trial 1. LV system with alga *a* and grazer *h*. TOP, Run(i): parameter values as in table 5. MID, Run(ii): Initial biomasses re-set to steady-state values, *B*_*a*,0_ = 3.125, *B*_*h*,0_ = 1.25. BOTTOM, Run(iii): Parameter values as in Table 5 except **𝓈**_*a*_ = **𝓈**_*h*_ = 0.125, **𝓇**_*a*_ = 0.375, **𝒽**_*a,h*_ = 0.2, 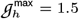. Key: black 1s=*a*, orange 2s=*h*.

##### Run(ii)

Fig. 5(mid). Starting with 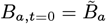 and 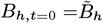, no oscillations occurred.

##### Run(iii)

Fig. 5(bottom). Halved values of time-dependent parameters slowed attainment of steady states. After 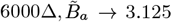 and 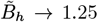 as expected. Wavelengths for peaks of *a* and *h* both gradually declined to 71Δ, 1.4% less than the expected doubling of that found in Run(i); see **LV model** note (iii).

##### Run(iv)

Fig. 6. Simulations with varying **𝓇**_*a*_ found, as steady states were approached between 1000 and 5000Δ (depending on **𝓇**_*a*_), that the 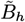 were close to, and the frequencies at the ends of series were proportional to theoretical values in Eq. (5), (7), respectively. Proportionality of frequencies is expected from theory based on eigenvalues (30, p. 219-20).

**Fig. 6.**
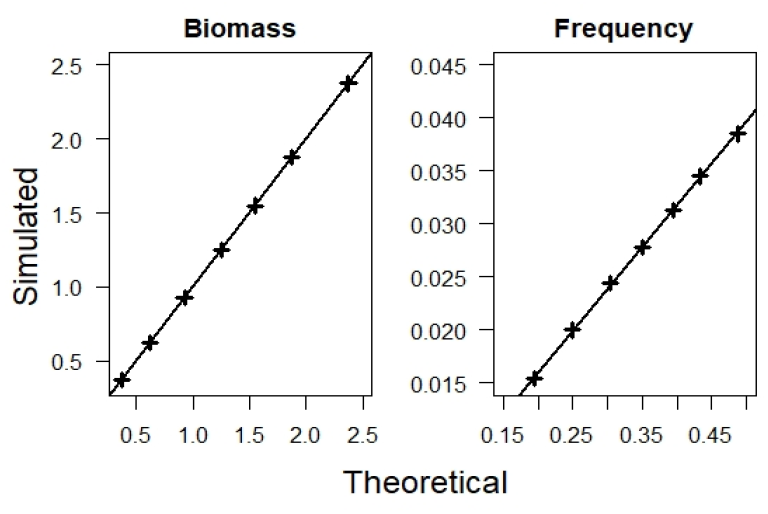
Trial 1, Run(iv). Comparisons of simulated values at steady states with theory at 7 points when **𝓇**_*a*_ −{ 0.4, 0.5, 0.62, 0.75, 0.87, 1, 1.2} ; 1000 to 5000Δs; other parameters in table 5. LEFT:- 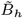, theory from Eq. (5). RIGHT:- oscillatory frequencies *ϕ*(*B*_*h*_) ≈ *ϕ*(*B*_*a*_), theory from Eq. (7).

##### Conclusions

Damped oscillations occur in minimal LV systems when initial displacements, 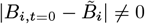 . *B*_*a,t*_ and *B*_*h,t*_ then progress with reducing amplitudes towards steady states. Their values, and the LV frequencies found by ECOLPS were consistent with theory, including when one parameter, **𝓇**_*a*_, was varied over a wide range, or when the time unit was doubled.

#### Trial 2. Grazer satiation in LV cycles

**Aim:** To simulate grazer satiation in a minimal LV system. **Method:** T1(i) was repeated with low values of 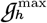 to activate a.SATE.h. **Runs:** (i) 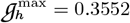. (ii) 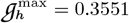.

##### Results

Fig. 7. GP.a, and either a.pGP.h or a.SATE.h were active in both runs. **Run(i)** Marginal activity of a.SATE.h was found; times are marked as green 3s in Fig. 7. Initial cycles of *B*_*a*_ and *B*_*h*_ peaked slightly higher than in T1(i) and with constant amplitudes. Peaks of *B*_*a*_ occurred while a.SATE.h was active, those of *B*_*h*_ 4 or 5Δ afterwards. Damping then led, at 3000Δ, to steady-state and wavelength values as in T1(i). **Run(ii)** Similar marginal activity of a.SATE.h occurred initially but followed by amplification, not damping, of cycles. *B*_*a*_ grew unrestrained by *h* until 357Δ when *B*_*h*_ collapsed, and run (ii) terminated prematurely.

**Fig. 7.**
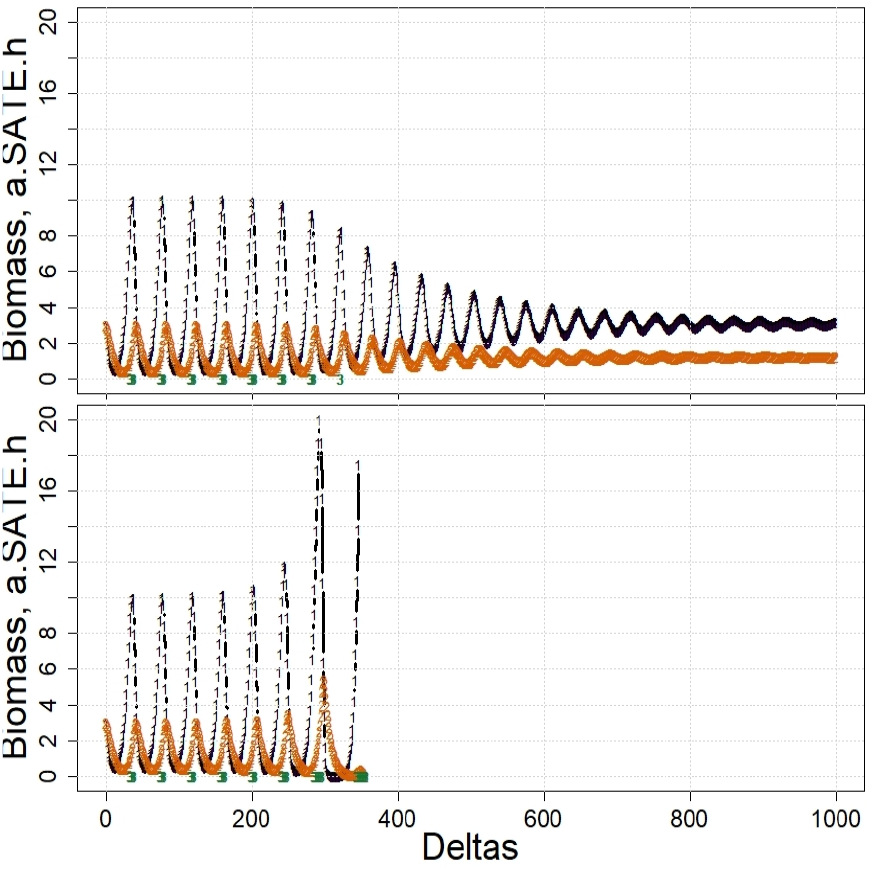
Trial 2. T1(i) repeated with active a.SATE.h and slightly altered 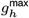 . TOP, Run(i): 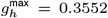. BOTTOM, Run(ii): 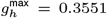. Key: *a*=black 1s, *h*=orange 2s, active a.SATE.h=green 3s at arbitrary ordinate.

##### Conclusions

Initial cycles in both runs showed constant amplitude, in contrast to damping seen with unsatiated grazers in T1(i). Satiation may help explain undamped cycles reported for predator-prey pairs (41), for example if predators migrate away from hunting grounds when sated. Marginally lower **ℊ** ^max^ ⇒ tighter control of harvesting ⇒ more release of growth by *a* ⇒ grazer satiation may be a trigger for algal blooms. The final collapse of *B*_*h*_ despite an excess of food *a* present in run(ii) was a numerical artifact of low 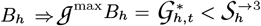 in Eq. (23) (with *w* ← *h* and without the harvesting term) because 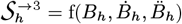, see Annex S1. So 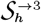 was large because both derivatives were increasing in response to sharply increasing *B*_*a*_.

#### Trials 3 and 4. Competition with one LV wild duplicated

**Aim:** (i) To check that ECOLPS simulated duplicated components as expected. (ii) To simulate competition between duplicates. **Method:** T1(i) was repeated with either *a* (in T3) or *h* (in T4) duplicated. Then one of the duplicates was given a slight competitive advantage. **Trial 3 Runs:** (i) *a*1 and *a*2 exactly duplicated *a* in Trial 1. (ii) **𝓇**_*a*1_ = 0.70, **𝓇**_*a*2_ = 0.75 giving *a*2 the advantage. **Trial 4 Runs** (i) *h*1 and *h*2 exactly duplicated *h* in Trial 1, *h*1<|>*h*2. (ii) **𝒽**_*a,h*1_ = 0.4, **𝒽**_*a,h*2_ = 0.35 giving *h*1 the advantage.

##### Results

**T3 and T4: Runs (i)** Figures 8, 9(top), Table 7. Duplicated components behaved exactly as one. **T3 and T4: Runs (ii)** Figures 8, 9(bottom), Table 7. The slight disadvantage of one component, *a*1 in T3, *h*2 in T4, was revealed in the first cycle then amplified until, by the fourth, the weaker component was almost eliminated.

**Table 7.**
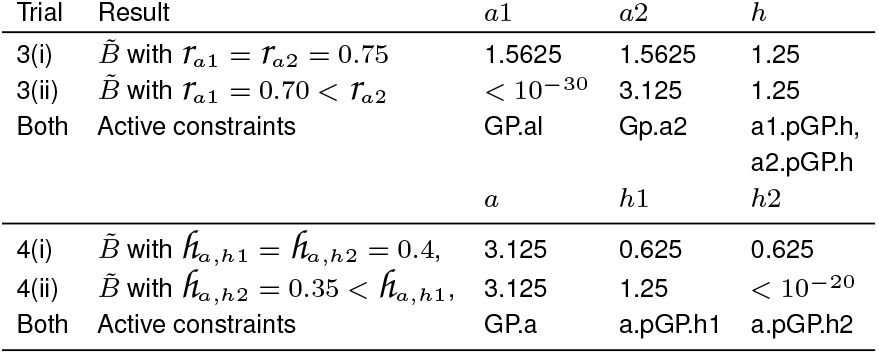
Trials 3 and 4: steady-state biomasses and active constraints with 3 wilds, one duplicated.

| Trial | Result | $a1$ | $a2$ | $h$ |
| --- | --- | --- | --- | --- |
| 3(i) | $\bar{B}$ with $r_{a1} = r_{a2} = 0.75$ | 1.5625 | 1.5625 | 1.25 |
| 3(ii) | $\bar{B}$ with $r_{a1} = 0.70 < r_{a2}$ | $< 10^{-30}$ | 3.125 | 1.25 |
| Both | Active constraints | Gp.a1 | Gp.a2 | a1.pGP.h,<br>a2.pGP.h |
| | | $a$ | $h1$ | $h2$ |
| 4(i) | $\bar{B}$ with $\hat{h}_{a,h1} = \hat{h}_{a,h2} = 0.4$, | 3.125 | 0.625 | 0.625 |
| 4(ii) | $\bar{B}$ with $\hat{h}_{a,h2} = 0.35 < \hat{h}_{a,h1}$, | 3.125 | 1.25 | $< 10^{-20}$ |
| Both | Active constraints | GP.a | a.pGP.h1 | a.pGP.h2 |

**Fig. 8.**
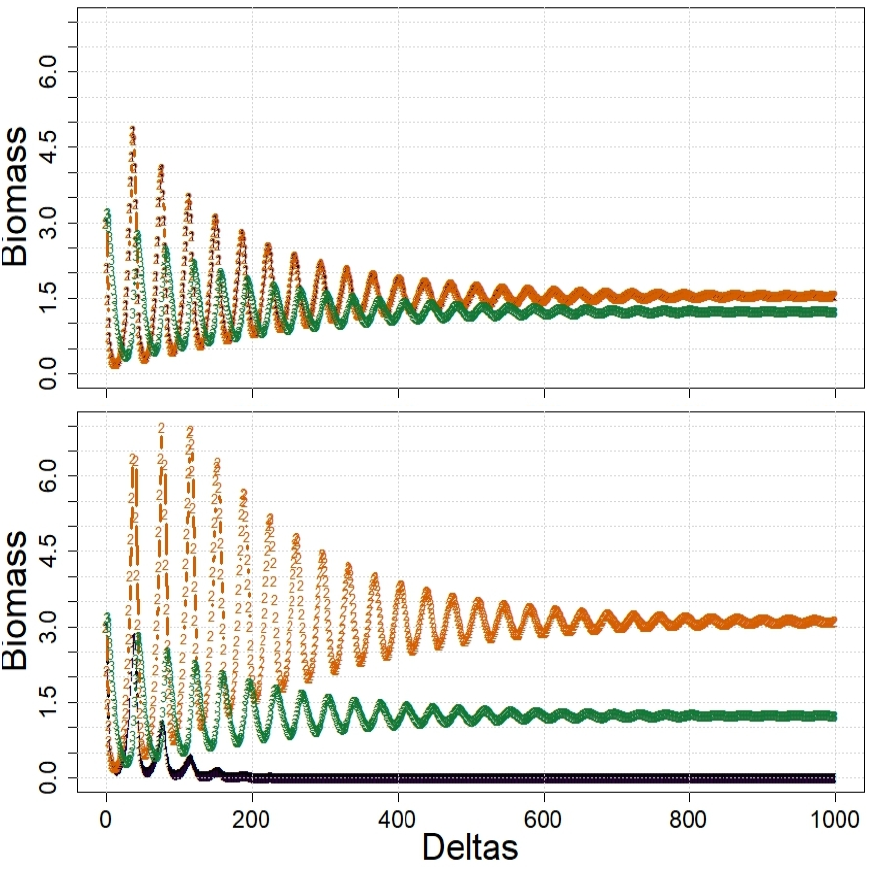
Trial 3. Competition between 2 algas with 1 grazer present simulated over 1000Δs. TOP, Run (i):- Algas *a*1, *a*2 identical (orange 2s masking black 1s). BOTTOM, Run (ii):- *a*1 has reduced **𝓇**_*a*1_ = 0.70. Key: *a*1=black 1s, *a*2=orange 2s, *h*=green 3s.

**Fig. 9.**
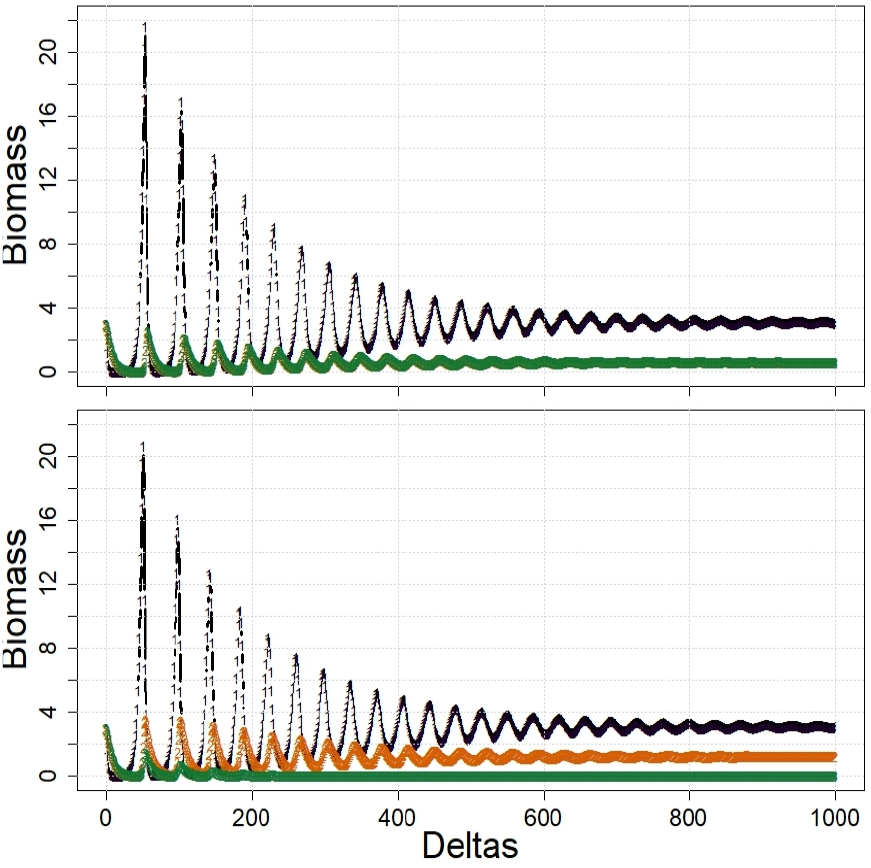
Trial 4. Competition between 2 grazers not eating each other with 1 alga present simulated over 1000Δs. TOP, Run (i):- Grazers *h*1, *h*2 identical (green 3s masking orange 2s). BOTTOM, Run (ii):- *h*2 has reduced **𝒽**_*a,h*2_ = 0.35. Key: *a*=black 1s, *h*1=orange 2s, *h*2=green 3s.

#### Conclusions

Identical replicated *a* or *h* in an LV system behaved as a single component, indicating reliability of ECOLPS with 3 components. A slightly disadvantaged duplicate in this system is competitively excluded (42) over early LV cycles.

#### Trial 5. Competition with both LV wilds duplicated

**Aim:** To simulate competition among paired algas and heterotrophs. **Method:** Both T1 components were duplicated; {*a*1,*a*2} *>* {*h*1, *h*2}, *h*1 *<* | *> h*2. **Runs:** (i) {*a*1, *a*2} and {*h*1, *h*2} pairs identical; see Table 6. (ii) Two different harvesting rates by *h*2:- **𝒽** _*a*2, *h*2_ 035, 0.45 . (iii) As (i) again but now *h*1 *>h*2 with **ℊ**_*h*1,*h*2_ = 0.01.

##### Results

**Run (i)** Fig. 10 (top). Equal grazing gave overlapping, damped LV series. **Run (ii)** Fig. 10 (upper middle). With **𝒽**_*a*2,*h*2_ = 0.35, reduced harvesting by *h*2 on *a*2 ⇒ faster growth of *a*2 ⇒ *h*1 harvested more of *a*1 + *a*2 than *h*2 ⇒ *h*1 responded more vigorously than *h*2 just after *a*1 and *a*2 peaked ⇒ gradual elimination of *h*2, and increase of *a*2 relative to *a*1. Fig. 10 (lower middle). With **𝒽**_*a*2,*h*2_ = 0.45, increased harvesting by *h*2 ⇒ reduction of *a*2 relative to *a*1 but without severe losses of *h*1. **Run (iii)** Fig. 10 (bottom). Low predation of *h*2 on *h*1 ⇒ *h*1 nearly eliminated over 4 cycles. The 2 algas showed stable overlap because *h*2 replaced lost *h*1 and consumed at the same rate.

**Fig. 10.**
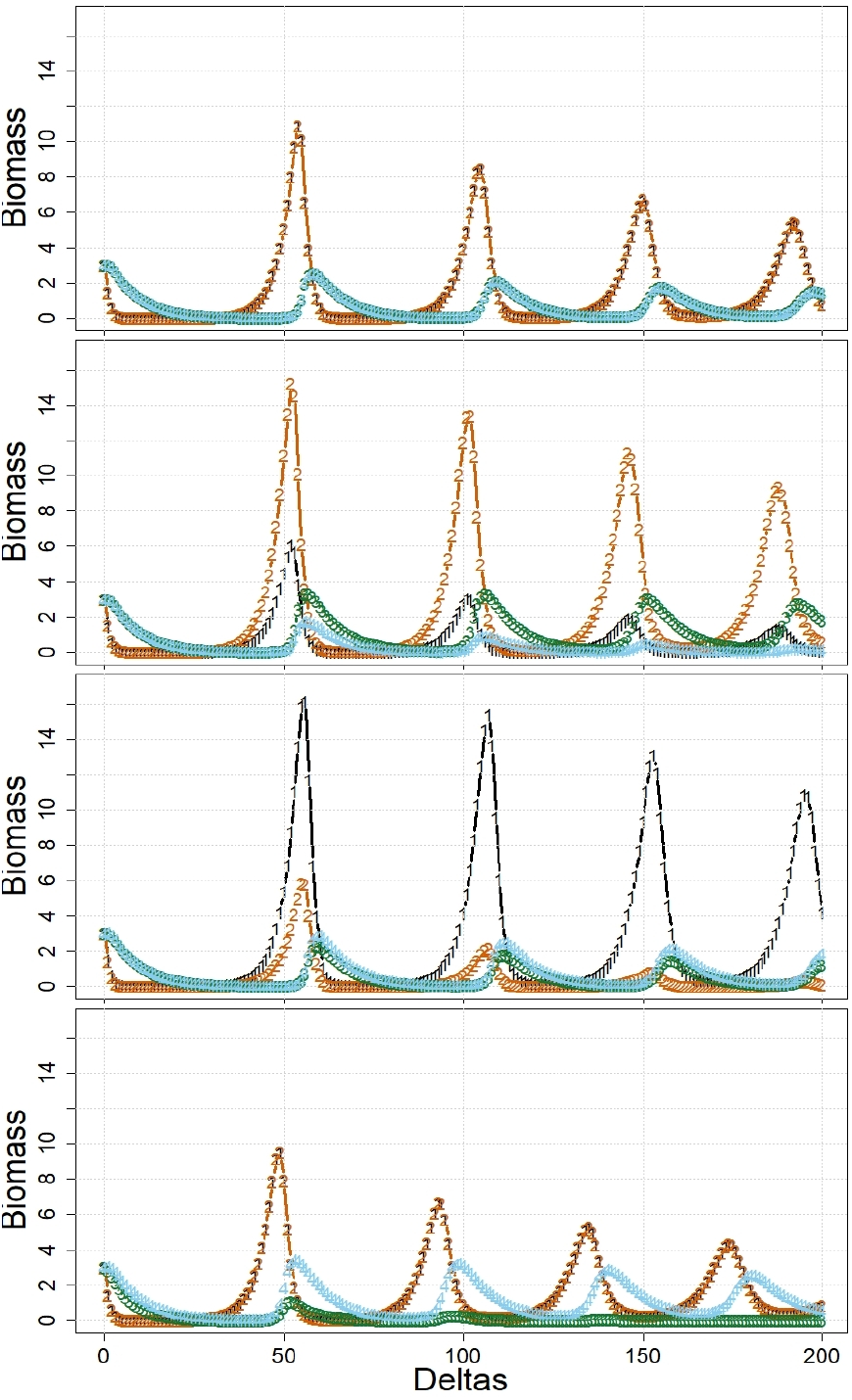
Trial 5. Competition between duplicated alga *a* and grazer *h* from T1(i) ⇒ *a*1, *a*2, *h*1, *h*2, with harvesting abilities of *h*_2_ varying. TOP, Run (i):- *h*1, *h*2 identical, not eating each other, *h*_*a*2,*h*2_ = 0.4 (orange 2s mask black 1s, blue 4s mask green 3s). MID TOP, Run (ii):- Reduced **𝒽**_*a*2,*h*2_ = 0.35. MID BOTTOM, Run (ii):- Increased **𝒽**_*a*2,*h*2_ = 0.45. BOTTOM, Run (iii):- As run(i) except slight consumption of *h*1 by *h*2, **𝒽**_*h*1,*h*2_ = 0.01 (orange 2s mask black 1s). Key: *a*1=black 1s, *a*2=orange 2s, *h*1=green 3s, *h*2=blue 4s.

##### Conclusions

Identical paired heterotrophs not eating each other and competing for the same paired autotrophic foods behaved as one heterotroph and one autotroph, indicating reliability of ECOLPS for 4 components. A small harvesting disadvantage for *h*2 on *a*2 led to demise of *h*2 and the equally preferred food, *a*1, of both grazers. A comparable advantage for *h*2 did not disadvantage *h*1 so severely because, in this case, *h*1 could feed equally on either algal component. One grazer in this system slowly eating the other competing for the same algal foods is soon at an advantage.

#### Trial 6. 3-level food chain: alga *a*, **grazer** *h*, **predator** *p*

##### Aim: To simulate a 3-level food chain; *h>p, a>h, p*| *>h, a*| *>p*

##### Method

Theory for LV steady states, Eq. (5) and (6), was extended to a 3-level food chain, yielding two solutions for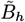, and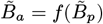

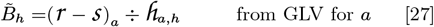

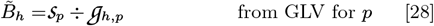

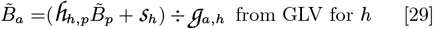

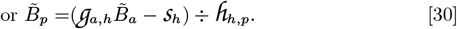

Simulations used only minimal constraints with pvs for *a* and *h* as in Table 6. **Runs** (i) For *p*, **𝓈**_*p*_ = 0.1, **𝒶**_*h,p*_ = 0.15, **𝒽**_*h,p*_ = 0.5333 and *B*_*p*,0_ = 0.3. (ii) **𝒽**_*a,h*_ was increased from 0.4 to 0.5 ⇒ Eq. (28) ≠Eq. (27). (iii) As in (ii) but **𝒽**_*a,h*_ = 0.3.

##### Results

**Run(i)** Figure 11(top). A 3-component steady state found at 3000Δ had 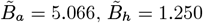 and 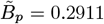. These values are consistent with Eq. (27) to Eq. (30). **Run(ii)** Figure 11(bottom). Increased **𝒽**_*a,h*_ ⇒ an LV system with *B*_*p*_ →0. **Run(iii)** (no figure) Reduced **𝒽**_*a,h*_ ⇒*h* lost more biomass to *p* than it could gain from eating *a* ⇒ *B*_*p*_ increased to 11.2 at 270Δ ⇒ *B*_*h*_, followed by *B*_*p*_, collapsed to 0 ⇒ *B*_*a*_ increased without restraint.

**Fig. 11.**
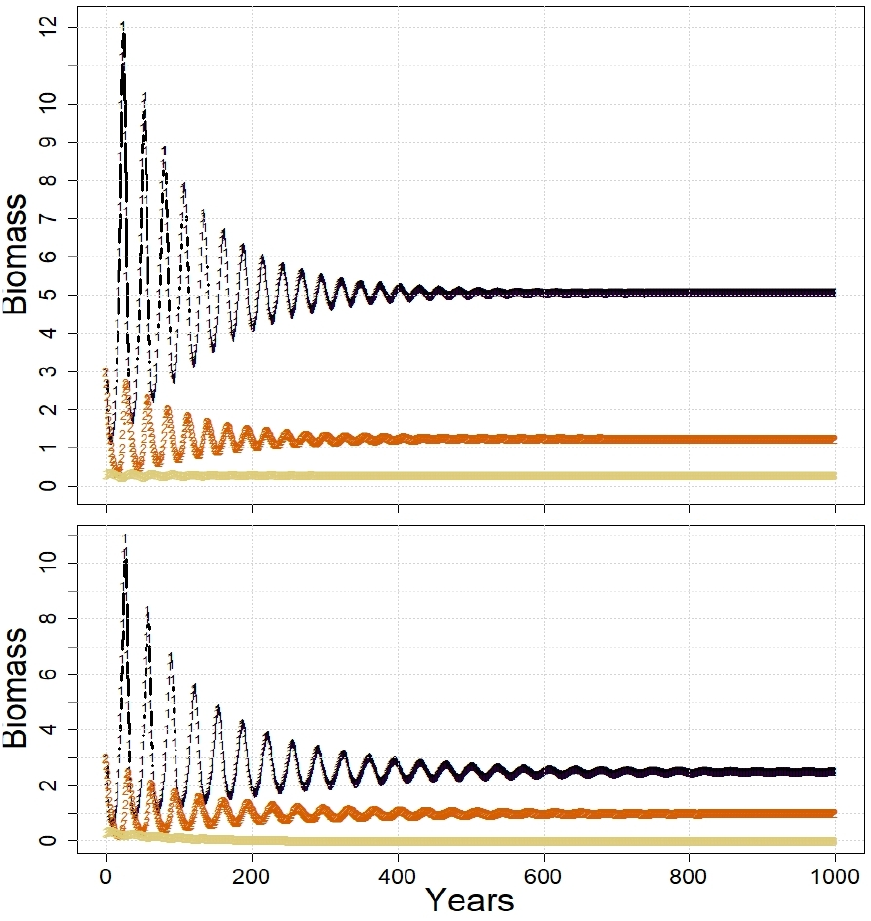
Trial 6. 3-level food chain simulated over 1000Δs. TOP: Run (i) Parameter values set in compliance with Eq. (27)-(30). BOTTOM: Run (ii) Non-compliant **𝒽**_*a,h*_ = 0.5. Key: *a*=black 1s, *h*=orange 2s, *p*=straw 3s.

##### Conclusion

A 3-level food chain is only consistent with the GLV if parameter values are set exactly according to theory. If not, the system reverts to 2 components or to *a* only.

#### Trial 7. Effects of constrained essential habitat on an LV system

**Aim:** To simulate effects of limited essential habitat. **Method:** The minimal LV system with alga *a* and grazer *h* of T1 and Table 6, was simulated over 200Δ with limited essential habitat *z*. **Runs:** (i) z.HAB.a constrained rGP of alga *a* with occupancy **𝓋**_*z,a*_ = 1, habitat size *A*_*z*_ = 3.05 – just too small to accommodate the steady-state biomass, 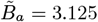, found in T1. (ii) z.HAB.h constrained rGP of grazer *h* with **𝓋**_*z,h*_ = 1, and *A*_*z*_ = 1.2 – just too small for 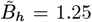. To prevent activity of z.HAB.h initially, *B*_*h,t*=0_ = 1 was reduced from 3 in T1.

##### Results

**Run(i)** Figure 12(top). Following initial instability brought about by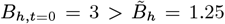, habitat *z* was filled with *B*_*a*_ at 26Δs ⇒ z.HAB.a constrained *B*_*a*_ ⇒ indirect constraint of 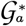by reduction of the RS limit of GP.a, that is 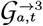; see Table 5 and Eq. (22). **Analysis:** In this case, referring to terms in Eq. (23) with *w* ← *a*, the start of activity by z.HAB.h 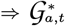 was initially larger than needed to maintain constrained 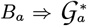 not fitting in *z* was available for consumption by *h* ⇒ *B*_*h*_ gradually increased subtraction of a larger harvest of *B*_*a*_ by *B*_*h*_, see RS term 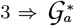, the RS term 4, could also gradually increase. The two variables converged asymptotically.

**Fig. 12.**
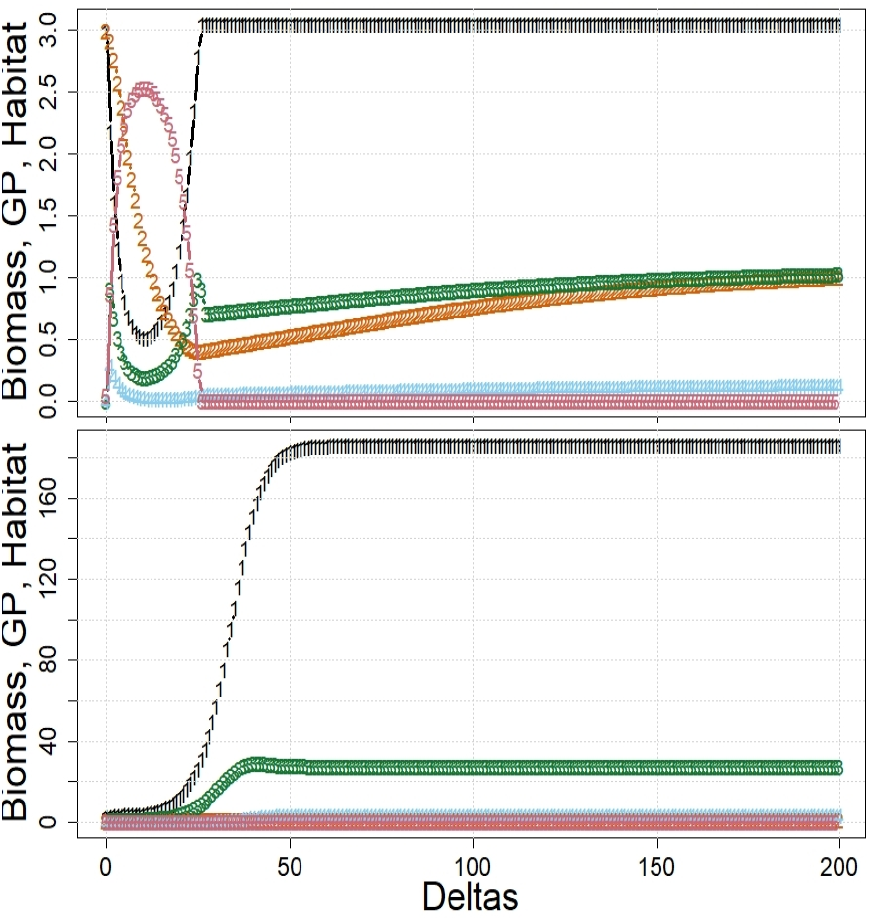
Trial 7. Effects of essential habitat. TOP, Run(i): Alga *a* constrained by z.HAB.a with *B*_*a*,0_ = 3, *A*_*z*_ = 3.05, **𝓋**_*z,a*_ = 1. BOTTOM, Run(ii): Grazer *h* constrained by z.HAB.h with *B*_*h*,0_ = 1.0, *A*_*z*_ = 1.24, and **𝓋**_*z,h*_ = 1. Key, *B*_*a*_=black 1s; *B*_*h*_=orange 2s;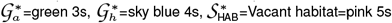.

**Run(ii)** Figure 12(bottom). GP.a and a.pGP.h were active over the first 9Δs ⇒*B*_*h*_ rose from 1 to the z.HAB.h limit= 1.24, and 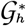 rose gradually from 0.12 reaching an asymptote at 3.68 after approximately 60Δ (obscured by ordinate scaling). *B*_*a*_ rose slowly at first, then rapidly after 10Δ when *B*_*h*_ was constant, reaching asymptote *B*_*a*_ ≈ 186 at 60Δs.

##### Conclusions

A habitat-constrained food in an LV system can allow gradual increases of both 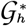 and *B*_*h*_ because *h* feeds on GP of *a* that cannot be accommodated as rGP in *z*. A habitat-constrained consumer can allow the food to grow to a higher asymptote. HAB. constraints primarily control rGP, secondarily GP. They remove periodicity from an LV system.

#### Trial 8. LV system with unreplenished essential nutrient N1

**Aim:** To simulate responses of algal component *a* to low levels of nutrient N1 required for photosynthesis. **Method:** Pvs are from T1(i) in Table 6 and, for N1, in Annex S4: Table S21.

##### Result

Figure 13. *B*_*a*_ and *B*_*h*_ collapsed to zero only when n.NUTR.a took over activity from GP.a.

**Fig. 13.**
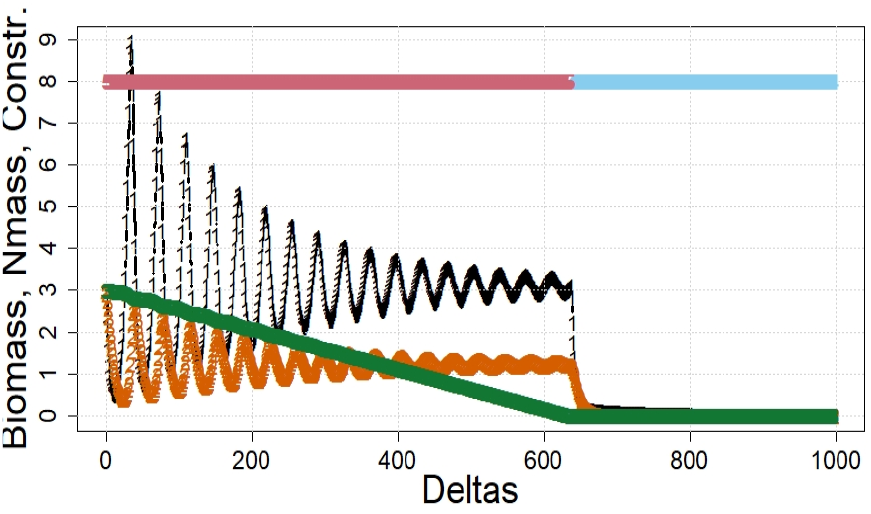
Trial 8. Effects of consumption of nutrient *n* in water on dependent alga *a* in an LV system. Key: *B*_*a*_=black 1s; *B*_*h*_=orange 2s; *Q*_N1,H2O_=green 3s. Active constraints (at arbitrary ordinate): GP.a=pink, *n*.NUTR.a=blue.

##### Conclusion

N1 had no effect on simulated GP of a dependent autotroph until none was available ⇒ collapse.

#### Trial 9. 3-level food chain with inputs of N1, and fishery *f*

##### Aims

(a) To test whether expected instability of a 3-level food chain with arbitrary pvs, as found in T6, is stabilized by regular inputs to H2O of nutrient N1 essential to *a*, and (b) to identify possible effects of fishery *f* on the food chain.

##### Method

An LV model with grazer *h* and alga *a* was supplemented with predator *p* eating only *h*. **𝒽**_*h,p*_ = 0 was set initially, then increased gradually until *B*_*h*_ was cropped sufficiently to allow *B*_*a*_ to increase when low *B*_*h*_ and constraints permitted. Fisheries were bled in similarly with initial low values of **𝒽**_*•,f*_ until effects on the food chain were evident. _N1,*a*_ was adjusted so that N1 inputs on day 1 of each year of 12 ‘months’ (=12Δs) would sustain positive **𝒢**_*a*_ only for the early part of each year before N1 was totally depleted and N1.NUTR.a became active. **𝒸**_N1,*h*_ and **𝒸**_N1,*p*_ were set low to prevent activity by N1.NUTR.h or N1.NUTR.p. Final pvs are in Annex S4: Tables S22-S25. **Runs:** (i) A low initial N1, *Q*_N1,H2O,*t*=0_ = 0.001*γ* (mass units), was set merely to signal nutrient presence to ECOLPS. 0.7*γ* of N1 was added once annually. (ii) fishery *f* with effort 1 unit per Δ was added; *p>f, h*|*>f* . (iii) As in (ii) but *h>f, p*|*>f* .

##### Results

**Run (i)** Figure 14(left). Annual inputs of N1 ⇒ annual cycles of biomass because N1.NUTR.a was active (when *Q*_N1,H2O_ = 0*γ*) for 6 or 7 months in every year ⇒ grazing by *h* exceeded GP of *a* ⇒ increases of *B*_*a*_ to peaks were reversed. *h* itself was controlled by food *a* and predator *p*. Cycles of *B*_*p*_ had small amplitude, sustained above 0 when food *h* was low by low self-loss rates, **𝓈**_*p*_ = 0.2.

**Fig. 14.**
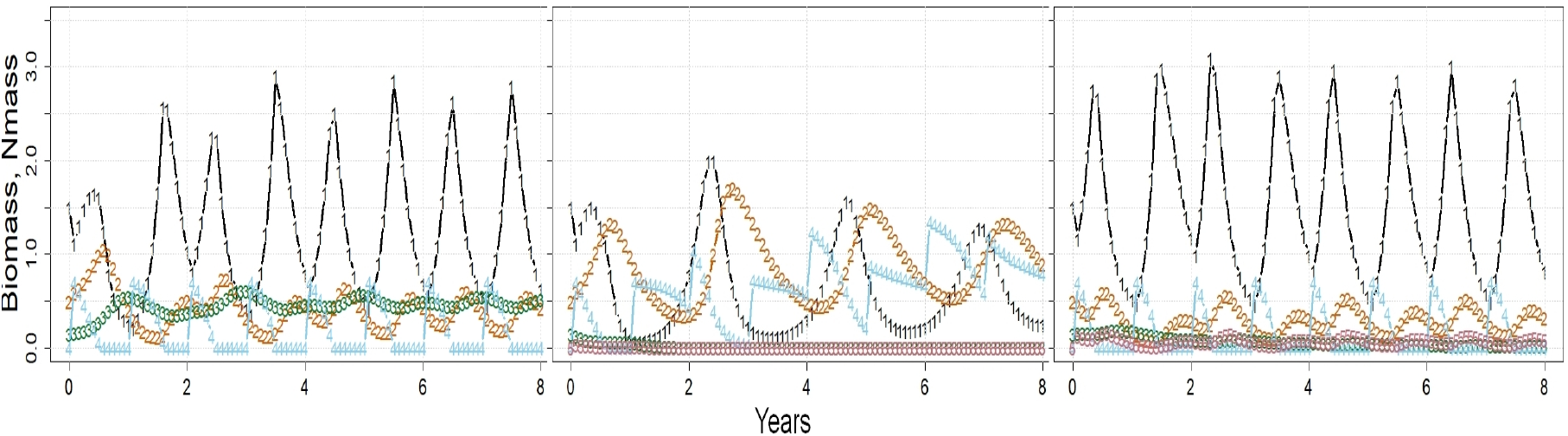
Trial 9. Biomasses of alga *a*, grazer *h* eating *a*, predator *p* eating *h*, with inputs to water of 0.7*γ* (mass units) of nutrient N1, essential to *a*, on day 1 of each year. LEFT:- Run (i) No fishing. CENTER:- Run (ii) Fishing on *p* only. RIGHT:- Run (iii) Fishing comparably on *h* only. Key: *B*_*a*_ =black 1s, *B*_*h*_ =orange 2s, *B*_*p*_ =green 3s, *Q*_N1,,H2O_ =blue 4s; **𝒞**_*p,f*_ in (ii), and **𝒞**_*h,f*_ in (iii)=pink 5s.

**Run (ii)** Figure 14(center). Catches of *p* by *f* depleted *B*_*p*_ to almost zero; see pink 5s masking green 3s in the figure ⇒ increase of *B*_*h*_ compared to Run(i) ⇒ substantially reduced peaks, and valleys of *B*_*a*_ cycles ⇒ slower response of *a* to N1 inputs ⇒ biennial cycling of both *B*_*a*_ and *B*_*h*_ ⇒ slower consumption of N1 in H2O ⇒ lack of constraint of *B*_*a*_ by N1.NUTR.a after year 2 ⇒ increasing *Q*_N1,H2O_.

**Run (iii)** Figure 14(right). Catches of *h* by *f* ⇒ lowered cycles of *B*_*h*_ ⇒ reduced grazing on *a* ⇒ slightly higher peaks of *B*_*a*_ than in Run (i) ⇒ *Q*_N1,H2O_ reached 0*γ* in every year N1.NUTR.a constrained GP of *a* annually ⇒ annual cycles of *B*_*a*_. Also, reduced *B*_*h*_ ⇒ less food and much reduced *B*_*p*_ compared to Run (i) though not fished in either run.

##### Conclusions

The unstable 3-level food chain of T6 can be stabilized if GP of *a* depends on nutrient N1 supplied and exhausted annually, thereby limiting growth of *a* each year. T9(ii), with fishing on top predator *p* ⇒ high *B*_*h*_⇒ low *B*_*a*_, consistently simulated a top-down trophic cascade (8, 43). T9(iii), with fishing on intermediate predator *h* ⇒ starved and depleted *B*_*p*_ despite high *B*_*a*_ – a ‘wasp-waist’ system (44).

#### Trial 10. 3-level food chain re-cycling variable mass of N1

**Aim:** To simulate a system like T9(i) but with recycling of N1 through water, NLO *d* and scavenger *s*, instead of with annual inputs. **Method:** *Q*_N1,,H2O,*t*=0_ = *Q*_N1,,NLO,*t*=0_ = 0.35*γ*, were set equally to match the single annual input of N1 in T9(i), 0.7*γ*. Final pvs are in Annex S4: Table S26.

##### Results

Figure 15. A self-sustaining recycling system highly sensitive to pvs was found. Initial oscillations resolved after 2 years to approximately steady states for most series including *Q*_N1,Σ_, in contrast to the annual oscillations seen in T9(i), Figure 11(left). The 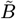 were as high (*B*_*h*_) or higher (*B*_*a*_ and *B*_*p*_) than peaks seen in T9(i). N1.NUTR.a was active from 14 months. **Analysis:** in T9(i), growth of *B*_*a*_ started each year from a low level after exhaustion of N1, whereas in T10, N1 could be re-used as it was released, dissolved, by self losses ⇒ *B*_*a*_ reached a higher level than the peaks in T9(i) even though *Q*_N1,H2O_ reached zero in year 1.

**Fig. 15.**
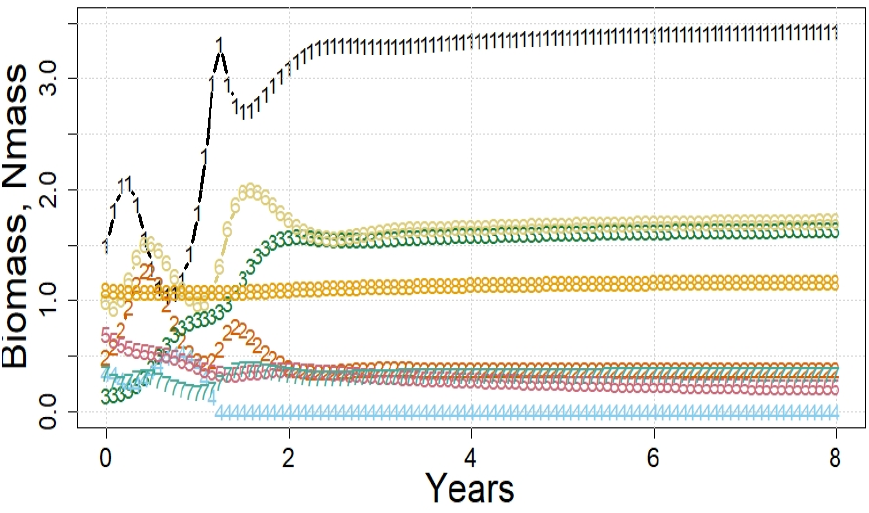
Trial 10. Trial 9(i) repeated with recycling of N1 via NLO *d* and scavenger *s*; annual inputs of N1 removed. Key: as in Figure 11 with, also, *B*_*s*_ =pink 5s, *B*_*d*_ =straw 6s, *Q*_N1,NLO_ =turquoise 7s, *Q*_N1,Σ_ =gold 8s.

##### Conclusions

An unstable 3-level food chain can be sustained by recycling N1 via H2O, NLO and a scavenger. *B*_*a*_ can then reach higher levels than when N1 is input annually. The rate of recycling stabilized the limit of active N1.NUTR.a near 0 ⇒ stable *B*_*a*_ and other biomasses dependent on it.

#### Trial 11. 3-level food chain, seasonal light cycles, recycling N1, and fishery *f*

**Aims:** (a) To simulate T10 with seasons using annual sinusoidal cycles of light index *I*_*t*_. (b) To compare effects of fishing when N1 is being recycled to those found in T9 without recycling. **Method:** Applying Eq. (26) with {*l*_*T*_, *h*_*T*_} = 0, 1 to an annual cycle of *I*_*t*_ from Trial11.Envir.txt, **𝓇**_*a*_(*I*_*t*_) cycled between 0.199 at *I*_*t*=*x*_ = 0.131 and 1.324 at *I*_*t*=*x*+0.5year_ = 0.870 in all runs. Pvs for all runs of T11 were initially from T10, adjusted as necessary for stability; final pvs are in SI Tables S27-S30. **Runs:** (i) As T10 but with light-driven seasons. (ii) fishery *f* with effort 1 unit per Δ was added; *p>f, h*|*>f* . (iii) As in (ii) but *h>f, p*|*>f* .

##### Results

**Run (i)** Figure 16(left); cf. Figure 14left), Figure 15. Seasonal **𝓇**_*a*_(*I*_*t*_) altered the steady states found in T10 to annual cycles. To improve comparability with T9(i) given the changed conditions: (a) **𝒶** was reduced from 0.6 to 0.47⇒ peaks of *B*_*h*_ *> B*_*p*_ as in T9(i). (b) **𝓈**_*h*_ was increased from 0.25 to 0.5 ⇒ steadier annual peaks of *B*_*a*_ more comparable with T9(i). (c) **𝒽**_*d,s*_ was increased from 0.176 to 0.2994 ⇒ flatter *Q*_N1,Σ_ to 16 years, meaning better N1 accounting. *Q*_N1,H2O_ was depleted annually at peak *B*_*a*_ for 1 to 3 months and, as in T10, peaks of *B*_*a*_ were slightly higher in this recycling system than in T9(i).

**Fig. 16.**
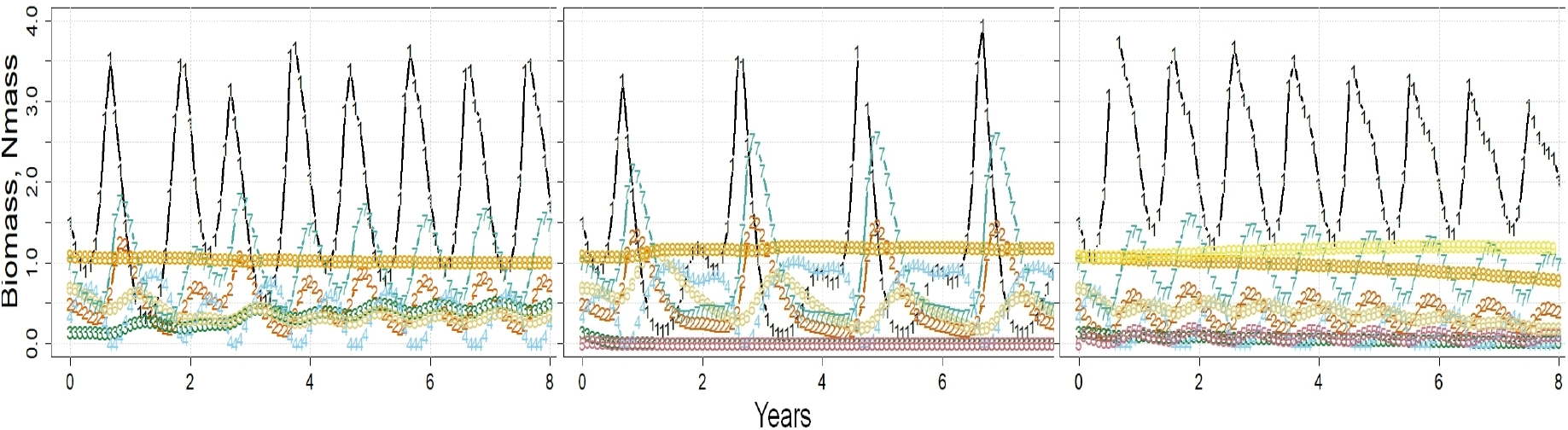
Trial 11. Trial 9(i) repeated but with recycled nutrient N1 instead of annual inputs, and with self-growth of alga *a*, **𝓇**_*a*_(*I*_*t*_), varying with a seasonal light cycle, low in ‘Jan’, high in ‘June’ (not shown). LEFT:- No fishery. CENTER:- Fishing on predator *p* only. RIGHT:- Fishing comparably on grazer *h* only. Key: see Fig. 14; also *B*_*s*_=straw 6s, *B*_*d*_=turquoise 7s *Q*_N1,Σ_ =gold 8s; in right panel only, *Q*_N1,Σ_ + *Q*_N1,Landings_=yellow 9s.

**Run (ii)** Figure 16(center); cf. Figure 14(center). Fishing on *p* reduced *B*_*p*_ and **𝒞**_*p,f*_ to almost zero ⇒ negligible N1 was removed by fishing (not shown). A decline of *Q*_N1,Σ_ was flattened from year 1 by decreasing **𝓈**_*d*_ from 0.8 to 0.425. Comparing peak heights of biomasses of Run(ii) with those of the unfished Run(i), those in T11 were closer in value than those in T9, suggesting that recycling of N1 reduced disturbance caused by fishing *p* with the same effort. Low *B*_*p*_ ⇒ slower declines of *B*_*h*_, mainly by self losses. Annual valleys *B*_*a*_ were alternately high and low ⇒ simultaneous levels *B*_*h*_ were low and high ⇒ the next ‘summer’ peak of *B*_*a*_ was high or low respectively because of the altered grazing rates. Biennial cycles also spread to *B*_*d*_ and *B*_*s*_. The rate of N1 recycling in T11(ii) was held steady by biennial activity of N1.NUTR.a, in contrast to the unchecked rise of N1 seen from year 3 in T9(ii) in response to annual inputs that exceeded needs.

**Run (iii)** Figure 16(right); cf. Figure 14(right). Fishing on *h* ⇒ declining annual cycles of most series including *Q*_N1,Σ_. They were caused by removal of N1 in landings of *h* which were larger and had higher **𝒸**_N1_ than landings of *p* in T11(ii) ⇒ *Q*_N1,Σ_+ cumulative *Q*_N1,Landings_ (yellow 9s in Figure 16(right)) was nearly flat. The period when *Q*_N1,H2O_ = 0 increased with decreasing *Q*_N1,Σ_ from 4Δ in year 0 to 5Δ in year 7 ⇒ the gradual decline of peaks of *B*_*a*_ and dependent *B* series. Peaks of *B*_*a*_ were higher than in T9(iii) because activity of N1.NUTR.a was longer, 7 to 8Δ in each year in T9(iii). *p* was not fished in T11(i) or (iii) but levels of *B*_*p*_ were higher in T11(i) suggesting that high **𝒢**_*a*_ in T11(iii) was not being transferred efficiently via *h* to *p*.

##### Conclusions

Light-driven seasonal growth with recycling N1 can be simulated by ECOLPS with approximately constant *Q*_N1,Σ_ though it is sensitive to pvs. Low recycling rates in T11(i) ⇒ exhaustion of N1 annually by **𝒢**_*a*_ ⇒ annual biomass cycles. Trophic cascades occurred biennially in T11(ii) when fishing on *p* resulted in relatively high levels of *B*_*h*_ and low levels of *B*_*a*_ in the next ‘summer’ season. Fishing on *h* in T11(iii) ⇒ lower *B*_*p*_ than when unfished in T11(i), implying that T11(iii) was a wasp-waist system like T9(iii). Allowance for cumulative removals of nutrient in landings of a significant fishery can account for downward trends of biomass cycles. Simulated nutrient recycling can reproduce similar but not identical ecological effects to those when N1 is added externally, as in Trial 9.

#### Trial 12. Food web with seasonal light and temperature cycles, recycled N1, and fishery *f*

**Aim:** To simulate a more complex, nutrient-recycling ecosystem than previous trials together with seasonal succession of algal components based on seasonal light and temperature indices. (Succession caused by seasonal mixing of nutrients was not investigated.) **Method:** Annually cycling temperature indices *T*_*t*_ were present in Trial12.Envir.txt along with the light indices *I*_*t*_ previously used in Trial11; see Annex S4: Table S31. *T*_*t*_ cycles lagged *I*_*t*_ by 63 days. Three copies of the 3-level food chain of Trial11.Run1.txt were made, named Trial12.Run1(a,b,c).txt. Component and nutrient names were numbered 1-3 to match. Light-dependent variations of photosynthesis in Trial 11(i) were supplemented with different temperature-dependent restrictions for Alga1, 2 and 3 respectively: {*l*_*T*_, *H*_*t*_ } = {0.2, 0.4}, {0.4, 0.7},{ 0.65, 1.0} ; see Eq. (26). Thus Alga1 could grow only in early spring, Alga2 twice yearly in early and late summer, and Alga3 only in high in summer. These restrictions necessitated adjustments of pvs to sustain each system independently over 16 years with approximately constant *Q*_{N1, N2, N3},Σ_. Final pvs of the 3 separately run files are in Annex S4: Table S32. For Run(ii), the 3 adjusted food chains were placed side-by-side in Trial12.Run2.txt, with 3 algas, 3 grazers and 3 predators; NLO, scavenger, and H2O components were unique, serving each chain jointly. Further adjustments of pvs were needed to sustain the complete system over 16 years with seasonal successions and approximately constant *Q*_{N1, N2, N3},Σ_. Final pvs are in Annex S4, Table S33.

##### Results

Looking only at seasonal succession of algas, Figure 17 shows that *B*_A_ peaks occurred in the same order as the photosynthetically active periods, Alga1 before Alga2 before Alga3, with significant secondary peaks for Alga2, as expected in every year. On one occasion (5.417y) N1.NUTR.a1 was activated by Alga1 using N1 faster than supplied by recycling. N2.NUTR.a2 was activated after the first peak of Alga2 but not after the second, lower peak in each year. N3.NUTR.a3 was active at each peak of *B*_*a*3_ and for 3 years after. Not shown are the approximately level *Q*_N1,2,3,Σ_ and purely annual cycles of grazers and predators.

**Fig. 17.**
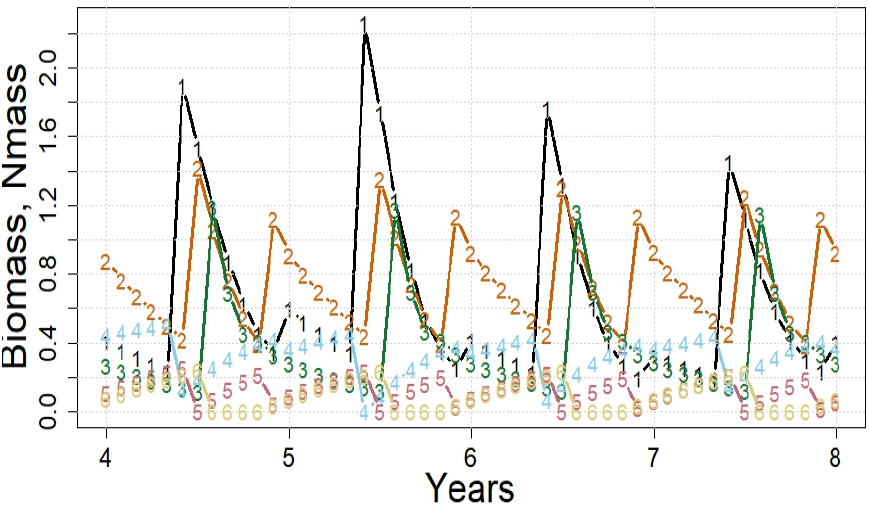
Trial 12. Simulated succession of 3 algas brought about by programmed seasonal cycles of temperature and light indices, with recycling of nutrients via NLO and a scavenger. Key: *B*_*a*1_=black 1s, *B*_*a*2_=red 2s, *B*_*a*3_=green 3s, *Q*_N1_ =blue 4s, *Q*_N2_ =pink 5s, *Q*_N3_ =straw 6s.

##### Conclusions

Multi-chain systems can be simulated with light- and temperature-related seasonal annual or biannual succession. Setting pvs for individual food chains, then combining them side-by-side was a feasible but awkward method of progressing towards an ecosystem with a food web, rather than separate food chains. Readers may re-run this system to see other outputs or, possibly, to try to elaborate it with top predators, nutrient sharing across chains, and/or fisheries.

## Discussion

Trial simulations have shown that the constraint-based algorithm implemented by ECOLPS can simulate time series of wild biomasses consistently with expectations from theory [T1, T6(i)], and from simple logic when wilds are exactly duplicated [T3(i), T4(i), T5]. Utility of ECOLPS is further supported by its ability to generate, and reveal explanations for, ecological events occurring in the simulation. They included LV cycles [T1(i, ii)], grazer satiation possibly triggering algal blooms [T2], competitive inhibition (42) from marginal disadvantages [T3,4], alteration of diet in response to reduced competition [T5(ii)], food chain stabilization with nutrients [T9(i)], essential habitat affecting food and consumer differently [T7], on-off control of photosynthesis by an essential nutrient [T8], alteration of biomass cycles and food chains by fisheries and/or a nutrient whether input or recycled [T9,11], loss of nutrient via fishery landings [T11] and, lastly, seasonal successions of algas [T12(ii)]. Given the broad scope of these simulated events, a constrained-GP hypothesis for individual organisms may serve to explain varied behavior of aquatic ecosystems:-

> *Every living aquatic organism seeks to maximize its rate of GP, and thus its survival and reproductive success, subject to other behavioural requirements to, for example, reproduce, escape predators, migrate, digest food, or rest. Its maximum GP rate is limited by:- either the rate of the slowest contributing process, or a lack of food or other essential resource, or predation or other impediment to its healthy existence. The identity of the active constraint (with, possibly, redundant active constraints) and its limit for each individual vary over time and space*.

The truth of this hypothesis, and of findings of trials on which it is based, might be assessable practically using, as one suggestion, laboratory microcosms. Note that an individual need have only one active constraint at maxGP whereas ecosystems must have at least one for every component contributing to maxΣGP; see Eq. (1). Constraints on rGP, not on GP directly, are special. In the case of z.HAB.w [T7], an individual is either alive in essential habitat *z* with its GP limited by another constraint or, if not in *z*, will not survive.

ECOLPS has demonstrated several useful features. Predator diets are automatically altered by FOOD constraints to reflect the availability of their preferred foods. All component types present in the simulated ecosystem are dealt with by automatically bringing in the relevant constraints. Nutrient recycling, seasonality and successions are carried out if appropriate components and parameters are supplied. Quick, stepped calculations, with pvs modifiable over time, permit thorough exploration of model ecosystems without the need, in function-based modeling, to select functions and error distributions, possibly to adjust formulations, then to solve equations, some of which may present non-linear or multivariate complications.

Nevertheless, more complex simulations are not yet straightforward:-

- Choosing pvs so that all intended wild and fishery components are viable is difficult using only ecological logic and trial-and-error, as for the reported trials. This is unsurprising because real ecosystems are not a haphazard mix of component types but evolve into a compatible group of components over long periods by competition, migrations, acclimatizations and adaptations. Possibly, a system might be set up to simulate repeatedly with different pvs, then to select the set consistently yielding the highest ΣGP, implying that the community is exploiting the ecosystem to the fullest extent. The search is restricted to relative pvs and may not need to include resource constraints.
- Maintaining all *Q*_*n*,Σ_ constant in a closed ecosystem here required awkward *ad hoc* adjustment of pvs. An equality constraint in LP is ineffective because *Q*_*n*,H2O_ and *Q*_*n*,NLO_, Eq. (20), are not in the objective function, Eq. (1). Also, corrections to *Q*_*n*_ between Δ_*t*_ interfered with nutrient recycling calculations. Real, closed ecosystems, logically, should have constant *Q*, implying that a method of achieving this in a simulation also exists.
- Untrialed code in ECOLPS permits transfers of biomass among components between Δ_*t*_. When fully tested, this could allow ontogenetic transfers between components, as needed when a wild has life stages that fit into different component types.
- Other untrialed code allows projections of biomasses harvested but not assimilated (**𝒶**_*w*,*f*_ = 0) and thus not estimable by LP. These could allow estimates of totally discarded fishery catches, for example when demersal trawls strip environmentally important epibenthos from the sea floor.
- Symbioses and parasitism are important in many aquatic ecosystems but cannot yet be processed by ECOLPS.
- Lastly, restriction of simulations to closed ecosystems could be removed by dividing an aquatic region into contiguous cells with net mass transfers after each Δ_*t*_. Transfer terms would then have to be included in constraint-limit formulas.

ECOLPS might assist ecological fieldwork by suggesting priorities for, or explanations of observations. Another application could be for environmental risk assessment (19, 20) as part of an ecological approach to management (45). Simulations might help find risks of high nutrient levels, algal blooms, trophic cascades, loss of important species, depletion of fish stocks etc. Use for predicting future states of the ecosystem is not recommended (11). They could be wrong due to inappropriate pvs, omission of relevant constraints and trophic interactions, or due to unknown effects of environmental factors, or fluxes into or out of the ecosystem. At best, predictions could only provide relative biomasses, inappropriate for absolute management measures such as fishery quota.

Development of constraint-based ecosystem modeling may have been delayed historically by lack of an agreed objective function. ΣGP, used here, is simpler to maximize than other production functions because of its monotonicity and its independence from other processes such as respiration (for net production) and mortality (for standing crop). Wild individuals do not, themselves, have foreknowledge of respiratory or biomass losses and must, therefore, consistently maximize their GP for best chance of survival through their uncertain futures. This argument implies that ΣGP is both practical and realistic as an objective function.

Simulated ecosystems were found to be very sensitive to small adjustments of pvs, for example, the precise pivotal value of **ℊ** ^max^ found to separate damping from amplification of LV cycles in Trial 2, and the competitive exclusion (42) of marginally disadvantaged living components in T3 – T6. This finding suggests that equilibrium biomasses, 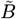, used here for comparisons with theory and, elsewhere, in function-based models for calculation of pvs, have little relevance to real ecosystems because values of parameters and environmental variables are unlikely to be constant long enough for steady states to be reached. Step-wise analyses of simulated time series offer better information because an event at one step may be explicable by previous steps in the same or other series. Also, amplification of small changes may be visible over a few cycles, as in Trials 3 and 4. Thus interpretations are less reliant on extended constant conditions. Addition of stochasticity to the **𝒢**^∗^ might enhance the apparent realism of simulations since very precise responses, as seen in the trials, are unlikely in reality. However, causes would then be harder to identify.

The LV model, central here, has been much criticized and modified (18, 46) possibly as a consequence of the low flexibility of some function-based modeling, or of over-wide domains assumed for functions. In support of the LV model, the harvesting term, **𝒽***B*_*i*_*B*_*j*_ becomes zero when either *B*_*i*_ or *B*_*j*_ is zero, harvesting increases in proportion to food and consumer presence, and exponential growth occurs when a biomass is unconstrained, all of which are reasonable expectations for short time intervals Δ. Extra terms in the LV model would complicate formulation of LP constraints and appear unnecessary for step-wise constraint-based modeling. Other advantages of LV, used here, are that it can be simply generalized to **𝓃**_*W*_ wilds as in Eq. (10), then differentiated analytically to provide approximations of integrated quantities over short time periods as in Annex S1.

The present ecological application of LP contrasts with typical commercial applications involving industrial processes. There, the optimized objective function and weightings of terms in it are critical for calculating costs and profits whereas here, the focus is on the 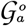 and 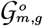 and the calculated biomasses. Unit weights given to all sources of GP are consistent with ΣGP as the objective function but, if some other function is preferred, weights should reflect what drives the system, not what Man wants from it. Financial weightings, for example, have no effect on ecological processes.

## Supporting information

Annexes

ECOLPS simulator in R

Trials1-12 parameters

## Acknowledgements

Fadia, my wife, tolerated, supported, even encouraged this oddity on my bucket list. Providers of R, Tinn-R, Zotero, IrfanView, L^A^T_E_X 2*ε*, Overleaf, Open Office, WinMerge, Google Scholar and research papers freely on the internet are thanked wholeheartedly for enabling my unfunded research. This pre-print was formatted with a PNAS L^A^T_E_X 2*ε*template but is not a PNAS publication.

