## Supplementary material for "Virtual aquatic ecology: stepped simulations of gross production within constraints explain biomass changes": Annexes

December 2, 2025

Note: Table, figure and equation numbers should be S1, S2 etc in these Annexes with 'S' for 'supplementary' to distinguish from Table 1, Figure 2 etc in the main paper. Numbers should start freshly from 1 in each annex. However, I have not found ways to typeset these features here in Latex.

#### 1 Annex S1. Mathematical supplement

##### 1.1 Projecting and integrating GLV rates over $\Delta_t$

Two constraint types, GP.a and m.pGP.g require projected integrals of gross production (GP or $\mathcal{G}$) terms over short time period $\Delta_t$ to serve as upper limits for GP in that period, starting from zero at $t$. Other constraints require projections of self losses of biomass ($\mathcal{S}$), also starting from zero. Taylor's theorem (Abbott 1940) permits approximate projections of a variable's value (Ayres 1972) at $t + \Delta$ using its known value and derivatives at $t$ treated as constant over $\Delta_t$. For example, a 3[rd]-order projection of biomasses in an $n_{\mathrm{W}}$-vector $\mathbf{b}_t$ from $t$ to $t + \Delta$ is

$$\mathbf{b}_{t+\Delta}^{\to 3} = \mathbf{b}_t + \dot{\mathbf{b}}_t \Delta + \ddot{\mathbf{b}}_t \frac{\Delta^2}{2} + \dddot{\mathbf{b}}_t \frac{\Delta^3}{6}. \tag{1}$$

Equation (6) to (10) below supply formulas for the biomass differentials. Covariation among the different component biomasses of the $\mathbf{b}$ vector over time is allowed for by the conversion $\mathcal{g}$ and harvesting $\mathcal{h}$ terms in these formulas. $\mathbf{b}_t^{\to 3} - \mathbf{b}_t$ approximates net changes in biomasses over $\Delta_t$. In this study, $\Delta_t = 1$ time unit always.

For a 3[rd]-order projection of *integrated* biomasses, let $\tau$ track time locally and continuously between 0 at $t$ and $\Delta$ at $t+\Delta$:-

$$\begin{aligned}\int_0^\Delta \mathbf{b}_{t+\tau}.d\tau &\approx \int_0^\Delta \mathbf{b}_{t+\tau}^{\rightarrow 3}.d\tau \\ &= \int_0^\Delta \left(\mathbf{b}_t + \dot{\mathbf{b}}_t\tau + \ddot{\mathbf{b}}_t\frac{\tau^2}{2} + \dddot{\mathbf{b}}_t\frac{\tau^3}{6}\right).d\tau \\ &= \mathbf{b}_t\Delta + \dot{\mathbf{b}}_t\frac{\Delta^2}{2} + \ddot{\mathbf{b}}_t\frac{\Delta^3}{6} + \dddot{\mathbf{b}}_t\frac{\Delta^4}{24} \end{aligned} \tag{2}$$

by solution of the definite integrals in the second line. Integrated biomasses divided by time have meaning as the average biomass over that period.

Similarly, for an $n_w$-vector $\mathbf{g}_t$ of elements $G_{w,t}$ from Eq. 11 of Cotter 2026, a projected integral from $\mathbf{g}_t = \mathbf{0}$ at $t$ to $t+\Delta$ is

$$\int_0^\Delta \mathbf{g}_t^{\rightarrow 3}.d\tau = \dot{\mathbf{g}}_t\frac{\Delta^2}{2} + \ddot{\mathbf{g}}_t\frac{\Delta^3}{6} + \dddot{\mathbf{g}}_t\frac{\Delta^4}{24}. \tag{3}$$

Differentials needed to evaluate Equation (3) are calculated by ECOLPS using gain- and loss-rate matrices, one pair for each order of differentials. Foods are in rows, consumers in columns. If all components of the simulated ecosystem are wild, these matrices are $n_{\mathrm{W}} \times n_{\mathrm{W}}$. If there is a single non-living organics (NLO) component $d \in \mathrm{D}$ and fishery components $f \in \mathrm{F}$ as well, the matrices are $n_{\mathrm{W}\cup\mathrm{d}} \times n_{\mathrm{W}\cup\mathrm{F}\cup\mathrm{d}}$. Rows and columns are filled as follows.

The 1[st]-order gain-rate matrix $\dot{\mathbf{G}}$ is filled with zeroes, except as follows. Self-growth rates, $rB_a$, are stored along the autotrophic diagonal, and partial conversion rates, $gB_mB_g$, for food $m$ and consumer $g$ in row $m$ and column $g$, respectively. See Eq. (11) of Cotter 2026. Any fisheries columns present will store the partial landing rates, $gB_mE_f$, of each food. See Eq.(14) of Cotter 2026. If NLO is present, row $d$ holds partial conversion rates, $hB_dB_s$, of NLO by scavengers $s$; see RS term 5 of Eq. (15) of Cotter 2026. Column $d$ of row $d$ is zero, representing the absence of self gain by NLO. Column $d$ of wild rows holds row-summed rates of gain of NLO from particulate self-losses of $w$ and from any particulate harvests of $w$ dropped by consumers; these are RS terms 1 or $psB_w$, 2 or $(h-g)B_wB_h$ and 3 or $(h-g)B_wE_f$, respectively, of Eq. (15) of Cotter 2026. $p$ is the particulate proportion of self loss.

The 1[st]-order loss matrix $\dot{\mathbf{L}}$ is also zero except as follows. Self-loss rates, $sB_w$, are stored along the wild diagonal, and partial harvesting-rates for food $m$ and consumer $g$, in row $m$ and column $g$. See Eq. (10), (12) and (13) of Cotter 2026. Any fisheries columns present will store the partial catch rates of each food or unproductive harvest. See the last, harvesting term in Eq.(13) of Cotter 2026. If NLO is present, row-$d$ elements are the same as in row-$d$ of $\dot{\mathbf{G}}$ because $a_{d,s} = 1$; see NLO pathway (iv) in Cotter 2026. Column-$d$ elements are all zero except in row $d$ which holds self loss of NLO $sB_d$, RS term 4 in Eq. (15) of Cotter 2026.

The higher-order matrices, for example $\ddot{\mathbf{G}}$ and $\ddot{\mathbf{L}}$, are structured as for these 1[st]-order matrices. Formulas for higher differentials are set out in the next sub-section.

The projected 1[st]-order gain rates for use on the RS of Equation (3) are the appropriate elements of $\dot{\mathbf{G}}$. As a column vector,

$$\dot{\mathbf{g}} = \left(\mathbf{1}^T.\dot{\mathbf{G}}\right)^T \tag{4}$$

where $^T$ is transpose, and $\mathbf{1}$ is a unit column vector of length the same as the number of rows in $\dot{\mathbf{G}}$. $\ddot{\mathbf{g}}$ and $\dddot{\mathbf{g}}$ are found likewise.

Projected rates for the unassimilated fraction of partial harvests dropped as particulates to NLO come from off-diagonal elements of $(\dot{\mathbf{L}} - \dot{\mathbf{G}})$. A $d$-row, if present at the bottom of this matrix, is filled with zeros except in the $d$-column which holds $\mathcal{s}B_d$. Projected particulate self losses dropped to NLO can be calculated as $\mathbf{p}.\mathrm{diag}(\dot{\mathbf{L}} - \dot{\mathbf{G}})_{\mathrm{W}\cup\mathrm{D}}$ where $\mathbf{p}$ is the vector of particulate self-loss proportions $\mathcal{p}_{\mathrm{W}\cup\mathrm{D}}$ (Cotter 2026). Projected partial discarding rates of fisheries, if required, come from the fishery elements (which have no relevant diagonal):

$$\dot{\mathbf{d}}_{\mathrm{F}} = (\dot{\mathbf{L}} - \dot{\mathbf{G}})_{\mathrm{F}} \ . \tag{5}$$

Let $\dot{\mathbf{l}} = \dot{\mathbf{L}}.\mathbf{1}$ be the column vector of row sums of $\dot{\mathbf{L}}$ where $\mathbf{1}$ is conformable with columns in $\dot{\mathbf{L}}$. From Equation (10) of Cotter 2026, projected rates of change of biomass with time, (1), needed for higher differentials discussed below, are

$$\dot{\mathbf{b}} = \dot{\mathbf{g}} - \dot{\mathbf{l}} \tag{6}$$

Any fishery elements present in $\dot{\mathbf{g}}$ obtained with Equation (4) represent the rate of change of total landings of all food components of $f$, $\dot{\mathcal{G}}_f$; see Eq. (14) of Cotter 2026. Likewise, in $\dot{\mathbf{l}}$, they refer to total catches. Since catches $\geq$ landings, the element of $\dot{\mathbf{b}}$ for any $f$ will be negative total discarded-biomass rate (which has no role in formulas for higher differentials below). Equation (4) to (6) may also be written for 2nd- and 3rd-order differentials needed for the Taylor series above.

#### 1.2 GLV formulas for differentials

Equation (6) provides the $\dot{B}$ used below. Fishery effort is assumed constant over $\Delta_t$: $\dot{E}_t = 0$.

$$\ddot{\mathcal{G}}_{i,j} = \begin{cases} \mathcal{r}\dot{B}_j & \text{for } i = j, j \in \mathrm{A} \\ \mathcal{g}(\dot{B}_i B_j + B_i \dot{B}_j) & \text{for } i \in \mathrm{M}_j, j \in \mathrm{H} \\ \mathcal{g}\dot{B}_i E_j & \text{for } i \in \mathrm{M}_j, j \in \mathrm{F} \\ \mathcal{ps}\dot{B}_i + \sum_{i>\mathrm{H}} (\mathcal{h} - \mathcal{g})(\dot{B}_i B_h + B_i \dot{B}_h) & \\ \quad + \sum_{i>\mathrm{F}} (\mathcal{h} - \mathcal{g})\dot{B}_i E_f & \text{for } i \in \mathrm{W}, j = d \end{cases} \tag{7}$$

$$\ddot{\mathcal{L}}_{i,j} = \begin{cases} \mathcal{s}\dot{B}_j & \text{for } i = j, j \in \mathrm{W} \\ \mathcal{h}(\dot{B}_i B_j + B_i \dot{B}_j) & \text{for } i \in \mathrm{M}_j, j \in \mathrm{H} \\ \mathcal{h}\dot{B}_i E_j & \text{for } i \in \mathrm{M}_j, j \in \mathrm{F} \\ \mathcal{s}\dot{B}_{i,j} & \text{for } i = d, j = d \end{cases} \tag{8}$$

The 2nd-order matrix equation is solved for $\ddot{\mathbf{b}}$ as for the 1st-order equation in Equation (6). 3rd derivatives are

$$\dddot{\mathcal{G}}_{i,j} = \begin{cases} \mathcal{r}\ddot{B}_j & \text{for } i = j, j \in \mathrm{A} \\ \mathcal{g}\left\{\ddot{B}_i B_j + 2\dot{B}_i \dot{B}_j + B_i \ddot{B}_j\right\} & \text{for } i \in \mathrm{M}_j, j \in \mathrm{H} \\ \mathcal{g}\ddot{B}_i E_j & \text{for } i \in \mathrm{M}_j, j \in \mathrm{F} \\ \mathcal{ps}\ddot{B}_i + \sum_{i>\mathrm{H}} (\mathcal{h} - \mathcal{g})\left\{\ddot{B}_i B_h + 2\dot{B}_i \dot{B}_h + B_i \ddot{B}_h\right\} & \\ \quad + \sum_{i>\mathrm{F}} (\mathcal{h} - \mathcal{g})\ddot{B}_i E_f & \text{for } i \in \mathrm{W}, j = d \end{cases} \tag{9}$$

$$\dddot{\mathcal{L}}_{i,j} = \begin{cases} s\ddot{B}_j & \text{for } i = j, j \in \mathrm{W} \\ \hbar\Big\{\ddot{B}_i B_j + 2\dot{B}_i\dot{B}_j & \\ \quad +B_i\ddot{B}_j\Big\} & \text{for } i \in \mathrm{M}_j, j \in \mathrm{H} \\ \hbar\ddot{B}_i E_j & \text{for } i \in \mathrm{M}_j, j \in \mathrm{F} \\ s\ddot{B}_{i,j} & \text{for } i = d, j = d \end{cases} \tag{10}$$

The 3rd-order matrix equation for $\dddot{\mathbf{b}}$ is not needed unless unless projection orders higher than 3 are required.

### 2 Annex S2. Setting parameter values

#### 2.1 Using steady-state solutions

Using LV steady-state biomasses presented in Eq. (5) and (6) of Cotter 2026 to assist assignment of values to parameters was suggested in Cotter 2026. A 3-level food chain with, say, alga $a$, grazer $h$ eating $a$, and predator $p$ eating $h$, may be solved in the same way and applied to achieve diminishing biomasses going up that food chain if active constraints permit. Solving finds two solutions for $\tilde{B}_h$, and $\tilde{B}_a = f(\tilde{B}_p)$:

$$\tilde{B}_h = (r - s)_a \div h_{a,h} \qquad \text{from GLV for } a \tag{11}$$

$$\tilde{B}_h = s_p \div g_{h,p} \qquad \text{from GLV for } p \tag{12}$$

$$\tilde{B}_a = (h_{h,p}\tilde{B}_p + s_h) \div g_{a,h} \quad \text{from GLV for } h \tag{13}$$

$$\text{or } \tilde{B}_p = (g_{a,h}\tilde{B}_a - s_h) \div h_{h,p}. \tag{14}$$

For Equation (13) $>$ (11), set values so that

$$\frac{h_{h,p}\tilde{B}_p + s_h}{a_{a,h}} > (r - s)_a \quad \text{when } \tilde{B}_a > \tilde{B}_h \text{ is expected,} \tag{15}$$

and for (12) $>$ (14), set

$$\frac{s_p}{a_{h,p}} > g_{a,h}\tilde{B}_a - s_h \quad \text{when } \tilde{B}_h > \tilde{B}_p \text{ is expected.} \tag{16}$$

Possibly other multi-component systems could be usefully solved in this way.

#### 2.2 Using ranges for rate coefficients

These were little used but may be of interest. Suppose that values of time-dimensioned parameters, see Table 3 in Cotter 2026, are intended individually to cause no more than a halving or doubling of any biomass between adjacent simulated points for an LV model (with $a > h$ = '$a$ eaten by $h$') as an aid to the accuracy of projections. This idea can provide 1[st]-order ranges for the values of these positive parameters, given that they are constant for any non-negative values of biomasses including zeros.

With all $B_{h \in \mathrm{H}} = 0$, a value for the self-change coefficient of autotroph $a$ complies with

$$\begin{aligned} 0.5B_a &< B_a + \Delta(r - s)B_a < 2B_a, \\ \text{or, cancelling } B_a, \qquad 0 &< (r - s)_a < \Delta^{-1} \end{aligned} \tag{17}$$

because $0 < s_a < r_a$ if $a$ grows in the absence of any grazing.

With all $B_w = 0$ except $B_h$, and all $E_f = 0$, the self loss of heterotroph $h$ complies with

$$\begin{aligned} 0.5B_h &< B_h - \Delta s B_h \\ \text{or, cancelling } B_h, \qquad 0 &< s_h < 0.5\Delta^{-1} \end{aligned} \tag{18}$$

because all $s_h > 0$.

With all $B_w = 0$ except $B_{m\downarrow h}$ and $B_h$, and all $E_f = 0$, harvest coefficient $\mathcal{h}_{m,h}$ complies with

$$0.5B_h < B_h - \Delta \mathcal{s} B_h + \Delta \mathcal{a}\mathcal{h} B_m B_h < 2B_h,$$
$$\text{or, cancelling } B_h, \qquad \frac{\mathcal{s}_h - 0.5\Delta^{-1}}{\mathcal{a}_{m,h} B_m} < 0 < \mathcal{h}_{m,h} < \frac{\Delta^{-1} + \mathcal{s}_h}{\mathcal{a}_{m,h} B_m}. \tag{19}$$

The left side, being negative by Equation (18), can be deleted. So, noting $\mathcal{a} \leq 1$ (table 3 of Cotter 2026),

$$0 < \mathcal{h}_{m,h} < \frac{\Delta^{-1} + \mathcal{s}_h}{B_m}. \tag{20}$$

### 3 Annex S3. ECOLPS software

#### 3.1 Introduction

The theory in this paper was developed and tested using ECOLPS, a program written in R (R Core team, 2016), available with Trial.txt and Controls.txt files from [URL to be decided] . (ECOLPS was once known as LINSIM before another program of that name was found). ECOLPS is documented with set-up and introductory text, sectional texts, and line-by-line comments intended to assist usage, maintenance and adjustments. They add detail to this short description. There are two code files, ECOLPS.R and ECOLPSfunctions.R, the latter holding repeatedly used functions. Both must be stored in the same R folder for operation. ECOLPS is conveniently operated from a text editor called Tinn-R (https://tinn-r.org/en/download) because it highlights R commands, brackets and comments, and it allows direct submission of code or parts of it (for de-bugging) to R's graphic user interface (GUI). This is described in a READ.ME.txt file. An unsuccessful attempt to run ECOLPS from Rstudio was not pursued.

#### 3.2 Inputs

Two or three text files are read by ECOLPS. They have row and/or column headers. ECOLPS searches the row headers for specific, minimal text included in the row or column name, eg. 'Envir' for finding the row holding the Enviroment.file, and 'View.Env' for finding the switch to screen the Envir.file after processing it. Note that the search text is designed so that a search for one piece of text does not find another or both. With this input system, the user omits variables if they are not needed, may abbreviate variable names as long as they still include the searched-for text, and may order rows and columns freely in input files. Unused rows in input files may be temporarily removed with # at the beginning of the line. Unused columns can be temporarily removed by changing their column headers to NA. The three input file types are:

- Controls.txt contains the names of data files, and run-time options. Controls.txt must be stored in the same directory=folder as ECOLPS.R and ECOLPSfunctions.R. For naming and explanations of Control options, please refer to the ECOLPS.R file in section #1 of the introductory notes. The \ECOLPS\TRIALinputs\folder contains only one Controls.txt file. Lines in this must be adjusted *without changing the file name* for each of the reported trial simulations. Most of the different runs only require one or two characters to be changed in Controls.txt, eg. for the run number in the file name. YearLength and Years.sim must be changed when proceeding from simple LV models to others with years and seasons. Annex S4 provides hard copies of the controls and parameter files used for every trial and run if needed for reference.

- 'Params.file' holds a table of parameters(rows)$\times$components(columns). They can be named freely with a .txt ending, eg. Trial1.Run1.txt. ECOLPS categorizes each column of Params.file as a type of living or non-living component, see Table 1 of Cotter 2026, using positive values supplied for specific parameters; see section #3 of the code where the rules are shown as comments at the beginning. 0 or NA in a column of Params.file means that the parameter value (pv) in the row is not applicable to the component of that column. NAs are converted by ECOLPS to zeros for arithmetic purposes. So, if a zero value is required (unusual), it should be written as a very small positive value $\approx 0$ eg. 0.0000001, so that it has negligible influence on the simulation. ECOLPS sorts rows of parameters relating to components so that their ordering matches that of the given component columns. This is done to simplify and standardise indexing in the R code. As a result, the data matrices may acquire many zero values.

- An optional 'Envir.file' holds a table of dates (rows)$\times$updates (columns) permitting adjustments at specific times to parameter values or variables used in a simulation. Dates in this file are always measured in days from the start of the simulation, $1 \ldots 365 \ldots 730 \ldots$, etc.. This locates events within imaginary 'years' using the YearLength variable in

Controls.txt which supplies the number of $\Delta$s per 365 days. ECOLPS stacks the .envir file if necessary to cover all simulated years, thereby permitting seasonal or multi-annual cycles, and convenient extension of simulations one or more times using MainMenu options 8 or 9.

The formats of these files are described in prefacing text in ECOLPS.R, along with advice on downloading and running R and Tinn-R.

#### 3.3 Outputs

Simulated time series for variables, active constraints, slack variables and constraint limits, may be graphed, tabulated on screen, and analyzed for peaks and valleys if appropriate. Time series may be extended in length by appending to, or overwriting existing series. These facilities are helpful for identifying causes and effects in the series, and for continuation of series showing still-developing features.

At the end of each simulated block of time, ECOLPS presents the 'Main' menu. Selecting the 'Variables' sub-menu allows selection of simulated time series for graphing or printing on screen. Alternatively, the 'Constraints' sub-menu may be selected to show constraint properties over time. Three types of constraint property shown can be cycled through by repeatedly pressing 0:- (i) Active constraints, those that are that are active at any time during the simulation, have periods of activity shown by horizontal bars over time, each different constraint being shown at an arbitrary integer height up the ordinate axis. Constraints that were never active are omitted (but see option 6 of the Main Menu). (ii) Slack variables are shown as time series, with zero values indicating activity of the constraint. (iii) Constraint limits are time series of limits projected at time $t$; the size of the limits may be instructive about the effects of a constraint. Having viewed either simulated variables or constraints, pressing [Enter] without a menu option takes you back to the Main Menu. The selected output series are preserved across sub-menus, making it possible to show simulated variables and constraints together on the same time-series graph (though such presentations can easily become cluttered). Option C clears all stored time series; usually you use this when moving from one menu to another. Option R removes the current active graph. graphics.off() typed at the R> prompt clears all graphs.

When graphing or printing time series, obscuring series can be edited out with E, maximum ordinate values may be set for editing or comparative purposes with O, beginning (B) and finishing (F) times may be narrowed to improve clarity at selected times, and time-series points thinned to improve clarity with I (though sharp peaks and dips may be truncated by thinning). Very small and large values are best seen by printing the data (above the last visible menu on screen) using the same sub-setting facilities as for graphs. Titles, axis labels and keys may be altered, or simple text labels added, for reporting purposes.

Graphs may be saved from R as .jpegs or .pdfs, etc for adjustments and juxtaposing, eg. using panoramas in Irfanview (quick loading, free picture-editing and file-format changing software). Data printed on screen may be copied and pasted from R into a text editor, or read into a spreadsheet.

#### 3.4 Debugging and program maintenance

Variables calculated for the last projection step can be printed on screen for checking of internal calculations. Another option prints internal calculations for every step after a given date; the final step may be brought forward to shorten voluminous output. Alternatively, stop() or print(variable) statements may be temporarily written into the code, and then selected code can be sent separately to the R GUI to check the values at those places in the code. Variables remain in memory after a simulation is terminated and so may be printed on screen if the name of the variable in the code is looked up and typed in at the R prompt. Unfortunately this does not work for variables in written functions(); they can be viewed by placing print(variable) in the function code.

##### 3.5 Unpublished feature: totally unassimilated harvests

ECOLPS has code for features not described in Cotter 2026. Attention is drawn to its presence. The code has not been considered or tested as extensively as the code used for Trials.

Unassimilated harvesters, $u \in \mathrm{U} \subset \mathrm{G}$, are defined as those that kill wild components without assimilating any nourishment from them at all, ie. $a_{w,u} = 0$. As an example, demersal trawlers may lift large quantities of epibenthos of no market value and land none of it. Unassimilated harvests are UAH or ${}_{o}\mathcal{H}_{w,u}$. UAH relevant for simulations would typically arise from fisheries. UAH cannot be calculated like other harvests during LP as $\mathcal{G}^{*}/a$ because $a_{w,u} = 0$. Instead, a projection is used, as for self losses Eq.(22) in Cotter 2026. For example, to first order projection:

$$
{}_{o}\mathcal{H}^{\rightarrow 1}_{w,g} = \begin{cases} \frac{1}{2}\Delta^2 h B_{w,t} B_{h,t} & \text{for } h \in \mathrm{H},\ a_{w,h} = 0 \\ \frac{1}{2}\Delta^2 h B_{w,t} E_{f,t} & \text{for } f \in \mathrm{F},\ a_{w,f} = 0 \end{cases} \tag{21}
$$

An additional term for UAH dropped to NLO is needed on the RS of Eq.(15) of Cotter 2026 is $+\sum_{u \in \mathrm{G}_w} \big( h B_w B_{u \in \mathrm{H}} + h B_w E_{u \in \mathrm{F}} \big) \Big\}$. Modifications of constraints needed for UAH are listed in Table 1.

Table 1: Additional linear constraints on GP ($\mathcal{G}^{o}_{w \in \mathrm{W}}$ and $\mathcal{G}^{o}_{f \in \mathrm{F}}$) over $\Delta_t$ needed for LP simulations with unassimilated harvests

| Name [code] | Const-rained | Process affec-ted | Left side | Right-side limit | Sources for formulas | Number of constraints | Slack variable measures:- |
|---|---|---|---|---|---|---|---|
| Living food† [m.FOOD.G] | $g \in \mathrm{G}_{\mathrm{M} \cap \mathrm{W}}$ | harvest of $m \in \mathrm{M}$ | $\sum_{m > \mathrm{G}} \frac{\mathcal{G}^{o}_{m,g,t}}{a_{m,g}} - \mathcal{G}^{o}_{m,t}$ | $\leq B_{m,t} - \mathcal{S}^{\rightarrow 1}_{m,t} - \sum_{m > \mathrm{U} \subset \mathrm{G}} {}_{o}\mathcal{H}^{\rightarrow 1}_{m,u,t}$ | Eq. (10,12,22) of Cotter 2026 | $n_{\mathrm{M}}$ | $B_{m,t+\Delta}$ |
| Dummy food† [w.FOOD] | None | None | $-\mathcal{G}^{o}_{w,t}$ | $\leq B_{w,t} - \mathcal{S}^{\rightarrow 1}_{w,t} - \sum_{w > \mathrm{U} \subset \mathrm{G}} {}_{o}\mathcal{H}^{\rightarrow 1}_{w,u,t}$ | Eq. (10) of Cotter 2026 | $n_{w \notin \mathrm{M}}$ | $B_{w,t+\Delta}$ for $w \notin \mathrm{M}$ |
| Non-living food [d.FOOD.S] | $s \in \mathrm{S}_d$ | harvest of $d \in \mathrm{D}$ | $\sum_{d > \mathrm{S}} \frac{\mathcal{G}^{o}_{d,s,t}}{a_{d,s}} - \sum_{\mathrm{M}} \sum_{m > \mathrm{G}} (a^{-1}_{m,g} - 1)\, \mathcal{G}^{o}_{m,g,t}$ | $\leq B_{d,t} - \mathcal{S}^{\rightarrow 1}_{d,t} + \sum_{w \in \mathrm{W}} p_w \mathcal{S}^{\rightarrow 1}_{w,t} + \sum_{\mathrm{W}} \sum_{w > \mathrm{U} \subset \mathrm{G}} {}_{o}\mathcal{H}^{\rightarrow 1}_{w,u,t}$ | Eq.21 &(12,13,22) of Cotter 2026 | $n_d$ | $B_{d,t+\Delta}$ |
| Habitat [z.HAB.W] | $w \in \mathrm{W}_z$ | RP of $w$ need-ing $z$ | $\sum_{w \in \mathrm{W}_z} v_{z,w} \Big\{ \mathcal{G}^{o}_{w,t} - \sum_{w > \mathrm{G}} \frac{\mathcal{G}^{o}_{w,g,t}}{a_{w,g}} \Big\}$ | $\leq A_{z,t} + \sum_{w \in \mathrm{W}_z} v_{z,w} \mathcal{S}^{\rightarrow 1}_{w,t} + \sum_{w > \mathrm{U}} \sum_{w \in \mathrm{W}_z} v_{z,w}\, {}_{o}\mathcal{H}^{\rightarrow 1}_{w,u,t}$ | Eq. (21) of Cotter 2026 | $n_{\mathrm{Z}}$ | $A_{z,t+\Delta}$ |

### 4 Annex S4. Trial simulations

This annex presents ECOLPS inputs and outputs not included in the main texts of Cotter 2026 for reasons of space. Parameter files are copies of .txt files downloadable from Zenodo at https://tinyurl.com/ECOLPS.

#### 4.1 Trial 1. Basic LV example

##### 4.1.1 Run (i)

Demonstration with Lotka-Volterra example. See Tables 2 and 3.

Table 2: Left: Trial1.Run1.txt . Right: Controls.txt

| Parameter | Alga | Grazer | Control | Value | Text |
|---|---|---|---|---|---|
| SelfLoss | 0.25 | 0.25 | AdviceLevel | 2 | NA |
| SelfGrow | 0.75 | 0 | YearLength | 1 | NA |
| Harvest.Alga | 0 | 0.4 | Years.sim | 1000 | NA |
| Assim.Alga | 0 | 0.2 | proj.SLorder | 3 | NA |
| Bmass.init | 3 | 3 | View.params | 1 | NA |
| Gmax | NA | 3 | View.comps | 1 | NA |
| | | | View.controls | 0 | NA |
| | | | KeyItemWidth | 25 | NA |
| | | | KeyLineLength | 90 | NA |
| | | | WinHeight | 5 | NA |
| | | | WinWidth | 9.5 | NA |
| | | | Progress.rep | 500 | NA |
| | | | Parameters.file | NA | C:\\ECOLPS\TRIALinputs\ Trial1.Run1.txt |

Table 3 shows the existence of stationary points for $B_{h,t}$ and $B_{a,t}$ at peak biomasses of the other component at the start and end of the LV series. 'Theory' is from Eq. (5), (6) of Cotter 2026.

Table 3: Trial 1(i). $B_{a,t}$ and $B_{h,t}$ found at peak biomasses of the other component near the start and end of simulation over 1000Δs.

| Time, Δs | Peak $B_a$ | $B_{h,t}$ | | Time, Δs | Peak $B_h$ | $B_{a,t}$ |
|---|---|---|---|---|---|---|
| 34 | 8.95 | 1.23 | | 40 | 2.62 | 3.14 |
| 72 | 7.63 | 1.17 | | 79 | 2.33 | 2.94 |
| 110 | 6.62 | 1.30 | | 116 | 2.11 | 3.36 |
| . | . | . | | . | . | . |
| 895 | 3.17 | 1.25 | | 904 | 1.26 | 3.12 |
| 930 | 3.16 | 1.25 | | 939 | 1.26 | 3.13 |
| 966 | 3.15 | 1.25 | | 975 | 1.26 | 3.12 |
| Theory | - | 1.25 | - | - | - | 3.125 |

##### 4.1.2 Run (ii)

Basic Lotka-Volterra example starting at steady-state biomasses found from Run (i). See Table 4.

Table 4: Left: Trial1.Run2.txt . Right: Controls.txt

| Parameter | Alga | Grazer | Control | Value | Text |
|---|---|---|---|---|---|
| SelfLoss | 0.25 | 0.25 | AdviceLevel | 2 | NA |
| SelfGrow | 0.75 | 0 | YearLength | 1 | NA |
| Harvest.Alga | 0 | 0.4 | Years.sim | 1000 | NA |
| Assim.Alga | 0 | 0.2 | proj.SLorder | 3 | NA |
| Bmass.init | 3.125 | 1.25 | Parameters.file | NA | C:\\ECOLPS\TRIALinputs \Trial1.Run2.txt |
| Gmax | NA | 3 | | | |

###### 4.1.3 Run (iii)

Basic Lotka-Volterra example with time-dependent parameters halved. See Table 5.

Table 5: Left: Trial1.Run3.txt . Right: Controls.txt

| Parameter | Alga | Grazer | Control | Value | Text |
|---|---|---|---|---|---|
| SelfLoss | 0.125 | 0.125 | AdviceLevel | 2 | NA |
| SelfGrow | 0.375 | 0 | YearLength | 1 | NA |
| Harvest.Alga | 0 | 0.2 | Years.sim | 1000 | NA |
| Assim.Alga | 0 | 0.2 | proj.SLorder | 3 | NA |
| Bmass.init | 3 | 3 | Parameters.file | NA | C:\\ECOLPS\TRIALinputs\ Trial1.Run3.txt |
| Gmax | NA | 3 | | | |

###### 4.1.4 Run (iv)

Basic Lotka-Volterra example with Alga's SelfGrow set variably from {0.4, 0.5, 0.62, 0.75, 0.87, 1.0, 1.2}. See * in Table 6.

Table 6: Left: Trial1.Run4.txt . Right: Controls.txt

| Parameter | Alga | Grazer | Control | Value | Text |
|---|---|---|---|---|---|
| SelfLoss | 0.25 | 0.25 | YearLength | 1 | NA |
| SelfGrow | 0.4* | 0 | Years.sim | 3000 | NA |
| Harvest.Alga | 0 | 0.4 | proj.SLorder | 3 | NA |
| Assim.Alga | 0 | 0.2 | Parameters.file | NA | C:\\ECOLPS\TRIALinputs\ Trial1.Run4.txt |
| Bmass.init | 3 | 3 | | | |
| Gmax | NA | 3 | | | |

Code in R, Trial1.CompareLVsim2Theor.R , used to prepare Fig. 4 in Cotter 2026 can be downloaded from Zenodo at https://tinyurl.com/LINSIMsimulator/TRIALinputs/ .

#### 4.2 Trial 2. Grazer satiation in LV cycles

##### 4.2.1 Run (i)

This is a re-run of Trial 1(i) with a.SATE.h activated by reducing g.max from 3. See * in Table 7

Table 7: Left: Trial2.Run1.txt . Right: Controls.txt

| Parameter | Alga | Grazer | Control | Value | Text |
|---|---|---|---|---|---|
| SelfLoss | 0.25 | 0.25 | YearLength | 1 | NA |
| SelfGrow | 0.75 | 0 | Years.sim | 1000 | NA |
| Harvest.Alga | 0 | 0.4 | proj.SLorder | 3 | NA |
| Assim.Alga | 0 | 0.2 | Parameters.file | NA | C:\\ECOLPS\TRIALinputs\ Trial2.Run1.txt |
| Bmass.init | 3 | 3 | | | |
| Gmax | NA | 0.3552* | | | |

##### 4.2.2 Run (ii)

Trial 1(i) is re-run again with a.SATE.h activated marginally by g.max. See * in Table 8.

Table 8: Left: Trial2.Run2.txt . Right: Controls.txt

| Parameter | Alga | Grazer | Control | Value | Text |
|---|---|---|---|---|---|
| SelfLoss | 0.25 | 0.25 | YearLength | 1 | NA |
| SelfGrow | 0.75 | 0 | Years.sim | 1000 | NA |
| Harvest.Alga | 0 | 0.4 | proj.SLorder | 3 | NA |
| Assim.Alga | 0 | 0.2 | Parameters.file | NA | C:\\ECOLPS\TRIALinputs\ Trial2.Run2.txt |
| Bmass.init | 3 | 3 | | | |
| Gmax | NA | 0.3551* | | | |

#### 4.3 Trial 3. One LV component duplicated (algae)

##### 4.3.1 Run (i)

Trial 1(i) is re-run with identical alga1 and alga2. See Table 9.

Table 9: Left: Trial3.Run1.txt . Right: Controls.txt

| Parameter | Alga1 | Alga2 | Grazer | Control | Value | Text |
|---|---|---|---|---|---|---|
| SelfLoss | 0.25 | 0.25 | 0.25 | YearLength | 1 | NA |
| SelfGrow | 0.75 | 0.75 | 0 | Years.sim | 1000 | NA |
| Harvest.Alga1 | 0 | 0 | 0.4 | proj.SLorder | 3 | NA |
| Assim.Alga2 | 0 | 0 | 0.2 | Parameters.file | NA | C:\\ECOLPS\ TRIALinputs\Trial3.Run1.txt |
| Harvest.Alga1 | 0 | 0 | 0.4 | | | |
| Assim.Alga2 | 0 | 0 | 0.2 | | | |
| Bmass.init | 3 | 3 | 3 | | | |
| Gmax | NA | NA | 3 | | | |

##### 4.3.2 Run (ii)

Trial 3(i) is re-run with alga1 less competitive than alga2. See * in Table 10.

Table 10: Left: Trial3.Run2.txt . Right: Controls.txt

| Parameter | Alga1 | Alga2 | Grazer | Control | Value | Text |
|---|---|---|---|---|---|---|
| SelfLoss | 0.25 | 0.25 | 0.25 | YearLength | 1 | NA |
| SelfGrow | 0.70* | 0.75 | 0 | Years.sim | 1000 | NA |
| Harvest.Alga1 | 0 | 0 | 0.4 | proj.SLorder | 3 | NA |
| Assim.Alga1 | 0 | 0 | 0.2 | Parameters.file | NA | C:\\ECOLPS\ TRIALinputs\Trial3.Run2.txt |
| Harvest.Alga2 | 0 | 0 | 0.4 | | | |
| Assim.Alga2 | 0 | 0 | 0.2 | | | |
| Bmass.init | 3 | 3 | 3 | | | |
| Gmax | NA | NA | 3 | | | |

#### 4.4 Trial 4. One LV component duplicated (grazers)

##### 4.4.1 Run (i)

Trial 1(i) is re-run with identical grazer1 and grazer2. See Table 11.

.

Table 11: Left: Trial4.Run1.txt . Right: Controls.txt

| Parameter | Alga | Grazer1 | Grazer2 | Control | Value | Text |
|---|---|---|---|---|---|---|
| SelfLoss | 0.25 | 0.25 | 0.25 | YearLength | 1 | NA |
| SelfGrow | 0.75 | 0 | 0 | Years.sim | 1000 | NA |
| Harvest.Alga | 0 | 0.4 | 0.4 | proj.SLorder | 3 | NA |
| Assim.Alga | 0 | 0.2 | 0.2 | Parameters.file | NA | C:\\ECOLPS\TRIALinputs\ Trial4.Run1.txt |
| Bmass.init | 3 | 3 | 3 | | | |
| Gmax | NA | 3 | 3 | | | |

###### 4.4.2 Run (ii)

Trial 4(i) is re-run with grazer2 less effective than grazer1. See * in Table 12

Table 12: Left: Trial4.Run2.txt . Right: Controls.txt

| Parameter | Alga | Grazer1 | Grazer2 | Control | Value | Text |
|---|---|---|---|---|---|---|
| SelfLoss | 0.25 | 0.25 | 0.25 | YearLength | 1 | NA |
| SelfGrow | 0.75 | 0 | 0 | Years.sim | 1000 | NA |
| Harvest.Alga | 0 | 0.4 | 0.35* | proj.SLorder | 3 | NA |
| Assim.Alga | 0 | 0.2 | 0.2 | Parameters.file | NA | C:\\ ECOLPS\TRIALinputs\Trial4.Run2txt |
| Bmass.init | 3 | 3 | 3 | | | |
| Gmax | NA | 3 | 3 | | | |

#### 4.5 Trial 5. Both LV components duplicated

###### 4.5.1 Run (i)

Trial 1(i) is re-run with an extra identical alga, and an extra identical grazer. See Table 13

Table 13: Left: Trial5.Run1.txt . Right: Controls.txt

| Parameter | Alga1 | Alga2 | Grazer1 | Grazer2 | Control | Value | Text |
|---|---|---|---|---|---|---|---|
| SelfLoss | 0.25 | 0.25 | 0.25 | 0.25 | YearLength | 1 | NA |
| SelfGrow | 0.75 | 0.75 | 0 | 0 | Years.sim | 200 | NA |
| Harvest.Alga1 | 0 | 0 | 0.4 | 0.4 | proj.SLorder | 3 | NA |
| Assim.Alga1 | 0 | 0 | 0.2 | 0.2 | Parameters.file | NA | C:\\ECOLPS\TRIALinputs\Trial5.Run1.txt |
| Harvest.Alga2 | 0 | 0 | 0.4 | 0.4 | | | |
| Assim.Alga2 | 0 | 0 | 0.2 | 0.2 | | | |
| Harvest.Grazer1 | 0 | 0 | 0 | 0 | | | |
| Assim.Grazer1 | 0 | 0 | 0 | 0 | | | |
| Bmass.init | 3 | 3 | 3 | 3 | | | |
| Gmax | NA | NA | 3 | 3 | | | |

##### 4.5.2 Run (ii)

Trial 5(i) was re-run twice with grazer2 the less, or more effective grazer. See * in Table 14

Table 14: Left: Trial5.Run2.txt . Right: Controls.txt

| Parameter | Alga1 | Alga2 | Grazer1 | Grazer2 | Control | Value | Text |
|---|---|---|---|---|---|---|---|
| SelfLoss | 0.25 | 0.25 | 0.25 | 0.25 | YearLength | 1 | NA |
| SelfGrow | 0.75 | 0.75 | 0 | 0 | Years.sim | 200 | NA |
| Harvest.Alga1 | 0 | 0 | 0.4 | 0.4 | proj.SLorder | 3 | NA |
| Assim.Alga1 | 0 | 0 | 0.2 | 0.2 | Parameters.file | NA | C:\\ECOLPS\TRIALinputs\Trial5.Run2.txt |
| Harvest.Alga2 | 0 | 0 | 0.4 | 0.35 or 0.45* | | | |
| Assim.Alga2 | 0 | 0 | 0.2 | 0.2 | | | |
| Harvest.Grazer1 | 0 | 0 | 0 | 0 | | | |
| Assim.Grazer1 | 0 | 0 | 0 | 0 | | | |
| Bmass.init | 3 | 3 | 3 | 3 | | | |
| Gmax | NA | NA | 3 | 3 | | | |

##### 4.5.3 Run (iii)

Trial 5(i) was re-run with grazer2 now eating grazer1 at a low rate for comparison with Run (i). See * in Table 15.

Table 15: Left: Trial5.Run3.txt . Right: Controls.txt

| Parameter | Alga1 | Alga2 | Grazer1 | Grazer2 | Control | Value | Text |
|---|---|---|---|---|---|---|---|
| SelfLoss | 0.25 | 0.25 | 0.25 | 0.25 | YearLength | 1 | NA |
| SelfGrow | 0.75 | 0.75 | 0 | 0 | Years.sim | 200 | NA |
| Harvest.Alga1 | 0 | 0 | 0.4 | 0.4 | proj.SLorder | 3 | NA |
| Assim.Alga1 | 0 | 0 | 0.2 | 0.2 | Parameters.file | NA | C:\\ECOLPS\TRIALinputs\Trial5.Run3.txt |
| Harvest.Alga2 | 0 | 0 | 0.4 | 0.4 | | | |
| Assim.Alga2 | 0 | 0 | 0.2 | 0.2 | | | |
| Harvest.Grazer1 | 0 | 0 | 0 | 0.05* | | | |
| Assim.Grazer1 | 0 | 0 | 0 | 0.2* | | | |
| Bmass.init | 3 | 3 | 3 | 3 | | | |
| Gmax | NA | NA | 3 | 3 | | | |

#### 4.6 Trial 6. 3-level food chain: alga, grazer, and predator eating grazer

##### 4.6.1 Run (i)

Parameter values complied with steady-state Eq. (27)–(30) in Cotter 2026. See Table 16 and **??**.

Table 16: Left: Trial6.Run1.txt . Right: Controls.txt

| Parameter | Alga | Grazer | Predator | Control | Value | Text |
|---|---|---|---|---|---|---|
| SelfLoss | 0.25 | 0.25 | 0.1 | YearLength | 1 | NA |
| SelfGrow | 0.75 | 0 | 0 | Years.sim | 3000 | NA |
| Harvest.Alga | 0 | 0.4 | 0 | proj.SLorder | 3 | NA |
| Assim.Alga | 0 | 0.2 | 0 | Parameters.file | NA | C:\\ECOLPS\TRIALinputs\Trial6.Run1.txt |
| Harvest.Grazer | 0 | 0 | 0.5333 | | | |
| Assim.Grazer | 0 | 0 | 0.15 | | | |
| Bmass.init | 3 | 3 | 0.3 | | | |
| Gmax | NA | 3 | 3 | | | |

##### 4.6.2 Run (ii)

Trial 6(i) was re-run with non-compliant $\hbar_{a,h} = 0.5$. See * in Table 17 and **??**.

Table 17: Left: Trial6.Run2.txt . Right: Controls.txt

| Parameter | Alga | Grazer | Predator | Control | Value | Text |
|---|---|---|---|---|---|---|
| SelfLoss | 0.25 | 0.25 | 0.1 | YearLength | 1 | NA |
| SelfGrow | 0.75 | 0 | 0 | Years.sim | 1000 | NA |
| Harvest.Alga | 0 | 0.5* | 0 | proj.SLorder | 3 | NA |
| Assim.Alga | 0 | 0.2 | 0 | Parameters.file | NA | C:\\ECOLPS\TRIALinputs\Trial6.Run2.txt |
| Harvest.Grazer | 0 | 0 | 0.5333 | | | |
| Assim.Grazer | 0 | 0 | 0.15 | | | |
| Bmass.init | 3 | 3 | 0.3 | | | |
| Gmax | NA | 3 | 3 | | | |

##### 4.6.3 Run (iii)

Run 6(i) was re-run with non-compliant $\hbar_{a,h} = 0.3$. See * in Table 18.

Table 18: Left: Trial6.Run3.txt . Right: Controls.txt

| Parameter | Alga | Grazer | Predator | Control | Value | Text |
|---|---|---|---|---|---|---|
| SelfLoss | 0.25 | 0.25 | 0.1 | YearLength | 1 | NA |
| SelfGrow | 0.75 | 0 | 0 | Years.sim | 1000 | NA |
| Harvest.Alga | 0 | 0.3* | 0 | proj.SLorder | 3 | NA |
| Assim.Alga | 0 | 0.2 | 0 | Parameters.file | NA | C:\\ECOLPS\TRIALinputs\Trial6.Run3.txt |
| Harvest.Grazer | 0 | 0 | 0.5333 | | | |
| Assim.Grazer | 0 | 0 | 0.15 | | | |
| Bmass.init | 3 | 3 | 0.3 | | | |
| Gmax | NA | 3 | 3 | | | |

#### 4.7 Trial 7. Effects of constrained essential habitat on an LV system

##### 4.7.1 Run (i)

Trial 1(i) re-run with Habitat, Hab1, constraining retained GP of alga $a$. See Table 19.

Table 19: Left: Trial7.Run1.txt . Right: Controls.txt

| Parameter | Alga | Grazer | Hab1 | Control | Value | Text |
|---|---|---|---|---|---|---|
| SelfLoss | 0.25 | 0.25 | NA | YearLength | 1 | NA |
| SelfGrow | 0.75 | 0 | NA | Years.sim | 200 | NA |
| Harvest.Alga | 0 | 0.4 | NA | proj.SLorder | 3 | NA |
| Assim.Alga | 0 | 0.2 | NA | Parameters.file | NA | C:\\ECOLPS\TRIALinputs\Trial7.Run1.txt |
| Bmass.init | 3 | 3 | NA | | | |
| Gmax | NA | 3 | NA | | | |
| Occup.Hab1 | 1 | 0 | NA | | | |
| HabSize | NA | NA | 3.05 | | | |

###### 4.7.2 Run (ii)

Trial 1(i) re-run with habitat, Hab2, constraining rGP of grazer $h$. Note: Grazer biomass reduced from 3 for initial compliance with z.HAB.h. See * in Table 20.

Table 20: Left: Trial7.Run2.txt . Right: Controls.txt

| Parameter | Alga | Grazer | Hab2 | Control | Value | Text |
|---|---|---|---|---|---|---|
| SelfLoss | 0.25 | 0.25 | NA | YearLength | 1 | NA |
| SelfGrow | 0.75 | 0 | NA | Years.sim | 200 | NA |
| Harvest.Alga | 0 | 0.4 | NA | proj.SLorder | 3 | NA |
| Assim.Alga | 0 | 0.2 | NA | Parameters.file | NA | C:\\ECOLPS\TRIALinputs\Trial7.Run2.txt |
| Bmass.init | 3 | 1* | NA | | | |
| Gmax | NA | 3 | NA | | | |
| Occup.Hab2 | 0 | 1 | NA | | | |
| HabSize | NA | NA | 1.24 | | | |

##### 4.8 Trial 8. LV system with unreplenished essential nutrient $n$

###### 4.8.1 Run (i)

Trial 1(i) re-run with an initial supply of $n$ for photosynthesis of alga $a$. See * in Table 21. Trial 8 had only one run.

Table 21: Left: Trial8.Run1.txt . Right: Controls.txt

| Parameter | Alga | Grazer | H2O | Control | Value | Text |
|---|---|---|---|---|---|---|
| SelfLoss | 0.25 | 0.25 | NA | YearLength | 1 | NA |
| SelfGrow | 0.75 | 0 | NA | Years.sim | 1000 | NA |
| Harvest.Alga | 0 | 0.4 | NA | proj.SLorder | 3 | NA |
| Assim.Alga | 0 | 0.2 | NA | Parameters.file | NA | C:\\ECOLPS\TRIALinputs\Trial8.Run[illegible] |
| Bmass.init | 3 | 3 | NA | | | |
| Gmax | NA | 3 | NA | | | |
| Nconc.$n$ | 0.0068 | 0.002 | NA | | | |
| Nmass.$n$ | NA | NA | 3* | | | |
| ParticSL | 0.08 | 0.08 | NA | | | |

##### 4.9 Trial 9. 3-level food chain with annual inputs of essential nutrient N1

Trial9.Envir.txt is needed for all 3 runs. Inputs of 0.7$\gamma$ (mass units) of N1 occur on day 1 of each year of 12$\Delta$s. See Table 22.

Table 22: Trial9.Envir.txt .

| Date.day | Variable | Value | Component | TransferTo |
|---|---|---|---|---|
| 1 | Nmass.N1 | 0.7 | H2O | NA |

##### 4.9.1 Run (i)

Low initial Nmass.N1 was set to signal nutrient presence to ECOLPS. See * in Table 23.

Table 23: Left: Trial9.Run1.txt . Right: Controls.txt

| Parameter | Alga | Graz | Pred | H2O | Control | Value | Text |
|---|---|---|---|---|---|---|---|
| SelfLoss | 0.15 | 0.25 | 0.2 | NA | YearLength | 12 | NA |
| SelfGrow | 1.1 | 0 | 0 | NA | Years.sim | 8 | NA |
| Harvest.Alga | 0 | 1.05 | 0 | NA | proj.SLorder | 3 | NA |
| Assim.Alga | 0 | 0.4 | 0 | NA | Parameters.file | NA | C:\\ECOLPS\TRIALinputs\Trial9.Run1.txt |
| Harvest.Graz | 0 | 0 | 0.9 | NA | Envir.file | NA | C:\\ECOLPS\TRIALinputs\Trial9.Envir.txt |
| Assim.Graz | 0 | 0 | 0.6 | NA | | | |
| Harv.Pred | 0 | 0 | 0 | NA | | | |
| Assim.Pred | 0 | 0 | 0 | NA | | | |
| Bmass.init | 1.5 | 0.5 | 0.15 | NA | | | |
| Nconc.N1 | 0.23 | 0.06 | 0.01 | NA | | | |
| Nmass.N1 | NA | NA | NA | 0.001* | | | |
| ParticSL | 0.33 | 0.17 | 0.05 | NA | | | |

##### 4.9.2 Run (ii)

Fishery, F1 was added to Trial 9(i), catching Pred, not Graz. See * in Table 24.

Table 24: Left: Trial9.Run2.txt . Right: Controls.txt

| Parameter | Alga | Graz | Pred | H2O | F1 | Control | Value | Text |
|---|---|---|---|---|---|---|---|---|
| SelfLoss | 0.15 | 0.25 | 0.2 | NA | NA | YearLength | 12 | NA |
| SelfGrow | 1.1 | 0 | 0 | NA | 0 | Years.sim | 8 | NA |
| Harvest.Alga | 0 | 1.05 | 0 | NA | NA | proj.SLorder | 3 | NA |
| Assim.Alga | 0 | 0.4 | 0 | NA | 0 | Parameters.file | NA | C:\\ECOLPS\TRIALinputs\Trial9.Run2.txt |
| Harvest.Graz | 0 | 0 | 0.9 | NA | 0 | Envir.file | NA | C:\\ECOLPS\TRIALinputs\Trial9.Envir.txt |
| Assim.Graz | 0 | 0 | 0.6 | NA | 0 | | | |
| Harv.Pred | 0 | 0 | 0 | NA | 0.5* | | | |
| Assim.Pred | 0 | 0 | 0 | NA | 0.7 | | | |
| Bmass.init | 1.5 | 0.5 | 0.15 | NA | NA | | | |
| Nconc.N1 | 0.23 | 0.06 | 0.01 | NA | NA | | | |
| Nmass.N1 | NA | NA | NA | 0.001 | NA | | | |
| ParticSL | 0.33 | 0.17 | 0.05 | NA | NA | | | |
| F.effort | NA | NA | NA | NA | 1 | | | |

##### 4.9.3 Run (iii)

Fishery, F2 was added to Trial 9(i), catching Graz, not Pred. See * in Table 25.

Table 25: Left: Trial9.Run3.txt . Right: Controls.txt

| Parameter | Alga | Graz | Pred | H2O | F2 | Control | Value | Text |
|---|---|---|---|---|---|---|---|---|
| SelfLoss | 0.15 | 0.25 | 0.2 | NA | NA | YearLength | 12 | NA |
| SelfGrow | 1.1 | 0 | 0 | NA | 0 | Years.sim | 8 | NA |
| Harvest.Alga | 0 | 1.05 | 0 | NA | NA | proj.SLorder | 3 | NA |
| Assim.Alga | 0 | 0.4 | 0 | NA | 0 | Parameters.file | NA | C:\\ECOLPS\TRIALinputs\Trial9.Run3.txt |
| Harvest.Graz | 0 | 0 | 0.9 | NA | 0.5* | Envir.file | NA | C:\\ECOLPS\TRIALinputs\Trial9.Envir.txt |
| Assim.Graz | 0 | 0 | 0.6 | NA | 0.7 | | | |
| Harv.Pred | 0 | 0 | 0 | NA | 0 | | | |
| Assim.Pred | 0 | 0 | 0 | NA | 0 | | | |
| Bmass.init | 1.5 | 0.5 | 0.15 | NA | NA | | | |
| Nconc.N1 | 0.23 | 0.06 | 0.01 | NA | NA | | | |
| Nmass.N1 | NA | NA | NA | 0.001 | NA | | | |
| ParticSL | 0.33 | 0.17 | 0.05 | NA | NA | | | |
| F.effort | NA | NA | NA | NA | 1 | | | |

#### 4.10 Trial 10. 3-level food chain re-cycling variable mass of essential nutrient N1

This trial repeats Trial 9 Run(i) without annual inputs of essential nutrient N1 (so there is no Trial10.Envir.txt). Instead, N1 is recycled via NLO $d$ and scavenger $s$. See Cotter 2026. Initial N1 masses in H2O and NLO were set equally to match the single annual input of N1 in Trial 9; see * in Table 26.

Table 26: Left: Trial10.Run1.txt . Right: Controls.txt

| Parameter | Alga | Graz | Pred | Scav | H2O | NLO | Control | Value | Text |
|---|---|---|---|---|---|---|---|---|---|
| SelfLoss | 0.15 | 0.25 | 0.2 | 0.3045 | NA | 0.8 | YearLength | 12 | NA |
| SelfGrow | 1.1 | 0 | 0 | 0 | NA | NA | Years.sim | 8 or 16 | NA |
| Harvest.Alga | 0 | 1.05 | 0 | 0 | NA | NA | proj.SLorder | 3 | NA |
| Assim.Alga | 0 | 0.4 | 0 | 0 | NA | NA | Parameters.file | NA | C:\\ECOLPS\TRIALinputs\Trial10.Run1. |
| Harvest.Graz | 0 | 0 | 0.9 | 0 | NA | NA | | | |
| Assim.Graz | 0 | 0 | 0.6 | 0 | NA | NA | | | |
| Harv.NLO | 0 | 0 | 0 | 0.176 | NA | NA | | | |
| Assim.NLO | 0 | 0 | 0 | 1 | NA | NA | | | |
| Bmass.init | 1.5 | 0.5 | 0.15 | 0.7 | NA | 1 | | | |
| Nconc.N1 | 0.23 | 0.06 | 0.01 | 0.03 | NA | NA | | | |
| Nmass.N1 | NA | NA | NA | NA | 0.35* | 0.35* | | | |
| ParticSL | 0.33 | 0.17 | 0.05 | 0.05 | NA | 0 | | | |

#### 4.11 Trial 11. 3-level food chain re-cycling of N1, seasonal light cycles, and fisheries

This trial develops Trial 10 with inclusion of a seasonal, approximately sinusoidal light cycle from Trial11.Envir.txt in all 3 runs; See Table 27.

Table 27: Trial11.Envir.txt .

| Date.day | Variable | Value | Component | TransferTo |
|---|---|---|---|---|
| 10 | L.index | 0.149 | H2O | NA |
| 40 | L.index | 0.260 | H2O | NA |
| 71 | L.index | 0.441 | H2O | NA |
| 101 | L.index | 0.631 | H2O | NA |
| 132 | L.index | 0.792 | H2O | NA |
| 162 | L.index | 0.870 | H2O | NA |
| 192 | L.index | 0.852 | H2O | NA |
| 223 | L.index | 0.737 | H2O | NA |
| 253 | L.index | 0.563 | H2O | NA |
| 284 | L.index | 0.366 | H2O | NA |
| 314 | L.index | 0.210 | H2O | NA |
| 344 | L.index | 0.131 | H2O | NA |

##### 4.11.1 Run (i)

No fishery. L.index and alphaSG.L were applied to pvs initially from Trial 10(i), Table 26, in order to vary self growth of Alga with an annual cycle of light. Some pvs had to be adjusted to obtain level $Q_{\Sigma,N1}$ and biomass cycles; see * in Table 28, comparing with Table 26.

Table 28: Left: Trial11.Run1.txt . Right: Controls.txt

| Parameter | Alga | Graz | Pred | Scav | H2O | NLO | Control | Value | Text |
|---|---|---|---|---|---|---|---|---|---|
| SelfLoss | 0.15 | 0.5* | 0.2 | 0.3045 | NA | 0.8 | YearLength | 12 | NA |
| SelfGrow | 1.1 | 0 | 0 | 0 | NA | NA | Years.sim | 8 or 16 | NA |
| Harvest.Alga | 0 | 1.05 | 0 | 0 | NA | NA | proj.SLorder | 3 | NA |
| Assim.Alga | 0 | 0.4 | 0 | 0 | NA | NA | Parameters.file | NA | C:\ECOLPS\TRIALinputs\Trial11.Run1.txt |
| Harvest.Graz | 0 | 0 | 0.9 | 0 | NA | NA | Envir.file | NA | C:\ECOLPS\TRIALinputs\Trial11.Envir.txt |
| Assim.Graz | 0 | 0 | 0.47* | 0 | NA | NA | | | |
| Harv.NLO | 0 | 0 | 0 | 0.2994* | NA | NA | | | |
| Assim.NLO | 0 | 0 | 0 | 1 | NA | NA | | | |
| Bmass.init | 1.5 | 0.5 | 0.15 | 0.7 | NA | 1 | | | |
| Nconc.N1 | 0.23 | 0.06 | 0.01 | 0.03 | NA | NA | | | |
| Nmass.N1 | NA | NA | NA | NA | 0.35 | 0.35 | | | |
| ParticSL | 0.33 | 0.17 | 0.05 | 0.05 | NA | 0 | | | |
| L.index | NA | NA | NA | NA | 0.131* | NA | | | |
| alphaSG.L | 1.384* | NA | NA | NA | NA | NA | | | |

##### 4.11.2 Run(ii)

As in Run(i) but with fishery $f$ catching $p$.

Table 29: Left: Trial11.Run2.txt . Right: Controls.txt

| Parameter | Alga | Graz | Pred | Scav | H2O | NLO | F1 |
|---|---|---|---|---|---|---|---|
| SelfLoss | 0.15 | 0.5 | 0.2 | 0.3045 | NA | 0.425 | NA |
| SelfGrow | 1.1 | 0 | 0 | 0 | NA | NA | NA |
| Harvest.Alga | 0 | 1.05 | 0 | 0 | NA | NA | NA |
| Assim.Alga | 0 | 0.4 | 0 | 0 | NA | NA | NA |
| Harvest.Graz | 0 | 0 | 0.9 | 0 | NA | NA | NA |
| Assim.Graz | 0 | 0 | 0.47 | 0 | NA | NA | NA |
| Harv.NLO | 0 | 0 | 0 | 0.2994 | NA | NA | NA |
| Assim.NLO | 0 | 0 | 0 | 1 | NA | NA | NA |
| Harv.Pred | 0 | 0 | 0 | 0 | NA | NA | 0.5 |
| Assim.Pred | 0 | 0 | 0 | 0 | NA | NA | 0.7 |
| Bmass.init | 1.5 | 0.5 | 0.15 | 0.7 | NA | 1 | NA |
| Nconc.N1 | 0.23 | 0.06 | 0.01 | 0.03 | NA | NA | NA |
| Nmass.N1 | NA | NA | NA | NA | 0.35 | 0.35 | NA |
| ParticSL | 0.33 | 0.17 | 0.05 | 0.05 | NA | NA | 0 |
| L.index | NA | NA | NA | NA | 0.131 | NA | NA |
| alphaSG.L | 1.384 | NA | NA | NA | NA | NA | NA |
| F.effort | NA | NA | NA | NA | NA | NA | 1 |

| Control | Value | Text |
|---|---|---|
| YearLength | 12 | NA |
| Years.sim | 8 or 16 | NA |
| proj.SLorder | 3 | NA |
| Parameters.file | NA | C:\ECOLPS \TRIALinputs\Trial11.Run2.txt |
| Envir.file | NA | C:\ECOLPS\TRIALinputs \Trial11.Envir.txt |

###### 4.11.3 Run(iii)

As in Run(ii) but with fishery $f$ catching $h$, not $p$.

Table 30: Left: Trial11.Run3.txt . Right: Controls.txt

| Parameter | Alga | Graz | Pred | Scav | H2O | NLO | F2 | Control | Value | Text |
|---|---|---|---|---|---|---|---|---|---|---|
| SelfLoss | 0.15 | 0.5 | 0.2 | 0.3045 | NA | 0.8 | NA | YearLength | 12 | NA |
| SelfGrow | 1.1 | 0 | 0 | 0 | NA | NA | NA | Years.sim | 8 or 16 | NA |
| Harvest.Alga | 0 | 1.05 | 0 | 0 | NA | NA | NA | proj.SLorder | 3 | NA |
| Assim.Alga | 0 | 0.4 | 0 | 0 | NA | NA | NA | Parameters.file | NA | C:\ECOLPS \TRIALinputs\Trial11.Run3.txt |
| Harvest.Graz | 0 | 0 | 0.9 | 0 | NA | NA | 0.5 | Envir.file | NA | C:\ECOLPS\TRIALinputs \Trial11.Envir.txt |
| Assim.Graz | 0 | 0 | 0.47 | 0 | NA | NA | 0.7 | | | |
| Harv.NLO | 0 | 0 | 0 | 0.2994 | NA | NA | NA | | | |
| Assim.NLO | 0 | 0 | 0 | 1 | NA | NA | NA | | | |
| Harv.Pred | 0 | 0 | 0 | 0 | NA | NA | 0 | | | |
| Assim.Pred | 0 | 0 | 0 | 0 | NA | NA | 0 | | | |
| Bmass.init | 1.5 | 0.5 | 0.15 | 0.7 | NA | 1 | NA | | | |
| Nconc.N1 | 0.23 | 0.06 | 0.01 | 0.03 | NA | NA | NA | | | |
| Nmass.N1 | NA | NA | NA | NA | 0.35 | 0.35 | NA | | | |
| ParticSL | 0.33 | 0.17 | 0.05 | 0.05 | NA | NA | NA | | | |
| L.index | NA | NA | NA | NA | 0.131 | NA | NA | | | |
| alphaSG.L | 1.384 | NA | NA | NA | NA | NA | NA | | | |
| F.effort | NA | NA | NA | NA | NA | NA | 1 | | | |

#### 4.12 Trial 12. 3-level food chain re-cycling of N1, seasonal light and temperature cycles, and fisheries

This trial develops Trial 11(i) which included annual, approximately sinusoidal light cycles. Trial 12 adds annual temperature cycles lagging light by approximately 63 days. Both cycles were supplied from Trial12.Envir.txt, see Table 31 below.

Table 31: Trial12.Envir.txt .

| Date.day | Variable | Value | Component | TransferTo |
|---|---|---|---|---|
| 10 | L.index | 0.149 | H2O | NA |
| 40 | L.index | 0.260 | H2O | NA |
| 71 | L.index | 0.441 | H2O | NA |
| 101 | L.index | 0.631 | H2O | NA |
| 132 | L.index | 0.792 | H2O | NA |
| 162 | L.index | 0.870 | H2O | NA |
| 192 | L.index | 0.852 | H2O | NA |
| 223 | L.index | 0.737 | H2O | NA |
| 253 | L.index | 0.563 | H2O | NA |
| 284 | L.index | 0.366 | H2O | NA |
| 314 | L.index | 0.210 | H2O | NA |
| 344 | L.index | 0.131 | H2O | NA |
| 1 | T.index | 0.05 | H2O | NA |
| 5 | T.index | 0.04 | H2O | NA |
| 55 | T.index | 0.07 | H2O | NA |
| 75 | T.index | 0.18 | H2O | NA |
| 100 | T.index | 0.32 | H2O | NA |
| 125 | T.index | 0.47 | H2O | NA |
| 150 | T.index | 0.59 | H2O | NA |
| 175 | T.index | 0.78 | H2O | NA |
| 200 | T.index | 0.85 | H2O | NA |
| 225 | T.index | 0.89 | H2O | NA |
| 245 | T.index | 0.83 | H2O | NA |
| 275 | T.index | 0.67 | H2O | NA |
| 300 | T.index | 0.60 | H2O | NA |
| 325 | T.index | 0.35 | H2O | NA |
| 350 | T.index | 0.06 | H2O | NA |

##### 4.12.1 Run (1)

Parameter values from Trial 11(i) were used initially, with temperature effects on self growth of Algas implemented by setting lower and upper temperature bounds outside which $r_a(I,T) = 0$, see Eq.(26) of Cotter 2026. These and other modifications were made to obtain steady biomass cycles and total system nutrient in 3 differently parameterized runs, Trial12 Run(i)a,b and c. Parameters varying between a,b and c are marked * in Table 32.

Table 32: Left: Trial12.Run1.txt with variants {a,b,c}. Right: Controls.txt

| Parameter | Alga | Graz | Pred | Scav | H2O | NLO | Control | Value | Text |
|---|---|---|---|---|---|---|---|---|---|
| SelfLoss | 0.25 | {0.1,0.2, 0.15}* | 0.1 | {0.17,0.17, 0.24}* | NA | 0.1* | YearLength | 12 | NA |
| SelfGrow | 2.25 | 0 | 0 | 0 | NA | NA | Years.sim | 8 or 16 | NA |
| Harvest.Alga | 0 | {0.2,0.3, 0.2}* | 0 | 0 | NA | NA | proj.SLorder | 3 | NA |
| Assim.Alga | 0 | {0.8,0.8, 0.4}* | 0 | 0 | NA | NA | Para-meters.file | NA | C:\ECOLPS\TRIALinputs\Trial12.Run1a.txt |
| Harvest.Graz | 0 | 0 | {0.22,0.2, 0.22}* | 0 | NA | NA | Envir.file | NA | C:\ECOLPS\TRIALinputs\Trial12.Envir.txt |
| Assim.Graz | 0 | 0 | 0.5 | 0 | NA | NA | | | |
| Harv.NLO | 0 | 0 | 0 | {0.25,0.25, 0.23}* | NA | NA | | | |
| Assim.NLO | 0 | 0 | 0 | 1 | NA | NA | | | |
| Bmass.init | 1.0* | 0.5 | 0.15 | 0.7 | NA | 0.3* | | | |
| Nconc.N1 | {0.23,.205, .206}* | 0.06 | 0.01 | 0.03 | NA | NA | | | |
| Nmass.N1 | NA | NA | NA | NA | 0.35 | 0.07 | | | |
| ParticSL | 0.33 | 0.17 | 0.05 | 0.05 | NA | 0 | | | |
| L.index | NA | NA | NA | NA | 0.131 | NA | | | |
| alphaSG.L | 3.0 | NA | NA | NA | NA | NA | | | |
| T.index | NA | NA | NA | NA | 0.06* | NA | | | |
| LoBoundSG.T | {0.2,0.4, 0.65}* | NA | NA | NA | NA | NA | | | |
| HiBoundSG.T | {0.4,0.7, 1.0}* | NA | NA | NA | NA | NA | | | |
| alphaH.T | {3.8,2.5, 2.5}* | {3.0,3.0, 2.0}* | NA | NA | NA | NA | | | |

##### 4.12.2 Run (2)

Parameter values from Trial 12(i) incorporated as 3 separate food chains side-by-side with numbered component names in a single parameters file with addition of separate essential nutrients. Some pvs were adjusted to improve presentation of resulting time series. See Table 33.

Table 33: Left: Trial12.Run2.txt with 3 independent food chains with different essential nutrients; H2O and NLO components are common to all 3. Right: Controls.txt .

| Parameter | Alga1 | Alga2 | Alga3 | Graz1 | Graz2 | Graz3 | Pred1 | Pred2 | Pred3 | Scav | H2O | NLO | Control | Value | Text |
|---|---|---|---|---|---|---|---|---|---|---|---|---|---|---|---|
| SelfLoss | 0.25 | 0.25 | 0.25 | 0.1 | 0.2 | 0.15 | 0.1 | 0.1 | 0.1 | 0.17 | NA | 0.1 | YearLeng | 12 | NA |
| SelfGrow | 2.25 | 2.25 | 2.25 | 0 | 0 | 0 | 0 | 0 | 0 | 0 | NA | NA | Years.sim | 8 or 16 | NA |
| Harvest.Alga1 | 0 | 0 | 0 | 0.2 | 0 | 0 | 0 | 0 | 0 | 0 | NA | NA | proj.SLord | 3 | NA |
| Assim.Alga1 | 0 | 0 | 0 | 0.8 | 0 | 0 | 0 | 0 | 0 | 0 | NA | NA | Para-meters.file | NA | C:\ECOLPS \TRIALinputs \Trial12.Run2.txt |
| Harvest.Alga2 | 0 | 0 | 0 | 0 | 0.3 | 0 | 0 | 0 | 0 | 0 | NA | NA | Envir.file | NA | C:\ECOLPS \TRIALinputs \Trial12.Envir.txt |
| Assim.Alga2 | 0 | 0 | 0 | 0 | 0.8 | 0 | 0 | 0 | 0 | 0 | NA | NA | | | |
| Harvest.Alga3 | 0 | 0 | 0 | 0 | 0 | 0.2 | 0 | 0 | 0 | 0 | NA | NA | | | |
| Assim.Alga3 | 0 | 0 | 0 | 0 | 0 | 0.4 | 0 | 0 | 0 | 0 | NA | NA | | | |
| Harvest.Graz1 | 0 | 0 | 0 | 0 | 0 | 0 | 0.22 | 0 | 0 | 0 | NA | NA | | | |
| Assim.Graz1 | 0 | 0 | 0 | 0 | 0 | 0 | 0.5 | 0 | 0 | 0 | NA | NA | | | |
| Harvest.Graz2 | 0 | 0 | 0 | 0 | 0 | 0 | 0 | 0.2 | 0 0 | 0 | NA | NA | | | |
| Assim.Graz2 | 0 | 0 | 0 | 0 | 0 | 0 | 0 | 0.5 | 0 | 0 | NA | NA | | | |
| Harvest.Graz3 | 0 | 0 | 0 | 0 | 0 | 0 | 0 | 0 | 0.22 | 0 | NA | NA | | | |
| Assim.Graz3 | 0 | 0 | 0 | 0 | 0 | 0 | 0 | 0 | 0.5 | 0 | NA | NA | | | |
| Harv.NLO | 0 | 0 | 0 | 0 | 0 | 0 | 0 | 0 | 0 | 0.0335 | NA | NA | | | |
| Assim.NLO | 0 | 0 | 0 | 0 | 0 | 0 | 0 | 0 | 0 | 1 | NA | NA | | | |
| Bmass.init | 1.0 | 1.0 | 1.0 | 0.5 | 0.5 | 0.5 | 0.15 | 0.15 | 0.15 | 0.7 | NA | 0.3 | | | |
| Nconc.N1 | 0.23 | 0 | 0 | 0.06 | 0 | 0 | 0.01 | 0 | 0 | 0.03 | NA | NA | | | |
| Nconc.N2 | 0 | 0.23 | 0 | 0 | 0.06 | 0 | 0 | 0.01 | 0 | 0.03 | NA | NA | | | |
| Nconc.N3 | 0 | 0 | 0.23 | 0 | 0 | 0.06 | 0 | 0 | 0.01 | 0.03 | NA | NA | | | |
| Nmass.N1 | NA | NA | NA | NA | NA | NA | NA | NA | NA | NA | 0.35 | 0.07 | | | |
| Nmass.N2 | NA | NA | NA | NA | NA | NA | NA | NA | NA | NA | 0.35 | 0.07 | | | |
| Nmass.N3 | NA | NA | NA | NA | NA | NA | NA | NA | NA | NA | 0.35 | 0.07 | | | |
| ParticSL | 0.33 | 0.33 | 0.33 | 0.17 | 0.17 | 0.17 | 0.05 | 0.05 | 0.05 | 0.05 | NA | 0 | | | |
| L.index | NA | NA | NA | NA | NA | NA | NA | NA | NA | NA | 0.131 | NA | | | |
| alphaSG.I | 3.0 | 3.0 | 3.0 | NA | NA | NA | NA | NA | NA | NA | NA | NA | | | |
| T.index | NA | NA | NA | NA | NA | NA | NA | NA | NA | NA | 0.06 | NA | | | |
| LoBoundSG.T | 0.2 | 0.4 | 0.65 | NA | NA | NA | NA | NA | NA | NA | NA | NA | | | |
| HiBoundSG.T | 0.4 | 0.7 | 1 | NA | NA | NA | NA | NA | NA | NA | NA | NA | | | |
| alphaH.T | NA | NA | NA | 3.8 | 2.5 | 2.5 | 3.0 | 3.0 | 2.0 | NA | NA | NA | | | |

ECOLPS.bib
