## Supplementary material for "Virtual aquatic ecology: stepped simulations of gross production within constraints explain biomass changes": ECOLPS simulator in R: ECOLPS.v32.R

```
#Ecosystem Linear Programming simulator of a fished aquatic ecosystem written by A J R Cotter [].  Published Version 1.0.0; JC's Version 32 last edited 01/Dec/2025 for bugs, tidiness and general good behaviour

#CONTENTS (Search for the text items as spelled below)
   #INSTRUCTIONS
   #MAIN PROGRAM
   #1. PROGRAM CONTROLS
   #2. READING AND IDENTIFICATION OF COMPONENTS AND THEIR ATTRIBUTES
   #3. IDENTIFICATION OF COMPONENT TYPES
   #4. RE-ORDERING OF COMPONENTS, CREATION OF PARAMETERS AND INTERNAL LOCATION VECTORS
   #5. ENVIRONMENTAL VARIABLES THAT CHANGE WITH TIME (if supplied)
   #6. OPTIONAL PRINTING OF INPUT FILES
   #7. INITIALISE DATA STRUCTURES FOR SIMULATIONS
   #8. MENUS AND SIMULATION LOOP
                  
                                      
#INSTRUCTIONS
   #CONTENTS:
      #READ.ME
      #TIPS
      #ABBREVIATIONS USED IN VARIABLE NAMES AND COMMENTS
      #NOTES AND CONVENTIONS
      #INPUT FILES
      #ROUTE MAP
   
   
   #READ.ME
#Disclaimer: This program is made available for simulations of biomasses in aquatic ecosystems as presented in an accompanying paper, Cotter A J R 2026.  Much effort has been applied to making it do what is expected from that paper, and to use similar terms and formulae consistently.  However, it is not possible to declare that software is perfect.  One can only test for expected errors, then wait for unexpected errors to be discovered.  Therefore, USERS are solely responsible for checking all inputs and outputs themselves, and for all consequences of their use of this program.   Copyright details are given in the banner presented by MAIN PROGRAM on the code below.

#WARNING!!!  Running this program COMPLETELY CLEARS !!! the .RData file to prevent any interference by variables left from previous runs.  You must therefore run this program in a different .RData file from any other already on your system associated with other applications of R.

#Familiarity with R is not necessary for running published Trials using ECOLPS, only if you wish to change, test or investigate something in the code.  

#I used Tinn-R in Windows 10 for writing, de-bugging and running ECOLPS. It is a compact text editor that conveniently allows you to submit the entire program of code, selected lines, or just variable names for direct processing by R in its graphic user interface (GUI).  R code words, the cursor line, and paired brackets are all highlighted after you have set [Options][Syntax highlighter][Set][R]. Of course, you can also use Tinn-R to modify data, text and other language files. An unusual but useful feature for editing individual columns of data is the column-editing button (at the bottom of the window). Another is the [Editing] button (also at the bottom of the window) which allows you to protect files you don't wish to alter unintentionally, eg. ECOLPS. 

#R-studio:- I briefly experimented with R-studio but ECOLPS did not run, possibly because of the source("ECOLPSfunctions.R") statement at the beginning, intended simply to clear the main code of written functions that seldom needed attention. If you prefer R-studio, maybe you only have to shift all of the functions to the top of the main code as one file to get it to work. I had all I needed in Tinn-R and made no further investigations of R-studio.

#Other text editors are suitable for editing R code, including that supplied with R itself.  Please refer to manuals to find how best to run R code in these situations.

#The following instructions are for Tinn-R.  They are intended to assist you to make simulations with ECOLPS.  Please note that I have only a basic understanding of the linking of Tinn-R with the R GUI. You may prefer other approaches, or to use Tinn-R's Help file to assist with any problems. The ECOLPS.R code file has further extensive text to help you understand the simulation system and its code, for which, see below.

#METHOD
#i) Download Tinn-R, a text editor intended to pair with R, available from https://tinn-r.org/en/download.  There are alternatives to Tinn-R but I have had no reason to investigate them. Tinn-R highlights R code words and differentiates code from #text comments thus greatly enhancing readability.  It also allows direct submission of all or selected parts of the code to the R interface, making a convenient platform for development and debugging. Another feature allows selection of vertical columns (see button in bottom margin).This is useful when removing, duplicating, or adding entire component columns in the Params.file. [Alternatively, components can be temporarily removed by renaming their column as NA.]  Next to this button, another button toggles between editing and read-only which is helpful for preventing accidental corruption of the code. I work by pasting all lines of this program, ECOLPSfunctions.R, Controls.txt and any Trial files into separate tabbed pages in Tinn-R. 

#ii) Download R from https://www.r-project.org/  if you have not already got it on your machine.  My copy of R was installed to C:\Program Files, the default storage location. 

#iii) ****NOT SUCCESSFULLY OPENED AT 2 Dec 2025****Open an account for Zenodo at https://tinyurl.com/ECOLPS.  There you should find ECOLPS.v32.R, ECOLPSfunctions.R (both code files are needed for simulations), Controls.txt, and Trial.txt files numbered as in the paper. The input files for each trial all go together in this folder; they will be in alphabetic order.  However, I am new to, and somewhat confused by Zenodo.  If I have left access problems for new users, please try me at or the Zenodo help resource.

#iv) Download ECOLPS.R, ECOLPSfunctions.R, and Controls.txt from (location to be decided).  All three should be stored in the same folder so that they are found by ECOLPS, for example C:\ECOLPS\ .  Then you will not have to change the basic file refs in the ECOLPS code. ECOLPSfunctions.R is automatically read with source() at the start of the Main Program near line 177, and Controls.txt is read with read.table() in section 1.0 near line 200.

#v) Open Tinn-R and click the [R] button (near the right end of the lower icon ribbon in my unmodified installation). Wait to see >startSocketServer(port=xxx) then, 2 or 3 secs later, [1] TRUE. Next, arrange Tinn-R and R so that they both share the screen using the [Visualization: organize screen (Tinn-R/Rgui)] button in the top ribbon centrally. R code, loaded into Tinn-R, can now be sent directly to R for execution using the [R send:file] or the [R send:selection] buttons at the left of the lower Tinn-R icon ribbon. Hover the cursor over these buttons and they identify themselves. [R echo off] starts/stops R echoing submitted code (which is confusing for a long program and unnecessary). 

#vi) If, now, you load the ECOLPS code into Tinn-R then send it to R using [R send:file], the program will start but it may not find the files it needs, ie. ECOLPSfunctions.R, Controls.txt, and .txt data files . R announces "Error in file(filename, "r", etc)". The first two must be in the same folder as ECOLPS if they will be found automatically by ECOLPS without augmenting file references in the code (eg as " \\folders\\file").  Additionally, R's own directory should be set to the ECOLPS folder. To do this:-  In the R GUI, press [File][Change dir...] . If you've followed the advice so far, set it to C:\ECOLPS\ where ECOLPSfunctions.R or Controls.txt have been stored alongside ECOLPS.R, and you should get no further bother from this. The .txt data files must be given full file references, without quotes, in the text (2nd) column of Controls.txt .  

#vii) To run different simulations with ECOLPS, edit fields of Controls.txt in C:\ECOLPS\, notably those related to the parameters and environment .txt file references, the number of projections, year length, reports, graphic window size, etc. This is easy when you are familiar with what is needed. To assist with learning, specific Controls.txt data for each Trial and Run are printed in Supplementary Information Annex D.
#DONT FORGET to save Controls.txt (or Trial.txt files) after editing, then switch tabs in Tinn-R back to ECOLPS.R to run ECOLPS. Otherwise you'll be puzzling over R error messages; ECOLPS only reads from saved files.  Please also look through the extensive advisory notes lower down in this ECOLPS code file. 
 
#viii) If ECOLPS gives problems, after closing ECOLPS, you can find the values of variables by typing their exact name or by selecting the name in the code in Tinn-R and pressing [R send:selection].  The result appears in the R GUI. It may help explain what ECOLPS has done.  If you are confident that there is a coding bug, please think about it for a day or two with a colleague if possible, then email to me the relevant Controls.txt, Parameters.file, Environ.file (if used), a list of any menu choices made, plus copies of the ECOLPS.R and ECOLPSfunctions.R you were using if at all modified from source.  Then, no promises, I might consider it. You could also tell me about code improvements you make yourself but please test them extensively with both LV and multi-component ecosystems first.  Proposed edits to these guidance notes would also be welcome.

#ECOLPS will open Controls.txt, the Params file (see below), and an Environ file if supplied.  YearLength X Years.sim projections, supplied in Controls.txt are then run automatically (only 1 sec for 1000 LV projections in Trial1.Run1.txt) before the Main Menu appears on screen inviting you to look at constraints (active, slack variables, limits) affecting selected components, or at simulated variables such as biomass, harvest, landings, etc. You choose from the Constraints Menu or the Variables Menu. There, your selections are put as rows in Output.set.  They are graphed as time series with G or g, printed on screen as data with P or p.  Numerous other options for output are in the menus.  Output.set may be edited (E or e) to remove rows, or cleared altogether (C or c) ready for a new start. Remember to clear Output.set when moving from the Constraints to the Variables menu, or vice versa, unless you want both types in output.  An empty line + ENTER takes you1 back to Main Menu. Options 8 and 9 run another block of simulations, over-writing or appending them to those already present in Output.set. Press Escape to terminate.  Output.set or other variables can be typed at the R > after termination to investigate the workings of the last simulation. Highlighting text shown on the R GUI allows you to copy and paste to another programme or file for printing.

#A 'Controls.txt' file supplies controlling variables to ECOLPS, for example specifying source data files, the number of simulations, how to print or graph resultson your screen, etc. 'Controls.txt' is a fixed name coded into this program (near line 200).  The file has 3 columns: 1)Row headers 2)LEFT column for numeric variables 3)RIGHT column for text variables such as file references.  Every row must have a row header and 2 column elements so that R knows when to move to the next line. Elements that are inapplicable must have NA (R code for Not Available/Applicable) in place of a value. For example, in the row headed Parameters.file, the LEFT value is NA, the RIGHT value is the file reference on your system (written without quotes). Rows may be omitted where defaults can be accepted. Defaults are coded in MAIN PROGRAM#1 below.
 
#A 'Params.file' supplies the required quantities and coefficients for simulating an ecosystem with given 'components'(=living or non-living parts of the system) , where 'Params.file' refers to a file reference decided by the user that is included in Controls.txt.  The Params file has at least 3 columns: 1) Row headers 2) Autotrophic component, 3) Heterotrophic component. ECOLPS requires at least one autotroph and one heterotroph eating it. This is called a Lotka-Volterra (LV) 'minimal' system here.  After column 2, 1 column is added for each additional living or non-living component, the latter possibly including water, non-living organics (NLO), and essential habitats. [Essential nutrients are treated as properties of water etc so nutrient masses and concentrations are variables that go into rows.] The ordering of rows and columns does not matter because they are re-ordered in a standard way in #4.  The row headers name the variables, often with a dot and a suffix specifying the component, or other special category, to which it applies. Examples: 'Harv.alga3' names the row of harvest coefficients for heterotrophic components harvesting alga3 as a food; 'NutrConc.N3' names the row of constant concentrations of nutrient N3 in the tissues of living and NLO components.
 
#NAs (NA=not available) fill unused element positions in Parameters.file so as to allow R to read the same number of values from each row. For example, biomass is not applicable to fisheries so NA is inserted in the fisheries columns of the Bmass row. 0s may also be used for 'not applicable'.  Inf=infinity is appropriate for some params, eg quota that do not restrict landings can be Inf.  NAs are converted to 0s by ECOLPS because NA values prevent calculations. 
 
#Component types are defined and automatically put into the simulation model by ECOLPS according to the rules listed at the start of code section #3.  From those rules, it can be seen that most variables supplied in the Params.file must be given positive values because this is how component types are identified.  If a zero is thought most appropriate, for example for a value of T.index in H2O, the input value can be approximated as a very small positive value, say 0.000001. This will allow ECOLPS to identify the H2O column but have no effect on the simulation; later, this initial value might be replaced by a 0 from an Environ.file - see below.
 
#Phytoheterotrophs, cannibals, parasites, and mutualism among living components, though important ecologically, are NOT yet handled explicitly by ECOLPS. Untried 'fixes' are suggested here-and-there in the code below.

#Much of ECOLPS's initial code is for categorising cols of the Params.file into component types, and for warning users of combinations of supplied parameter values that are incompatible with the way ECOLPS is currently implemented. Imposing error statements are screened if you contravene requirements. The explanatory text with each error should help you to find the problem, probably in the Params file. In this way, categorisation of components is conveniently done for you, and should achieve consistency among users.  'Wild' or 'Living' components include autotrophs and heterotrophs. 'Foods' are any wild or NLO (non-living organic) component that is eaten or landed. Consumers are heterotrophs and fisheries.  Non-living components include NLO, H2O, Habitats, and Fisheries. 

#The following descriptors of components are used:-
 #(1) Bmass or biomass is organic weight of living or NLO components consistently in the dried, or wet condition
 #(2) Nmass.N1 is weight of nutrient N1 in H2O and in NLO components; this is why the simulated ecosystem must be closed to external inputs (code for transfers among neighbouring systems is not present in this version of ECOLPS)
 #(3) Nconc.N1 is concentration (wt/wt) of N1 in living components and NLO; it is not used for H2O where only Nmass.N1 is relevant
 #(4) Effort is fishing nets in the water, hours of trawling, hooks set, etc.  Effort is assumed constant over each short projection interval, Delta.
 #(5) T.index, L.index refer to H2O only
 
#An Environ.file, if required for alterations over time, is supplied with a file reference in the last column of Controls.txt. Environ.file has 5 columns: date (=day number, minimum value is 1), variable, value, component affected, and 'transfer-to' refering to a recipient component.  This file allows variation of parameters after Date=0 as functions of Date in days, variations of T.index and/or L.index (from which seasons may be simulated), and/or transfers of biomass between components at specific seasonal dates, for example migrations or growth through different life stages thought to merit different component identities. 
 
   #TIPS 
#TIP1. To check that calculations and processing by ECOLPS are proceeding as expected, insert a stop("line number") command at any line in the main code. [Note: The stop() function automatically prints out an 'error' statement of its own plus whatever you add in quotes in the brackets.  You can over-write the general error statement with 7 back spaces, \b ].  When execution stops, you can submit selected lines or variable names (from Tinn-R), or paste variables into R at the prompt to discover their current value. However, this only works for the main code, not for functions, most of which are stored out of the way in ECOLPSfunctions.R.  To evaluate variables in functions,  you must insert `print(variable)' `stop()' into the function.  This is because functions store variables at a higher level in R which cannot be accessed directly (so far as I know).  Functions read variables in the main code but do not directly write to them, even if the name is the same; the value must be explicitly returned to the main program from the function using either the last statement in the function or an explicit return() statement.

#TIP2. Switch off any unrequired code in a line with # at the beginning. Useful for de-bugging.
   
#TIP4. If you wish to adjust or supplement the code below in any way, make sure to test the product on Lotka-Volterra 2-component systems as well as on varied, more complicated multicomponent systems to verify that the new code allows for components being present and/or absent. Some functions() in R only work with matrices having 2 or more rows and 2 or more cols. Minimal LV models may have data in single rows or cols, sometimes causing an R error. Additionally, it would be advisable to familiarise yourself thoroughly with the structure of ECOLPS before making any changes because, like an ecosystem, just about everything is connected to something else! If you distribute the adjusted code, please state clearly that your code is an adjustment of this version as requested in the full opening banner at the start of Main Program below. Please also describe tests made on the adjusted code to check for bugs.


   #ABBREVIATIONS USED IN VARIABLE NAMES AND COMMENTS
#A = autotroph; assim = assimilation (of harvest) coefficient; Bmass = organic biomass; C = constraint or column; Cat=caught or catch by fisheries; coef = coefficient; col or C = column; Cpar = col number of Params matrix;  Cgain = col number of gainers matrix, Co = col (or element) number of Objective function; Cv = col number of Variables matrix; comp = component of ecosystem (living or non-living) ; conc = concentration wt/wt; D1. D2. D3 = 1st, 2nd and 3rd derivatives with respect to time; eff = effort by fishery; F = fishery; GL = gross loss; GP = gross production; grow = growth; harv = harvest; H = heterotroph (may include predators and scavengers but not fisheries); Inf = infinity; Loc = location of an element in a vector; N. = number of; nutr = nutrient; NLO = non-living organics; proj = projected; Q = quota; Rb = row number of biomass matrices; Rc=row number(s) of coeff matrix; RCc = row and column numbers of a coeff matrix; RCpar = row and col of Params matrix; RCv = row and col number of Variables matrix; S = scavenger (eating NLO, possibly living prey also); var = variable; TGP = total gross production (all GPs + landings); W = wild (living components, i.e. not fisheries or physical components); wts = weights;

   #NOTES AND CONVENTIONS
#NOTE 1. Time is measured internally as a positive, integer number of Delta units. 1 Delta=1 projection, or simulation short-interval.  'Years' may be used for output, and for Years.sim supplied in Controls.txt to specify how long the initial simulation should be; it can be extended from the Main Menu.  Years are related to Deltas with YearLength as set in Controls.txt.  YearLength may be altered so as to vary the length of 1 Delta unit relative to a year without affecting the length of the simulation or the timing of interventions. Coefficients with a per.time dimension are affected by YearLength and must be adjusted to have the same effect: eg 1 tonne per day = 7 tonnes per week when Delta is changed from 1 day to 1 week (by changing YearLength from 365 to 52).  Standard 'Days' are used in the Environ.file to time interventions. All years have 365 days.

#NOTE 2. Input tables or matrices of trophic-type variables have rows = foods, and cols = consumers, by convention.  Otherwise it is easy to get confused which is which.  However, transpositions using t(matrix) occur as necessary mathematically within the program code.  

#NOTE 3. Unexpected input to the program gives either an ERROR or an ADVICE message.  ERROR terminates the program.  ADVICE is not necessarily critical and does not stop the program. It is easy to ignore advice but, if you have a problem, go back and read it. It may help!  The location in the code where an error is found is usually marked with a unique, searchable code, eg "#4.1" .  Advice messages may be bothersome.  Their number can be reduced with Advice.level as set in Controls.txt. 0=almost no messages, 1=quite a lot, 2=All. The if(AL > ) code at the beginning of cat("\nADVICE:" . . . ) statements below could also be altered individually to adjust which advice appears for each level of AL.

#NOTE 4. ECOLPS finds variable names in input files by searching for occurrences of the labelling text in the row or column names using R's grep() [gives all occurrences] or grep()[1] [gives 1st occurrence only] functions.  The searched-for text may be all or part of the variable name, prefix or suffix in lower or upper case.  This means that the rows and columns in input files can be in any order, and prefixes and suffixes are OK in the row names.  Take care, however, to spell names exactly right if the name is used in two places, and not to have one text string in two different items, eg. Parameters and Params; they will both be found with grep("Param" ..).  [R note: If grep("text", ...) does not find "text" it gives logical[0]. If grep("text", ...)[1] does not find "text" it gives NA.  This affects if() statements using the result.  Therefore, removing [1] may affect dependent if() statements.]

#NOTE 5. Suffixes separated by a '.' are used in var names when, for example, a variable has an associated component or type, e.g. ConcNutr.Nitrate.    ECOLPS uses the '.' to separate the variable name from the component or type and tie it in with other components or suffixes. Remove or re-position the dot at your peril!
   
#NOTE 6. Code was written on a screen with 230-character lines with comments adjusted to fit on one line if possible.  This wide screen assisted programming and debugging but, unfortunately, means that users with narrower screens will have many line overflows.
 

   #INPUT FILES
#Three different input files with different formats are used by ECOLPS.
 #Controls.txt
 #PARAMETERS FILE
 #ENVIRONMENT FILE

#Controls.txt (NO OTHER NAME is accepted!)
 #Controls.txt MUST be supplied in the same file folder as ECOLPS.R and ECOLPSfunctions.R [or the file ref near line 200 could be adjusted].  The main program searches for each type of control by applying grep() to the row names in Controls.txt.  Spelling exactly as shown in the list below should prevent unwanted variables being found (but case is unimportant).
 #The 3 columns of Controls.txt are headed Control, Value, Text.  The variable's name is the row header under Control.  Numerical values go under Value; text values (e.g. file names) go under Text (without quotes "").  R has problems when text and numerical data are mixed in one column, hence the need for separate Value and Text columns. You MUST insert NA (=not available/applicable) in every unused col of the row for either type of variable.  Any comments at the end of the line must be preceded by #; this way, R can count and locate the values correctly in each row.
     
      #Controls that may be included are:

#Row name                 Meaning                                                             Default value

#Advice.level             level of advice about inputs, and non-terminal errors               0=no advice, 1=some advice, 2=all advice (default)
#YearLength               number of Deltas per year (for annual cycles etc)                   eg. 52 means Delta is equivalent to one week
#Years.sim                duration of simulation in years                                     eg. 3
#Progress.rep             interval for showing simulation progress                            100 deltas.  How often do you need to know what the program is doing?  Each time, one line of info is put on screen
#proj.SLorder             order of differentials in Taylor-series projections of self loss    optional, 3=default, or 1, or 2 are possible; only needed for testing
#View.params              switch to show on screen the Params.file after processing           optional, 0=no, 1=yes
#View.Envir               switch to show the Environment.file after processing                optional, 0=no, 1=yes
#View.controls            switch to show the Controls.txt file after processing               optional, 0=no, 1=yes
#View.components          switch to show the component-type table formed from Harv coeffs     optional, 0=no, 1=yes
#KeyItemWidth             N chars for key names on graphs; stops truncated names, bulky keys  14 or 19 suggested; depends on WinHeight, WinWidth and the number and names of variables plotted
#KeyLineLength            N characters per line of the key (to fit WinWidth of graph)         eg. 60
#WinHeight                usual height of graph windows in nominal inches                     eg. 7
#WinWidth                 usual width of graph windows in nominal inches                      eg. 10
#Parameters.file          input file reference for ecological components                      no default, a file reference is REQUIRED
#Environ.file             input file reference for changing variables over time               optional with no default; if not present params are constant, and biomasses unadjusted, throughout the simulation
#GraphLabel               default label for graphs                                            label could be text, or date, time, Params.file (no quotes) in col 2
#ShowCalcsFromStep        show all calculations for every projection from step number in col 1 to YearLength * Years.sim.  Output for checking appears above the first Main Menu. May be voluminous. Default=NA or Inf or omit row
#Col.plots                graphs are in colour or black and white only                        optional, default 1=colour, 0=B & W

#PARAMETERS FILE (variably named but always with a .txt suffix eg. Trial1.1.txt)
 #The Params.file MUST be supplied and named as shown in the text col of Controls.txt; use eg C:\folders\file.txt.  It supplies parameters and initial values in columns, one for each ecological component.  Note that, at input, a zero is equivalent to NA where a positive value is required; this is so that relevant parameter values can be identified; if zero is required, use a very small positive value, eg. 0.000000001.
#Components may be 'living wild' (e.g. groups of species and/or life-stages), 'non-wild' (e.g. fisheries), or 'non-living' (e.g. NLO, essential habitats, water).  Column headers name the component; they should be unique and short for compact labelling of tables, graphs and keys. Take care that any component names found in col headers and in row.suffixes of this or any other input files are all spelled exactly the same so that they can be matched by ECOLPS.  If unmatchable, error messages will be shown. A component may be temporarily removed from a simulation by simply replacing its header with NA.
#The Params.file may include the following rows in any order, not all of which are required or relevant for every component.  When a variable is not relevant to a component, an NA or zero must be inserted in its column.  Optional rows can be omitted for some components but not all: eg. you must supply Effort.init if fisheries are given. ECOLPS tells you if you have omitted something essential.  The rows include (after the row of headers):

#Row name       N rows        Meaning                                                             Notes
#SelfGrow       1             self-growth coefficient.                                            Positive, finite for all autotrophs
#SelfLoss       1             self-loss coefficient.                                              Positive, finite for all wild and NLO components
#Harvest.food   1 per food    harvest coefficient.                                                Positive, finite for each harvested food in all Hetero and fishery cols
#Assim.food     1 per food    assimilation efficiency coeff.                                      0 < Assim < 1 in all H or F cols of each .food row to match positive Harv coeffs
#Effort         1             initial fishing effort                                              Positive, finite for all fisheries
#HabSize        1             total empty size of each essential habitat.                         Positive, finite for all habitat components
#Occup.hab      1             per habitat area-per-organic mass useage coeff.                     Positive, finite for all living components requiring the habitat; others have zero
#Nmass.nutr     1 per nutr    initial mass of .nutr in water.                                     Positive, finite for all nutrients in H2O and NLO cols (1 each, if present). REQUIRES H2O comp; and an Nconc row
#NConc.nutr     1 per nutr    conc of .nutr in tissues or substance of components.                Positive, finite for all nutrients in wild columns. REQUIRES H2O component with Nmass
#Bmass.init     1             initial organic mass of each living comp. and NLO                   Positive, finite for all living components and NLO
#Gmax           1             maximum growth rate regardless of food.                             for any heterotroph/predator; optional. 0 stops growth; Inf removes the .SATE. constraint
#Hmax           1             maximum harvest rate regardless of prey.                            for any heterotroph/predator; optional. 0 stops harvesting; Inf removes the .HARV. constraint
#ParticulateSL  1             particulate self loss coefficient.                                  0 < ParticulateSL < 1 for wild comps, 0 for NLO if present. REQUIRED if nutrients or NLO are present; otherwise optional
#Quota.F        1 per fishery initial permitted landings/year, organic mass units. 0<=real<=Inf   'F' names the fishery.  REQUIRES fishery F; otherwise optional; 0 and Inf are OK
#T.index        1             temperature index                                                   Coolest=0 < T.index <= 1=hottest in an H2O col. Optional. REQUIRES H2O component
#L.index        1             light index                                                         Positive, darkest=0 < L.index <= 1=brightest light in an H2O col; optional. REQUIRES H2O component
#alphaSG.I      1             slope of SelfGrow function over L.index.  0<real<Inf                Adjusts SG of autotrophs with L.index and T.index values in Envir.file; optional for seasonal effects
#LoBoundSG.T    1             lowest T.index permitting positive SelfGrow after adjustment        0 <= LoBoundSG.T < T.index.init <= HiBoundSG.T <= 1
#HiBoundSG.T    1             highest T.index permitting positive SelfGrow after adjustment       0 <= LoBoundSG.T < T.index.init <= HiBoundSG.T <= 1
#SlopeHvst.T    1             slope of temperature index over T.index.  0<real<Inf                Adjusts Harvest coeff of heterotrophs with T.index; optional [spelled with Hvst so grep("Harv") finds Harv, not Hvst as well]
#LoBoundHvst.T  1             lowest T.index permitting positive Harv coef after adjustment       0 <= LoBoundHvst.T < T.index.init <= HiBoundHvst.T <= 1
#HiBoundHvst.T  1             highest T.index permitting positive Harv coef after adjustment      0 <= LoBoundHvst.T < T.index.init <= HiBoundHvst.T <= 1


#ENVIRONMENT FILE (variably named eg. Trial10.Envir.txt)
 #The Environment file holds data that vary with date (always a day number) after the start of the simulation. The Envir.file is optional but, without it, you can only simulate with the initial fixed parameters in Params.file.  The Date.day column is used to time interventions relative to a year.  Hence, eg. Days=465 represents day 100 of year 1 (the first year being 0).  The Vars file is automatically stacked as necessary in units of 1 year to cover the full length of the simulation.  Each required block is stacked at the beginning of a year so that any seasonal cycles present in the Vars file have effect at the same time of year.
 #Cols of Environ.file should be named Date.day, Variable, Value, Component, and TransferTo.  'Date.Day' is used to prevent confusion with R function date() and other similarly named vars
 #Variable names given in the 2nd col of Envir.txt must exactly match the spelling of those in the Params.file, possibly as prefixes. or .suffixes .
 #Initial values of each variable, applicable at Days=0, should be put in the Params.file, NOT in the Environment.file.  That way, the initial values are not recycled when the Environment.file is stacked.  e.g. Initial nutrient  masses may not be the same as what is added on January 1st (Days=1) of each subsequent year.  For this reason, the first value of Date.day in the Environ.file should be greater than 0.
 #Values read from a row of Envir.txt may represent a one-off addition eg. for a nutrient, or a value that remains constant until the next time a value is read, eg. for light and fishing effort.  Light (also called irradiance) and Temperature indexes can provide seasonality or other time-related variation for plant growth or harvesting activities.
 #The columns of the Environ.file are:-

#Name           N cols        Meaning                                                             Notes
#Date.day       1             Calendar days from start of simulation.                             Positive: 0 < Date < Inf for all rows.  ECOLPS sorts all rows into Date.day order.
#Variable       1             System-wide name of the variable                                    REQUIRED for all rows
#Value          1             Numeric value of the variable                                       REQUIRED for all rows
#Component      1             System-wide name of component affected by Variable                  REQUIRED for all rows
#TransferTo     1             System-wide name of component transfered to                         NA REQUIRED except when Variable=Trans


#ROUTE MAP FOR CODE
 #ECOLPS prepares the code scene in #0:- the R directory is cleared, the boot() function is read from R's libraries, functions called repeatedly are read into the front of the code from ECOLPS.R with a source() statement. 
 #ECOLPS next reads Controls.txt, a Params.txt file and, possibly, an Environ.txt file.  Input is checked extensively for inconsistencies or unacceptable values; data structures needed for the simulation are created.  All this takes a little time.  Processing then goes straight into section #8 where simulations are calculated and menus are implemented.
 #On the first visit to #8, the Main Menu is not shown before carrying out the number of simulation steps specified by YearLength and Years.sim in Controls.txt. After this, variables available from all simulation steps but not otherwise needed are calculated for possible display. Then the user visits Main Menu for the first time. Here, she/he can choose to look at constraint features with the Constraints Menu, or at simulated variables with the Variables Menu. Both types of data may be combined by switching to the other menu without clearing the Output.set (option C).  Main Menu option 9 allows the simulation to be extended as another block.  Option 8 also continues simulations but overwrites those already stored.  The user is returned to Main Menu when each selected option has been carried out. When finished with all options, press [ESCAPE] to close the program. 

#####################################################################################################################################################
###MAIN PROGRAM
   #SECTIONS
      #0. PREPARATORY
      #1. PROGRAM CONTROLS
      #2. READING AND IDENTIFICATION OF COMPONENTS AND THEIR ATTRIBUTES
      #3. IDENTIFICATION OF COMPONENT TYPES
      #4. RE-ORDERING OF COMPONENTS, CREATION OF PARAMETERS AND INTERNAL LOCATION VECTORS
      #5. ENVIRONMENTAL VARIABLES THAT CHANGE WITH TIME, if supplied
      #6. OPTIONAL PRINTING OF INPUT FILES
      #7. INITIALISE DATA STRUCTURES AND LOCATION VECTORS FOR SIMULATIONS                                        
      #8. MENUS AND SIMULATION LOOPS


#0. PREPARATORY 
rm(list=ls())                                                    #BEWARE!!! TOTALLY CLEARS R directory of internal data; necessary to avoid lingering variables from previous runs; don't have any other stuff stored there
library(boot)                                                    #source for simplex() linear programming function
options(scipen=4, max.print=99999)                               #discourages some scientific notation for numbers -> easier-to-read results; default max (lines?) of output screened (can cut out variables if too small)
source("C:/ECOLPS/ECOLPSfunctions.R")                            #functions called by ECOLPS are sourced from ECOLPSfunctions.R which must be in the same directory as ECOLPS, with R connected to that directory. [Change directory] if necessary. source() makes the coded functions available here without extensively cluttering the start of this main-program file. Alternatively, the contents of ECOLPSfunctions.R could inserted here if the source() statement gives trouble. (I suspect, possibly, it prevented me using Rstudio when I tried.)


#1. PROGRAM CONTROLS
#Program controls are read from Controls.txt, or defaults are used. See notes at the start of this file.
   #SUBSECTIONS
      #1.0 Read Controls.txt
      #1.1 Advice.level for run-time events that do not stop the program
      #1.2 YearLength (Deltas per year)
      #1.3 Duration of the simulation in years:- Years.sim
      #1.4 Order to use (1st, 2nd, 3rd) for Taylor-series projections of self loss
      #1.5 Progress reporting interval:- Progress.rep
      #1.6 Switch for displaying Params.file:- View.params
      #1.7 Switch for displaying Environment.file (if present):- View.Envir
      #1.8 Switch for displaying Controls.file:-  View.controls
      #1.9 Switch for displaying the component-type table constructed from harvest and other coefficients:- View.compType
      #1.10 Simulation number from which internal calculations are screened for EVERY projection:- ShowCalcsFromStep
      #1.11 Graph window height
      #1.12 Graph window width
      #1.13 Graph-key parameters: KeyItemWidth
      #1.14 Parameters file name
      #1.15 Environment file name, if supplied

#1.0 Read Controls.txt
#The short file ref below requires that Controls.txt is stored in the same folder as this program and ECOLPSfunctions.R . It can optionally by a full file ref, for example,    "C:\\ECOLPS\\Trial1\\Controls.txt"
Controls <- read.table("C:/ECOLPS/Controls.txt", header=TRUE,  row.names=1, , as.is=TRUE )                   #read file to Controls; as.is prevents R from making factors; 1st col becomes row names
if(class(Controls[ , 1]) == "character")                                                                     #If 1st data col is class(character), probably because of characters somewhere in it, advise error:-
   {AL <- 2                                                                                                  #AL is needed for Advise() which stops the program because Fatal=TRUE
   Advise(WithRefTo="Column 1 of Controls.txt", ErrText="The first data column is of class 'character'.  It should be numeric.", AdvText="Remove all non-numeric characters from this column.  \n\tNA MUST be used if a numeric value is not applicable in column 1. \n\tCharacter data are only read from column 2.", ShowData=NULL, CallSection="#1.0", Fatal=TRUE, MinAdviceLevel2Show=0) #\n gives new line, \t gives a tab
   }
Delta <- 1                                                                                                   #Delta might be varied but only for special investigation of effects
#----------------------------------------------------------------


#1.1 Advice.level for run-time events that do not stop the program
#Values of AL are used in the code to select which advisory messages you want to see, and which you don't.  Adjust AL and AdviceLevel values {0,1,2} in advisory code lines anywhere in this program to personalise screen outputs
#Advice is issued by AdviseAdvice() in ECOLPSfunctions.R if AL > MinAdviceLevel2Show                         #findOneLine() and is.DatumOK() below are in ECOLPSfunctions.R
AL <- 2                                                                                                      #Set default AL = 2 => screen ALL advisory messages for beginners
Advice.level.R <- findOneLine(SearchTxt="Adv", Headers=row.names(Controls), call.section="1.1a Advice.level", file="Controls.txt", Var.name="Advice.level", domain="row", ZeroOK=TRUE)  #find row.header with 'Adv' in it.
if(!is.na(Advice.level.R))                                                                                   #If the AL row is found:-
   {Temp <- Controls[Advice.level.R, 1]                                                                      #store the supplied value temporarily
   ERR <- is.DatumOK(Temp, "Advice level", NA.OK=FALSE, CharOK=FALSE, NumbOK=TRUE, LogicalOK=FALSE, MustBeInSet=c(0,1,2)) #test the value with is.DatumOK()
   if(ERR==TRUE)                                                                                             #If ERR=TRUE:-
      {Advise(WithRefTo="Advice.level in column 1 of Controls.txt", ErrText=paste0("The supplied value for Advice.level is ", Temp, ".  This is not in the acceptable set:- 0 -> no advice, 1 -> some advice, 2 -> full advice."), AdvText="Adjust the AL value given OR remove the Advice.level row in Controls.txt => default = 2.", ShowData=NULL, CallSection="#1.1b", Fatal=FALSE, MinAdviceLevel2Show=0)
      } else {AL <- Temp}                                                                                    #else over-write default AL
   }
#Display banner, given Advice level chosen
Version.JC <- "32"; Version.Zenodo <- "1.0.0" ; LastEdit <- "1 December 2025"
cat("\n**************************************************************************************************************************************************\nECOLPS:- Copyright CC BY-SA 2025 by A.John R.Cotter. See https://creativecommons.org/ This program is distributed WITHOUT ANY WARRANTY WHATSOEVER.\n")

if(AL>=1)
   {cat("Version.AJRC: ", Version.JC, ", last edited: ", LastEdit, ", Version.Zenodo:", Version.Zenodo, ".  Contact\n")
   }
if(AL==2)
   {cat("THIS PROGRAM simulates relative GP, organic biomasses, and related variables in an aquatic ecosystem using linear programming within constraints.\n")
   }                                                                                                                                
cat("**************************************************************************************************************************************************\n")
cat("Advice level = ", AL)                                                                                   #routinely notify of current advice level
#----------------------------------------------------------------                       \n


#1.2 YearLength (Deltas per year)
#YearLength sets the number of projections (Deltas) made within 1 year.  If years are unimportant, use YearLength=1.  For, eg., monthly results, use YearLength=12.
YearLength <- 1                                                                                              #set integer default, 1 "year" = 1 Delta; alternatives could be 52 for weeks, 365 for days
YL.R <- findOneLine(SearchTxt="YearL", Headers=row.names(Controls), call.section="1.2a YearLength", file="Controls.txt", Var.name="YearLength", domain="row", ZeroOK=TRUE)  #find row.header with 'YearL' in it
if(!is.na(YL.R))                                                                                             #If the YearLength row is found:-
   {Temp <- Controls[YL.R, 1]                                                                                #store the supplied value temporarily
   ERR <- is.DatumOK(Temp, "YearLength in Deltas", NA.OK=FALSE, CharOK=FALSE, NumbOK=TRUE, LogicalOK=FALSE, InfinOK=FALSE, MinNotOK=0, MaxOK=365, nonIntegerOK=FALSE)#test that value is OK
   if(ERR==TRUE)                                                                                             #If ERR=TRUE:-
      {Advise(WithRefTo="YearLength in column 1 of Controls.txt", ErrText=paste0("The supplied value for YearLength is ", Temp, " Deltas per year.  This is not an integer in the acceptable range:- 0 < YearLength <= 365"), AdvText="Adjust the value given OR remove the YearLength row in Controls.txt => default = 1; the number of years (Years.sim) could be increased instead.\n\tYearLength sets how many simulation steps (Deltas) will be carried out over Years.sim, the duration of the simulation in nominal years.", ShowData=NULL, CallSection="#1.2b", Fatal=TRUE, MinAdviceLevel2Show=1)
      }
   YearLength <- Temp                                                                                        #no error found in Temp; overwrite default YearLength
   }    
#----------------------------------------------------------------


#1.3 Duration of the simulation in years:- Years.sim
#The duration of the simulation is measured in `years'. If YearLength=1 and Years.sim=1000, the simulation has 1000 Deltas.  If YearLength=52 and Years.sim=10, the sim has 520 Deltas.
Years.sim <- 10                                                                                              #set default length of simulation in years
Years.sim.R <- findOneLine(SearchTxt="Years.sim", Headers=row.names(Controls), call.section="#1.3a Years.sim", file="Controls.txt", Var.name="Years.sim", domain="row", ZeroOK=TRUE)  #find row.header with 'Years.sim' in it
if(!is.na(Years.sim.R))                                                                                      #If the Years.sim row is found:-
   {Temp <- Controls[Years.sim.R, 1]                                                                         #store the supplied value temporarily and check that it is OK
   ERR <- is.DatumOK(Temp, "Years.sim (length of simulation in yrs)", NA.OK=FALSE, CharOK=FALSE, NumbOK=TRUE, LogicalOK=FALSE, nonIntegerOK=FALSE, InfinOK=FALSE, MinNotOK=0)#test the value
   if(ERR==TRUE)                                                                                             #If ERR=TRUE:-
      {Advise(WithRefTo="Years.sim in column 1 of Controls.txt", ErrText=paste0("The supplied value for Years.sim is ", Temp, " years.  This is not an integer in the acceptable range:- 0 < Years.sim < Infinity"), AdvText=paste0("Adjust the value given OR remove the Years.sim row in Controls.txt => default = ", Years.sim, ".\n\tYears.sim sets the duration of the simulation in nominal years, where each 'year' has YearLength simulation steps (Deltas)."), ShowData=NULL, CallSection="#1.3b", Fatal=TRUE, MinAdviceLevel2Show=1)
      }
   Years.sim <- Temp                                                                                         #no error found in Temp; overwrite default Years.sim
   }
N.LPproj <- Years.sim * YearLength                                                                           #number of LP projections required
#----------------------------------------------------------------


#1.4 Order to use (1st, 2nd, 3rd) for Taylor-series projections of self loss
#proj.SLorder=3 was found to provide the most credible simulation series though, if there are concerns that some constraints are seriously affecting projected values, proj.SLorder=1 could be tried for comparisons.
proj.SLorder <- 3                                                                                            #default Taylor-series order for calculating projected self losses generally
proj.SLorder.R <- findOneLine(SearchTxt="order", Headers=row.names(Controls), call.section="#1.4a proj.SLorder", file="Controls.txt", Var.name="proj.SLorder", domain="row", ZeroOK=TRUE) #find row.header with 'order' in it
if(!is.na(proj.SLorder.R))                                                                                   #If proj.SLorder.R is found:-
   {Temp <- Controls[proj.SLorder.R, 1]                                                                      #store the supplied value temporarily
   ERR <- is.DatumOK(Temp, "Taylor Series order for proj.SLorder", NA.OK=FALSE, CharOK=FALSE, NumbOK=TRUE, LogicalOK=FALSE, nonIntegerOK=FALSE, MustBeInSet=c(1, 2, 3))
   if(ERR==TRUE)                                                                                             #If ERR=TRUE:-
      {Advise(WithRefTo="proj.SLorder in column 1 of Controls.txt", ErrText=paste0("The supplied value for proj.SLorder is ", Temp, " .  This is not an integer in the acceptable set:- {1, 2, 3}.  \n\tAdjust the value given OR remove the proj.SLorder row in Controls.txt -> default = 1."), AdvText="proj.SLorder sets the order of Taylor-series used to project self losses of wild and NLO components for the simulation. \n\tproj.SLorder=1 is recommended for general use because 1st-order self loss does not depend on activity of constraints during the simulation.  \n\tHowever more accurate results can be obtained with proj.SLorder=3 for the minimal model with GP.a, a.pGP.h, a.FOOD.h and FOOD.h only.", ShowData=NULL, CallSection="#1.4b", Fatal=TRUE, MinAdviceLevel2Show=0)
      }
   proj.SLorder <- Temp                                                                                      #no error found in Temp; overwrite default proj.SLorder
   }
#----------------------------------------------------------------


#1.5 Progress reporting interval:- Progress.rep
#Sets the interval for reporting on screen how many Delta projections have been computed, and how much CPU time they have taken, during a simulation
Progress.rep <- 500                                                                                          #set default; units are Deltas
Progress.rep.R <- findOneLine(SearchTxt="Progr", Headers=row.names(Controls), call.section="#1.6a Progress.rep", file="Controls.txt", Var.name="Progress.rep", domain="row", ZeroOK=TRUE)  #find row.header with 'Progr' in it
if(!is.na(Progress.rep.R))                                                                                   #If the Progress.rep. row is found:-
   {Temp <- Controls[Progress.rep.R, 1]                                                                      #store the supplied value temporarily
   ERR <- is.DatumOK(Temp, "Progress reporting interval", NA.OK=TRUE, CharOK=FALSE, NumbOK=TRUE, LogicalOK=FALSE, InfinOK=TRUE, nonIntegerOK=FALSE, MinNotOK=0)
   if(ERR==TRUE)                                                                                             #If ERR=TRUE:-
      {Advise(WithRefTo="Progress.rep in column 1 of Controls.txt", ErrText=paste0("The supplied value for Progress.rep is ", Temp, " .  This is not NA or an integer in the acceptable range:- 1 <= Progress.rep < infinity"), AdvText=paste0("Adjust the value given OR remove the Progress.rep row in Controls.txt => default = ", Progress.rep, ".\n\tProgress.rep sets the number of simulation steps that must occur before a progress report is screened.  \n\tReports may be of low interest for simple, quick simulations."), ShowData=NULL, CallSection="#1.6b", Fatal=FALSE, MinAdviceLevel2Show=0)
      } else {Progress.rep <- Temp}                                                                          #error, if found in Temp, prevents overwrite of default Progress.rep
   }
#----------------------------------------------------------------

   
#1.6 Switch for displaying Params.file:- View.params
#Allows you to see how ECOLPS has processed parameter values that you supplied in Params.file
View.params <- 0                                                                                             #set default. 0 => don't view
View.params.R <- findOneLine(SearchTxt="View[.]pa", Headers=row.names(Controls), call.section="#1.7a", file="Controls.txt", Var.name="View.params", domain="row", ZeroOK=TRUE)  #find row.header with 'View.par' in it
if(!is.na(View.params.R))                                                                                    #If View.params.R is found:-
   {Temp <- Controls[View.params.R, 1]                                                                       #store the supplied value temporarily
   ERR <- is.DatumOK(Temp, "Parameters.file display switch", NA.OK=TRUE, CharOK=FALSE, NumbOK=TRUE, LogicalOK=FALSE, nonIntegerOK=FALSE, MustBeInSet=c(0, 1))
   if(ERR==TRUE)                                                                                             #If ERR=TRUE:-
      {Advise(WithRefTo="View.params in column 1 of Controls.txt", ErrText=paste0("The supplied value for View.params is ", Temp, " .  This is not an integer in the acceptable set:- {0, 1}."), AdvText=paste0("Adjust the value given OR remove the View.params row in Controls.txt => default = ", View.params, ".\n\tView.params = 1 displays a table of parameters read from the Params.file as processed by ECOLPS.  V.p = 0 omits the table.\n\tThe table should be checked to ensure that data have been read and interpreted as expected."), ShowData=NULL, CallSection="#1.7b", Fatal=FALSE, MinAdviceLevel2Show=0)
      } else {View.params <- Temp}                                                                           #error, if found in Temp, prevents overwrite of default View.params
   }
#----------------------------------------------------------------


#1.7 Switch for displaying Environment.file (if present):- View.Envir
#Allows you to see how ECOLPS has processed values of variables in Envir.file, if supplied
View.Envir <- 0                                                                                              #set default. 0 => don't view
View.Envir.R <- findOneLine(SearchTxt="View[.]env", Headers=row.names(Controls), call.section="#1.8a", file="Controls.txt", Var.name="View.Envir", domain="row", ZeroOK=TRUE)  #find row.header with 'View.env' in it
if(!is.na(View.Envir.R))                                                                                     #If View.Envir.R is found:-
   {Temp <- Controls[View.Envir.R, 1]                                                                        #store the supplied value temporarily
   ERR <- is.DatumOK(Temp, "Environment.file display switch", NA.OK=TRUE, CharOK=FALSE, NumbOK=TRUE, LogicalOK=FALSE, nonIntegerOK=FALSE, MustBeInSet=c(0, 1))
   if(ERR==TRUE)                                                                                             #If ERR=TRUE:-
      {Advise(WithRefTo="View.Envir in column 1 of Controls.txt", ErrText=paste0("The supplied value for View.Envir is ", Temp, " .  This is not an integer in the acceptable set:- {0, 1}."), AdvText=paste0("Adjust the value given OR remove the View.Envir row in Controls.txt => default = ", View.Envir, ".\n\tView.vars = 1 displays a table of date.days and values read from an Environment.file, if supplied. V.e. = 0 omits.  \n\tThese can be used to alter values of environmental variables or biomasses at the given date.days. \n\tThe table should be checked to ensure that data have been read and interpreted as expected."), ShowData=NULL, CallSection="#1.8b", Fatal=FALSE, MinAdviceLevel2Show=0)
      } else {View.Envir <- Temp}                                                                            #error, if found in Temp, prevents overwrite of default View.Envir
   }
#----------------------------------------------------------------


#1.8 Switch for displaying Controls.file:-  View.controls
#Allows you to see how ECOLPS has processed simulation control variables that you set in Controls.txt
View.controls <- 0                                                                                           #set default. 0 => don't view
View.controls.R <- findOneLine(SearchTxt="View[.]con", Headers=row.names(Controls), call.section="#1.9a", file="Controls.txt", Var.name="View.controls", domain="row", ZeroOK=TRUE)  #find row.header with 'View.con' in it
if(!is.na(View.controls.R))                                                                                  #If View.controls.R is found:-
   {Temp <- Controls[View.controls.R, 1]                                                                     #store the supplied value temporarily
   ERR <- is.DatumOK(Temp, "Controls.txt display switch", NA.OK=TRUE, CharOK=FALSE, NumbOK=TRUE, LogicalOK=FALSE, nonIntegerOK=FALSE, MustBeInSet=c(0, 1))
   if(ERR==TRUE)                                                                                             #If ERR=TRUE:-
      {Advise(WithRefTo="View.controls in column 1 of Controls.txt", ErrText=paste0("The supplied value for View.controls is ", Temp, " .  This is not an integer in the acceptable set:- {0, 1}."), AdvText=paste0("Adjust the value given OR remove the View.controls row in Controls.txt => default = ", View.controls, ".\n\tView.controls = 1 displays a table of the supplied variables that control simulations and output of ECOLPS.  \n\tThe table is read from Controls.txt. It should be checked, especially if ECOLPS is not operating as expected."), ShowData=NULL, CallSection="#1.9b", Fatal=FALSE, MinAdviceLevel2Show=0)
      } else {View.controls <- Temp}                                                                         #error, if found in Temp, prevents overwrite of default View.controls
   }
#----------------------------------------------------------------


#1.9 Switch for displaying the component-type table constructed from harvest and other coefficients:- View.compType
#Allows you to see how ECOLPS has translated values of parameters that you supplied in Params.file into component types (autotroph, fishery, NLO, etc).
#Interpretation of parameter values to give component types is performed in #3.
View.compType <- 0                                                                                           #set default. 0 => don't view
View.compType.R <- findOneLine(SearchTxt="View[.]comp", Headers=row.names(Controls), call.section="#1.10a", file="Controls.txt", Var.name="View.compType", domain="row", ZeroOK=TRUE)  #find row.header with 'View.comp' in it
if(!is.na(View.compType.R))                                                                                  #If View.compType.R is found:-
   {Temp <- Controls[View.compType.R, 1]                                                                     #store the supplied value temporarily
   ERR <- is.DatumOK(Temp, "Component.table viewing switch", NA.OK=TRUE, CharOK=FALSE, NumbOK=TRUE, LogicalOK=FALSE, nonIntegerOK=FALSE, MustBeInSet=c(0, 1))
   if(ERR==TRUE)                                                                                             #If ERR=TRUE:-
      {Advise(WithRefTo="View.compType in column 1 of Controls.txt", ErrText=paste0("The supplied value for View.compType is ", Temp, " .  This is not an integer in the acceptable set:- {0, 1}."), AdvText=paste0("Adjust the value given OR remove the View.compType row in Controls.txt => default = ", View.compType, ".\n\tView.compType = 1 displays a table of the component types prepared by ECOLPS from the values of parameters given in the Params.file.  \n\tThe table should be checked to ensure that ECOLPS has assigned component types to components named in the column headers, as intended."), ShowData=NULL, CallSection="#1.10b", Fatal=FALSE, MinAdviceLevel2Show=0)
     } else {View.compType <- Temp}                                                                          #error, if found in Temp, prevents overwrite of default View.compType
   }
#----------------------------------------------------------------


#1.10 Simulation number from which internal calculations are screened for EVERY projection:- ShowCalcsFromStep
#This may be useful for checking internal calculations or explaining simulation results. The output on screen is voluminous so start near the end or insert in #8. a line like: if(M>=xx) stop()
#The output is put above the first MainMenu.  The results shown can be altered by editing ShowCalcs() in ECOLPSfunctions.R
ShowCalcsFromStep <- Inf                                                                                     #default value = Inf = never show internal calculations
ShowCalcsFromStep.R <- findOneLine(SearchTxt="ShowCalcs", Headers=row.names(Controls), call.section="#1.11a", file="Controls.txt", Var.name="ShowCalcsFromStep", domain="row", ZeroOK=TRUE)  #find row.header with 'ShowCalcs' in it
if(!is.na(ShowCalcsFromStep.R))                                                                              #If ShowCalcs is found,
   {Temp <- Controls[ShowCalcsFromStep.R, 1]                                                                 #store the supplied value temporarily
   ERR <- is.DatumOK(Temp, "Internal-calculations viewing switch", NA.OK=TRUE, CharOK=FALSE, NumbOK=TRUE, LogicalOK=FALSE, nonIntegerOK=FALSE, MinOK=0, InfinOK=TRUE)
   if(ERR==TRUE)                                                                                             #If ERR=TRUE:-
      {Advise(WithRefTo="ShowCalcsFromStep in column 1 of Controls.txt", ErrText=paste0("The supplied value for ShowCalcsFromStep is ", Temp, " .  This is not an integer in the acceptable range:-  0 <= ShowCalcsFromStep <= infinity."), AdvText="Adjust the value given OR remove the ShowCalcsFromStep row in Controls.txt => default = Inf\n\tShowCalcsFromStep = 100, for example, displays values calculated for each simulation step of biomasses from  step 100.  \n\tThis permits internal calculations to be checked or may help explain simulation results. To prevent voluminous, unmanageable output, \n\tset ShowCalcsFromStep close to the end of the simulation, if necessary reducing Years.sim to bring the end temporarily forward.", ShowData=NULL, CallSection="#1.11b", Fatal=TRUE, MinAdviceLevel2Show=0)
      }
   ShowCalcsFromStep <- Temp                                                                                 #no error found in Temp; overwrite default ShowCalcsFromStep
   }
#----------------------------------------------------------------


#1.11 Graph window height
#Refers to the height of the graph window used to plot simulated time series; adjust it for your screen
#WinHeight <= MaxWinHeight is adjustable in graph menus with the W option. MWH depends on screen size to avoid folding over
WinHeight <- 10; MaxWinHeight <- 15                                                                          #default heights in "inches".
WH.R <- findOneLine(SearchTxt="WinHeight", Headers=row.names(Controls), call.section= "#1.12a", file="Controls.txt", Var.name="WinHeight", domain="row", ZeroOK=TRUE)  #find row.header with 'WinHeight' in it
if(!is.na(WH.R))                                                                                             #If WinHeight is found:-
   {Temp <- Controls[WH.R, 1]                                                                                #store the supplied value temporarily
   ERR <- is.DatumOK(Temp, "HEIGHT for graph windows", NA.OK=TRUE, CharOK=FALSE, NumbOK=TRUE, LogicalOK=FALSE, nonIntegerOK=TRUE, MaxOK=MaxWinHeight, MinOK=2) #check validity
   if(ERR==TRUE)                                                                                             #If ERR=TRUE:-
      {Advise(WithRefTo="WinHeight in column 1 of Controls.txt", ErrText=paste0("The supplied value for WinHeight is ", Temp, " .  This is not in the acceptable range:-  2 <= WinHeight <= ", MaxWinHeight, "."), AdvText=paste0("Adjust the value given OR remove the WinHeight row in Controls.txt => default = ", WinHeight, ".\n\tWinHeight sets the height of time-series graph windows in nominal inches. \n\tWindow height can be adjusted up to ", MaxWinHeight, " using the 'W' option in output menus. \n\tAdjust MaxWinHeight in code for screen height to prevent reflection."), ShowData=NULL, CallSection="#1.12b", Fatal=TRUE, MinAdviceLevel2Show=0)
      }
   WinHeight <- Temp                                                                                         #no error found in Temp; overwrite default maxHeight
   }
#----------------------------------------------------------------   

       
#1.12 Graph window width
#Refers to the width of the graph window used to plot simulated time series; adjust it for your screen
#WinWidth <= MaxWinWidth is adjustable in graph menus with the W option. MWW depends on screen size to avoid folding over
WinWidth  <- 10 ; MaxWinWidth <- 25                                                                          #default widths in "inches".
WW.R <- findOneLine(SearchTxt="WinWidth", Headers=row.names(Controls), call.section= "1.13a", file="Controls.txt", Var.name="WinWidth", domain="row", ZeroOK=TRUE)  #find row.header with 'WinWidth' in it
if(!is.na(WW.R))                                                                                             #If WinWidth is found:-
   {Temp <- Controls[WW.R, 1]                                                                                #store the supplied value temporarily
   ERR <- is.DatumOK(Temp, "WIDTH for graph windows", NA.OK=TRUE, CharOK=FALSE, NumbOK=TRUE, LogicalOK=FALSE, nonIntegerOK=TRUE, MaxOK=MaxWinWidth, MinOK=5) #check validity
   if(ERR==TRUE)                                                                                             #If ERR=TRUE:-
      {Advise(WithRefTo="WinWidth in column 1 of Controls.txt", ErrText=paste0("The supplied value for WinWidth is ", Temp, " .  This is not in the acceptable range:-  5 <= WinWidth <= ", MaxWinWidth, "."), AdvText=paste0("Adjust the value given OR remove the WinWidth row in Controls.txt => default = ", WinWidth, ".\n\tWinWidth sets the width of time-series graph windows in nominal inches. \n\tWindow width can be adjusted up to ", MaxWinWidth, " using the 'W' option in output menus. \n\tAdjust MaxWinWidth in code for screen width to prevent reflection."), ShowData=NULL, CallSection="#1.13b", Fatal=TRUE, MinAdviceLevel2Show=0)
      }
   WinWidth <- Temp                                                                                          #no error found in Temp; overwrite default maxWidth
   }
#----------------------------------------------------------------   


#1.13 Graph-key parameters: KeyItemWidth
#Keys on graphs can be adjusted to allow more items per line or longer item names, given that the key should occupy as few lines as possible so as to leave most space for the graph itself
#Automatic key sizing, the default, depends on KeyItemWidth and KeyLineLength; it works OK for the published Trial sets      
#Refer to Plot.results() section #PR1 in ECOLPSfunctions.R for automatic setting of key dimensions; K option of Constraints and Variables menus selects auto or manual key sizing 
#KeyItemWidth sets how many characters are allowed for the name of each plotted variable, an = sign, and minimal spaces between; short component names prevent confusing truncations in the key.
#KeyItemWidth= 18 is suggested for 3 items per line in a Windows(width=7, height=7) graph; 14 is suggested for 4 items/line
KeyItemWidth <- 22                                                                                           #default width of one key item as a number of characters.  Depends on window width and variables chosen
KIW.R <- findOneLine(SearchTxt="KeyItemW", Headers=row.names(Controls), call.section= "#1.14a", file="Controls.txt", Var.name="KeyItemWidth", domain="row", ZeroOK=TRUE)  #find row.header with 'KeyItemW' in it
if(!is.na(KIW.R))                                                                                            #If KIW.R is found:-
   {Temp <- Controls[KIW.R, 1]                                                                               #store the supplied value temporarily
   ERR <- is.DatumOK(Temp, "KeyItemWidth for graphs", NA.OK=TRUE, CharOK=FALSE, NumbOK=TRUE, LogicalOK=FALSE, nonIntegerOK=FALSE, MaxOK=50, MinOK=1) #check validity
   if(ERR==TRUE)                                                                                             #if ERR=TRUE:-
      {Advise(WithRefTo="KeyItemWidth in column 1 of Controls.txt", ErrText=paste0("The supplied value for KeyItemWidth is ", Temp, " .  This is not an integer in the acceptable range:-  1 <= KeyItemWidth <= 50."), AdvText=paste0("Adjust the value given OR remove the KeyItemWidth row in Controls.txt => default = ", KeyItemWidth, ".\n\tKeyItemWidth sets the number of characters allotted to each key item identifying a simulated time series in a graph.  \n\tToo short causes truncated identifiers; too long causes the total key to need too many lines, cramping the graph above.  \n\tThe default often works for simple simulations."), ShowData=NULL, CallSection="#1.14b", Fatal=TRUE, MinAdviceLevel2Show=0)
     }
   KeyItemWidth <- Temp                                                                                      #no error found in Temp; overwrite default KeyItemWidth
   }
#----------------------------------------------------------------   

   
#1.14 Graph-key parameters: KeyLineLength
#KeyLineLength can be adjusted to allow more characters in each line of the key for a time-series graph; this will depend on the window width, see #1.12, set for the graph
KeyLineLength <- WinWidth * 8                                                                                #default length of each line of the key (number of characters); the constant worked for me
KLL.R <- findOneLine(SearchTxt="KeyLineL", Headers=row.names(Controls), call.section="#1.15a", file="Controls.txt", Var.name="KeyLineLength", domain="row", ZeroOK=TRUE)  #find row.header with 'KeyLineL' in it
if(!is.na(KLL.R))                                                                                            #If KLL.R is found:-
   {Temp <- Controls[KLL.R, 1]                                                                               #store the supplied value temporarily
   ERR <- is.DatumOK(Temp, "KeyLineLength for graphs", NA.OK=TRUE, CharOK=FALSE, NumbOK=TRUE, LogicalOK=FALSE, nonIntegerOK=FALSE, MaxOK=WinWidth * 12, MinOK=20) #check validity; WWx12 worked for me
   if(ERR==TRUE)                                                                                             #If ERR=TRUE:-
      {Advise(WithRefTo="KeyLineLength in column 1 of Controls.txt", ErrText=paste0("The supplied value for KeyLineLength is ", Temp, " .  This is not an integer in the acceptable range:-  1 <= KeyLineLength <= WinWidth * 12."), AdvText=paste0("Adjust the value given OR remove the KeyLineLength row in Controls.txt => default = ", WinWidth * 8, ".\n\tKeyLineLength sets the number of characters allotted to each key item identifying a simulated time series in a graph.  \n\tToo short causes truncated identifiers; too long causes the total key to need too many lines, cramping the graph above.  \n\tThe default often works for simple simulations."), ShowData=NULL, CallSection="#1.15b", Fatal=TRUE, MinAdviceLevel2Show=0)
      }
   KeyLineLength <- Temp                                                                                     #no error found in Temp; overwrite default KeyItemWidth
   }
#----------------------------------------------------------------   


#1.15 Parameters file name
#The parameters file is required.  Its columns are the components used by ECOLPS; its rows are the parameters.  Entries are values, always NA if no value is appropriate
Params.file.R <- findOneLine(SearchTxt="Paramet", Headers=row.names(Controls), call.section="#1.16a", file="Controls.txt", Var.name="Parameters.file", domain="row", ZeroOK=FALSE)  #find row.header with 'Paramet' in it
if(!is.na(Params.file.R))                                                                                    #If one Params.file.Row is found in the Controls:-
   {Temp <- as.character(Controls[Params.file.R, 2])                                                         #temporarily store the string supplied in the 2nd col
   ERR <- is.DatumOK(Temp, "Parameters.file", NA.OK=FALSE, CharOK=TRUE,  NumbOK=FALSE, LogicalOK=FALSE, MustBeFile=TRUE)#set ERR flag if value not a valid file name
   if(ERR == FALSE) Params.file <- Temp                                                                      #If all checks are passed, store file ref permanently
   } else                                                                                                    #Else, no Params.file.R is found:-
   {ERR <- TRUE                                                                                              #set ERR flag
   }
if(ERR==TRUE)                                                                                                #If ERR=TRUE, i.e. no file ref row was found, or supplied value is not a file.ref:-
   {Advise(WithRefTo="Parameters.fileref in column 2 of Controls.txt", ErrText=paste0("The supplied value for Params.file is ", Temp, " .  \n\tThis is not a valid file reference."), AdvText="The file reference must be corrected. \n\tNA must be in the 1st col, and a file reference (a string without quotes, eg. C:/parameters.txt) in the 2nd col.\n\tThe Parameters.file is essential for supplying all initial values for parameters of ecological components.", ShowData=NULL, CallSection="#1.16b", Fatal=TRUE, MinAdviceLevel2Show=0)                                     #terminate the program
   }
#----------------------------------------------------------------   
                                                          

#1.16 Environment file name, if supplied
#The Envir.file is optional. Its purpose is to change parmeters or migrate biomasses between components during the simulation. It has 5 standard columns.  Rows are dates as days, ie. 365 per year.
#See #5. where Envir.file is read and processed
Environ.file <- NA                                                                                           #initialise with NA because Environ.file is optional
Environ.file.R <- findOneLine(SearchTxt="Envir[.]", Headers=row.names(Controls), call.section="#1.17a", file="Controls.txt", Var.name="Environment.file", domain="row", ZeroOK=TRUE)  #find row.header with 'Envir' in it
if(!is.na(Environ.file.R))                                                                                   #If Environ.file.R is found:-
   {Temp <- as.character(Controls[Environ.file.R, 2])                                                        #temporarily store the string supplied in the 2nd col of Controls.txt
   if(!is.na(Temp))                                                                                          #if the file name is not NA, check that it is a valid file ref:-
      {ERR <- is.DatumOK(Temp, "Environment.file", NA.OK=FALSE, CharOK=TRUE,  NumbOK=FALSE, LogicalOK=FALSE, MustBeFile=TRUE)#set ERR if value is not a valid file name
      if(ERR==TRUE)                                                                                          #If ERR=TRUE, i.e. the supplied value is not a file.ref:-
         {Advise(WithRefTo="Environment.fileref in column 2 of Controls.txt", ErrText=paste0("The supplied value for Environ.file is ", Temp, " .  This is not a valid file reference."), AdvText="Correct the file reference  OR remove the optional Envir.file row in Controls.txt. \n\tNA must be in the 1st col.  The 2nd col may either hold a file reference (a string without quotes, eg. C:/Environ.txt) OR be NA.\n\tAn Environment.file permits changes to initial values of parameters, and/or transfers of biomass between components, on given dates.", ShowData=NULL, CallSection="#1.17b", Fatal=TRUE, MinAdviceLevel2Show=0)
         }
      }   
   Environ.file <- Temp                                                                                      #no error found in Temp; store the Envir file name
   }                                                             
#----------------------------------------------------------------   
#################################################################End of section 1. PROGRAM CONTROLS


#2. READING AND IDENTIFICATION OF COMPONENTS AND THEIR ATTRIBUTES
#This section reads and extensively checks the unsorted columns of Params.file into Params0, then processes Params0 into named arrays of components, parameters, and logical variables. Most of their names have 0 suffixed to indicate "as read" and "unsorted". They are finalised in #3. #Checks are carried out to minimise occurrences of generalised R error messages and to standardise user inputs.
#The type of component (autotroph, habitat, etc) represented by each column of Params0 is determined from the presence or absence of positive values of specific parameters supplied. 0s are equivalent to NA for the purpose of classifying to type. If you would like a value to be 0, put in a very small positive number instead, eg. 0.000001, to prevent misclassification. Most NAs in the input must be changed to 0s to permit matrix arithmetic.
#Unwanted components in the Params.file can be removed temporarily by changing the column name to NA; thus the effects of a component are easily checked
#Only one NLO and one H2O component are allowed in this version of ECOLPS to avoid complicating physical transfers. Code sometimes allows for multiple NLO and H2O col(s) which may assist any later developments.
#Row names in the Params.file may include dot-suffixes to specify identity, eg relating to a food.  Suffixes must be spelled exactly everywhere, to permit matchings during computations. Non-matches are flagged as errors.
#Arrays and variables are defined even for components and nutrients not present in the supplied ecosystem model; these dummies prevent R errors if called needlessly, eg as outputs.
#Redundant rows and cols filled with zeros are often included in stored arrays. This allows the same row and col indexing variable to be used in multiple ways, thereby greatly simplifying code.
   #SUBSECTIONS
       #2.0 Read the parameters file to Params0 array
       #2.1 Self-growth coeffs; positive values define Autotrophs
       #2.2 SelfLoss coeffs; positive values define wild and non-living organic components (NLO)
       #2.3 Harv coeffs; positive values define harvesters, ie. heterotrophs and fisheries
       #2.4 Assimilation coeffs (efficiencies)
       #2.5 Organic biomasses of wild and NLO components:- Bmass
       #2.6 Maximum total growth rate of a wild component not dependent on food types:- Gmax
       #2.7 Maximum total harvest rate of a wild component not dependent on food types:- Hmax
       #2.8 Fishing effort
       #2.9 Fisheries quota
       #2.10 Essential habitat sizes
       #2.11 Essential habitat occupancies
       #2.12 Temperature of water
       #2.13 Light in water
       #2.14 Nutrient masses in H2O, and NLO (if in model)
       #2.15 Nutrient concentrations:- Nconc
       #2.16 Particulate SelfLoss coeffs
       #2.17 Light-dependent adjustments of maximum SelfGrowth rates of autotrophs
       #2.18 Temperature bands within which positive SelfGrow rates occur
       #2.19 Temperature-dependent adjustments of harvesting rates by heterotrophs
       #2.20 Temperature band within which positive harvest rates occur


#2.0 Read the parameters file to Params0 array
if(AL >= 0) cat("\nADVICE:-  The Params.file found in Controls.txt is: ", Params.file, "\n")                 #display the file name as a screen record of the source of data
Params0 <- read.table(file=Params.file, header=TRUE, row.names=1, as.is=TRUE)                                #Read data. [In R: as.is=T => don't create factors. row.names=1 => use col 1 for row.names]
Params0 <- array(data=unlist(Params0), dim=dim(Params0), dimnames=dimnames(Params0))                         #unlist() removes the automatically assigned data.frame class which can obstruct arithmetic code
UnwantedCols <- grep("NA.", colnames(Params0))                                                               #R changes NA colnames:- the 1st 'NA' to 'NA.', 2nd to 'NA..1", etc; grep() finds col names including "NA.", "NA.."
if(length(UnwantedCols) > 0) Params0 <- Params0[ , -UnwantedCols]                                            #remove any unwanted columns
Components0 <- colnames(Params0)                                                                             #component names are taken from the remaining columns
N.Components <- ncol(Params0)                                                                                #numbers of cols of Params0 = number of all components, wild or non-living
Rpar0.names <- rownames(Params0)                                                                             #rownames = parameter names in the Params0.file
#----------------------------------------------------------------


#2.1 Self-growth coeffs; positive values define Autotrophs
#The given values of SelfGrow are treated as maximum SG coeffs, unless variation of the coefficients with temperature or light over time is specified with an Environ.file (see #5.)
SearchText <- "SelfG"                                                                                        #text expected in row headers of SelfGrowth coeffs
SelfGrow0.Rpar <- findOneLine(SearchTxt=SearchText, Headers=Rpar0.names, call.section="#2.1a", file=Params.file, Var.name="SelfGrow", domain="row", ZeroOK=FALSE)  #find row.header with SearchText in it. Note: if 0 or more than 1 line is found, a fatal error message is delivered by findOneLine() 
if(is.na(SelfGrow0.Rpar)) Advise(WithRefTo=paste0("Absence of a SelfGrow row in \n\t", Params.file), ErrText="One row of SelfGrow coeffs is required for every simulation.  None has been found.", AdvText="SelfGrow coeffs must be in the range,  0 <= SelfGrow < infinity, or NA.\n\tPositive self growth (photosynthesis) identifies autotrophic components.  \n\tNon-autotrophic components should have SelfGrow = 0 or NA.\n\tAt least one autotroph is required in every simulation.", ShowData=NULL, CallSection="#2.1b", Fatal=TRUE, MinAdviceLevel2Show=1) 
for(Cpar in 1:N.Components)                                                                                  #For each col of the Params.file,
   {ERR <- is.DatumOK(Params0[SelfGrow0.Rpar, Cpar], "SelfGrow", NA.OK=TRUE, CharOK=FALSE, NumbOK=TRUE, LogicalOK=FALSE, MinOK=0, InfinOK=FALSE)#check coeff values
   if(ERR == TRUE)                                                                                           #If an invalid value is found:-
      {Advise(WithRefTo=paste0("SelfGrow in row ", SelfGrow0.Rpar, " column ", Cpar, " of ", Params.file), ErrText=paste0("A supplied value: SelfGrow = ", Params0[SelfGrow0.Rpar, Cpar], ", is not in the accepted range:  NA, or 0 <= SelfGrow < Inf. The value must be adjusted."), AdvText=paste0("One row of SelfGrowth coeffs must be supplied with row header including  '", SearchText, "'. \n\tSelf growth (photosynthesis) coefficients > 0 identify autotrophic components.  Non-autotrophs must have SelfGrow = 0 or NA.  \n\tAt least one autotroph is required in every simulation.  Note: Input values must be class(numeric); R may change input to \n\tclass(character) because of a single non-numeric in the column => this error.  Check the supplied row of data below:-\n\t"), ShowData=array(Params0[SelfGrow0.Rpar, ], dim=c(1, N.Components), dimnames=list("SelfGrow", Components0)), CallSection="#2.1c", Fatal=TRUE, MinAdviceLevel2Show=1)                                                                                      #The last advice, about R changing class, is necessary because ECOLPS will not go past the first wrong class
      }                                                                                                      #Advice level, AL given in Controls.txt, signals whether or not to show 'Advice' text. The rest is always shown
   }
SelfGrow0 <- array(data=Params0[SelfGrow0.Rpar, ], dim=c(1, N.Components), dimnames=list("SelfGrow0", Components0)) #store SelfGrow0 as a 1-row array, the full width of Params.file
SelfGrow0 <- ifelse(!is.na(SelfGrow0), SelfGrow0, 0)                                                         #swap NAs for 0s to permit matrix arithmetic later
is.SelfGrow0 <- SelfGrow0 > 0                                                                                #logical variable to identify col locations of positive values of SelfGrow0
#----------------------------------------------------------------


#2.2 SelfLoss coeffs; positive values define wild and non-living organic components (NLO)
#Wild and NLO components are both 'Self losers' (of organic mass); they are identified jointly here, then distinguished in #3.  Fisheries and other component types have 0 or NA as supplied SelfLoss coeffs
#SelfLoss coeffs are maximum rates assuming that exceptional conditions are not present, eg. pollution, lack of oxygen
SearchText <- "SelfLoss"                                                                                     #text expected in row headers of SelfLoss coeffs
SelfLoss0.Rpar <- findOneLine(SearchTxt=SearchText, Headers=Rpar0.names, call.section="#2.2a", file=Params.file, Var.name="SelfLoss", domain="row", ZeroOK=FALSE)  #find row.header of Params0 with SearchText in it 
if(is.na(SelfLoss0.Rpar)) Advise(WithRefTo=paste0("Absence of a SelfLoss row in \n\t", Params.file), ErrText="One row of SelfLoss coeffs is required for every simulation.  None has been found.", AdvText="SelfLoss coeffs must be in the range,  0 <= SelfLoss < infinity, or NA.\n\tPositive self Loss identifies wild and NLO components.  \n\tOther components should have SelfLoss = 0 or NA.\n\tAt least one autotroph and one heterotroph, 'wilds', are required in every simulation.", ShowData=NULL, CallSection="#2.2a", Fatal=TRUE, MinAdviceLevel2Show=0)
for(Cpar in 1:N.Components)                                                                                  #For each col of the Params.file,
   {ERR <- is.DatumOK(Params0[SelfLoss0.Rpar, Cpar], "SelfLoss", NA.OK=TRUE, CharOK=FALSE, NumbOK=TRUE, LogicalOK=FALSE, MinOK=0, MaxNotOK=2) #check coeffs for validity
   if(ERR == TRUE)                                                                                           #If an invalid value is found:-
      {Advise(WithRefTo=paste0("Invalid SelfLoss coeffs in row ", SelfLoss0.Rpar, " column ", Cpar, " of ", Params.file), ErrText=paste0("A supplied value: SelfLoss = ", Params0[SelfLoss0.Rpar, Cpar], ", is not in the accepted range:  NA, or 0 <= SelfLoss < 2 . The value must be adjusted."), AdvText=paste0("One row of SelfLoss coeffs must be supplied with row header including '", SearchText, "'.  SelfLoss coefficients > 0 identify both wild and \n\tnon-living organic (NLO) components.  At least two wild components (1 auto- & 1 heterotroph) are required for every simulation.  \n\tNLO is optional unless nutrients are in the model. Non-wild and non-NLO components have SelfLoss = 0 or NA.\n\t"), ShowData=array(Params0[SelfLoss0.Rpar, ], dim=c(1, N.Components), dimnames=list("SelfLoss", Components0)), CallSection="#2.2c", Fatal=TRUE, MinAdviceLevel2Show=1)
      }
   }
SelfLoss0 <- array(data=Params0[SelfLoss0.Rpar, ], dim=c(1, N.Components), dimnames=list("SelfLoss", Components0)) #create SelfLoss0 as a 1-row array, full width of Params.file initially
SelfLoss0 <- ifelse(!is.na(SelfLoss0), SelfLoss0, 0)                                                         #change NAs to zeroes
is.SelfLoss0 <- SelfLoss0 > 0                                                                                #logical variable to identify col locations of positive values of SelfLoss0
#----------------------------------------------------------------


#2.3 Harv coeffs; positive values define harvesters, ie. heterotrophs and fisheries
#These are maximum Harv coeffs, used by default unless any of the coefficients are reduced by sub-optimal temperatures at times specified using an Environ.file (see #5. )
#All autotrophs and NLO should have zero Harv coeffs because they do not harvest.  Phytoheterotrophs are NOT coded for
#All consumers must have a +ve Harv coeff AND a +ve Assim coeff to be classified as a consumer, including fisheries.
#Harvesters are not consumers if they don't assimilate what they harvest. ie. Their Assim=0. Their harvests are called unassimilated harvests (UAHs); they are not considered in Parts 1 or 2 of the write-up. UAHs must be estimated by Taylor-series projection because no Gross Production (GP), needed for LP, eventuates from them. Special code and matrices with rows for them included has been written in below but not tested extensively.
#Harv coeffs are given .suffixes in rows of the Params.file to indicate the food or UAH they refer to, eg. Harv.alga . Non-food, non-UAH components receive a row of zero Harv coeffs automatically in #4 if not supplied.
#Self harvesting, ie. cannibalism, has not been tested so Self-Harv coeffs should be set to zero in this version of ECOLPS to avoid possible over-writing or double counting. Or, if cannibalism is on juveniles, separate the juvs and adults into different components; adults could be recruited from juveniles by transferring a proportion of them with an Envir.file (code present in #5. but not thoroughly tested)

SearchText <- "Harv"                                                                                         #text expected in row headers of Harv coeffs
Harv0.Rpar <- grep(SearchText, ignore.case=T, Rpar0.names)                                                   #find the row headers in Params0 with the SearchText in them
N.Harv0.Rpar <- length(Harv0.Rpar)                                                                           #count the number of harvest rows given
#There must be at least 1 Harv row in Params.file,
if(N.Harv0.Rpar == 0) {Advise(WithRefTo=paste0("Absence of harvest coeff rows in \n\t", Params.file), ErrText="One row of Harv coeffs must be supplied for every harvested component. There must be at least one but none has been found. The file must be adjusted.", AdvText=paste0("Harv row-headers should include ' ", SearchText, ". ' with a food (or unassimilated-harvest, UAH) name suffixed; eg. 'Harv.alga3' .  \n\tEach food name must be spelled so as to exactly match a component column name. \n\tPositive harvest coeffs identify harvesters, namely heterotrophs and fisheries. \n\tAt least one harvesting grazer or predator is required for every simulation. \n\tIn this version of ECOLPS, autotrophs (SelfGrowth > 0) must not have any positive Harv coeffs in their column."), ShowData=NULL, CallSection="#2.3a", Fatal=TRUE, MinAdviceLevel2Show=1)
   }
Harv0.Rnames <- Rpar0.names[Harv0.Rpar]                                                                      #temporarily store Harv row names; Rpar0.names was created in #2.0
Harv0.suffixes <- substr(Harv0.Rnames, regexpr("[.]", Harv0.Rnames) +1, 100)                                 #get the .food or .UAH suffixes
#Check validity of supplied Harv coeffs:
for(Rharv0 in 1:N.Harv0.Rpar)                                                                                #For each row of Harv0,
   {#Check that Harv0.suffixes match component names supplied as column headers in Params.file
   ERR <- is.DatumOK(Harv0.suffixes[Rharv0], "Harvest suffixes", NA.OK=FALSE, CharOK=TRUE, NumbOK=FALSE, LogicalOK=FALSE, MustBeInSet=Components0)
   if(ERR == TRUE)
      {Advise(WithRefTo=paste0("Invalid suffixes for Harvest coefficients in row ", Harv0.Rpar[Rharv0], " of \n\t", Params.file), ErrText=paste0("A supplied Harvest suffix is ", Harv0.suffixes[Rharv0], " . This does not match any component name given in the file. The suffix must be adjusted."), AdvText="Harvest suffixes, written Harv.suffix, must match one of the components, supplied as column headers. \n\tThe suffixes identify food, and unassimilated, harvested components. \n\tCheck the component names found in the file, shown below:-\n", ShowData=Components0, CallSection="#2.3b", Fatal=TRUE, MinAdviceLevel2Show=2)
      }
   for(Cpar in 1:N.Components)                                                                              #For all potentially harvesting cols,
      {ERR <- is.DatumOK(Params0[Harv0.Rpar[Rharv0], Cpar], "Harvest coeff", NA.OK=TRUE, CharOK=FALSE, NumbOK=TRUE, LogicalOK=FALSE, MinOK=0, InfinOK=FALSE) #check supplied values
      if(ERR == TRUE)                                                                                        #If an invalid value is found:-
         {Advise(WithRefTo=paste0("Invalid harvest coeffs in row ", Harv0.Rpar[Rharv0], " column ", Cpar, " of \n\t", Params.file), ErrText=paste0("A supplied value for Harv is ", Params0[Harv0.Rpar[Rharv0], Cpar], " . This is not in the accepted ranges:  NA, or 0 <= Harv < infinity. The value must be adjusted."), AdvText="Positive Harv coeffs identify harvested wild and non-living organic (NLO) food components in rows, and their heterotrophic or fishery \n\tharvesters in columns. Unharvested rows of Harv, if provided in the Params.file, should have Harv = 0 or NA in all columns. \n\tCheck the input row below:-", ShowData=array(Params0[Harv0.Rpar[Rharv0],  ], dim=c(1, N.Components), dimnames=list(Harv0.Rnames[Rharv0], Components0))  , CallSection="#2.3c", Fatal=TRUE, MinAdviceLevel2Show=2)
         }
      }
   }                                                             
Harv0 <- array(data=Params0[Harv0.Rpar, ], dim=c(N.Harv0.Rpar, N.Components), dimnames=list(Harv0.Rnames, paste("by.", Components0, sep="")))#fill Harv0 array with the given rows
Harv0 <- ifelse(!is.na(Harv0), Harv0, 0)                                                                     #convert any NAs to 0 for arithmetic purposes
is.Harv0 <- Harv0 > 0                                                                                        #logical variable (a matrix) to identify row and col locations of positive values of Harv0
#----------------------------------------------------------------


#2.4 Assimilation coeffs (efficiencies)
#Assimilation efficiencies are proportions ranging from 0 to 1. Every food x consumer pair must have at least one +ve harvest paired with a +ve assim coeff. If no matching Assim, harvested component is a UAH (see #2.3)
#Assim = 1 for scavengers eating NLO because unassimilated NLO harvested by them is assumed to drop directly back to NLO
SearchText <- "Assim"                                                                                        #text expected in row header of Assim coeffs
Assim0.Rpar <- grep(SearchText, ignore.case=T, Rpar0.names)                                                  #find the row headers of Params0 with this SearchText in them
N.Assim0.Rpar <- length(Assim0.Rpar)                                                                         #count the number of assim rows given
#There must be 1 Assim row to match each supplied Harv row.  This will define foods and any UAH
if(N.Assim0.Rpar != N.Harv0.Rpar) {Advise(WithRefTo=paste0("Absence of assimilation coeff rows in \n\t", Params.file), ErrText="One row of Assim coeffs must be supplied for every food. None has been found. The file must be adjusted.", AdvText=paste0("Assim row headers should include ' ", SearchText, ". ' with a food name suffixed; eg. 'Assim.alga3' .  \n\tEach food name must be spelled so as to exactly match a component-column name. \n\tPositive assimilation coeffs identify consumers, namely heterotrophs and fisheries. \n\tAt least one consuming grazer or predator is required for every simulation."), ShowData=NULL, CallSection="#2.4a", Fatal=TRUE, MinAdviceLevel2Show=1)
   }
Assim0.Rnames <- Rpar0.names[Assim0.Rpar]                                                                    #temporarily store Assim row names
Assim0.suffixes <- substr(Assim0.Rnames, regexpr("[.]", Assim0.Rnames) +1, 100)                              #get the .food suffixes located after the dot in the name
for(Rassim0 in 1:N.Assim0.Rpar)                                                                              #For all potentially assimilated col comps in Params0, ie. rows in Assim
   {#Check that Assim0.suffixes match Harv0.suffixes read in #2.3
   ERR <- is.DatumOK(Assim0.suffixes[Rassim0], "Assimilation suffixes", NA.OK=FALSE, CharOK=TRUE, NumbOK=FALSE, LogicalOK=FALSE, MustBeInSet=Harv0.suffixes)
   if(ERR == TRUE)
      {Advise(WithRefTo=paste0("Invalid suffixes for Assimilation coefficients in row ", Assim0.Rpar[Rassim0], " of \n\t", Params.file), ErrText=paste0("A supplied Assim suffix is ", Assim0.suffixes[Rassim0], " . This does not match any Harv.suffix given in the file. The suffix must be adjusted."), AdvText="Assim suffixes, written Assim.suffix, must match one Harv.suffix, supplied as rows in the file. \n\tThe suffixes identify food (Assim>0), and unassimilated (Assim=0), harvested components. \n\tCheck the Harv.suffixes found in the file, shown below:-\n", ShowData=Harv0.suffixes, CallSection="#2.4b", Fatal=TRUE, MinAdviceLevel2Show=1)
      }
   for(Cpar in 1:N.Components)                                                                              #For all potentially assimilating col comps in Params0, ie. cols in Assim
      {ERR <- is.DatumOK(Params0[Assim0.Rpar[Rassim0], Cpar], "Assim coefficient", NA.OK=TRUE, CharOK=FALSE, NumbOK=TRUE, LogicalOK=FALSE, MinOK=0, MaxOK=1)#check supplied values
      if(ERR == TRUE)                                                                                        #If an invalid value is found:-
         {Advise(WithRefTo=paste0("Invalid assimilation coeffs in row ", Rassim0, " column ", Cpar, " of of \n\t", Params.file), ErrText=paste0("A supplied value for Assim is ", Params0[Assim0.Rpar[Rassim0], Cpar], " . This is not in the acceptable ranges: NA, or 0 <= Assim <= 1. The file must be adjusted."), AdvText="Assim coeffs measure the proportion of harvested biomass that is converted into GP of a consumer, or into landings of a fishery.  \n\t
 Positive Assim coeffs identify harvesters (columns) that are consumers of foods (rows). Unassimilated rows of Assim, if provided, should have \n\tAssim = 0 or NA in all columns. Positive Harv paired with zero Assim defines an unassimilated harvest (UAH).  \n\tProductive harvests are estimated by linear programming; unAssimilated harvests are estimated by projection using Taylor series.\n\tCheck the input row below:-", ShowData=array(Params0[Assim0.Rpar[Rassim0], ], dim=c(1, N.Components), dimnames=list(Assim0.Rnames[Rassim0], Components0)), CallSection="#2.4c", Fatal=TRUE, MinAdviceLevel2Show=1)
         }
      }
   }
Assim0 <- array(data=Params0[Assim0.Rpar, ], dim=c(N.Assim0.Rpar, N.Components), dimnames=list(Assim0.Rnames, paste("by.", Components0, sep="")))#fill preliminary Assim array
Assim0 <- ifelse(!is.na(Assim0), Assim0, 0)                                                                  #convert any NAs to 0 for arithmetic
#Put Assim0 into the same row order as Harv0
Assim00 <- array(data=0, dim=c(nrow2(Assim0, N.Components), N.Components))                                   #copy the Assim0 structure
ERR <- FALSE                                                                                                 #cancel any previous error
for(Rharv in 1:N.Harv0.Rpar)                                                                                 #For each row of the Harv0 matrix,
   {Rassim <- which(Assim0.suffixes == Harv0.suffixes[Rharv])                                                #find the matching row of the Assim0 matrix
   if(length(Rassim) == 1)  {Assim00[Rharv, ] <- Assim0[Rassim, ]} else {ERR <- TRUE}                        #If one row is found, move the Assim coef; else flag an error if 0 or > 1 rows are found
   }
Assim0[ , ] <- Assim00                                                                                       #overwrite Assim0 with the sorted Assim00
if(ERR == TRUE)                                                                                              #If an error is flagged:-
   {Advise(WithRefTo=paste0("Invalid Harv and Assim suffixes supplied in row names of ",  Params.file) , ErrText= "The sets of harvest- and assimilation-coefficient suffixes are not the same. Correct the row suffixes in Harv and Assim rows of the file. ", AdvText="ECOLPS matches the ordering of rows of harvest and assimilation coefficients in order to determine which harvests are assimilated \n\tso as to nourish the harvester. If the suffixes, referring to foods, cannot be matched exactly by order, the matching of coefficients \n\tmay be wrong. Check the input below. Above: suffixes in Assim, not in Harv.  Below: suffixes in Harv, not in Assim. :-", ShowData= list(setdiff(Assim0.suffixes, Harv0.suffixes), setdiff(Harv0.suffixes, Assim0.suffixes)),  CallSection="#2.4c", Fatal=TRUE, MinAdviceLevel2Show=0)
   }
is.Assim0 <- Assim0 > 0                                                                                      #logical variable (a matrix) to identify row and col locations of positive values of Assim0
#----------------------------------------------------------------


#2.5 Organic biomasses of wild and NLO components:- Bmass
#A Bmass row is required for every simulation
SearchText <- "Bmass"                                                                                        #text expected in row header for initial biomasses in Params0
Bmass0 <- array(data=0, dim=c(1, N.Components), dimnames=list("Bmass.init", Components0))                    #1-row array for initial biomasses; default data are 0s not NAs
Bmass0.Rpar <- findOneLine(SearchTxt=SearchText, Headers=Rpar0.names, call.section=NULL, file=Params.file, Var.name="Bmass.init", domain="row", ZeroOK=FALSE)  #find row.header with SearchText in it 
if(is.na(Bmass0.Rpar)) Advise(WithRefTo=paste0("Absence of a Biomass row in \n\t", Params.file), ErrText="One row of Biomasses is required for every simulation.  None has been found. The file must be adjusted", AdvText=paste0("Biomass row headers should include ' ", SearchText, ". 'Biomasses must be in the range, NA, or 0 <= Bmass < infinity."), ShowData=NULL, CallSection="#2.5c", Fatal=TRUE, MinAdviceLevel2Show=0) 
for(Cpar in 1:N.Components)                                                                                  #For each col of the Params.file,
   {ERR <- is.DatumOK(Params0[Bmass0.Rpar, Cpar], "Bmass", NA.OK=TRUE, CharOK=FALSE, NumbOK=TRUE, LogicalOK=FALSE, MinOK=0, MaxNotOK=Inf) #check coeffs for validity
   if(ERR == TRUE)                                                                                           #If an invalid value is found:-
      {Advise(WithRefTo=paste0("Invalid Bmass values in row ", Bmass0.Rpar, " column ", Cpar, " of ", Params.file), ErrText=paste0("A supplied value: Bmass = ", Params0[Bmass0.Rpar, Cpar], ", is not in the accepted range:  NA, or 0 =< Bmass < Inf. The value must be adjusted."), AdvText=paste0("One row of Bmass must be supplied with row header including '", SearchText, "'. Bmass > 0 identify both wild and \n\tnon-living organic (NLO) components.  At least two wild components (1 auto- & 1 heterotroph) are required for every simulation.  \n\tNLO is optional unless nutrients are in the model.\n\t"), ShowData=array(Params0[Bmass0.Rpar, ], dim=c(1, N.Components), dimnames=list("Bmass", Components0)), CallSection="#2.2c", Fatal=TRUE, MinAdviceLevel2Show=0)
      }
   }
Bmass0[1, ] <- Params0[Bmass0.Rpar , ]                                                                       #transfer initial biomasses from Params0 to their own row vector
Bmass0[1, ] <- ifelse(!is.na(Bmass0[1, ]), Bmass0[1, ], 0)                                                   #convert any NAs to zeros for non wild or NLO components
is.Bmass0 <- Bmass0 > 0                                                                                      #logical variable to identify col locations of positive values of Bmass0
#----------------------------------------------------------------


#2.6 Maximum total growth rate of a wild component not dependent on food types:- Gmax 
#Gmax is an optional parameter applied to heterotrophs to constrain their total GP from ALL of their foods
#Individual heteros not intended for control by a provided row of Gmax should have their Gmax=Inf or NA if the row is supplied in Params.file
Gmax0 <- array(data= Inf, dim=c(1, N.Components), dimnames = list("Gmax.coef", Components0))                 #default Gmax
SearchText <- "Gmax"
Gmax0.Rpar <- findOneLine(SearchTxt=SearchText, Headers=Rpar0.names, call.section="#2.7a", file=Params.file, Var.name="Gmax", domain="row", ZeroOK=TRUE)  #find row.header with 'Gmax' in it 
if(!is.na(Gmax0.Rpar))                                                                                       #If an Gmax row is present:-
   {for(Cpar in 1:N.Components)                                                                              #For each col of the Params.file,
      {ERR <- is.DatumOK(Params0[Gmax0.Rpar, Cpar], "Gmax", NA.OK=TRUE, CharOK=FALSE, NumbOK=TRUE, LogicalOK=FALSE, MinOK=0, MaxOK=Inf) #check coeffs for validity
      if(ERR == TRUE)                                                                                        #If an invalid value is found:-
         {Advise(WithRefTo=paste0("Invalid Gmax values in row ", Gmax0.Rpar, " column ", Cpar, " of ", Params.file), ErrText=paste0("A supplied value: Gmax = ", Params0[Gmax0.Rpar, Cpar], ", is not in the accepted range:  NA, or 0 < Gmax =< Inf. The value must be adjusted."), AdvText=paste0("One row of Gmax may optionally be supplied with row header including '", SearchText, "'. Gmax > 0 restricts the maximum total GP for \n\ta heterotrophic component from all of its foods.\n\t"), ShowData=array(Params0[Gmax0.Rpar, ], dim=c(1, N.Components), dimnames=list("Gmax", Components0)), CallSection="#2.7b", Fatal=TRUE, MinAdviceLevel2Show=0)
         }
      }
   Gmax0[1, ] <- Params0[Gmax0.Rpar , ]                                                                      #transfer Gmaxes from Params0 to their own row vector
   Gmax0[1, ] <- ifelse(!is.na(Gmax0[1, ]), Gmax0[1, ], Inf)                                                 #convert NAs to infinities
   }
is.Gmax0 <- Gmax0 < Inf                                                                                      #logical variable to identify col locations of sub-infinite values of Gmax
#----------------------------------------------------------------


#2.7 Maximum total harvest rate of a wild component not dependent on food types:- Hmax
#Hmax is an optional parameter applied to heterotrophs to constrain their total harvest of all foods. Hmax should be independent of Gmax to avoid duplicative effects; in that case, use Hmax=NA or omit Hmax input row
Hmax0 <- array(data= Inf, dim=c(1, N.Components), dimnames = list("Hmax.coef", Components0))                 #default Hmax
SearchText <- "Hmax"
Hmax0.Rpar <- findOneLine(SearchTxt=SearchText, Headers=Rpar0.names, call.section="#2.8a", file=Params.file, Var.name="Hmax", domain="row", ZeroOK=TRUE)  #find row.header with 'Hmax' in it 
if(!is.na(Hmax0.Rpar))                                                                                       #If an Hmax row is present:-
   {for(Cpar in 1:N.Components)                                                                              #For each col of the Params.file,
      {ERR <- is.DatumOK(Params0[Hmax0.Rpar, Cpar], "Hmax", NA.OK=TRUE, CharOK=FALSE, NumbOK=TRUE, LogicalOK=FALSE, MinOK=0, MaxNotOK=Inf) #check coeffs for validity
      if(ERR == TRUE)                                                                                        #If an invalid value is found:-
         {Advise(WithRefTo=paste0("Invalid Hmax values in row ", Hmax0.Rpar, " column ", Cpar, " of ", Params.file), ErrText=paste0("A supplied value: Hmax = ", Params0[Hmax0.Rpar, Cpar], ", is not in the accepted range:  NA, or 0 < Hmax < Inf. The value must be adjusted."), AdvText=paste0("One row of Hmax may optionally be supplied with row header including '", SearchText, "'. Hmax > 0 restricts the maximum total harvest by a \n\theterotrophic component of all of its foods.\n\t"), ShowData=array(Params0[Hmax0.Rpar, ], dim=c(1, N.Components), dimnames=list("Hmax", Components0)), CallSection="#2.8b", Fatal=TRUE, MinAdviceLevel2Show=0)
         }
      }
   Hmax0[1, ] <- Params0[Hmax0.Rpar , ]                                                                      #transfer Hmaxes from Params0 to their own row vector
   Hmax0[1, ] <- ifelse(!is.na(Hmax0[1, ]), Hmax0[1, ], Inf)                                                 #convert NAs to infinities
   }
is.Hmax0 <- Hmax0 < Inf                                                                                      #logical variable to identify col locations of sub-infinite values of Hmax
#----------------------------------------------------------------


#2.8 Fishing effort
#Fishing effort has one row of values. It is required if any fishery is in the model. Values in the fishery columns are initial effort values, modifiable after the first projection with an Envir.txt file (see #5)
Effort0 <- array(data= 0, dim=c(1, N.Components), dimnames = list("FishingEffort", Components0))             #default Effort
SearchText <- "Eff"
Effort0.Rpar <- findOneLine(SearchTxt=SearchText, Headers=Rpar0.names, call.section="#2.8a", file=Params.file, Var.name="Effort", domain="row", ZeroOK=TRUE)  #find row.header with 'Effort' in it 
is.Fishery0 <- rep(FALSE, N.Components)                                                                      #default
if(!is.na(Effort0.Rpar))                                                                                     #If an effort row is found:-
   {for(Cpar in 1:N.Components)                                                                              #For each col of the Params.file,
      {ERR <- is.DatumOK(Params0[Effort0.Rpar, Cpar], "Effort", NA.OK=TRUE, CharOK=FALSE, NumbOK=TRUE, LogicalOK=FALSE, MinOK=0, MaxNotOK=Inf) #check coeffs for validity
      if(ERR == TRUE)                                                                                        #If an invalid value is found:-
         {Advise(WithRefTo=paste0("Invalid fishing effort dat in row ", Effort0.Rpar, " column ", Cpar, " of ", Params.file), ErrText=paste0("A supplied value: Effort = ", Params0[Effort0.Rpar, Cpar], ", is not in the accepted range:  NA, or 0 =< Effort < Inf. The value must be adjusted."), AdvText=paste0("One row of Effort must be supplied with row header including '", SearchText, " ' if any fishery is in the model. Values in the fishery columns are initial effort values, modifiable later with an Envir.txt file. Below: Initial fishing effort, as read from the file:-\n\t"), ShowData=array(Params0[Effort0.Rpar, ], dim=c(1, N.Components), dimnames=list("FishingEffort", Components0)), CallSection="#2.8b", Fatal=TRUE, MinAdviceLevel2Show=0)
         }
      }
   Effort0[1, ] <- Params0[Effort0.Rpar , ]                                                                  #transfer Effort0 from Params0 to their own row vector
   Effort0[1, ] <- ifelse(!is.na(Effort0[1, ]), Effort0[1, ], 0)                                             #convert NAs to zeros
   }
is.Fishery0 <- Effort0 > 0                                                                                   #logical variable (matrix) to identify col locations of positive values of Effort; all FALSE if no fishery
Fisheries <- Components0[is.Fishery0]                                                                        #names of fisheries; this ordering of fishery column names takes precedence in the rest of the simulation
N.Fisheries <- sum(is.Fishery0)                                                                              #number of fisheries in the model by effort data; other fishery properties checked in #3.
#----------------------------------------------------------------


#2.9 Fisheries quota
#Quota may be 0 = no landings allowed, or Inf=no restrictions
#Initial quota, if supplied, are in 1 row per fishery in the Params.file; rows are re-ordered by fishery, if necessary, to match the ordering of fishery columns
#If any quota rows are supplied, 1 quota row is provided for every fishery whether or not restricting quota are present for any target component; the quota rows are ordered the same as the fisheries columns of Params.file. Some storage space is unused but coding is simpler without having to allow for all combinations of fisheries present, quota present, quota fished or not, etc. 
#All quota must be landable as shown by a positive Growth coefficient.  Those not landed give a fatal error at #
#PS. I sometimes use quotum for the singular, and quota for the plural (from Latin at school, yawn).
Quota0 <- array(data= NA, dim=c(1, N.Components), dimnames = list("Quota.NoFishery", Components0))           #default quota
is.QuotumSet0 <- rep(FALSE, N.Components)                                                                    #default logical variable to indicate col locations of sub-infinite quota
ParamsQuota0.Rnames <- NA; ParamsQuota0.suffixes <- NA                                                       #default quota row names and suffixes (ie fisheries) in Quota row names of Params
SearchText <- "Quot"                                                                                         #text expected in row header of Quota coeffs
Quota0.Rpar <- grep(SearchText, ignore.case=T, Rpar0.names)                                                  #find the row numbers of Params0 with this SearchText in their row headers
N.Quota0.Rpar <- length(Quota0.Rpar)                                                                         #count the number of quota rows given (whether or not they have any values < Inf)
if(N.Fisheries > 0 & N.Quota0.Rpar > 0)                                                                      #If fisheries with quota are present in the model,
   {Quota0 <- array(data= NA, dim=c(N.Fisheries, N.Components), dimnames = list(paste0("Quota.", Fisheries), Components0))#create quota rows, 1 per fishery whether having sub-infinite quota or not, ordered as fishery columns
   ParamsQuota0.Rnames <- row.names(Params0)[Quota0.Rpar]                                                    #names of quota rows in Params0
   ParamsQuota0.suffixes <- substr(ParamsQuota0.Rnames, regexpr("[.]", ParamsQuota0.Rnames) +1, 100)         #get the .fishery suffixes located after the dot in each name
   for(Qrow in 1:N.Quota0.Rpar)                                                                              #For each supplied quotum row,
      {ERR <- is.DatumOK(ParamsQuota0.suffixes[Qrow], "Quota suffixes", NA.OK=FALSE, CharOK=TRUE, NumbOK=FALSE, LogicalOK=FALSE, MustBeInSet=Fisheries)   #check that all supplied quota fisheries correspond to fishery components
      if(ERR == TRUE)                                                                                        #If not,
         {Advise(WithRefTo=paste0("Invalid suffixes for Quota values in row ", Quota0.Rpar[Qrow], " of \n\t", Params.file), ErrText=paste0("A supplied Quota suffix is ", ParamsQuota0.suffixes[Qrow], " . This does not match any component name identified as a fishery by positive fishing effort in the file. \n\tThe suffix must be adjusted."), AdvText="Quota suffixes, written Quota.suffix, must match one of the components, supplied as column headers, and having a positive value in the \n\trow for fishing effort. Suffixes identify the quota-controlled fishery. Remove unwanted quota with # at col 1 of quota.fishery row.  \n\tCheck names of fisheries in the model, shown below, for unmatched quota:-\n", ShowData=Fisheries, CallSection="#2.9a", Fatal=TRUE, MinAdviceLevel2Show=1)
         }  
      for(Cpar in 1:N.Components)                                                                            #For each component possibly landed by this quota fishery,
         {ERR <- is.DatumOK(Params0[Quota0.Rpar[Qrow], Cpar], "Quotum", NA.OK=TRUE, CharOK=FALSE, NumbOK=TRUE, LogicalOK=FALSE, MinOK=0, MaxOK=Inf)#check validity of supplied values
         if(ERR == TRUE)                                                                                     #If an invalid value is found:-
            {Advise(WithRefTo=paste0("Invalid quotum values in row ", Quota0.Rpar[Qrow], " column ", Cpar, " of \n\t", Params.file), ErrText=paste0("A supplied value for a quotum is ", Params0[Quota0.Rpar[Qrow], Cpar], " . This is not in the acceptable ranges: NA, or 0 =< Quota <= Inf. The file must be adjusted."), AdvText=paste0("A quota row, if supplied, may restrict landings of a fishery. If there is no quotum for a target component, use NA or Inf. \n\tCheck the input row below:-"), ShowData=array(Params0[Quota0.Rpar[Qrow], ], dim=c(1, N.Components), dimnames=list(ParamsQuota0.Rnames[Qrow], Components0)), CallSection="#2.9c", Fatal=TRUE, MinAdviceLevel2Show=0)
            }
         }
      OrderedQrow <- which(Fisheries == ParamsQuota0.suffixes[Qrow])                                         #find the row in the Quota0 matrix corresponding to a fishery
      Quota0[OrderedQrow, ] <- Params0[Quota0.Rpar[Qrow], ]                                                  #put the supplied quota row into the row of Quota0 ordered by fishery; all fisheries have a row wheth regulated/not
      }
   Quota0[ , ] <- ifelse(is.na(Quota0), Inf, Quota0)                                                         #replace NAs with infinities
   }
#----------------------------------------------------------------                                 


#2.10 Essential habitat sizes
#If essential habitats are in the model, there should be 1 HabSize row with +ve values in habitat columns
HabSize0 <- array(data= 0, dim=c(1, N.Components), dimnames = list("HabitatSize", Components0))              #default HabSize
SearchText <- "HabSize"
HabSize0.Rpar <- findOneLine(SearchTxt=SearchText, Headers=Rpar0.names, call.section="#2.10a", file=Params.file, Var.name="HabSize", domain="row", ZeroOK=TRUE)  #find row.header with 'HabSize' in it 
is.Habitat0 <- rep(FALSE, N.Components)                                                                      #default
if(!is.na(HabSize0.Rpar))                                                                                    #If a HabSize row is present:-
   {for(Cpar in 1:N.Components)                                                                              #For each col of the Params.file,
      {ERR <- is.DatumOK(Params0[HabSize0.Rpar, Cpar], "HabSize", NA.OK=TRUE, CharOK=FALSE, NumbOK=TRUE, LogicalOK=FALSE, MinOK=0, MaxNotOK=Inf) #check coeffs for validity
      if(ERR == TRUE)                                                                                        #If an invalid value is found:-
         {Advise(WithRefTo=paste0("Invalid HabSize values in row ", HabSize0.Rpar, " column ", Cpar, " of ", Params.file), ErrText=paste0("A supplied value: HabSize = ", Params0[HabSize0.Rpar, Cpar], ", is not in the accepted range:  NA, or 0 =< HabSize < Inf. The value must be adjusted."), AdvText=paste0("One row of HabSize must be supplied with row header including '", SearchText, " ' if any essential habitat is in the model. Values in the habitat columns are initial sizes, reduced as biomass fills them. Below: Habitat sizes, as read from the file:-\n\t"), ShowData=array(Params0[HabSize.Rpar, ], dim=c(1, N.Components), dimnames=list("HabitatSize", Components0)), CallSection="#2.10b", Fatal=TRUE, MinAdviceLevel2Show=0)
         }
      }
   HabSize0[1, ] <- array(data=Params0[HabSize0.Rpar, ], dim=c(1, N.Components), dimnames=list("HabitatSize", Components0))#fill preliminary Occup array
   HabSize0[1, ] <- ifelse(!is.na(HabSize0[1, ]), HabSize0[1, ], 0)                                          #convert NAs to zeros
   }
is.Habitat0 <- HabSize0 > 0                                                                                  #logical variable to identify col locations of positive habitat size, ie habitat components
Habitats <- Components0[is.Habitat0]                                                                         #names of habitats; the ordering of the habitat column names takes precedence in the rest of the simulation
N.Habitats <- sum(is.Habitat0)                                                                               #number of habitats in the model
#----------------------------------------------------------------

#2.11 Essential habitat occupancies
#If essential habitats are in the model, 1 row of occupancy coeffs (Biomass per unit size) is coded for every habitat whether or not positive values are present; rows are re-ordered to match ordering of the habitat columns.
#Occupancy rows supplied for non-existent habitat columns give a fatal error at #2.11a
HabOccup0 <- array(data= 0, dim=c(1, N.Components), dimnames = list("HabitatOccupancy", Components0))        #default occupancies
is.HabOccup0 <- rep(FALSE, N.Components)                                                                     #default logical variable to indicate col locations of positive occcupancies
ParamsOccup0.Rnames <- NA; ParamsOccup0.suffixes <- NA                                                       #default occupancy row names and suffixes (ie. habitats) in Occup row names of Params
N.OccupHabitat.pairs <- 0                                                                                    #default number of occupancy-habitat pairs as components, eg. Alga1-Hab2
SearchText <- "Occup"                                                                                        #text expected in row headers of Occup coeffs if(N.Habitats > 0)
Occup0.Rpar <- grep(SearchText, ignore.case=T, Rpar0.names)                                                  #find the row numbers of Params0 with this SearchText in them
N.Occup0.Rpar <- length(Occup0.Rpar)                                                                         #count the number of occupancy rows given (whether or not they have any positive values)
if(N.Habitats > 0 & N.Occup0.Rpar > 0)                                                                       #If habitats with occupancies are in the model:-
   {HabOccup0 <- array(data=NA, dim=c(N.Habitats, N.Components), dimnames=list(paste0("Occup.", Habitats), Components0)) #fill preliminary Occup array
   ParamsOccup0.Rnames <- row.names(Params0)[Occup0.Rpar]                                                    #names of Occup rows in Params0
   ParamsOccup0.suffixes <- substr(ParamsOccup0.Rnames, regexpr("[.]", ParamsOccup0.Rnames) +1, 100)         #get the .habitat suffixes located after the dot in each name
   for(Orow in 1:N.Occup0.Rpar)                                                                              #For each supplied occupancy row,
      {ERR <- is.DatumOK(ParamsOccup0.suffixes[Orow], "Habitat occupancy suffixes", NA.OK=FALSE, CharOK=TRUE, NumbOK=FALSE, LogicalOK=FALSE, MustBeInSet=Habitats) #check all supplied occupancies correspond to habitat components
      if(ERR == TRUE)
         {Advise(WithRefTo=paste0("Invalid suffixes for Habitat occupancy values in row ", Occup0.Rpar[Orow], " of \n\t", Params.file), ErrText=paste0("A supplied occupancy suffix is ", Occup0.suffixes[Orow], " . This does not match any component name identified as an essential habitat by positive habitat size, HabSize, in the file. The suffix must be adjusted."), AdvText="Occupancy suffixes, written Occup.suffix, must match one of the components, supplied as column headers, and having a positive value in the row for HabSize. \n\tSuffixes identify essential habitats. \n\tCheck names of habitats found in the file, shown below, for unmatched occupancies:-\n", ShowData=Habitats, CallSection="#2.11a", Fatal=TRUE, MinAdviceLevel2Show=0)
         }
      for(Cpar in 1:N.Components)                                                                            #For all potentially habitat-restricted wild comps in Params0, ie. cols in Occup0,
         {ERR <- is.DatumOK(Params0[Occup0.Rpar[Orow], Cpar], "HabitatOccupancy", NA.OK=TRUE, CharOK=FALSE, NumbOK=TRUE, LogicalOK=FALSE, MinOK=0, MaxNotOK=Inf)#check validity of supplied values
         if(ERR == TRUE)                                                                                     #If an invalid value is found:-
            {Advise(WithRefTo=paste0("Illegal habitat-occupancy values in row ", Orow, " column ", Cpar, " of of \n\t", Params.file), ErrText=paste0("A supplied value for Occup is ", Params0[Occup0.Rpar[Orow], Cpar], " . This is not in the acceptable ranges: NA, or 0 =< Occup < Inf. The file must be adjusted."), AdvText=paste0("Occup, if essential habitats are supplied, restrict the total biomass in an essential habitat.  \n\tCheck the input row below:-"), ShowData=array(Params0[Occup0.Rpar[Orow], ], dim=c(1, N.Components), dimnames=list(Occup0.Rnames[Orow], Components0)), CallSection="#2.11b", Fatal=TRUE, MinAdviceLevel2Show=1)
            }
         }
      OrderedOrow <- which(Habitats == ParamsOccup0.suffixes[Orow])                                          #find the corresponding row in the HabOccup0 matrix ordered by habitats
      HabOccup0[OrderedOrow, ] <- Params0[Occup0.Rpar[Orow], ]                                               #put the supplied occupancy row into the correct row of HabOccup0
      }
   Occup.suffixes <-  Components0[is.Habitat0]                                                               #order occup.suffixes the same as habitat components in Params column headers
   HabOccup0[ , ] <- ifelse(is.na(HabOccup0), 0, HabOccup0)                                                  #replace NAs with zeroes
   is.HabOccup0 <- HabOccup0 > 0                                                                             #logical variable with N.Habitat rows to identify col locations of positive habitat occupancies
   N.OccupHabitat.pairs <- sum(is.HabOccup0)                                                                 #N occupancy-habitat pairs set
   }   
#----------------------------------------------------------------                                 


#2.12 Temperature of water
#A water component is required if Temperature, Light or nutrients are in the model.  Otherwise, omit a water component col.
#A temperature index, T.index, varies from coolest=0 to hottest=1.
#If water temperature is to be varied during a simulation using an Environ.file, one T.index row with one initial temperature index in the water component col should be in Params.file.  Values in other columns should be NA
PhysicalVars <- NULL                                                                                         #default vector of physical vars, eg. T.index, L.index
N.PhysicalVars <- 0                                                                                          #default number of them
T.index0 <- array(data= 0, dim=c(1, N.Components), dimnames = list("T.index0", Components0))                 #default T.index
SearchText <- "T.ind"
T.index0.Rpar <- findOneLine(SearchTxt=SearchText, Headers=Rpar0.names, call.section="#2.12a", file=Params.file, Var.name="T.index0", domain="row", ZeroOK=TRUE)  #find row.header with 'T.ind' in it 
if(!is.na(T.index0.Rpar))                                                                                    #If a T.index row is present:-
   {for(Cpar in 1:N.Components)                                                                              #For each col of the Params.file,
      {ERR <- is.DatumOK(Params0[T.index0.Rpar, Cpar], "T.index0", NA.OK=TRUE, CharOK=FALSE, NumbOK=TRUE, LogicalOK=FALSE, MinOK=0, MaxOK=1) #check coeffs for validity
      if(ERR == TRUE)                                                                                        #If an invalid value is found:-
         {Advise(WithRefTo=paste0("Initial T.index in row ", T.index0.Rpar, " column ", Cpar, " of ", Params.file), ErrText=paste0("A supplied value: T.index0 = ", Params0[T.index0.Rpar, Cpar], ", is not in the accepted range:  NA, or 0 =< T.index =< 1. The value must be adjusted."), AdvText=paste0("One row of initial T.index is needed if temperature is to be varied during the simulation.  The row header must include '", SearchText, " '. Values should be NA in all columns other than that for the water component. This is the initial value; it can be varied using an Environ.file. Below: T.index values, as read from the file:-\n\t"), ShowData=array(Params0[T.index0.Rpar, ], dim=c(1, N.Components), dimnames=list("T.index0", Components0)), CallSection="#2.12b", Fatal=TRUE, MinAdviceLevel2Show=0)
         }
      }
   T.index0[1, ] <- array(data=Params0[T.index0.Rpar , ], dim=c(1, N.Components), dimnames=list("T.index0", Components0))#fill prelimary array
   T.index0[1, ] <- ifelse(!is.na(T.index0[1, ]), T.index0[1, ], 0)                                          #convert NAs to zeros
   }
is.T.index0 <- T.index0 > 0                                                                                  #logical variable with TRUE wherever T.index0 has a positive value
PhysicalVars <- c(PhysicalVars, "T.index")                                                                   #include T.index in a vector of physical variables
N.PhysicalVars <- N.PhysicalVars + 1                                                                         #increment their number
#----------------------------------------------------------------


#2.13 Light in water
#A water component is required if Temperature, Light or nutrients are in the model.  Otherwise, omit a water component col.
#A light (or irradiance) index, L.index, varies from darkest=0 to brightest=1.
#If light is to be varied during a simulation using an Environ.file, one L.index row with one initial light index in the water component col should be in Params.file.  Values in other columns should be NA
L.index0 <- array(data= 0, dim=c(1, N.Components), dimnames = list("L.index0", Components0))                 #default L.index
SearchText <- "L.ind"                                                                                        #search text
L.index0.Rpar <- findOneLine(SearchTxt=SearchText, Headers=Rpar0.names, call.section="#2.13a", file=Params.file, Var.name="L.index0", domain="row", ZeroOK=TRUE)  #find row.header with 'L.ind' in it 
if(!is.na(L.index0.Rpar))                                                                                    #If a L.index row is present:-
   {for(Cpar in 1:N.Components)                                                                              #For each col of the Params.file,
      {ERR <- is.DatumOK(Params0[L.index0.Rpar, Cpar], "L.index0", NA.OK=TRUE, CharOK=FALSE, NumbOK=TRUE, LogicalOK=FALSE, MinOK=0, MaxOK=1) #check coeffs for validity
      if(ERR == TRUE)                                                                                        #If an invalid value is found:-
         {Advise(WithRefTo=paste0("Initial L.index in row ", L.index0.Rpar, " column ", Cpar, " of ", Params.file), ErrText=paste0("A supplied value: L.index0 = ", Params0[L.index0.Rpar, Cpar], ", is not in the accepted range:  NA, or 0 =< L.index =< 1. The value must be adjusted."), AdvText=paste0("One row of initial L.index is needed if light is to be varied during the simulation.  The row header must include '", SearchText, " '. Values should be NA in all columns other than that for the water component. This is the initial value; it can be varied using an Environ.file. Below: L.index values, as read from the file:-\n\t"), ShowData=array(Params0[L.index.Rpar, ], dim=c(1, N.Components), dimnames=list("L.index0", Components0)), CallSection="#2.12b", Fatal=TRUE, MinAdviceLevel2Show=0)
         }
      }
   L.index0[1, ] <- array(data=Params0[L.index0.Rpar , ], dim=c(1, N.Components), dimnames=list("L.index0", Components0))#fill preliminary array
   L.index0[1, ] <- ifelse(!is.na(L.index0[1, ]), L.index0[1, ], 0)                                          #convert NAs to zeros
   }
is.L.index0 <- L.index0 > 0                                                                                  #logical variable with TRUE wherever there is a positive value of L.index
PhysicalVars <- c(PhysicalVars, "L.index")                                                                   #include L.index in a vector of physical variables
N.PhysicalVars <- N.PhysicalVars + 1                                                                         #increment their number
#----------------------------------------------------------------


#2.14 Nutrient masses in H2O, and NLO (if in model)
#A water component is required if Temperature, Light or nutrients are in the model.  
#If essential nutrients are in the model, there should be 1 row of nutrient masses for each nutrient with positive values in water and NLO
#Additionally, there should be 1 row of nutrient concentrations for each nutrient with positive values in component columns for wilds.  (Nconc in NLO is calculated from Bmass and Nmass present. Nconc in H2O is not used.)
N.Nutr <- 0                                                                                                  #default
Nmass0 <- array(data= 0, dim=c(1, N.Components), dimnames = list("Nmass", Components0))                      #default Nmass
SearchText <- "Nmass"                                                                                        #text expected in row header of Nmass coeffs
Nmass0.Rpar <- grep(SearchText, ignore.case=T, Rpar0.names)                                                  #find the row headers of Params0 with this SearchText in them
N.Nmass0.Rpar <- length(Nmass0.Rpar)                                                                         #count the number of Nmass rows given
Nmass0.Rnames <- Rpar0.names[Nmass0.Rpar]                                                                    #temporarily store the Nmass row names from those given in Params0
Nmass0.suffixes <- substr(Nmass0.Rnames, regexpr("[.]", Nmass0.Rnames) +1, 100)                              #get the Nmass.comp suffixes located after the dot in the name
if(N.Nmass0.Rpar > 0)                                                                                        #if Nmass variables were supplied:-
   {for(Rnmass0 in 1:N.Nmass0.Rpar)                                                                          #For all potential Nmass nutrients in Params0, ie. rows in Nmass0,
      {for(Cpar in 1:N.Components)                                                                           #For all potentially Nmass-containing components in Params0, ie. cols in Nmass0,
         {ERR <- is.DatumOK(Params0[Nmass0.Rpar[Rnmass0], Cpar], "Nmass", NA.OK=TRUE, CharOK=FALSE, NumbOK=TRUE, LogicalOK=FALSE, MinOK=0, MaxNotOK=Inf)#check supplied values
         if(ERR == TRUE)                                                                                     #If an invalid value is found:-
            {Advise(WithRefTo=paste0("Illegal nutrient-mass values in row ", Rnmass0, " column ", Cpar, " of of \n\t", Params.file), ErrText=paste0("A supplied value for Nmass is ", Params0[Nmass0.Rpar[Rnmass0], Cpar], " . This is not in the acceptable ranges: NA, or 0 =< Nmass < Inf. The file must be adjusted."), AdvText=paste0("Nmass indicates that essential nutrients are in the model; Nmass may restrict GP of autotrophs requiring the essential nutrient.  \n\tCheck the input row below:-"), ShowData=array(Params0[Nmass0.Rpar[Rnmass0], ], dim=c(1, N.Components), dimnames=list(Nmass0.Rnames[Rnmass0], Components0)), CallSection="#2.14a", Fatal=TRUE, MinAdviceLevel2Show=1)
            }
         }
      }
   Nmass0 <- array(data=Params0[Nmass0.Rpar, ], dim=c(N.Nmass0.Rpar, N.Components), dimnames=list(Nmass0.Rnames, Components0))#fill preliminary Nmass array
   Nmass0 <- ifelse(!is.na(Nmass0), Nmass0, 0)                                                               #convert any NAs to 0 for arithmetic purposes
   }
is.Nmass0 <- Nmass0 > 0                                                                                      #logical variable with TRUE wherever there is a positive value of Nmass
Nutrients <- Nmass0.suffixes                                                                                 #names of nutrients in the model in the order of the Nmasses
N.Nutr <- N.Nmass0.Rpar                                                                                      #store the number of nutrients from reading Nmasses; this value has priority of any other values read (eg. from Nconcs)
#----------------------------------------------------------------                                 


#2.15 Nutrient concentrations:- Nconc
#If essential nutrients are in the model, as indicated by Nmass rows in Params.file, there must also be 1 row of Nconc values for each nutrient in wild  components
#Note: Nconc in NLO component(s) is calculated after Params0 has been sorted to Params column ordering later
Nconc0 <- array(data= 0, dim=c(1, N.Components), dimnames = list("Nconc", Components0))                      #default Nconc
SearchText <- "Nconc"                                                                                        #text expected in row header of Nconc coeffs
Nconc0.Rpar <- grep(SearchText, ignore.case=T, Rpar0.names)                                                  #find the row headers of Params0 with this SearchText in them
N.Nconc0.Rpar <- length(Nconc0.Rpar)                                                                         #count the number of Nconc rows given
Nconc0.Rnames <- Rpar0.names[Nconc0.Rpar]                                                                    #temporarily store the Nconc row names from those given in Params0; may be re-sorted
Nconc0.suffixes <- substr(Nconc0.Rnames, regexpr("[.]", Nconc0.Rnames) +1, 100)                              #get the Nconc.comp suffixes located after the dot in the name
if(N.Nconc0.Rpar > 0)                                                                                        #If Nconc rows were supplied:-
   {for(Rnconc0 in 1:N.Nconc0.Rpar)                                                                          #For all potential Nconc nutrients in Params0, ie. rows in Nconc0
      {#Check that Nconc.suffixes match Nmass.suffixes=Nutrients, #2.14
      ERR <- is.DatumOK(Nconc0.suffixes[Rnconc0], "Nutrient-concentration suffixes", NA.OK=FALSE, CharOK=TRUE, NumbOK=FALSE, LogicalOK=FALSE, MustBeInSet=Nutrients)
      if(ERR == TRUE)
         {Advise(WithRefTo=paste0("Invalid suffixes for Nutrient concentration values in row ", Nconc0.Rpar[Rnconc0], " of \n\t", Params.file), ErrText=paste0("A supplied nutrient-concentration suffix is ", Nconc0.suffixes[Rnconc0], " . This does not match any nutrient name identified from Nmass.suffixes in the file. \n\tThe Nconc suffix must be adjusted."), AdvText="Nconc suffixes, written Nconc.suffix, must match one of the nutrient mass suffixes, written Nmass.suffix, in input rows for Nmass. \n\tNconc.suffix gives the concentration of the suffixed nutrient in living components and in non-living organic matter. \n\tCheck the nutrient names found in the Nmass rows, shown below:-\n", ShowData=Nmass0.suffixes, CallSection="#2.15a", Fatal=TRUE, MinAdviceLevel2Show=1)
         }
      for(Cpar in 1:N.Components)                                                                            #For all potentially Nconc-containing comps in Params0, ie. cols in Nconc0
         {ERR <- is.DatumOK(Params0[Nconc0.Rpar[Rnconc0], Cpar], "Nconc", NA.OK=TRUE, CharOK=FALSE, NumbOK=TRUE, LogicalOK=FALSE, MinOK=0, MaxNotOK=Inf)#check supplied values
         if(ERR == TRUE)                                                                                     #If an invalid value is found:-
            {Advise(WithRefTo=paste0("Illegal nutrient-concentration values in row ", Rconc0, " column ", Cpar, " of of \n\t", Params.file), ErrText=paste0("A supplied value for Nconc is ", Params0[Nconc0.Rpar[Rconc0], Cpar], " . This is not in the acceptable ranges: NA, or 0 =< Nconc < Inf. The file must be adjusted."), AdvText=paste0("Nconc specifies requirements of wild components for essential nutrients, if present in the model; high Nconc values increase the requirement for a nutrient for the GP of autotrophs.  \n\tCheck the input row below:-"), ShowData=array(Params0[Nconc0.Rpar[Rnconc0], ], dim=c(1, N.Components), dimnames=list(Nconc0.Rnames[Rnconc0], Components0)), CallSection="#2.15a", Fatal=TRUE, MinAdviceLevel2Show=0)
            }
         }
      }
   Nconc0 <- array(data=Params0[Nconc0.Rpar, ], dim=c(N.Nconc0.Rpar, N.Components), dimnames=list(Nconc0.Rnames, Components0))#fill preliminary Nconc array
   Nconc0 <- ifelse(!is.na(Nconc0), Nconc0, 0)                                                               #convert any NAs to 0 for arithmetic purposes
   }
is.Nconc0 <- Nconc0 > 0                                                                                      #logical variable with TRUE wherever Nconc0 has a positive value
#----------------------------------------------------------------                                 


#2.16 Particulate SelfLoss coeffs
#Particulate selfLoss coeffs are required if NLO or nutrients are present; they control recycling of nutrients and/or biomass via particulates falling to NLO
ParticulateSL0 <- array(data=1, dim=c(1, N.Components), dimnames = list("ParticulateSL0", Components0))      #default ParticulateSL
SearchText <- "Partic"
ParticulateSL0.Rpar <- findOneLine(SearchTxt=SearchText, Headers=Rpar0.names, call.section="#2.16a", file=Params.file, Var.name="ParticulateSL0", domain="row", ZeroOK=TRUE)  #find row.header with 'Partic' in it 
if(!is.na(ParticulateSL0.Rpar))                                                                              #If a ParticulateSL row is present:-
   {for(Cpar in 1:N.Components)                                                                              #For each col of the Params.file,
      {ERR <- is.DatumOK(Params0[ParticulateSL0.Rpar, Cpar], "ParticulateSL0", NA.OK=TRUE, CharOK=FALSE, NumbOK=TRUE, LogicalOK=FALSE, MinOK=0, MaxOK=1) #check coeffs for validity
      if(ERR == TRUE)                                                                                        #If an invalid value is found:-
         {Advise(WithRefTo=paste0("Initial ParticulateSL in row ", ParticulateSL0.Rpar, " column ", Cpar, " of ", Params.file), ErrText=paste0("A supplied value: ParticulateSL0 = ", Params0[ParticulateSL0.Rpar, Cpar], ", is not in the accepted range:  NA, or 0 =< ParticulateSL =< 1. The value must be adjusted."), AdvText=paste0("One row of initial ParticulateSL is needed if NLO or nutrients are present; they control recycling of nutrients and/or biomass via \n\tparticulates falling to NLO. The row header must include '", SearchText, " '. Positive values should be present for all wild components; all \n\tother cols have NA or 0. Below: ParticulateSL values, as read from the file:-\n\t"), ShowData=array(Params0[ParticulateSL0.Rpar, ], dim=c(1, N.Components), dimnames=list("ParticulateSL0", Components0)), CallSection="#2.16b", Fatal=TRUE, MinAdviceLevel2Show=0)
         }
      }
   ParticulateSL0[1, ] <- array(data=Params0[ParticulateSL0.Rpar , ], dim=c(1, N.Components), dimnames=list("ParticulateSL0", Components0))#fill prelimary array
   ParticulateSL0[1, ] <- ifelse(!is.na(ParticulateSL0[1, ]), ParticulateSL0[1, ], 0)                        #convert NAs to zeros
   }
is.ParticulateSL0 <- ParticulateSL0 > 0                                                                      #logical variable with TRUE wherever ParticulateSL0 has a positive value
#----------------------------------------------------------------


#2.17 Light-dependent adjustments of maximum SelfGrowth rates of autotrophs
#Environ.file can be used to alter the values of parameters or of bio- or nutrient-masses over time if required.  
#Light-dependent changes of SelfGrow use alphaSG.I; otherwise the value of SelfGrow supplied for each autotroph in Params.file is a maximum rate that applies over the entire simulation
alphaSG.I0 <- array(data=0, dim=c(1, N.Components), dimnames=list("alphaSG.I0", Components0))                #default alphaSG.I
SearchText <- "alphaSG"                                                                                      #text in Params file to be searched for
alphaSG.I0.Rpar <- findOneLine(SearchText, Headers=Rpar0.names, call.section="#2.17a", Params.file, Var.name="alphaSG.I", domain="row", ZeroOK=TRUE) #find one line in Params.file with SearchText in it
if(!is.na(alphaSG.I0.Rpar))                                                                                  #If a row of alphaSG.I0 has been supplied:-
   {for (Cpar in 1:N.Components)                                                                             #For each col of the Params.file,
      {ERR <- is.DatumOK(Params0[alphaSG.I0.Rpar, Cpar], SearchText, NA.OK=TRUE, CharOK=FALSE, NumbOK=TRUE, LogicalOK=FALSE, MinOK=0, MaxNotOK=Inf) #check value for validity
      if(ERR == TRUE)                                                                                        #If an invalid value is found:-
         {Advise(WithRefTo=c(paste0("alphaSG.I, given in ", Params.file), " , row ", alphaSG.I0.Rpar, " , column ", Cpar), ErrText=paste0("A value of alphaSG.I, ", Params0[alphaSG.I0.Rpar, Cpar], " , supplied for autotrophs is not in the acceptable range NA, or 0 <= alphaSG.I < Inf.  \n\tThe value must be adjusted."), AdvText="alphaSG.I is the slope coeff of SelfGrow over L.index used to adjust SelfGrow with varying light. \n\tUse of it requires an initial positive L.index value supplied in the water column of the Params file; it can be varied later with new \n\tL.index values in an Environ.file.  L.index varies between 0=darkest to 1=brightest over the simulation.  \n\tBelow:- alphaSG.I as read from the file: ", ShowData= Params0[alphaSG.I0.Rpar, ], CallSection="#2.17b", Fatal=TRUE, MinAdviceLevel2Show=0)
         }
      }
   alphaSG.I0[1, ] <- array(data=Params0[alphaSG.I0.Rpar, ], dim=c(1, N.Components), dimnames=list("alphaSG.I0", Components0))  #fill prelimary array
   alphaSG.I0[1, ] <- ifelse(!is.na(alphaSG.I0[1, ]), alphaSG.I0[1, ], 0)                                    #convert NAs to zeros
   }
is.alphaSG.I0 <- alphaSG.I0 > 0                                                                              #logical variable with TRUE wherever alphaSG.I0 hqs a positive value
#----------------------------------------------------------------


#2.18 Temperature bands within which positive SelfGrow rates occur
#If required, SelfGrow may only be positive if T.index is within a pre-set temperature band favoured by the autotroph. This is done with LoBoundSG.T and HiBoundSG.T; each of these may range between 0 and 1.
LoBoundSG.T0 <- array(data=0, dim=c(1, N.Components), dimnames=list("LoBoundSG.T0", Components0))            #default row vector of values for LoBoundSG.T
HiBoundSG.T0 <- array(data=1, dim=c(1, N.Components), dimnames=list("HiBoundSG.T0", Components0))            #default row vector of values for HiBoundSG.T;  Lo=0 and Hi=1 -> SelfGrow can be +ve at all temps by default
SearchText <- "LoBoundSG.T"                                                                                  #text in Params file to be searched for
LoBoundSG.T0.Rpar <- findOneLine(SearchText, Headers=Rpar0.names, call.section="#2.16a", Params.file, Var.name="LoBoundSG.T", domain="row", ZeroOK=TRUE) #find one line in Params.file with SearchText in it; >1 lines gives an error. Because argument ZeroOK is set to TRUE, no LoBoundSG.T row in the Params.file returns an NA for LoBoundSG.T.Rpar; see findOneLine() in ECOLPSfunctions.R
SearchText <- "HiBoundSG.T"                                                                                  #next text in Params file to be searched for
HiBoundSG.T0.Rpar <- findOneLine(SearchText, Headers=Rpar0.names, call.section="#2.16b", Params.file, Var.name="HiBoundSG.T0", domain="row", ZeroOK=TRUE) #find one line in Params.file with SearchText in it; 
if(!is.na(LoBoundSG.T0.Rpar) | !is.na(HiBoundSG.T0.Rpar))                                                    #If a row of LoBoundSG.T0 AND/OR a row of HiBoundSG.T0 have been supplied:-
   {for (Cpar in 1:N.Components)                                                                             #for each col in Params0,
      {ERR1 <- ERR2 <- FALSE                                                                                 #clear ERR switches
      if(!is.na(LoBoundSG.T0.Rpar)) ERR1 <- is.DatumOK(Params0[LoBoundSG.T0.Rpar, Cpar], SearchText, NA.OK=TRUE, CharOK=FALSE, NumbOK=TRUE, LogicalOK=FALSE, MinOK=0, MaxNotOK=1) #check values for validity
      if(!is.na(HiBoundSG.T0.Rpar)) ERR2 <- is.DatumOK(Params0[HiBoundSG.T0.Rpar, Cpar], SearchText, NA.OK=TRUE, CharOK=FALSE, NumbOK=TRUE, LogicalOK=FALSE, MinOK=Params0[LoBoundSG.T0.Rpar, Cpar], MaxOK=1)
      if(ERR1 == TRUE | ERR2 == TRUE)                                                                        #If either value is found invalid:-
         {Advise(WithRefTo=paste0("LoBoundSG.T and HiBoundSG.T, in rows ", LoBoundSG.T0.Rpar, " and ", HiBoundSG.T0.Rpar, " of \n\t", Params.file), ErrText="Values of LoBoundSG.T or HiBoundSG.T  supplied for positive selfgrowth by autotrophs are not in the acceptable range NA, or 0 <= LoBoundSG.T < HiBoundSG.T <= 1, \n\t. Values must be adjusted.", AdvText="LoBoundSG.T and HiBoundSG.T bound the temperature band in which SelfGrow rates by autotrophs depend on L.index values \n\tsupplied in an Envir.file. An initial postive T.index value must be supplied in the water column of the Params file; \n\tit can be varied later with new T.index values in an Environ.file.  T.index varies between 0=coldest to 1=warmest over the simulation.  \n\tBelow:- LoBoundSG.T and HiBoundSG.T as read from the file: ", ShowData= Params0[c(LoBoundSG.T0.Rpar, HiBoundSG.T0.Rpar), Cpar], CallSection="#2.18c", Fatal=TRUE, MinAdviceLevel2Show=0)
         }
      }
   LoBoundSG.T0[1, ] <- array(data=Params0[LoBoundSG.T0.Rpar, ], dim=c(1, N.Components), dimnames=list("LoBoundSG.T0", Components0))  #fill prelimary array 
   HiBoundSG.T0[1, ] <- array(data=Params0[HiBoundSG.T0.Rpar, ], dim=c(1, N.Components), dimnames=list("HiBoundSG.T0", Components0))  #fill prelimary array 
   LoBoundSG.T0[1, ] <- ifelse(!is.na(LoBoundSG.T0[1, ]), LoBoundSG.T0[1, ], 0)                              #convert NAs to zeros
   HiBoundSG.T0[1, ] <- ifelse(!is.na(HiBoundSG.T0[1, ]), HiBoundSG.T0[1, ], 1)                              #convert NAs to ones; with Lo=0 and Hi=1, both bounds have no effect, consistent with Lo and Hi both being NA
   }
is.LoBoundSG.T0 <- LoBoundSG.T0 > 0                                                                          #logical variable with TRUE wherever LoBoundSG.T0 has a positive value
is.HiBoundSG.T0 <- HiBoundSG.T0 < 1                                                                          #same for HiBound wherever less than 1
#----------------------------------------------------------------


#2.19 Temperature-dependent adjustments of harvesting rates by heterotrophs
alphaHvst.T0 <- array(data=0, dim=c(1, N.Components), dimnames=list("alphaHvst.T0", Components0))            #default row vector of values for alphaHvst.T
SearchText <- "alphaH"                                                                                       #text in Params file to be searched for
alphaHvst.T0.Rpar <- findOneLine(SearchText, Headers=Rpar0.names, call.section="#2.19a", Params.file, Var.name="alphaHvst.T0", domain="row", ZeroOK=TRUE) #find one line in Params.file with SearchText in it; >1 lines gives an error. Because argument ZeroOK is set to TRUE, no alphaHvst.T row in the Params.file returns an NA for alphaHvst.T.Rpar; see findOneLine() in ECOLPSfunctions.R
if(!is.na(alphaHvst.T0.Rpar))                                                                                #If a row of alphaHvst.T0 has been supplied:-
   {for(Cpar in 1:N.Components)                                                                              #For all cols in Params0,
      {ERR <- is.DatumOK(Params0[alphaHvst.T0.Rpar, Cpar], SearchText, NA.OK=TRUE, CharOK=FALSE, NumbOK=TRUE, LogicalOK=FALSE, MinOK=0, MaxNotOK=Inf) #check values for validity
      if(ERR == TRUE)                                                                                        #If an invalid value is found:-
         {Advise(WithRefTo=paste0("alphaHvst.T, given in ", Params.file, " , row ", alphaHvst.T0.Rpar, " , column ", Cpar), ErrText=paste0("A value of alphaHvst.T, ", Params0[alphaHvst.T0.Rpar, Cpar], " , supplied for heterotrophs is not in the acceptable range NA, or 0 =< alphaHvst.T < Inf.  The value must be adjusted."), AdvText="alphaHvst.T is the slope coeff of Harv over T.index used to adjust harvest coeffs with temperature. An initial positive T.index \n\tvalue must be supplied in the water column of the Params file; it can be varied later with new T.index values in an Environ.file.  \n\tT.index varies between 0=coolest to 1=warmest over the simulation.  Below:- alphaHvst.T as read from the file: ", ShowData= Params0[alphaHvst.T0.Rpar, ], CallSection="#2.19b", Fatal=TRUE, MinAdviceLevel2Show=1)
         }
      }
   alphaHvst.T0[1, ] <- array(data=Params0[alphaHvst.T0.Rpar, ], dim=c(1, N.Components), dimnames=list("alphaHvst.T0", Components0))  #fill prelimary array
   alphaHvst.T0[1, ] <- ifelse(!is.na(alphaHvst.T0[1, ]), alphaHvst.T0[1, ], 0)                              #convert NAs to zeros
   }
is.alphaHvst.T0 <- alphaHvst.T0 > 0                                                                          #logical variable with TRUE wherever alphaHvst.T0 has a positive value
#----------------------------------------------------------------


#2.20  Temperature band within which positive harvest rates occur
#If required, harvesting rates may only be positive if T.index is within a pre-set temperature band favoured by the heterotroph. This is done with LoBoundHvst.T and HiBoundHvst.T; each of these may range between 0 and 1.
LoBoundHvst.T0 <- array(data=0, dim=c(1, N.Components), dimnames=list("LoBoundHvst.T0", Components0))        #default row vector of values for LoBoundHvst.T
HiBoundHvst.T0 <- array(data=1, dim=c(1, N.Components), dimnames=list("HiBoundHvst.T0", Components0))        #default row vector of values for HiBoundHvst.T;  Lo=0 and Hi=1 -> Harvest can be +ve at all temps by default
SearchText <- "LoBoundH.T"                                                                                   #text in Params file to be searched for
LoBoundHvst.T0.Rpar <- findOneLine(SearchText, Headers=Rpar0.names, call.section="#2.20a", Params.file, Var.name="LoBoundHvst.T0", domain="row", ZeroOK=TRUE) #find one line in Params.file with SearchText in it; >1 lines gives an error. Because argument ZeroOK is set to TRUE, no LoBoundHvst.T row in the Params.file returns an NA for LoBoundHvst.T.Rpar; see findOneLine() in ECOLPSfunctions.R
SearchText <- "HiBoundH.T"                                                                                   #next text in Params file to be searched for
HiBoundHvst.T0.Rpar <- findOneLine(SearchText, Headers=Rpar0.names, call.section="#2.20b", Params.file, Var.name="HiBoundHvst.T0", domain="row", ZeroOK=TRUE) #find one line in Params.file with SearchText in it; 
if(!is.na(LoBoundHvst.T0.Rpar) | !is.na(HiBoundHvst.T0.Rpar))                                                #If a row of LoBoundHvst.T0 AND/OR a row of HiBoundHvst.T0 have been supplied:-
   {for(Cpar in 1:N.Components)                                                                              #For each col in Params0,
      {ERR1 <- ERR2 <- FALSE                                                                                 #clear ERR switches; check values for validity:-
      if(!is.na(LoBoundHvst.T0.Rpar)) ERR1 <- is.DatumOK(Params0[LoBoundHvst.T0.Rpar, Cpar], SearchText, NA.OK=TRUE, CharOK=FALSE, NumbOK=TRUE, LogicalOK=FALSE, MinOK=0, MaxNotOK=1) #check values for validity
      if(!is.na(HiBoundHvst.T0.Rpar)) ERR2 <- is.DatumOK(Params0[HiBoundHvst.T0.Rpar, Cpar], SearchText, NA.OK=TRUE, CharOK=FALSE, NumbOK=TRUE, LogicalOK=FALSE, MinOK=Params0[LoBoundHvst.T0.Rpar, Cpar], MaxOK=1) #check values
      if(ERR1 == TRUE | ERR2 == TRUE)                                                                        #If either value is found invalid:-
         {Advise(WithRefTo=paste0("LoBoundHvst.T and HiBoundHvst.T, in rows ", LoBoundHvst.T0.Rpar, " and ", HiBoundHvst.T0.Rpar, " of \n\t", Params.file), ErrText="Values of LoBoundHvst.T or HiBoundHvst.T  supplied for positive harvesting by heterotrophs are not in the acceptable range NA, or 0 <= LoBoundHvst.T < HiBoundHvst.T <= 1, \n\t. Values must be adjusted.", AdvText="LoBoundHvst.T and HiBoundHvst.T bound the temperature band in which Harv rates by depend on T.index values \n\tsupplied in an Envir.file. An initial postive T.index value must be supplied in the water column of the Params file; \n\tit can be varied later with new T.index values in an Environ.file.  T.index varies between 0=coldest to 1=warmest over the simulation.  \n\tBelow:- LoBoundHvst.T and HiBoundHvst.T as read from the file: ", ShowData= Params0[c(LoBoundHvst.T0.Rpar, HiBoundHvst.T0.Rpar), Cpar], CallSection="#2.20c", Fatal=TRUE, MinAdviceLevel2Show=0)
         }
      }
   LoBoundHvst.T0[1, ] <- array(data=Params0[LoBoundHvst.T0.Rpar, ], dim=c(1, N.Components), dimnames=list("LoBoundHvst.T0", Components0))  #fill prelimary array 
   HiBoundHvst.T0[1, ] <- array(data=Params0[HiBoundHvst.T0.Rpar, ], dim=c(1, N.Components), dimnames=list("HiBoundHvst.T0", Components0))  #fill prelimary array 
   LoBoundHvst.T0[1, ] <- ifelse(!is.na(LoBoundHvst.T0[1, ]), LoBoundHvst.T0[1, ], 0)                        #convert NAs to zeros
   HiBoundHvst.T0[1, ] <- ifelse(!is.na(HiBoundHvst.T0[1, ]), HiBoundHvst.T0[1, ], 1)                        #convert NAs to ones; with Lo=0 and Hi=1, both bounds have no effect, consistent with Lo and Hi being NA
   }      
is.LoBoundHvst.T0 <- LoBoundHvst.T0 > 0                                                                      #logical variable with TRUE wherever LoBoundHvst.T0 has a positive value
is.HiBoundHvst.T0 <- HiBoundHvst.T0 < 1                                                                      #same for HiBoundHvst.T0 wherever < 1
#----------------------------------------------------------------
#################################################################End of section 2. READING AND IDENTIFICATION OF COMPONENTS AND THEIR ATTRIBUTES


#3. IDENTIFICATION OF COMPONENT TYPES
#The Variable0 arrays created in #2. are classified into component types (without sorting) in this section. 
#is.variable0s are logical, N.components wide
# Must be at least 1 auto in the model.                                                                
# Must be at least 1 hetero                                                                            
# Only 0 or 1 H2O, and 0 or 1 NLO, is allowed in this version of ECOLPS.
# Omitting Growth criterion for fisheries allows them to be in Params.file even if it takes no landings
ClassificationRules <- "(1) Autotrophs have SelfGrow > 0, SelfLoss > 0, and Bmass > 0                   
 (2) Heterotrophs have SelfGrow = 0, SelfLoss > 0, Bmass > 0, and at least 1 Growth > 0                 
 (3) Fisheries have fishing Effort > 0, SelfGrow = 0, SelfLoss = 0, Biomass = 0                          
 (4) Non-living organics (NLO) has SelfGrow = 0, SelfLoss > 0,  Bmass > 0, and all Growth = 0.           
 (5) Habitats have HabSize > 0, and a HabOccup > 0 must exist for at least 1 wild component.
 (6) Water (H2O) has at least one from initial {T.index, L.index, and/or Nmass.nutr} > 0.                
 (7) Wilds-and-NLOs have SelfLoss > 0 and Bmass > 0.
 (8) Scavengers are heterotrophs that have Harv.NLO > 0 and Assim.NLO > 0. Scavengers require NLO but may also eat wilds."     #all this text is used for an error message below

Growth0 <- Harv0 * Assim0                                                                                    #Harv0 and Assim0 were sorted in #2. so that their row orderings are the same
CompTypeList <- list("Autotroph", "Heterotroph", "Fishery", "NonLivingOrganics", "Habitat", "Water")         #list of all legal component types in the standard order to be used
is.CompType0 <- array(data=FALSE, dim=c(6, N.Components), dimnames=list(paste0("is.", CompTypeList), Components0))  #logical variable, true or false for members 1 to 6 of CompTypeList
is.WildOrNLO0 <- array(data=FALSE, dim=c(1, N.Components), dimnames=list("is.WildOrNLO0", Components0))      #wild or NLO components will be marked true
is.Harvester0 <- array(data=FALSE, dim=c(1, N.Components), dimnames=list("is.Harvester0", Components0))      #harvester comps will be marked true                                                  
is.Consumer0 <- array(data=FALSE, dim=c(1, N.Components), dimnames=list("is.Consumer0", Components0))        #consumer comps will be marked true                                                 
is.IdentComp0 <- array(data=0, dim=c(1, N.Components), dimnames=list("is.Identified0", Components0))         #marks how many times a component has been identified to type to check that its type is unique

is.CompType0[1, ] <- is.SelfGrow0 & is.SelfLoss0 & is.Bmass0                                                 #autotrophs are TRUE
is.CompType0[2, ] <- !is.SelfGrow0 & is.SelfLoss0 & is.Bmass0 & Csums(Growth0, N.Harv0.Rpar, N.Components) > 0  #heterotrophs are TRUE; Csums() is in ECOLPSfunctions.R => logical row vector ##
is.CompType0[3, ] <- is.Fishery0 & !is.SelfGrow0 & !is.SelfLoss0 & !is.Bmass0                                #fisheries are TRUE 
is.CompType0[4, ] <- !is.SelfGrow0 & is.SelfLoss0 & is.Bmass0 & Csums(Growth0, N.Harv0.Rpar, N.Components) == 0  #NLO is TRUE ; must be non-harvesting    ##& Csums(Harv0 * Assim0, N.Harv0.Rpar, N.Components) > 0
if(N.Habitats > 0) is.CompType0[5, ] <- is.Habitat0                                                          #essential habitats are TRUE from positive habitat sizes; occupancies may all be zero
is.CompType0[6, ] <- (is.T.index0 | is.L.index0 | is.Nmass0[1, ]) & !is.CompType0[4, ]                       #water is TRUE.  T.index, L.index, Nmass may all singly create a water component
is.WildOrNLO0 <- is.Bmass0                                                                                   #wild and NLO are TRUE
is.Harvester0[ , ] <- Csums(Harv0, N.Harv0.Rpar, N.Components) > 0                                           #harvesters incl fisheries = TRUE; Csums() sums TRUEs in comp col of Harv0;Csums() in ECOLPSfunctions
is.Consumer0[ , ] <-  Csums(Growth0, N.Harv0.Rpar, N.Components) > 0                                         #heterotrophs and fisheries are TRUE because they grow from harvests. Note, fisheries are non-wild
is.IdentComp0[ , ] <- Csums(is.CompType0, length(CompTypeList), N.Components)                                #components identified to type are TRUE. Note, TRUEs are summed as 1s, FALSEs as 0s

ErrMess <- NULL                                                                                                                #cancel any lingering error flags; NULL allows concatenation of text
if(sum(is.CompType0[1, ]) < 1) ErrMess <- c(ErrMess, "At least 1 autotroph is required but none were found. " )                #If no autos, error message
if(sum(is.CompType0[2, ]) < 1) ErrMess <- c(ErrMess, "At least 1 heterotroph is required but none were found.  ")              #If no heteros, error message
if(sum(ifelse(is.CompType0[3, ] == is.Fishery0[ ], TRUE, FALSE)) != N.Components) ErrMess <- c(ErrMess, "Classification to fisheries is not matched by supplied fishing-effort data.  ")  #If conflicting fishery id.s, error mess
if(sum(is.CompType0[4, ]) > 1) ErrMess <- c(ErrMess, "Only zero or one non-living-organics (NLO) component is permitted but more were found.  ") #If too many NLO components for this version of ECOLPS, error message
if(sum(ifelse(is.CompType0[5, ] == is.Habitat0[ ], TRUE, FALSE)) != N.Components) ErrMess <- c(ErrMess, "Classification to habitats by habitat-size data is not matched by supplied rows of Occup.habitat.  ")
if(sum(is.CompType0[6, ]) > 1) ErrMess <- c(ErrMess, "Only zero or one water (H2O) component is permitted but more were found.  ")#If too many H2O components for this version of ECOLPS, error message
if(!is.null(ErrMess))                                                                                                          #If an error has been found:-
   {Advise(WithRefTo=paste0("Errors found when classifying components supplied in ", Params.file, " to recognized types."), ErrText=c(ErrMess, "\n\tThe file must be adjusted."), AdvText="Outputs below show: Line 1 how many types were identified for each supplied component by ECOLPS; all should be 1.\n\tLines 2 onwards, the matrix of assigned types and, last, the rules for classification by type.\n\tAn unclassified component may be removed from the model if not required by changing its column name to NA", ShowData= rbind(is.IdentComp0, is.CompType0), CallSection="#3.a", Fatal=FALSE, MinAdviceLevel2Show=0)
   cat("\n\n",ClassificationRules, "\n")
   stop("\b\b\b\b\b\b\b\nFatal error: One or more components were misclassified.", call.=FALSE) 
   }
       
if(any(is.IdentComp0 != 1))                                                                                  #If any cols of Params0 identified as more than 1 component type:-
   {Advise(WithRefTo=paste0("Classifying components supplied in ", Params.file, " to recognized types."), ErrText="Supplied components do not all belong to one and only one type, as listed below, according to the values of certain parameters.  \n\tParameter values given in the file must be adjusted.", AdvText="Outputs below show: Line 1 how many types were identified for each supplied component by ECOLPS; all should be 1.\n\tLines 2 onwards, the matrix of assigned types and, last, the rules for classification by type.\n\tAn unclassified component may be removed from the model if not required by changing its column name to NA", ShowData= rbind(is.IdentComp0, is.CompType0), CallSection="#3.b", Fatal=FALSE, MinAdviceLevel2Show=0)
   cat("\n\n",ClassificationRules, "\n")
   stop("\b\b\b\b\b\b\b\nFatal error: Failed classification of components.", call.=FALSE) 
   }   


#4. RE-ORDERING OF COMPONENTS, CREATION OF PARAMETERS AND INTERNAL LOCATION VECTORS
#Column ordering for components found in the Params.file is altered to a standard = (Autotrophs, Heterotrophs, Fisheries, NLO, Habitats, H2O) if not already so ordered.  The same ordering is used for the rows and cols, referred to as suffix .RC, of matrix parameters such as Harv, Assim, Growth.  Parameter structures having the orderings as-read had 0 appended (in #2 above) to their names. Re-ordered structures now lose the 0.  Thus Harv0 becomes Harv, SelfGrow0 becomes SelfGrow, etc.  This standard ordering and sizing means that some rows and columns may be empty but it greatly simplifies indexing in calculations later. 
#Params is the re-ordered version of Params0; it stores all original parameters for later retrieval in case any will be altered for temperature, light etc. during execution
#A parameters table created in this section permits checks on inputs and processing. It can be viewed by setting View.params=1 in Controls.txt but the code for viewing is deferred until #5 where viewing of other tables is coded.
#A component-type table, also created in this section, permits checks that all components have been classified from their parameters as intended. It can be viewed by setting View.comp=1 in Controls.txt, with viewing code deferred until #5.  A stray value or NA in the input might have caused missclassification.
   #SUBSECTIONS
      #4.1 Markers of re-ordered row & column position in Params (.RC)
      #4.2 Standardise column- and row-orderings of matrix parameters
      #4.3 Logical variables relating to column ordering of Params
      #4.4 Numbers of each type of component found in Params
      #4.5 Names of each type of component in the same order as they are in Params
      #4.6 Create Params as an archive of supplied values
      #4.7 Component pairings
      #4.8 Adjustment of SelfGrow with L.index, and T.index if supplied
      #4.9 Adjustment of Harv with T.index if supplied
      #4.10 Create Component-type table
   
#4.1 Markers of re-ordered column positions in Params
#Note: logical vectors etc, dependent on matrices Harv, Assim, etc are re-ordered after the matrices have been re-ordered in #4.3
NewParamsOrder <- array(data=c(which(is.CompType0[1, ]), which(is.CompType0[2, ]), which(is.CompType0[3, ]), which(is.CompType0[4, ]), which(is.CompType0[5, ]), which(is.CompType0[6, ])), dim=c(1,N.Components))  #the sequence of column numbers, as read from Params.file, needed for the standard ordering by component type
Components <- Components0[NewParamsOrder]                                                                                              #Components0 now re-ordered

#4.2 Standardise column- and row-orderings of parameters
Harv <- array(data=0, dim=c(N.Components, N.Components), dimnames=list(paste0("HarvFrom.", Components), paste0("by.", Components)))    #the Harv matrix initially filled with zeros
Assim <- array(data=0, dim=c(N.Components, N.Components), dimnames=list(paste0("AssimOf.", Components), paste0("By.", Components)))    #the Assim matrix initially filled with zeros
Growth <- array(data=0, dim=c(N.Components, N.Components), dimnames=list(paste0("GrowthFrom.", Components), paste0("By.", Components)))#the Growth matrix initially filled with zeros
Harv0[ , ] <- Harv0[ , NewParamsOrder]                                                                                                 #re-order Harv0 cols to standard ordering
colnames(Harv0) <- paste0("by.", Components)                                                                                           #adjust the col names of Harv0
Assim0[ , ] <- Assim0[ , NewParamsOrder]                                                                                               #re-order Assim0 cols to standard ordering
colnames(Assim0) <- paste0("by.", Components)                                                                                          #adjust the col names of Assim0
#Move rows of Harv0 and Assim0 to their final row positions in Harv and Assim; the latter may have many all-zero rows
for(Rharv0 in 1:nrow2(Harv0, N.Components))                                                                                            #For each row of Harv0, [Note: nrow2() works like nrow() but also processes 1-row row vectors]
   {Rharv <- which(Components == Harv0.suffixes[Rharv0])                                                                               #Rharv is the row in Harv to receive row Rharv0 of Harv0
   Harv[Rharv, ] <- Harv0[Rharv0, ]                                                                                                    #move given coeffs from Harv0 to the ordered row of Harv matrix
   Rassim <- which(Components == Assim0.suffixes[Rharv0])                                                                              #Rassim is the row in Assim to receive row Rassim0 of Assim0
   Assim[Rharv, ] <- Assim0[Rharv0, ]                                                                                                  #move given coeffs from Assim0 to the ordered row of Assim matrix
   }
Growth[ , ] <- Harv * Assim                                                                                                            #Growth matrix now ordered as Harv and Assim.  +ve coeffs => food-consumer pair
InverseAssim <- array(data=1/Assim, dim=c(N.Components, N.Components), dimnames=list(paste0("InverseAssimOf.", Components), Components)) #InverseAssim estimates pHarvests from pGP; Inf replace zeros from Assim
Bmass.init <- array(data=Bmass0[1, NewParamsOrder], dim=c(1, N.Components), dimnames=list("Biomass.init", Components))                 #initial biomasses stored as a row.vec
Harv.suffixes <- Components                                                                                                            #new Harv. suffixes (referring to potential foods or UAH); also serve for Assim, Growth, etc
SelfGrow <- array(data=SelfGrow0[1, NewParamsOrder], dim=c(1, N.Components), dimnames=list("SelfGrow", Components))                    #SelfGrow coeffs stored as row.vec
SelfLoss <- array(data=SelfLoss0[1, NewParamsOrder], dim=c(1, N.Components), dimnames=list("SelfLoss", Components))                    #SelfLoss coeffs stored as row.vec
Gmax <- array(data=Gmax0[1, NewParamsOrder], dim=c(1, N.Components), dimnames=list("Gmax", Components))                                #row.vec; Gmax may be default values only
Hmax <- array(data=Hmax0[1, NewParamsOrder], dim=c(1, N.Components), dimnames=list("Hmax", Components))                                #row.vec; Hmax may be default values only
ParticulateSL <- array(data=ParticulateSL0[1, NewParamsOrder], dim=c(1, N.Components), dimnames=list("ParticulateSL", Components))     #row.vec; PSL may be default values only
if(N.Nutr > 0)                                                                                                                         #If there are nutrients in the model, order Nconc rows the same as Nmasses
   {Nutrients <- Nmass0.suffixes                                                                                                       #nutrients are named from the Nmass suffixes found in Params.file
   Nmass.init <- array(data=Nmass0[ , NewParamsOrder], dim=c(N.Nutr, N.Components), dimnames=list(Nutrients, Components))              #Nmass.init is made into a standardised array
   Nconc <- array(data=0, dim=c(N.Nutr, N.Components), dimnames=list(paste0("Nconc.", Nutrients), paste0("in.", Components)))          #create the Nconc matrix filled with zeros
#Make row-ordering of Nconc the same as Nmasses:-
   for(Rnconc0 in 1:N.Nutr)                                                                                                            #For each nutrient identified from Nmasses,
      {Rnconc <- which(Nutrients == Nconc0.suffixes[Rnconc0])                                                                          #the row in Nconc to receive row in Nconc0 
      Nconc[Rnconc, ] <- Nconc0[Rnconc0, NewParamsOrder]                                                                               #transfer Nconc0 row to ordered Nconc row
      }
   if(sum(is.CompType0[4, ]) > 0)                                                                                                      #If NLO is in the model:-
      {Nconc[ , is.CompType0[4, ][NewParamsOrder]] <- Nmass.init[ , is.CompType0[4, ][NewParamsOrder]] / Bmass.init[1, is.CompType0[4, ][NewParamsOrder]] #calculate nutr concs in NLO from Nmass and Bmass of NLO
      }
   }

Effort.init <- array(data=NA, dim=c(1, N.Components), dimnames=list("Effort.init", Components))                                        #dummy fishing effort data, in case called later
Quota.init <- array(data=NA, dim=c(1, N.Components), dimnames=list("Quota.init", Components))                                          #dummy quota data, in case called later
ActiveQuotaFisheries <- NA                                                                                                             #dummy act.q.f
ActiveQuotaFisheries.R <- NA                                                                                                           #dummy act.q.f
if(N.Fisheries > 0)                                                                                                                    #If there are fisheries in the model,
   {Effort.init <- array(data=Effort0[1, NewParamsOrder], dim=c(1, N.Components), dimnames=list("Effort.init", Components))            #sort the Effort0 row columnwise as for Fisheries, and put into Effort.init 
   if(N.Quota0.Rpar > 0)                                                                                                               #If quota rows are supplied for fisheries:-
      {Quota.init <- array(data=Quota0[ , NewParamsOrder], dim=c(N.Fisheries, N.Components), dimnames=list(Fisheries, Components))     #sort Quota0 (from #2.9) columnwise and transfer to Quota.init
      ActiveQuotaFisheries.R <- NULL                                                                                                   #stub to add to
      for(Q.R in 1:N.Fisheries)                                                                                                        #For each row in Quota0,
         {if(any(Quota0[Q.R, ] < Inf))                                                                                                 #If there is >0 sub-Inf quota, (+ve fishing effort has already been tested for),
             {ActiveQuotaFisheries.R <- c(ActiveQuotaFisheries.R, Q.R)                                                                 #this fishery is an active quota fishery
             }
         }
      ActiveQuotaFisheries <-  Fisheries[ActiveQuotaFisheries.R]                                                                       #names of active quota fisheries; same as corresponding quota.suffixes; ordered as fisheries
      }
   }                                                              

if(N.Habitats > 0)                                                                                                                     #If habitats are in the model,
   {HabSize.init <- HabSize0[1, NewParamsOrder]                                                                                        #row.vec N.Habitats wide
   HabOccup <- array(data=0, dim=c(N.Habitats, N.Components), dimnames=list(paste0("HabOccup.", Habitats), Components))                #create HabOccup array; habitats are row-ordered as in habitat component columns
   for(Rhabocc in 1:N.Habitats)                                                                                                        #For each Habitat,
      {Rhabocc0 <- which(Occup.suffixes == Habitats[Rhabocc])                                                                          #find the source row in HabOccup0,
      HabOccup[Rhabocc, ] <- HabOccup0[Rhabocc0, NewParamsOrder]                                                                       #transfer the source row to HabOccup
      }
   Occup.suffixes <- Habitats                                                                                                          #update occupancy suffixes to same as Habitats
   }

alphaSG.I <- array(data=alphaSG.I0[1, NewParamsOrder], dim=c(1, N.Components), dimnames=list("alphaSG.I", Components))                 #slopes of self-growth over light index between bounds
LoBoundSG.T <- array(data=LoBoundSG.T0[1, NewParamsOrder], dim=c(1, N.Components), dimnames=list("LoBoundSG.T", Components))           #lower T.index value below which self-growth is zero
HiBoundSG.T <- array(data=HiBoundSG.T0[1, NewParamsOrder], dim=c(1, N.Components), dimnames=list("HiBoundSG.T", Components))           #upper T.index above which s-g is zero
alphaHvst.T <- array(data=alphaHvst.T0[1, NewParamsOrder], dim=c(1, N.Components), dimnames=list("alphaHvst.T", Components))           #slopes of Harv coeffs (all in the col) over T.index between bounds
LoBoundHvst.T <- array(data=LoBoundHvst.T0[1, NewParamsOrder], dim=c(1, N.Components), dimnames=list("LoBoundHvst.T", Components))     #lower T.index value below which Harv is zero
HiBoundHvst.T <- array(data=HiBoundHvst.T0[1, NewParamsOrder], dim=c(1, N.Components), dimnames=list("HiBoundHvst.T", Components))     #upper T.index above which Harv is zero

#4.3 Logical variables relating to column ordering of Params
is.Auto <- array(data=is.CompType0[1, NewParamsOrder], dim=c(1, N.Components), dimnames=list("is.Autotroph", Components))      #is.Auto0 (from #2) re-ordered
is.Hetero <- array(data=is.CompType0[2, NewParamsOrder], dim=c(1, N.Components), dimnames=list("is.Heterotroph", Components))  #is.Hetero0 re-ordered
is.Grazer <- array(data=as.logical(Csums(Growth[is.Auto, ] > 0, sum(is.Auto), N.Components)), dim=c(1, N.Components), dimnames=list("is.Grazer", Components)) #grazers are heteros whose foods include autos
is.Fishery <- array(data=is.CompType0[3, NewParamsOrder], dim=c(1, N.Components), dimnames=list("is.Fishery", Components))     #is.Fishery0 re-ordered
is.Consumer <-  array(data=colSums(Growth) > 0, dim=c(1, N.Components), dimnames=list("is.Consumer", Components))              #TRUE if growing by harvesting and assimilating of any other; incl fisheries, scavengers
is.Wild <- array(data=is.Auto | is.Hetero, dim=c(1, N.Components), dimnames=list("is.Wild", Components))                       #is.Wild0 re-ordered
is.NLO <- array(data=is.CompType0[4, NewParamsOrder], dim=c(1, N.Components), dimnames=list("is.NLO", Components))             #is.NLO0 re-ordered
is.WildOrNLO <- array(data=is.Wild | is.NLO, dim=c(1, N.Components), dimnames=list("is.WildOrNLO", Components))                #used to locate biomasses and foods in Harv, Assim, Growth etc
is.Scavenger <- array(data=FALSE, dim=c(1, N.Components), dimnames=list("is.Scavenger", Components))                           #default is.Scavenger created
if(sum(is.CompType0[4, NewParamsOrder] > 0)) is.Scavenger[1, ] <- Growth[is.CompType0[4, NewParamsOrder], ] > 0                #TRUE if comp has +ve Growth coeff for NLO
is.Harvester <- array(data=is.Harvester0[1, NewParamsOrder], dim=c(1, N.Components), dimnames=list("is.Harvester", Components))   #is.Harvester0 re-ordered
is.Harvest <- array(data=rowSums(Harv) > 0, dim=c(1, N.Components), dimnames=list("is.Harvest", Components))                   #TRUE if a component is harvested by any other; not necessarily eaten by it
is.WildHarvest <- array(data=is.Wild & is.Harvest, dim=c(1, N.Components), dimnames=list("is.WildHarvest", Components))        #TRUE if a component is harvested by any other; not necessarily eaten by it
is.DroppedHarvest <- array(data=Rsums(Harv[ , is.Harvester] > 0 & Assim[ , is.Harvester] < 1, N.Components, sum(is.Harvester)) > 0, dim=c(1, N.Components), dimnames=list("is.DroppedHarvest", Components)) #TRUE if comp harvested but not totally assimilated; Rsums() used because works for 1 or more Harvesters (rowSums does not work for 1)
is.UnAssimHarvest <- array(data=rowSums(Harv > 0 & Assim == 0) > 0, dim=c(1, N.Components), dimnames=list("is.UnAssimilatedHarvest", Components)) #TRUE if comp harvested by any other with zero assimilation
is.UnAssimHarvester <- array(data=ifelse(colSums(Harv > 0 & Assim == 0) > 0, TRUE, FALSE), dim=c(1, N.Components), dimnames=list("is.UnAssimilatingHarvester", Components)) #TRUE if comp harvesting others with zero assimilation
is.UnAssimHarv.pair <- Harv > 0 & Assim == 0                                                                                   #used to locate UAH.pairs in Comps X Comps matrices, eg. Harv
is.Gainer <- array(data=is.WildOrNLO | is.Fishery, dim=c(1, N.Components), dimnames=list("is.Gainer", Components))             #used to locate Gainers in Harv, Assim, etc ; includes NLO
is.Food <- array(data=ifelse(rowSums(Growth) > 0, TRUE, FALSE), dim=c(1, N.Components), dimnames=list("is.Food", Components))  #create vector to indicate food components, ie. Harvested and eaten cf. is.Harvest
is.WildFood <- array(data=is.Food & is.Wild, dim=c(1, N.Components), dimnames=list("is.WildFood", Components))                 #wild, living foods (ie. not NLO) defined
is.Predator <- array(data=is.Hetero & !is.Grazer & !is.Scavenger, dim=c(1, N.Components), dimnames=list("is.Predator", Components)) #Preds are heteros whose foods include other heteros
#is.Predator[1, is.Hetero] <- is.Hetero & !is.Grazer & !is.Scavenger       #ifelse(colSums(Growth[is.Hetero & !is.Grazer & !is.Scavenger, is.Hetero]) > 0, TRUE, FALSE)       #defined by +ve hetero col sums of wild, non-graz, non-scav rows of Growth; fisheries excluded
is.TopPredator <- array(data=rowSums(Growth)==0 & is.Predator, dim=c(1, N.Components), dimnames=list("is.TopPredator", Components)) #create array
is.Landed <- array(data=FALSE, dim=c(max(N.Fisheries, 1), N.Components), dimnames=list(paste0("is.LandedBy.", Components[is.Fishery]), Components))#dummy: components landable by any fishery
is.Caught <- array(data=FALSE, dim=c(max(N.Fisheries, 1), N.Components), dimnames=list(paste0("is.CaughtBy.", Components[is.Fishery]), Components))#default components catchable by any fishery
is.Discarded <- array(data=FALSE, dim=c(max(N.Fisheries, 1), N.Components), dimnames=list(paste0("is.DiscardedBy.", Components[is.Fishery]), Components))#default components discarded by any fishery
is.ActiveQuotum <- array(data=FALSE, dim=c(max(N.Fisheries, 1), N.Components), dimnames=list(paste0("is.ActiveQuotum.", Fisheries), Components)) #dummy logical var, in case called later
if(N.Fisheries > 0)                                                                                                            #If fisheries are in the model,
   {is.ActiveQuotum <- array(data=FALSE, dim=c(N.Fisheries, N.Components), dimnames=list(paste0("is.ActiveQuotum.", Fisheries), Components))                  #dummy logical var, in case called later
   is.Landed[ , ] <- t(Growth[ , is.Fishery] > 0 )                                                                             #Growth in fishery col is > 0 in rows that are landed; result is TRUE
   is.Caught[ , ] <- t(Harv[ , is.Fishery] > 0 )                                                                               #Harv in fishery col is > 0 in rows that are caught; result is TRUE
   is.Discarded[ , ] <- t((Harv - Growth)[ , is.Fishery] > 0 )                                                                 #Harv - Growth in fishery col is > 0 in rows that are discarded including UAH; result is TRUE
   if(N.Quota0.Rpar > 0)                                                                                                       #If quota are in the model,
      {is.QuotaFishery <- array(data=Components %in% ActiveQuotaFisheries, dim=c(1, N.Components), dimnames=list("is.QuotaFishery", Components)) #fisheries having active (fished) quota for > 0 wilds are TRUE
      is.ActiveQuotum[ActiveQuotaFisheries.R, ] <- ifelse(Quota.init[ActiveQuotaFisheries.R, ] < Inf, TRUE, FALSE)             #logical indicator of active quota
      }
   }
is.WildOrFishery <- array(data=(is.Wild | is.Fishery), dim=c(1, N.Components), dimnames=list("is.WildOrFishery", Components))  #is the component a Wild or a Fishery?
is.UnfishedTopPredator  <- array(data=is.TopPredator & !as.logical(Csums(is.Caught, N.Fisheries, N.Components)), dim=c(1, N.Components), dimnames=list("is.UnfishedTopPredator", Components)) #UFTPs=wilds not food or caught
is.Habitat <- array(data=is.Habitat0[1, NewParamsOrder], dim=c(1, N.Components), dimnames=list("is.Habitat", Components))      #is.Habitat0 re-ordered
is.H2O <- array(data=is.CompType0[6, NewParamsOrder], dim=c(1, N.Components), dimnames=list("is.H2O", Components))             #is.H2O0 re-ordered
is.alphaSG.I <- is.alphaSG.I0[NewParamsOrder]                                                                                  #is slope of SelfGrow0 over L.index re-ordered 
is.LoBoundSG.T <- is.LoBoundSG.T0[NewParamsOrder]                                                                              #is LoBound for SG0 adjustments with T.index re-ordered
is.HiBoundSG.T <- is.HiBoundSG.T0[NewParamsOrder]                                                                              #is HiBound for SG0 adjustments with T.index re-ordered
is.alphaHvst.T <- is.alphaHvst.T0[NewParamsOrder]                                                                              #is slope of Hvst0 over T.index re-ordered 
is.LoBoundHvst.T <- is.LoBoundHvst.T0[NewParamsOrder]                                                                          #is LoBound for Harv0 adjustments with T.index re-ordered
is.HiBoundHvst.T <- is.HiBoundHvst.T0[NewParamsOrder]                                                                          #is HiBound for Harv0 adjustments with T.index re-ordered


#4.4 Numbers of each type of component found in Params
N.Autos <- sum(is.Auto)                                                                                                        #number of autotrophs, etc.
N.Heteros <- sum(is.Hetero)
N.Grazers <- sum(is.Grazer)
N.Fisheries <- sum(is.Fishery)
N.Consumers <- sum(is.Consumer)
N.Wilds <- sum(is.Wild)
N.NLOs <- sum(is.NLO)                                                                                                          #only 1 NLO component allowed in this version of ECOLPS
N.WildOrNLOs <- sum(is.WildOrNLO)
N.Harvesters <- sum(is.Harvester)
N.Harvests <- sum(is.Harvest)
N.WildHarvests <- sum(is.WildHarvest)
N.DroppedHarvests <- sum(is.DroppedHarvest)
N.UnAssimHarvests <- sum(is.UnAssimHarvest)
N.UnAssimHarvesters <- sum(is.UnAssimHarvester)
N.UnAssimHarv.pairs <- sum(is.UnAssimHarv.pair)
N.Gainers <- sum(is.Gainer)
N.Foods <- sum(is.Food)
N.WildFoods <- sum(is.WildFood)
N.Predators <- sum(is.Predator)
N.TopPredators <- sum(is.TopPredator)
N.Landeds <- sum(is.Landed)
N.Caughts <- sum(is.Caught)
N.Discardeds <- sum(is.Discarded)
N.WildOrFisheries <- sum(is.WildOrFishery)
#N.LandingFisheries <- sum(is.LandingFishery)
N.QuotumFishery.pairs <- sum(is.ActiveQuotum)                                    
N.ActiveQuotaFisheries <- ifelse(is.na(ActiveQuotaFisheries[1]), 0, length(ActiveQuotaFisheries))                                                                           
N.ActiveQuota <- sum(is.ActiveQuotum)                                                                                            
N.UnfishedTopPredators <- sum(is.UnfishedTopPredator)
N.Scavengers <- sum(is.Scavenger)
N.Habitats <- sum(is.Habitat)
N.H2Os <- sum(is.H2O)                                                                                                         #only 1 H2O component allowed in this version of ECOLPS


#4.5 Names of each type of component in the same order as they are in Params
#Omitted component types (Fisheries, Habitats) were defined in #2. because they were needed previously; ditto for Nutrients
Autotrophs <- Components[is.Auto]                                                                            #names of autotrophs, etc.   is.Auto and others below from #4.3
Heterotrophs <- Components[is.Hetero]
Grazers <- Components[is.Grazer]
Consumers <- Components[is.Consumer]
Wilds <- Components[is.Wild]
NLOs <- Components[is.NLO]
WildOrNLOs <- Components[is.WildOrNLO]
Harvesters <- Components[is.Harvester]
Harvests <- Components[is.Harvest]
WildHarvests <- Components[is.WildHarvest]
DroppedHarvests <- Components[is.DroppedHarvest]                                                             
UnAssimHarvests <- Components[is.UnAssimHarvest]
UnAssimHarvesters <- Components[is.UnAssimHarvester]                                                         #define vector
UnAssimHarv.pairs <- vector("character", length=N.UnAssimHarv.pairs)                                         #vector of unassimilated harvested>harvester pairs
UnAssimHarv.pairs <- NULL                                                                                    #stub vector to be added to
if(N.UnAssimHarv.pairs > 0)                                                                                  #If some components are harvested but not assimilated (do not contribute to growth)
   {for(uu in 1:N.UnAssimHarv.pairs)
      {UnAssimHarv.pairs[uu] <-paste0(Components[which(is.UnAssimHarv.pair, arr.ind=TRUE)[uu, 1]], ">", Components[which(is.UnAssimHarv.pair, arr.ind=TRUE)[uu, 2]]) #create names for each pair as 'harvested>harvester'
      }
   }
ifelse(length(UnAssimHarvests[ ]) > 0 & length(UnAssimHarvesters[ ]) > 0, paste0(Components[which(is.UnAssimHarv.pair, arr.ind=TRUE)[ , 1]], ">", Components[which(is.UnAssimHarv.pair, arr.ind=TRUE)[ , 2]]), NA)
Gainers <- Components[is.Gainer]
Foods <- Components[is.Food]
WildFoods <- Components[is.WildFood]
Predators <- Components[is.Predator]
TopPredators <- Components[is.TopPredator]
Landeds <- Components[is.Landed]
Caughts <- Components[is.Caught]
Discardeds <- Components[is.Discarded]
WildOrFisheries <- Components[is.WildOrFishery]
#ActiveQuotaFisheries <- Components[is.QuotaFishery]                                                         #previously defined in #4.2
UnfishedTopPredators <- Components[is.UnfishedTopPredator]
Scavengers <- Components[is.Scavenger]
H2Os <- Components[is.H2O]
NewParamsOrder.names <- c(Autotrophs, Heterotrophs, Fisheries, NLOs, Habitats, H2Os)                         #Autotrophs, Fisheries etc are vectors of individual component names
dimnames(NewParamsOrder) <- list("Params0cols", NewParamsOrder.names)                                        #row and col headers for NPO


#4.6 Create Params as an archive of supplied values
#These are for display, and for reference purposes if any param values are changed later by an Environ.file
Params <- array(data=0, dim=c(3 + 3 * N.Components, N.Components), dimnames=list(c("SelfGrow", "SelfLoss", "Bmass.init", row.names(Harv), row.names(Assim), row.names(Growth)), Components))
Rpar.SG <- 1                                                                    #create Params with SG as the first row; Rpar is for row location in Params
Params[Rpar.SG, ] <- SelfGrow[1, ]                                              #store SelfGrow at Rpar.SG
Rpar.SL <- 2                                                                    #row for SeflLoss
Params[Rpar.SL, ] <- SelfLoss[1, ]                                              #store SelfLoss at Rpar.SL
Rpar.Bmass <- 3                                                                 #row for Bmass
Params[Rpar.Bmass, ] <- Bmass.init[1, ]                                         #store Bmass
Rpar.Harv <- 4:(3 + N.Components)                                               #Harvest matrix stored at these rows
Params[Rpar.Harv, ] <- Harv                                                     #store Harv in the Gainer cols; note- Harv not Harv0 is stored because Harv has a full set of rows, 1 per wild or NLO component
LastR <- max(Rpar.Harv)
Rpar.Assim <- (LastR + 1):(LastR + N.Components)                                #Assimilation matrix
Params[Rpar.Assim, ] <- Assim
LastR <- max(Rpar.Assim)
Rpar.Growth <- (LastR + 1):(LastR + N.Components)                               #Growth matrix
Params[Rpar.Growth, ] <- Growth
LastR <- max(Rpar.Growth)       
#Next: row-bind on optional parameters if present
if(!is.na(Gmax0.Rpar))                                                          #if a Gmax row is present, store
   {Rpar.Gmax <- LastR + 1
   Params <- rbind(Params, vector(length=N.Components, "numeric"))
   Params[Rpar.Gmax, ] <- Gmax 
   LastR <- Rpar.Gmax
   row.names(Params)[LastR] <- "Gmax"
   }
if(!is.na(Hmax0.Rpar))                                                          #if an Hmax row is present, store
   {Rpar.Hmax <- LastR + 1
   Params <- rbind(Params,  vector(length=N.Components, "numeric"))
   Params[Rpar.Hmax, ] <- Hmax
   LastR <- Rpar.Hmax
   row.names(Params)[LastR] <- "Hmax"
   }
if(!is.na(ParticulateSL0.Rpar))                                                 #if a particulate self loss row is present, store
   {ParticulateSL[is.NLO] <- 0                                                  #make sure that PSL is zero for NLO
   Rpar.ParticulateSL <- LastR + 1
   Params <- rbind(Params,  vector(length=N.Components, "numeric"))
   Params[Rpar.ParticulateSL, ] <- ParticulateSL
   LastR <- Rpar.ParticulateSL
   row.names(Params)[LastR] <- "ParticulateSL"
   }
if(N.Nutr > 0)                                                                  #if nutrient-mass rows are present, store
   {Params <- rbind(Params, array(data=0, dim=c(2 * N.Nutr, N.Components), dimnames=list(c(paste0("Nmass.init.", Nutrients), paste0("Nconc.", Nutrients)), Components)))
   Rpar.Nmass.init <- (LastR + 1):(LastR + N.Nutr)
   Params[Rpar.Nmass.init, ] <- Nmass.init[1:N.Nutr, ]
   LastR <- max(Rpar.Nmass.init)
   Rpar.Nconc <- (LastR + 1):(LastR + N.Nutr)                                   #also store nutrient-conc rows
   Params[Rpar.Nconc, ] <- Nconc[1:N.Nutr, ]
   LastR <- max(Rpar.Nconc)                                      
   }
if(N.Fisheries > 0)                                                             #if fishing-effort rows are present, store
   {Params <- rbind(Params, array(data=0, dim=c(1, N.Components), dimnames=list("Effort.init", Components)))
   Rpar.Effort.init <- LastR + 1
   Params[Rpar.Effort.init, ] <- Effort.init[1, ]
   LastR <- Rpar.Effort.init
   if(N.ActiveQuotaFisheries > 0)                                               #also store fisheries quota if any,
      {Params <- rbind(Params, array(data=Inf, dim=c(N.ActiveQuotaFisheries, N.Components), dimnames=list(paste0("Quota.init.", ActiveQuotaFisheries), Components)))
      Rpar.Quota <- (LastR + 1):(LastR + N.ActiveQuotaFisheries)
      Params[Rpar.Quota, ] <- Quota.init[ActiveQuotaFisheries.R, ]
      LastR <- max(Rpar.Quota)
      }
   }
if(N.Habitats > 0)                                                              #if Habitat rows are present, store
   {Params <- rbind(Params, array(data=0, dim=c(1+ N.Habitats, N.Components), dimnames=list(c("HabitatSize", paste0("HabOccup.", Habitats)), Components)))
   Rpar.HabSize <- (LastR + 1):(LastR + 1)
   Params[Rpar.HabSize, ] <- HabSize.init                                       #store habitat size 
   LastR <- Rpar.HabSize
   Rpar.Occup <- (LastR + 1):(LastR + N.Habitats)                 
   Params[Rpar.Occup, ] <- HabOccup[1:N.Habitats, ]                             #store occupancy coeffs
   LastR <- max(Rpar.Occup)
   }
if(!is.na(T.index0.Rpar))                                                       #if temperature-index rows are present, store
   {T.index.init <- T.index0[1, NewParamsOrder][is.H2O]                         #scalar
   Params <- rbind(Params, array(data=0, dim=c(1, N.Components), dimnames=list("T.index.init", Components)))
   Rpar.T.index <- (LastR + 1):(LastR + 1)
   Params[Rpar.T.index, is.H2O] <- T.index.init
   LastR <- Rpar.T.index
   }
if(!is.na(L.index0.Rpar))                                                       #if light-index rows are present, store
   {L.index.init <- L.index0[1, NewParamsOrder][is.H2O]                         #scalar
   Params <- rbind(Params, array(data=0, dim=c(1, N.Components), dimnames=list("L.index.init", Components)))
   Rpar.L.index <- (LastR + 1):(LastR + 1)
   Params[Rpar.L.index, is.H2O] <- L.index.init
   LastR <- Rpar.L.index
   }
if(!is.na(alphaSG.I0.Rpar))                                                     #if SelfGrow-adjustment rows are present, store
   {Params <- rbind(Params, array(data=0, dim=c(1, N.Components), dimnames=list("alphaSG.I", Components)))
   Rpar.alphaSG.I <- (LastR + 1):(LastR + 1)
   Params[Rpar.alphaSG.I, ] <- alphaSG.I                                                                                           
   LastR <- Rpar.alphaSG.I
   }
if(!is.na(LoBoundSG.T0.Rpar))                                                   #if SelfGrow-adjustment rows are present, store
   {Params <- rbind(Params, array(data=0, dim=c(1, N.Components), dimnames=list("LoBoundSG.T", Components)))
   Rpar.LoBoundSG.T <- (LastR + 1):(LastR + 1)
   Params[Rpar.LoBoundSG.T, ] <- LoBoundSG.T                                                                                           
   LastR <- Rpar.LoBoundSG.T
   }
if(!is.na(HiBoundSG.T0.Rpar))                                                   #if SelfGrow-adjustment rows are present, store
   {Params <- rbind(Params, array(data=0, dim=c(1, N.Components), dimnames=list("HiBoundSG.T", Components)))
   Rpar.HiBoundSG.T <- (LastR + 1):(LastR + 1)
   Params[Rpar.HiBoundSG.T, ] <- HiBoundSG.T                                                                                           
   LastR <- Rpar.HiBoundSG.T
   }
if(!is.na(alphaHvst.T0.Rpar))                                                   #if Harv-adjustment rows are present, store
   {Params <- rbind(Params, array(data=0, dim=c(1, N.Components), dimnames=list("alphaHvst.T", Components)))
   Rpar.alphaHvst.T <- (LastR + 1):(LastR + 1)
   Params[Rpar.alphaHvst.T, ] <- alphaHvst.T                                                                                           
   LastR <- Rpar.alphaHvst.T
   }
if(!is.na(LoBoundHvst.T0.Rpar))                                                 #if Harv-adjustment rows are present, store
   {Params <- rbind(Params, array(data=0, dim=c(1, N.Components), dimnames=list("LoBoundHvst.T", Components)))
   Rpar.LoBoundHvst.T <- (LastR + 1):(LastR + 1)
   Params[Rpar.LoBoundHvst.T, ] <- LoBoundHvst.T                                                                                           
   LastR <- Rpar.LoBoundHvst.T
   }
if(!is.na(HiBoundHvst.T0.Rpar))                                                 #if Harv-adjustment rows are present, store
   {Params <- rbind(Params, array(data=0, dim=c(1, N.Components), dimnames=list("HiBoundHvst.T", Components)))
   Rpar.HiBoundHvst.T <- (LastR + 1):(LastR + 1)
   Params[Rpar.HiBoundHvst.T, ] <- HiBoundHvst.T                                                                                           
   LastR <- Rpar.HiBoundHvst.T
   }
#----------------------------------------------------------------


#4.7 Component pairings
Temp1 <- expand.grid(Components, Components)                                                                 #Temp1 gives a 2-col matrix with all permutations of component names
AllPossHarvests <- noquote(as.character(Temp1[ , 1]))                                                        #AllPossHarvests is the first col of Temp1
AllPossHarvesters <- noquote(as.character(Temp1[ , 2]))                                                      #AllPossHarvesters is the second col of Temp1
Temp2 <- paste(Temp1[ , 1], Temp1[ , 2], sep=">")                                                            #Temp2 has the 2 names pasted together in each row with ">" between to symbolise 'consumed by' (or supplied to, in the case of organic biomass dropping to NLO)
AllPairs <- array(data=Temp2, dim=c(N.Components, N.Components))                                             #array filled with Temp2; the pairs are selected from below
AllPossHarvests.array <- array(data=AllPossHarvests, dim=c(N.Components, N.Components))                      #AllPossHarvests is now a matrix to match with Growth; row1 has all elements=Comp1, row2 has all=Comp2, etc
AllPossHarvesters.array <- array(data=AllPossHarvesters, dim=c(N.Components, N.Components))                  #AllPossHarvesters is now a matrix to match with Growth; col1 has all elements=Comp1, col2 has all=Comp2, etc
FoodConsumer.pairs <- AllPairs[which(Growth > 0, arr.ind=TRUE)]                                              #all pairs of the type "food>consumer" ; consumers defined as assimilating their harvests, ie. Assim > 0
FoodHeterotroph.pairs <- AllPairs[ , is.Hetero][which(Growth[ , is.Hetero] > 0, arr.ind=TRUE)]               #all pairs of the type "food>heterotroph" ; consumers that are not fisheries
AutoGrazer.pairs <- AllPairs[which(Growth[is.Auto, ] > 0, arr.ind=TRUE)]                                     #all pairs of the type "auto>grazer"
HarvestHarvester.pairs <- AllPairs[which(Harv > 0, arr.ind=TRUE)]                                            #a 'harvester' kills other components;  harvesters may/not assimilate harvests; Cons are a subset of Harvesters
UnAssimHarv.pairs <- AllPairs[which(Harv > 0 & Assim == 0, arr.ind=TRUE)]                                    #unassimilated harvest pairs with Assim=0 ; UAH union Foods = Harvests  SEE #4.5
FoodFishery.pairs <- AllPairs[which(Growth > 0 & AllPossHarvesters.array %in% Fisheries, arr.ind=TRUE)]      #foods>fisheries
QuotumFishery.pairs <- NA                                                                                    #dummy value
if(N.ActiveQuotaFisheries > 0) QuotumFishery.pairs <- paste0(row.names(t(Quota.init))[which(t(Quota.init) < Inf, arr.ind=TRUE)[ , 1]], ">", ActiveQuotaFisheries[which(t(Quota.init) < Inf, arr.ind=TRUE)[ , 2]]) #quotum food x fishery pairs, ordered by fisheries
NLOScavenger.pairs <- AllPairs[which(Growth > 0 & AllPossHarvesters.array %in% Scavengers, arr.ind=TRUE)]    #NLO>scavengers
N.FoodConsumer.pairs <- length(FoodConsumer.pairs)                                                           #number of food>consumer pairs, etc.
N.FoodHeterotroph.pairs <- length(FoodHeterotroph.pairs)
N.HarvestHarvester.pairs <- length(HarvestHarvester.pairs)
N.FoodFishery.pairs <- length(FoodFishery.pairs)
N.QuotumFishery.pairs <- ifelse(is.na(QuotumFishery.pairs[1]), 0, length(QuotumFishery.pairs))                                                        


#4.8. Adjustment of SelfGrow with L.index, and T.index if supplied
#If an alphaSG.I row was provided in Params.file, adjust some or all initial (t=0) autotrophic SelfGrow for L.index.init; adjustments only occur in auto cols where alphaSG.I is +ve
#If no alphaSG.I row was provided, all SelfGrow values remain unaltered by changing L.index values read from Environ.file
#The originally supplied values of SelfGrow were stored, unaltered, in Params for use in later adjustments; see #4.6
#First, adjust the initial SelfGrow value supplied in Params.file using L.index.init if provided
if(!is.na(L.index0.Rpar))                                                                                    #If a light index value for t=0 was provided,
   {Advise(WithRefTo=NA, ErrText=NA, AdvText=c(paste0(length(L.index0.Rpar), "  initial light.index value for water found: "), L.index.init), ShowData=NULL, CallSection=NA, Fatal=FALSE, MinAdviceLevel2Show=1) #advise finding
   Advise(WithRefTo=NA, ErrText=NA, AdvText=c("Autotrophic SelfGrow coeffs as read: ", SelfGrow[1, is.Auto]), ShowData=NULL, CallSection=NA, Fatal=FALSE, MinAdviceLevel2Show=2)
   if(!is.na(alphaSG.I0.Rpar))                                                                               #If slope(s) for adjustment over L.index values (0 to 1) were provided,
      {Advise(WithRefTo=NA, ErrText=NA, AdvText=c("alphaSG.I coeffs (-> rate of increase of autotrophic SelfGrow over L.index) as read, 1 per autotroph: ", alphaSG.I[1, is.Auto]), ShowData=NULL, CallSection=NA, Fatal=FALSE, MinAdviceLevel2Show=2)
      SelfGrow[1, is.Auto] <- ifelse(alphaSG.I[1, is.Auto] > 0, Params[Rpar.SG, is.Auto] * alphaSG.I[1, is.Auto] * L.index.init, SelfGrow[1, is.Auto])#adjust auto elements of SelfGrow using slopes > 0 over L.index or leave as was
      Advise(WithRefTo=NA, ErrText=NA, AdvText=c("Autotrophic SelfGrow coeffs after adjustment for initial L.index (given alphaSG.I > 0): ", round(SelfGrow[1, is.Auto], 4)), ShowData=NULL, CallSection=NA, Fatal=FALSE, MinAdviceLevel2Show=1)
      }
   }   
#Next, adjust SelfGrow within Lo and HiBounds of T.index if provided. This clipping of SelfGrow with T.index is independent of adjustment with alphaSG.I.
#This means: only use positive SelfGrow if LoBoundSG.T <= T.index.init <= HiBoundSG.T; ie. if T.index.init is outside bounds, SelfGrow is temporarily set to a very low +ve value (to permit bound changes later) 
#The originally supplied values of the bounds were stored, unaltered, in Params for use in later adjustments; see #4.6
#AdvisedT.index <- FALSE                                                                                     #this is just to stop repeated advice about the T.index
if(!is.na(T.index0.Rpar))                                                                                    #If a temperature index value for t=0 was provided,
   {Advise(WithRefTo=NA, ErrText=NA, AdvText=c(paste0(length(Rpar.T.index), "  initial temperature.index value for water found: "), T.index.init), ShowData=NULL, CallSection=NA, Fatal=FALSE, MinAdviceLevel2Show=1) #advise 
   AdvisedT.index <- TRUE                                                                                    #advice about T.index.init has now been given
   TestTemp <- rep(T.index.init, N.Autos)                                                                    #used to test T.index boundaries for autotrophs
   names(TestTemp) <- Autotrophs                                                                             #name the test T.index.init values against bounds
   if(!is.na(LoBoundSG.T0.Rpar))                                                                             #If lower T.index bounds were provided for SelfGrow,
      {SelfGrow[1, is.Auto] <- ifelse(TestTemp >= LoBoundSG.T[1, is.Auto], SelfGrow[1, is.Auto], 0.0000000001)  #If T.index.init >= LoBound of T.index, accept +ve SG, else 0.0000000001
      Advise(WithRefTo=NA, ErrText=NA, AdvText=c("LoBound T.index for SelfGrow coeffs of autotroph(s): ", LoBoundSG.T[1, is.Auto]), ShowData=NULL, CallSection=NA, Fatal=FALSE, MinAdviceLevel2Show=2) #allow checking of calcs.
      }
   if(!is.na(HiBoundSG.T0.Rpar))                                                                             #If higher T.index bounds were provided for SelfGrow,
      {SelfGrow[1, is.Auto] <- ifelse(TestTemp <= HiBoundSG.T[1, is.Auto], SelfGrow[1, is.Auto], 0.0000000001)#If T.index.init <= HiBound of T.index, accept +ve SG, else 0
      Advise(WithRefTo=NA, ErrText=NA, AdvText=c("HiBound T.index for SelfGrow coeffs of autotroph(s): ", HiBoundSG.T[1, is.Auto]), ShowData=NULL, CallSection=NA, Fatal=FALSE, MinAdviceLevel2Show=2)
      }
   if(!is.na(LoBoundSG.T0.Rpar) | !is.na(Rpar.HiBoundSG.T))                                                  #If either or both T.index bounds were set, advise results if AL allows,
      {Advise(WithRefTo=NA, ErrText=NA, AdvText=c("Autotrophic SelfGrow AFTER zero-ing values outside bounds set from initial T.index =", T.index.init, ": ", round(SelfGrow[1, is.Auto], 4)), ShowData=NULL, CallSection=NA, Fatal=FALSE, MinAdviceLevel2Show=1)
      }
   }
#----------------------------------------------------------------


#4.9 Adjustment of Harv with T.index if supplied
#4.9.1 Adjustment of Harv values for each harvester with T.index
#If an alphaHvst.T0 row was provided in Params.file, ie. !is.na(alphaHvst.T0.Rpar)=TRUE, adjust some or all initial (t=0) heterotrophic Harv coeffs for T.index.init; adjustments only occur in hetero cols where alphaHvst.T is +ve 
#If no alphaHvst.T row was provided, all Harv values remain unaltered by changing T.index values
#The originally supplied values of Harv are stored, unaltered, in Params for use in later adjustments; see #4.6
if(!is.na(T.index0.Rpar)) 
   {if(AdvisedT.index == FALSE) 
       {Advise(WithRefTo=NA, ErrText=NA, AdvText=c(paste0(length(T.index0.Rpar), "  initial temperature.index value for water found: "), T.index.init), ShowData=NULL, CallSection=NA, Fatal=FALSE, MinAdviceLevel2Show=1) 
       AdvisedT.index <- TRUE
       }
   Advise(WithRefTo=NA, ErrText=NA, AdvText=c("Heterotrophic harvest coeffs as read: "), ShowData=NULL, CallSection=NA, Fatal=FALSE, MinAdviceLevel2Show=2)  ; if(AL==2) print(Harv[is.Food, is.Hetero])
   if(!is.na(alphaHvst.T0.Rpar))                                                                             #If slope(s) for adjustment over T.index values (0 to 1) were provided:-
      {Advise(WithRefTo=NA, ErrText=NA, AdvText=c("alphaHvst.T coeffs (rate of increase of heterotrophic harvest coeffs over T.index) as read, 1 per hetero: ", alphaHvst.T[1, is.Hetero]), ShowData=NULL, CallSection=NA, Fatal=FALSE, MinAdviceLevel2Show=2)
      for(Rhv in 1:N.Components)                                                                             #For all food rows in Harv matrix,
         {Harv[Rhv, is.Hetero] <- ifelse(alphaHvst.T[1, is.Hetero] > 0, Harv[Rhv, is.Hetero] * alphaHvst.T[1, is.Hetero] * T.index.init, Harv[Rhv, is.Hetero]) #adjust heterotrophic columns of Harv using slopes > 0 over T.index
         }
      Advise(WithRefTo=NA, ErrText=NA, AdvText=c("Harvest coeffs AFTER adjustment for initial T.index (given alphaHvst.T > 0) = ", c(T.index.init, " :-  ")), ShowData=NULL, CallSection=NA, Fatal=FALSE, MinAdviceLevel2Show=1)  ; if(AL > 0) print(Harv[is.Food, is.Hetero])                                                                     #for checks
      }

#4.9.2 Adjustment of Harv within Lo and HiBounds of T.index
#Harv may have been adjusted for temperature in #4.9.1, or not
#Only use positive Harv if LoBoundHvst.T <= T.index.init <= HiBoundHvst.T; if T.index.init is outside bounds, Harv is temporarily set to zero 
   TestTemp <- rep(T.index.init, N.Heteros)                                                                  #used to test T.index boundaries
   names(TestTemp) <- Heterotrophs
   for(Rhv in 1:N.Components)                                                                                #For all wild and NLO foods, ie rows in Harv matrix,
      {if(!is.na(LoBoundHvst.T0.Rpar)) 
         {Harv[Rhv, is.Hetero] <- ifelse(TestTemp >= LoBoundHvst.T[1, is.Hetero], Harv[Rhv, is.Hetero], 0.0000000001)  #If T.index.init >= LoBound of T.index.init, accept +ve Harv, else v low +value (to avoid skipping indexed values later)
         }
      if(!is.na(HiBoundHvst.T0.Rpar)) 
         {Harv[Rhv, is.Hetero] <- ifelse(TestTemp <= HiBoundHvst.T[1, is.Hetero], Harv[Rhv, is.Hetero], 0.0000000001)  #If T.index.init <= HiBound of T.index.init, accept +ve Harv, else v low value
         }
      }                                                                                 
   if(!is.na(LoBoundHvst.T0.Rpar) | !is.na(HiBoundHvst.T0.Rpar))                                             #If either or both T.index bounds are present:-
      {Advise(WithRefTo=NA, ErrText=NA, AdvText=c("LoBound T.index for Harv coeffs in heterotrophic columns:  ", LoBoundHvst.T[1, is.Hetero]), ShowData=NULL, CallSection=NA, Fatal=FALSE, MinAdviceLevel2Show=2)
      Advise(WithRefTo=NA, ErrText=NA, AdvText=c("HiBound T.index for Harv coeffs in heterotrophic columns: ", HiBoundHvst.T[1, is.Hetero]), ShowData=NULL, CallSection=NA, Fatal=FALSE, MinAdviceLevel2Show=2)
      Advise(WithRefTo=NA, ErrText=NA, AdvText=c("Harvest coeffs AFTER zero-ing values outside bounds set from initial T.index =", T.index.init, ": "), ShowData=NULL, CallSection=NA, Fatal=FALSE, MinAdviceLevel2Show=1)  ; if(AL > 0) print(Harv[is.Food, is.Hetero])                                                                            #for checks
      }
Growth <- array(data=Harv * Assim, dim=c(N.Components, N.Components), dimnames=list(paste0("GrowthFrom.", Components), paste0("By.", Components))) #adjust Growth for any new Harv values, element x element
   }
#----------------------------------------------------------------


#4.10 Create Component-type table
#Optional printing of this table of component types (View.comp=1 in Controls.txt) is deferred until all advice and error messages related to input files have been displayed.  See #6
#The component-type table is quite ugly here (if printed now) because quotes are still present around all entries
ComponentType.table <- rbind(is.Wild, is.Auto, is.Hetero, is.Predator, is.TopPredator, is.UnfishedTopPredator, is.Scavenger, is.NLO, is.Harvest, is.Harvester, is.WildHarvest, is.DroppedHarvest, is.UnAssimHarvest, is.UnAssimHarvester, is.Food, is.WildFood, is.Fishery, is.WildOrFishery, is.Caught, is.Landed, is.Discarded, is.ActiveQuotum, is.Consumer, is.Gainer, is.Habitat, is.H2O)  #bind rows to create ComponentType.table ; others are added in later sections below
ComponentType.table <- ifelse(ComponentType.table, TRUE, "x")                                                #change FALSE into x to make the table easier to read
is.UnIDcomp <- !is.Wild & !is.Fishery & !is.NLO & !is.H2O & !is.Habitat                                      #unidentified component(s) are TRUE
if(sum(is.UnIDcomp) > 0)                                                                                     #If the component cannot be identified with one coded into this version of ECOLPS:-
   {Advise(WithRefTo=paste0("Components not assigned a type in \n\t", Params.file), ErrText=c(paste0("Component column  ", Components[which(!is.Wild & !is.Fishery & !is.NLO & !is.H2O & !is.Habitat)]), " in the Params.file could not be assigned a type from 'Wild', 'Fishery', 'NLO', 'H2O' or 'Habitat'. \n\tCheck that all components have appropriate parameter values for the component type intended, or remove the column by calling it NA."), AdvText=paste0("Wild columns have Bmass.init > 0, SelfLoss > 0, and SelfGrow > 0 (Autotroph) or at least 1 Growth (=Harv * Assim) > 0 (Heterotroph). \n\tFisheries columns have Bmass = 0, SelfLoss = 0, at least 1 Growth > 0, and Effort > 0. \n\tNLO column has Bmass.init > 0, SelfLoss > 0, SelfGrow = 0, and all Harv = 0 and all Assim=0, \n\tH2O column has everything = 0 or NA except 1 or more of T.index, L.index, N.mass which are > 0.  \n\tHabitat columns have everything = 0 except HabSize > 0, and at least 1 wild component column must have 1 HabOccup > 0.\n\tData below are for the unidentified component(s):-  ", Components[which(is.UnIDcomp > 0)]), ShowData= Params[ , which(!is.Wild & !is.Fishery & !is.NLO & !is.H2O & !is.Habitat > 0)], CallSection="#4.11", Fatal=TRUE, MinAdviceLevel2Show=1)
   }
#----------------------------------------------------------------
#################################################################End of section #4. RE-ORDERING OF COMPONENTS, CREATION OF PARAMETERS AND INTERNAL LOCATION VECTORS


#5. ENVIRONMENTAL VARIABLES THAT CHANGE WITH TIME, if supplied
#This section creates the Environ.table from the Environment.file, if one has been supplied.  The 5 cols are 
   #'Date.day' (integers or reals > 0 with 365 per year), 
   #'Variable' (entries named exactly as in row.headers of Params.file), 
   #'Value' (integers or reals), and 
   #'Component' (to which the variable value is applied).  
   #'TransferTo', should always be present but is only used when a transfer ('Variable' = Trans) of biomass or nutrient from one component to another is intended.  This, 5th, col is entirely filled with NAs unless 'Variable'=Trans in which case 'TransferTo shows the recipient component, 'Component' shows the donor component, and (0 < Value < 1) holds the proportion of the donor's mass to be transferred at that time.
#NOTE:- Transfer code has hardly been tested!
#Rows can be input in any order, eg. as blocks for each variable type.  They are re-ordered below across all variables by Date.day.  Several possible errors in the input are checked here to prevent non-specific R error messages.
#Components, and suffixes for nutrients, fisheries, habitats, etc. must be spelled exactly the same as the headings or suffixes in Params.file to enable complete linking between them all.
#Rows and cols of the Environment.file are referred to in the code as .Re and .Ce  
   #SUBSECTIONS
      #5.1 Environment.file (if supplied) and required variables
      #5.2 Check values supplied in Environ.file
      #5.3 Create Environ.table


#5.1  Environment.file (if supplied) and required variables
#After reading the Environ.file, the standardised variables (Date.Ce etc) are found and stored.  
Environ.table <- NA                                                                                          #Initialise storage for envir variables. NA is a switch to indicate the non-existence of Environ.file
Advise(WithRefTo=NA, ErrText=NA, AdvText=c("The Environ.file found in Controls.txt is:- ", Environ.file, "\n"), ShowData=NULL, CallSection=NA, Fatal=FALSE, MinAdviceLevel2Show=0 )
if(!is.na(Environ.file))                                                                                     #If a valid Environ.file ref is supplied:-
   {Temp.table <- read.table(file=Environ.file, header=T, as.is=c(T,T,T,T,T))                                #as.is=T prevents each column being read by R as a factor
   ncol.Environ.file <- ncol(Temp.table)                                                                     #test the number of cols in Environ.file
   if(ncol.Environ.file != 5)                                                                                #If N cols not equal to 5:-
      {Advise(WithRefTo=paste0("The number of columns in the Environment.file supplied in Controls.txt:- \n\t", Environ.file, " ."), ErrText="There should be exactly 5 columns.  The columns must be adjusted or, if time-variations of parameters or masses are \n\tnot wanted, replace the Environ.file ref with NA.", AdvText="Columns should be headed \n'Date.day'   (numbers > 0), \n'Variable'   (named as a variable or suffix in rows of the Params.file), \n'Value'      (integers or reals), \n'Component'  (to which the Variable Value is applied), and \n'TransferTo' (filled with NA unless Variable=Trans in which case TransferTo shows the recipient component, Component shows the donor component, \n\tand (0 < Value < 1) holds the proportion of the donor's biomass (or mass) to be transferred at that time. \n\t\n\tBelow: column names read from Environ.file:=\n\t ", ShowData=colnames(Temp.table), CallSection="#5.1a", Fatal=TRUE, MinAdviceLevel2Show=0)
      }
   Date.Ce <- findOneLine(SearchTxt="Date", Headers=colnames(Temp.table), call.section="#5.1", file=Environ.file, domain="column", ZeroOK=FALSE)    #find number of column.header with 'Date' in it
   Var.Ce <- findOneLine(SearchTxt="Var", Headers=colnames(Temp.table), call.section="#5.1", file=Environ.file, domain="column", ZeroOK=FALSE)      #find n of column.header with 'Var' in it, variable names
   Value.Ce <- findOneLine(SearchTxt="Val", Headers=colnames(Temp.table), call.section="#5.1", file=Environ.file, domain="column", ZeroOK=FALSE)    #find n of column.header with 'Val' in it, values
   Comp.Ce <- findOneLine(SearchTxt="Comp", Headers=colnames(Temp.table), call.section="#5.1", file=Environ.file, domain="column", ZeroOK=FALSE)    #find n of column.header with 'Comp' in it, component names
   Trans.Ce <- findOneLine(SearchTxt="Trans", Headers=colnames(Temp.table), call.section="#5.1", file=Environ.file, domain="column", ZeroOK=FALSE)  #find n of col.header with 'Trans', TransferTo comp names

#Init.names stores relevant variable names from those already present in the model, ie. from Params.file.  It is used to check matching of variable names across inputs.
   Init.names <- NULL                                                                                        #Initialise a vector for names of vars in Params; NULL allows incremental lengthening of this vector
   if(N.Nutr > 0) {Init.names <- c(Init.names, Nutrients)}                                                   #Fill Init.names with names of variables initialised by Params; first, nutrient names if any
   if(N.PhysicalVars > 0) {Init.names <- c(Init.names, PhysicalVars)}                                        #Prefixes of physical variable names, if any
   if(N.Fisheries > 0) {Init.names <- c(Init.names, "Effort")}                                               #'Effort' if fisheries are present
   if(N.Habitats > 0) {Init.names <- c(Init.names, "HabSize")}                                               #'HabSize' if habitats are present
   Init.names.withTrans <- c(Init.names, "Trans")                                                            #'Trans' is added to a special set because it is not initialised in Params
#Any other variables in Params that are to be adjusted over time using the Environment.file should be concatenated with Init.names or Init.names.withTrans here
#----------------------------------------------------------------


#5.2 Check values supplied in Environ.file
#All checks are looped around every variable in each row of Environ.file, one row at a time
   #SUBSUBSECTIONS
      #5.2.1 Is Date.day an integer >= 1 and < Inf ?
      #5.2.2 Do Variable names match those already present in the model?
      #5.2.3 Do Component names match those supplied in Params.file ?
      #5.2.4 Are Values valid for 'variable' ?
      #5.2.5 Do 'TransferTo' component names match those supplied in Params.file ?
   for(Re in 1:nrow(Temp.table))                                                                             #For each row of Environ.file as now in Temp.table, do all following checks,

#5.2.1 Is Date.day an integer >= 1 and < Inf ?                                                               #check Date.Ce column (date, measured in days)
      {ERR <- is.DatumOK(Temp.table[Re , Date.Ce], "Date.day", NA.OK=FALSE, CharOK=FALSE, NumbOK=TRUE, LogicalOK=FALSE, MinOK=1, InfinOK=FALSE, nonIntegerOK=FALSE)#check that it is a numeric integer
      if(ERR == TRUE)                                                                                        #If a non-integer is found:-
         {Advise(WithRefTo=paste0("A value of 'Date.day' supplied in the Environ.file:- \n\t ", Environ.file, "  , row ", Re, " ."), ErrText="Date.day may only be a finite numeric integer >= 1.  A non-compliant value has been found, possibly converted to NA when read.  \n\tThe Date.day values must be adjusted", AdvText="Date.day controls the timings of interventions tabulated in the Environ.file. Date.day must be a finite integer >= 1.\n\tBelow: Date.day read from Environ.file: ", ShowData=Temp.table[Re, ], CallSection="#5.2.1", Fatal=TRUE, MinAdviceLevel2Show=0)

         }                  

#5.2.2 Do Variable names match those already present in the model?
      ERR <- is.DatumOK(Temp.table[Re , Var.Ce], "Variable", NA.OK=FALSE, CharOK=TRUE, NumbOK=FALSE, LogicalOK=FALSE, MustBeInSet=Init.names.withTrans)#Check that variable name is in Init.names.withTrans
      if(ERR == TRUE)                                                                                        #If an invalid variable name is found:-
         {Advise(WithRefTo=paste0("A variable name in 'Variable' supplied in the Environ.file:- \n\t", Environ.file, "  , row ", Re, " ."), ErrText=paste0("The Variable name does not match any of those read from ROW headers of the Params.file:- \n\t", Params.file, " \n\tor the special variable, 'Trans', referring to transfers between components.  \n\tA previously unused name has been found, possibly written in a different case.  NAs are not permitted."), AdvText="1. A simple cause of this error is inclusion of an Environ.file in Controls.txt when it is not needed, or use of the wrong file.  \n\tComment out the Environ.file line there with a # at the start of the line, check file names, and/or put NA for the Environ.file name.  \n\t2. A second cause may be the inclusion of a variable that was not initiated with a value in the Parameters.file. Either add a new row\n\t in Params.file with the variable and an initial, dummy value, or remove all rows of that variable in the Environ.file. \n\t3. Otherwise, variable names may not have been spelled exactly the same everywhere they appeared in input files; this prevents matching.  \n\t4. When using an Environment.file, you need a water-component column, eg. 'H2O', in the Params.file for initial values \n\tof T.index, L.index, Nutr.mass and, perhaps, other variables. If there is no water column, these variables may not have been read.  \n\tCorrect this by adding a water component to the Params.file. \n\tBelow: the row of Environ.file with the new (unmatched) variable name, and all variable names read from row.headers of Params.file:- ", ShowData=list(Temp.table[Re, ], row.names(Params)),  CallSection="#5.2.2", Fatal=TRUE, MinAdviceLevel2Show=0)

         }

#5.2.3 Do Component names match those supplied in Params.file ?
      ERR <- is.DatumOK(Temp.table[Re , Comp.Ce], "Component", NA.OK=FALSE, CharOK=TRUE, NumbOK=FALSE, LogicalOK=FALSE, MustBeInSet= Components)
      if(ERR == TRUE)                                                                                        #If an invalid component name is found:-
         {Advise(WithRefTo=paste0("'Component' supplied in the Environ.file:- \n\t", Environ.file, "  , row ", Re, " ."), ErrText=paste0("The Component name does not match any of those read from column headers of the Params.file:-\n\t ", Params.file, ", \n\tAnother, non-matching name has been found, possibly written in a different case.  NAs are not permitted."), AdvText="Component names must be spelled exactly the same wherever they appear in input files so that they may be matched.  \n\tBelow: the row of Environ.file with the unmatched Component name, and Component names read from col.headers of Params.file:- ", ShowData=list(Temp.table[Re, ], Components),  CallSection="#5.2.3", Fatal=TRUE, MinAdviceLevel2Show=0)
         }

#5.2.4 Are Values valid for 'variable' ?
      if(is.element(Temp.table[Re , Var.Ce], c(Nutrients, "Effort", "effort", "Habsize", "habsize")))        #If this 'variable' (as written) should be 0 <= variable < Infinity:-
         {ERR <- is.DatumOK(Temp.table[Re , Value.Ce], "Value", NA.OK=FALSE, CharOK=FALSE, NumbOK=TRUE, LogicalOK=FALSE, MinOK=0, InfinOK=FALSE)
         if(ERR == TRUE)                                                                                     #check that Nutrients, Effort and Habsize are supplied with non-negative, finite numbers as Values
            {Advise(WithRefTo=paste0("'Value' supplied for variable '", Temp.table[Re , Var.Ce], "' in the Environ.file:- \n\t", Environ.file, "  , row ", Re, " ."), ErrText=paste0("The Value provided may only be a non-negative, finite number when Variable = a nutrient, 'Effort' or 'HabSize'. \n\tA non-compliant value has been found. The value must be adjusted.  NAs are not permitted."), AdvText="The Value must be compatible with the Variable name in the same row of Environ.file.  \n\tBelow: the row of Environ.file with the unmatched Component name, and Component names read from col.headers of Params.file:- ", ShowData=list(Temp.table[Re, ], Components),  CallSection="#5.2.4a", Fatal=TRUE, MinAdviceLevel2Show=0)
            }
         }
      if(is.element(Temp.table[Re , Var.Ce], c("T.index", "L.index", "Trans", "trans")))                     #If this 'variable' should be 0 <= variable <= 1:-  
         {ERR <- is.DatumOK(Temp.table[Re , Value.Ce], "Value", NA.OK=FALSE, CharOK=FALSE, NumbOK=TRUE, LogicalOK=FALSE, MinOK=0, MaxOK=1)#Check that it is a proportion
         if(ERR == TRUE)                                                                                    #If a non-proportion is found:-
            {Advise(WithRefTo=paste0("'Value' supplied for variable '", Temp.table[Re , Var.Ce], "' in the Environ.file:- \n\t", Environ.file, "  , row ", Re, " ."), ErrText=paste0("The Value provided may only be a proportion (0 to 1) when Variable = 'L.index', 'T.index' or 'Trans'.   \n\tA non-compliant value has been found. The value must be adjusted.  NAs are not permitted."), AdvText="The Value must be compatible with the Variable name in the same row of Environ.file.  \n\tBelow: the row of Environ.file with the unaccepted value and associated variable:- ", ShowData=Temp.table[Re, ],  CallSection="#5.2.4b", Fatal=TRUE, MinAdviceLevel2Show=0)
            }
         }            
      #Add other if(is.element(...) code in this section for any other variables that change with time and have other valid values

#5.2.5 Do 'TransferTo' component names match those supplied in Params.file ?
#NAs must fill this col except when Variable=Trans:- then the 'TransferTo' component is given
      if(Temp.table[Re , Var.Ce] != "Trans")                                                                 #If the variable name in this row is not 'Trans',
         {ERR <- is.DatumOK(Temp.table[Re, Trans.Ce], "Transfer-to component", NA.OK=TRUE, CharOK=FALSE, NumbOK=FALSE, LogicalOK=FALSE)
         if(ERR == TRUE)                                                                                     #If a non NA is found:-
            {Advise(WithRefTo=paste0("'TransferTo' supplied in the Environ.file:- \n\t", Environ.file, "  , row ", Re, " ."), ErrText=paste0("'TransferTo'  must be NA when 'Variable' is NOT = 'Trans', ie. not a transfer. "), AdvText="TransferTo supplies the component name of the recipient of bio- or nutrient-mass from Component but only if 'Variable' is specified as Trans.  If not, there is \n\tno transfer and TransferTo must be NA.  Below: the row of Environ.file with the illegal TransferTo value:- ", ShowData=Temp.table[Re, ],  CallSection="#5.2.5a", Fatal=TRUE, MinAdviceLevel2Show=0)

            }     
         } else                                                                                              #Else the variable is TransferTo (=Trans)
         {ERR <- is.DatumOK(Temp.table[Re, Trans.Ce], "Transfer-to component", NA.OK=FALSE, CharOK=TRUE, NumbOK=FALSE, LogicalOK=FALSE, MustBeInSet= Components)
         if(ERR == TRUE)                                                                                     #If a name not among Components is found,
            {Advise(WithRefTo=paste0("'TransferTo' supplied in the Environ.file:- \n\t", Environ.file, " row ", Re, " ."), ErrText=paste0("'TransferTo'  must be a component name from column-headers of the Parameters.file:-\n\t", Params.file, " \n\twhen 'Variable' = Trans .  Otherwise, the TransferTo column should only contain NA. "), AdvText="TransferTo gives the recipient of bio- or nutrient-mass from Component but only if 'Variable' is specified as Trans.  \n\tIf not, there is no transfer and TransferTo must be NA.  Below: the row of Environ.file with the illegal TransferTo value:- ", ShowData=Temp.table[Re, ],  CallSection="#5.2.5b", Fatal=TRUE, MinAdviceLevel2Show=0)
            }
         }
      }                                                                                                      #End of row-checking loop
#----------------------------------------------------------------


#5.3 Create Environ.table
#If the supplied Environ.table does not overlap the whole simulation period, Years.sim in Controls.txt, it will be stacked repeatedly to whole years until it does.
#Each original Environ.table, however short is considered to extend so as to complete the year of the last date; this permits stacking by whole years which, in turn, preserves annual cycles if present  
#One Date.day in the Environ.table is always 1/365th of a year regardless of Delta and YearLength set in Controls.txt 
   Environ.table1 <- Temp.table[order(Temp.table[ , Date.Ce]), ]                                             #re-order the input table by the date column and store in Environ.table
   Orig.Environ.table <- Environ.table1                                                                      #create a duplicate of the re-ordered original for stacking as required
   FirstDateInOrigEnvirTable <- Environ.table1[1, Date.Ce]                                                   #the first Date.Day in re-ordered original Environ.table
   LastDateInOrigEnvirTable <- Environ.table1[nrow(Environ.table1) , Date.Ce]                                #the last date in re-ordered original Environ.table
   YearEndOfOrigEnvirTable <- ceiling(LastDateInOrigEnvirTable / 365)                                        #ceiling number of integer years covered by origE.t; ceiling() permits stacking by whole years
   N.RinOrigEnvirTable <- nrow(Orig.Environ.table)                                                           #number of rows in original E.T
   ETstack <- 0                                                                                              #how many times the E.T has been stacked
   if(Years.sim >  YearEndOfOrigEnvirTable)                                                                  #If the simulation period is longer than the original supplied Environ.table:-
      {for(ETstack in 1:ceiling(Years.sim / YearEndOfOrigEnvirTable))                                        #For each stacked E.T until Years.sim is covered,
         {Environ.table1 <- rbind(Environ.table1, Orig.Environ.table)                                        #row-bind on one orig.E.T
         }
      }
   if(ETstack > 0)                                                                                           #If the E.T has been stacked at least once:-
      {StackDateIncrem <- ceiling(LastDateInOrigEnvirTable / 365) * 365                                      #number of days to be added to dates for each stacked Orig.Environ.table
      Increm <- StackDateIncrem                                                                              #the 1st date increment appropriate for the first stacking of the original env.t
      for(Stack.N in 1:ETstack)                                                                              #For each stacking of the original env.t until Years.sim is covered,
         {for(Stack.R in 1:N.RinOrigEnvirTable)                                                              #For each row in the original env.t,
            {Ris <- (N.RinOrigEnvirTable * Stack.N) + Stack.R                                                #Ris is the current row in the new stack
            Environ.table1[Ris, Date.Ce] <- Environ.table1[Ris, Date.Ce] + Increm                            #add the data increment to the date row of this stacked orig.env.t
            }
         Increm <- Increm + StackDateIncrem                                                                  #increase the increm for the next orig.env.t to be stacked
         }
      }   
   Environ.table <- Environ.table1                                                                           #copy.  E.t1 remains constant for use later; E.t is extended or shortened as signalled by MainMenuOptions 8 or 9
   LastDateInEnvirTable <- Years.sim * 365                                                                   #Environ.table[nrow(Environ.table) , Date.Ce] #The last date in the current stacked Environ.table
   YearEndOfEnvirTable <-  ceiling(LastDateInEnvirTable / 365)                                               #the year end of the current stacked E.T
   YearEndOfEnvirTable1 <- YearEndOfEnvirTable                                                               #copy is made; it remains constant and is used under MainMenuOption 8
   N.RinEnvirTable <- nrow(Environ.table)                                                                    #the number of rows in this first stacked Environ.table
   N.RinEnvirTable1 <- N.RinEnvirTable                                                                       #copy is made; it remains constant for any later use
   MonthStarts <- c(1, 32, 60, 91, 121, 152, 182, 213, 244, 274, 305, 335)                                   #day numbers at the start of each month; used to apply understandable seasonal dates in outputs. No leap years allowed! 
   Date.Month <- Date.Day <- vector("numeric", length=nrow(Environ.table))                                   #define vectors for loop below; Date.Day is the name of the time variable in Environ.file
   }                                                                                                         #end of if(!is.na(Environ.file))
#----------------------------------------------------------------
#################################################################End of section 5. ENVIRONMENTAL VARIABLES THAT CHANGE WITH TIME, if supplied


#6. OPTIONAL PRINTING OF INPUT FILES
#Screening of input files, if requested in Controls.txt with View.controls, View.params, View.Envir, View.comps=1) is placed here so as to follow all displayed advice and error messages related to input files
#SUBSECTIONS
   #6.1 View Controls.txt from #1
   #6.2 View Component-Type table from #3
   #6.3 View Parameters table from #4
   #6.4 View Environment table from #5


#6.1 View Controls.txt from #1
if(View.controls == 1)                                                                                       #If View.controls read from Controls.txt file is TRUE [spaces below aim for tidy screen presentation]:-
   {cat("\nECOLPS CONTROLS read from Controls.txt or accepted by default:-\nDelta                              =", Delta, "time unit", "\nDuration of simulation             =", Years.sim, "years", "\nYearLength                         =", YearLength, "delta per year", "\nNumber of projections              =", N.LPproj, "\nProgress reporting, every          =", Progress.rep, " deltas", "\nView program controls on screen?   =", View.controls, "  (0=No, 1=Yes)", "\nView parameters table on screen?   =", View.params, "  (0=No, 1=Yes)", "\nView Environment table on screen?  =", View.Envir, "  (0=No, 1=Yes)", "\nView component table?              =", View.compType, "  (0=No, 1=Yes)", "\nAdvice level                       =", AL, "  (0=no advice, 1=much advice, 2=all advice)", "\nGraphics in colour?                =", Col.plots, "  (0=Black and white, 1=colour)", "\nGraphic key item width             =", KeyItemWidth, " characters", "\nGraphic key characters/line        =", KeyLineLength, "\nParameters file                    =", Params.file, "\nEnvironment file                   =", Environ.file)
   cat("\n\n\n")                                                                                             #line breaks
   }
#----------------------------------------------------------------   


#6.2 View Component-Type table from #3
if(View.compType)                                                                                            #If View.compType read from Controls.txt file is TRUE:-
   {cat("\nCOMPONENT TYPES and their properties calculated from parameter values supplied in \n", Params.file, " :-\n") #write title on screen
   print(ComponentType.table, quote=FALSE)                                                                   #view the component-type table on screen 
   cat("\n\n\n")                                                                                             #line breaks
   }
#----------------------------------------------------------------


#6.3 View original parameters table from #4
if(View.params==1)                                                                                           #If View.params switch read from Controls.txt is True:-
   {cat("\nPARAMETERS as read from the Parameters.file, \n", Params.file, ".  \n#Conversion of NAs to zeroes has been applied. \n#Rows of multi-row parameters have been re-ordered if necessary to match column ordering of components.  \n#Standardised row names have been applied.  \n#Growth matrix is calculated as Harv X Assim:-\n")
   print(Params)                                                                                             #print out the data on screeen
   cat("\n\n\n")                                                                                             #line breaks
   }
#----------------------------------------------------------------


#6.4 View Environment table from #5
if(!is.na(Environ.file))                                                                                     #If an Environ.file was found and View.Envir (set in Controls.txt) is true:-
   {Date.Year <- 0 + floor(Environ.table[ , Date.Ce]/365)                                                    #make a years vector from the date index; years are counted from 0 at start of simulation
   DayInYear <- Environ.table[ , Date.Ce] - (Date.Year - 0) * 365                                            #make a day-in-year vector
   for(ii in 1:length(Date.Month))                                                                           #For each element of these vectors,
      {Date.Month[ii] <- max(which(MonthStarts <= DayInYear[ii]))                                            #make a month-in-year vector at ii
      MthStart <- MonthStarts[Date.Month[ii]]                                                                #store the start of this month
      Date.Day[ii] <- DayInYear[ii] - MthStart + 1                                                           #and make a day-in-month vector at ii
      }
   Date <-  paste(Date.Day, Date.Month, Date.Year, sep="/")                                                  #paste the date components together in DD/MM/YY format
   Vars.Out <- cbind(Environ.table, Date)                                                                    #bind with Environ.table as a temporary table for viewing.
   row.names(Vars.Out) <- as.numeric(row.names(Vars.Out)) + 1                                                #row names; +1 corrects for header row
   colnames(Vars.Out)[(ncol(Environ.table) + 1)] <- "DD/MM/YY"                                               #label the new col
   if(View.Envir==1)                                                                                         #If View.Envir read from Controls.txt file is TRUE:-
      {cat("\nVARIABLES OVER TIME, sorted by Date.day, and read from the Environment.file, \n", Environ.file, ":-  \nIf necessary, supplied rows will be stacked by integer years to cover the simulation period, Years.sim =", Years.sim, ".  \n1st col = row number before sorting.  \n\n")        
      print(Vars.Out)                                                                                        #view the Environ.table and calendar dates as requested
      cat("\n\n\n")                                                                                          #line spacers
      }
   }
#----------------------------------------------------------------
#################################################################End of section 6. OPTIONAL PRINTING OF INPUT FILES


#7. INITIALISE DATA STRUCTURES AND LOCATION VECTORS FOR SIMULATIONS
#Data from Params (stored in #3, re-ordered with standard row and column orderings) are used for initialisation at t=0 of data structures needed for simulations
   #SUBSECTIONS
      #7.1 Variables needed for data structures
      #7.2 Objective function and associated variables
      #7.3 Time series arrays for storing simulated results and varying parameters
      #7.4 Left-side matrices of constraints
      #7.5 Constraint-related time series
      #7.6 Create non-time-series structures for simulations
      #7.7 Other variables and constants needed for projection steps


#7.1 Variables needed for data structures
TS.length <- N.LPproj + 1                                                                                    #length of simulated time series to be created in #7.3
SimTime <- seq(0, N.LPproj)                                                                                  #vector of times of simulation points in Delta units
Years <- round(SimTime/YearLength, 3)                                                                        #vector of dates in years of simulation points rounded to 3 decimal places
names(SimTime) <- paste(as.character(Years), "y", sep="")                                                    #character vector for naming the elements of SimTime in terms of years 'y'
Last.Cts <- TS.length                                                                                        #initial values are put in the last column of time-series (Last.Cts); moved to col.1 at Main menu Option8 below
#----------------------------------------------------------------


#7.2 Objective function and associated variables
#The objective-function is a row vector with autos and consumers ordered as in columns of the re-ordered Params file.  Autos each get 1 col; consumers get 1 column for each of their foods, also so ordered.
#Because the obj fun contains food-consumer pairs, the ordering of obj fun elements cannot match that of cols in Params, special indexing is required
#The orderings of ObjFun elements and cols in the constraint matrix (All.C) must be the same, as required for the simplex() function used for LP

# .Loc vectors have components named where they occur, otherwise NAs are stored. So, a search for non-NAs in the appropriate vector locates the component types in the objective function.
ObjFun.names <- c(Autotrophs, FoodConsumer.pairs)
N.LPvars <- length(ObjFun.names)                                                                             #number of variables for linear programming
ObjFun <- array(data=rep(1, N.LPvars), dim=c(1, N.LPvars), dimnames=list("ObjFun.coeffs", ObjFun.names))     #all coeffs in the obj.fun =1 initially to give equal weighting of GP from each p.p and food-cons pair
ObjFun.Cons.Loc <- c(rep(NA, N.Autos), AllPossHarvesters.array[which(Growth > 0, arr.ind=TRUE)])             #consumers found in food x consumer pairs
ObjFun.Autos.Loc <- c(Autotrophs, rep(NA, N.FoodConsumer.pairs))                                             #autotrophs
ObjFun.Foods.Loc <- c(rep(NA, N.Autos), substr(FoodConsumer.pairs, 1, regexpr(">", FoodConsumer.pairs) - 1)) #foods. Autos have NA because don't eat food
ObjFun.GProducers.Loc <- c(Autotrophs, ObjFun.Cons.Loc[(N.Autos + 1):N.LPvars])                              #GProducers = all cols with auto or consumer
ObjFun.Heteros.Loc <- ifelse(ObjFun.Cons.Loc %in% Heterotrophs, ObjFun.Cons.Loc, NA)                         #heterotrophs
ObjFun.Grazers.Loc <- ifelse(ObjFun.Cons.Loc %in% Grazers, ObjFun.Cons.Loc, NA)                              #grazers
ObjFun.Scavengers.Loc <- ifelse(ObjFun.Cons.Loc %in% Scavengers, ObjFun.Cons.Loc, NA)                        #scavengers
ObjFun.Predators.Loc <- ifelse(ObjFun.Cons.Loc %in% Predators, ObjFun.Cons.Loc, NA)                          #predators
ObjFun.Fisheries.Loc <- ifelse(ObjFun.Cons.Loc %in% Fisheries, ObjFun.Cons.Loc, NA)                          #fisheries, if any
ObjFun.Quota.Loc <- ifelse(ObjFun.names %in% QuotumFishery.pairs, ObjFun.names, NA)                          #quota, if any; Note: some QF.pairs may not be landed against so have no effect on ObjFun, until they are landed

# .Obj vectors store the element positions in the objective function of members of the named component or productive pair-type
Consumer.Obj <- which(!is.na(ObjFun.Cons.Loc))                                                               #element numbers of consumers
Auto.Obj <- which(!is.na(ObjFun.Autos.Loc))                                                                  #element numbers of autos
ObjFun.Foods.Obj <- which(!is.na(ObjFun.Foods.Loc))                                                          #element numbers of foods
GProducer.Obj <- which(!is.na(ObjFun.GProducers.Loc))                                                        #element numbers of GProducers
Hetero.Obj <- which(!is.na(ObjFun.Heteros.Loc))                                                              #element numbers of heteros
Grazers.Obj <- which(!is.na(ObjFun.Grazers.Loc))                                                             #element numbers of grazers
Scavengers.Obj <- which(!is.na(ObjFun.Scavengers.Loc))                                                       #scavengers
Predators.Obj <- which(!is.na(ObjFun.Predators.Loc))                                                         #element numbers of predators
Fishery.Obj <- which(!is.na(ObjFun.Fisheries.Loc))                                                           #element numbers of fisheries
Quota.Obj <- which(!is.na(ObjFun.Quota.Loc))                                                                 #element numbers of fisheries quota
#----------------------------------------------------------------


#7.3 Time series arrays for storing simulated results and varying parameters
#R note: array(..., dim=c(1, ) ...) formats as row vectors and provides row and col names even if there is only one row; other R structures (vector, matrix) are less flexible for 1 or 2+ rows or columns
#Initial values read from Params.file are stored in last col of time-series (Last.Cts); they are moved to col.1 in MainMenu option 8 below => 1st block of LP steps processed the same as any later blocks requested in M.Menu 8 or 9
#Note: Gainers, see #4.3, include components gaining by GP (all wilds), by dropping of organic biomass (NLO), and by landing fish (fisheries)
# pGains = partial Gains are from 1 food or source only; some of these may be calculated by projection, indicated by suffix '.Byproj'. All are calculated by LP if possible (if Assim > 0) indicated by '.ByLP'
#pHarvs = harvests taken by each HarvestHarvester.pair; zero for autos and NLOs; if pGain is by projection, so is pHarv
#NOTE: gains and harvests etc estimated by projection are not discussed in Cotter Parts I or II but are left in the code here for possible use
   #SUBSUBSECTIONS
      #7.3.1 Gross-production and harvest time series, and locators for rows in them
      #7.3.2 Grouped structures for outputs
      #7.3.3 Projected time series
      #7.3.4 Organic-biomass time series for wild and NLO components
      #7.3.5 Nutrient time series
      #7.3.6 Fishery time series
      #7.3.7 Habitat usage
      #7.3.8 Physical time series for Temperature and Light
      #7.3.9 Time series of varying parameters

#7.3.1 Gross-production and harvest time series, and locators for rows in them
   #SUBSUBSUBSECTIONS
      #7.3.1.1 Create vectors and matrices
      #7.3.1.2 .Loc vectors for rows in pGains, pHarvs etc
      #7.3.1.3 .RpG vectors to store the row numbers for .Loc vectors in pGains etc
      
#7.3.1.1 Create vectors and matrices      
AllPossGainers <- c(Autotrophs, HarvestHarvester.pairs)                                                                          #possible gainers including UAH pairs but not NLO yet
if(N.NLOs > 0) AllPossGainers <- c(AllPossGainers, NLOs)                                                                         #concatenate any NLO row(s); gains are dropped particulate self losses and harvests
N.PossGainers <- length(AllPossGainers)                                                                                          #the number of them
pGains <- array(data=0, dim=c(N.PossGainers, TS.length), dimnames=list(paste0("pGainFrom.", AllPossGainers), names(SimTime)))    #Auto rows, then pGains for each Harvest-harvester pair and any NLO
pHarvs <- array(data=0, dim=c(N.PossGainers, TS.length), dimnames = list(paste0("pHarvOf.", AllPossGainers), names(SimTime)))    #Auto rows, then pHarvs taken by each Harvest-Harvester.pair and any NLO; some rows redundant
Dropped.pHarvs <- array(data=0, dim=c(N.PossGainers, TS.length), dimnames = list(paste0("DroppedHarvOf.", AllPossGainers), names(SimTime))) #Auto rows, then droppedHarvs for ditto
pGainsToDropped.factor <- array(data=0, dim=c(1, N.PossGainers), dimnames=list("pGainsToNLO.factor", AllPossGainers))            #conv.factor for pGains -> dropped waste to NLO (for Assim > 0)
pGainsToDropped.factor[1, (N.Autos + which(HarvestHarvester.pairs %in% FoodConsumer.pairs))] <- as.vector(InverseAssim[which(InverseAssim < Inf, arr.ind=TRUE)] - 1)  #conv.factor is (InverseAssim - 1) except NLOs which=0

#7.3.1.2  .Loc vectors for rows in pGains, pHarvs etc
#pHarvs and others use pGains.Loc locators.  They work like ObjFun.Autos.Loc. Elements are either the name of a component or trophic pair, or NA
pGainsByLP.Loc <- ifelse(AllPossGainers %in% ObjFun.names, AllPossGainers, NA)                                                   #pGains loc for all autos and food>consumer pairs estim. by LP
pGainsByProj.Loc <- ifelse(AllPossGainers %in% UnAssimHarv.pairs, AllPossGainers, NA)                                            #pGains loc for all autos and food>consumer pairs estim. by projection; all are expected to be 0
pGains.Loc <- ifelse(is.na(pGainsByLP.Loc), pGainsByProj.Loc, pGainsByLP.Loc)                                                    #pGains loc for all autos and food>consumer pairs estim. either way
pGains.Autos.Loc <- ifelse(AllPossGainers %in% Autotrophs, Autotrophs, NA)                                                       #pGains Loc for autotrophs as photosynthesists
pGains.Harvesters.Loc <- ifelse(AllPossGainers %in% HarvestHarvester.pairs, substr(AllPossGainers, regexpr(">", AllPossGainers) + 1, 100), NA) #pGains Loc for Harvesters including when harvesting UAHs
pGains.NLO.Loc <- rep(NA, N.Gainers)                                                                                             #initial vector
if(N.NLOs > 0)                                                                                                                   #If NLO(s) is in the model:-
   {pGains.NLO.Loc <- ifelse(AllPossGainers %in% NLOs, AllPossGainers, NA)                                                       #pGains Loc for NLO(s)
   pGains.Loc <- c(pGains.Loc[1:N.PossGainers], NLOs)                                                                            #concatenate NLOs with previous pGains.Loc
   }
pGains.Consumers.Loc <- ifelse(AllPossGainers %in% FoodConsumer.pairs, substr(AllPossGainers, regexpr( ">", AllPossGainers) + 1, 100), NA)  #pGains loc for all consumers
pGains.Foods.Loc  <- ifelse(AllPossGainers %in% FoodConsumer.pairs, substr(AllPossGainers, 1, regexpr( ">", AllPossGainers) - 1), NA)       #pGains loc for all foods
pGains.Scavengers.Loc <- ifelse(AllPossGainers %in% NLOScavenger.pairs, substr(AllPossGainers, regexpr( ">", AllPossGainers) + 1, 100), NA) #pGains loc for all scavengers
pGains.Fisheries.Loc <- ifelse(AllPossGainers %in% FoodFishery.pairs, substr(AllPossGainers, regexpr( ">", AllPossGainers) + 1, 100), NA)   #pGains loc for all fisheries
pGains.UnAssimHarvesters.Loc <- c(rep(NA, N.Autos), ifelse(HarvestHarvester.pairs %in% UnAssimHarv.pairs, substr(HarvestHarvester.pairs, regexpr(">", HarvestHarvester.pairs) + 1, 100), NA), rep(NA, N.NLOs))  #for UAHarvesters
pGains.UnAssimHarvests.Loc <- c(rep(NA, N.Autos), ifelse(HarvestHarvester.pairs %in% UnAssimHarv.pairs, substr(HarvestHarvester.pairs, 1, regexpr(">", HarvestHarvester.pairs) - 1), NA), rep(NA, N.NLOs))  #for UAHarvesters
pGains.UnAssimHarv.pairs.Loc <- c(rep(NA, N.Autos), ifelse(HarvestHarvester.pairs %in% UnAssimHarv.pairs, HarvestHarvester.pairs, NA), rep(NA, N.NLOs))  #for UAHarvestHarvester.pairs        

#'Harvester' is a component that kills other components;  harvesters may or may not assimilate harvests
pHarvestersByLP.Loc <- pGains.Consumers.Loc                                                                                      #harvesters estimated by LP are all, and the only, consumers
pHarvestersByProj.Loc <-  pGains.UnAssimHarvesters.Loc                                                                           #for harvesters of HH pairs  by projection; all are UAHarvests
pHarvesters.Loc <- ifelse(is.na(pHarvestersByLP.Loc), pHarvestersByProj.Loc, pHarvestersByLP.Loc)                                #for harvesters of all HarvestHarvester pairs; not autos or NLOs
pGains.Comps.Loc <- c(pGains.Autos.Loc, pHarvesters.Loc, pGains.NLO.Loc)                                                         #pGains.Loc shows pairs but pGains.Comps.Loc shows only the gaining component
pGains.Comps.Loc <- pGains.Comps.Loc[which(!is.na(pGains.Comps.Loc))]                                                            #calculate it from initial values put in pG.Comps.Loc

#'Harvest' is a component that is killed (NLO is eaten) by others; may or may not be assimilated by harvester; if not, it is UAH
pHarvestsByLP.Loc <- c(ifelse(c(Autotrophs, HarvestHarvester.pairs) %in% ObjFun.names, c(rep(NA, N.Autos), substr(HarvestHarvester.pairs, 1, regexpr(">", HarvestHarvester.pairs) - 1)), NA), rep(NA, N.NLOs)) #for harvs of HHpairs
pHarvestsByProj.Loc <-  ifelse(is.na(pHarvestsByLP.Loc), c(rep(NA, N.Autos), substr(HarvestHarvester.pairs, 1, regexpr(">", HarvestHarvester.pairs) - 1)), rep(NA, N.NLOs))  #for harvests of HHpairs est.by projection
pHarvests.Loc <- ifelse(is.na(pHarvestsByLP.Loc), pHarvestsByProj.Loc, pHarvestsByLP.Loc)                                        #for harvests of all HH pairs
pHarvests.NLO.Loc <- ifelse(pHarvests.Loc %in% NLOs, pHarvests.Loc, NA)                                                          #for harvests of NLO by scavengers     

#7.3.1.3 .RpG vectors to store the row numbers for .Loc vectors in pGains etc
pGainsByLP.RpG <- which(!is.na(pGainsByLP.Loc))                                                                                  #row numbers in pGains etc estimated by LP   
pGainsByProj.RpG <- which(!is.na(pGainsByProj.Loc))                                                                              #rows in pGains etc estimated by projection
pHarvestersByLP.RpG <- which(!is.na(pHarvestersByLP.Loc))                                                                        #rows in pGains etc for the harvesters in food-consumer pairs by LP
pHarvestersByProj.RpG <- which(!is.na(pHarvestersByProj.Loc))                                                                    #rows in pGains etc for harvesters in HHpairs by projection
pHarvesters.RpG <- which(!is.na(pHarvesters.Loc))                                                                                #rows in pGains etc holding any harvesters
pHarvestsByLP.RpG <- which(!is.na(pHarvestsByLP.Loc))                                                                            #rows in pGains etc for harvests in food-consumer pair
pHarvestsByProj.RpG <- which(!is.na(pHarvestsByProj.Loc))                                                                        #rows in pGains etc for harvests in HHpairs by projection
pHarvests.RpG <- which(!is.na(pHarvests.Loc))                                                                                    #rows in pGains etc holding harvests
pHarvests.NLO.RpG <- which(!is.na(pHarvests.NLO.Loc))                                                                            #rows in pGains etc holding NLO
Autos.RpG <- which(!is.na(pGains.Autos.Loc))                                                                                     #rows in pGains etc for autos     
Consumers.RpG <- which(!is.na(pGains.Consumers.Loc))                                                                             #rows in pGains etc for consumers with Assim > 0
Scavengers.RpG <- which(!is.na(pGains.Scavengers.Loc))                                                                           #rows in pGains etc for scavengers assimilating NLO
UnAssimHarv.pairs.RpG <- which(!is.na(pGains.UnAssimHarv.pairs.Loc))                                                             #rows in pGains etc for unassimilated harvests estimated by projection
NLO.RpG <- which(!is.na(pGains.NLO.Loc))                                                                                         #rows in pGains etc for NLO
Landings.RpG <- which(pGains.Consumers.Loc %in% Fisheries)                                                                       #rows in pGains holding landings of individual fisheries 
Fisheries.RpG <- which(pHarvesters.Loc %in% Fisheries)                                                                           #rows in pGains holding catches=harvests by individual fisheries (some may not be landed)
Components.RpG <- c(Autos.RpG, Consumers.RpG, NLO.RpG)                                                                           #rows in pGains for all components

#7.3.2 Grouped structures for outputs
#These have 1 component per row and so can be indexed with is.Auto, is.NLO, etc. as for Bmass, Components, etc
Gains <- array(data=0, dim=c(N.Gainers, TS.length), dimnames=list(paste0("GainsTo.", Gainers), names(SimTime)))                              #totalled pGains for each gainer
HarvestsFrom <- array(data=0, dim=c(N.Gainers, TS.length), dimnames=list(paste0("HarvestsFrom.", Gainers), names(SimTime)))                  #total pHarvests taken from each gainer, including any projected
HarvestsBy <- array(data=0, dim=c(N.Gainers, TS.length), dimnames=list(paste0("HarvestsBy.", Gainers), names(SimTime)))                      #total pHarvests taken by each gainer, including any projected
Grouped.Gain.names <- c("Autos", "Predators", "Scavengers", "Heteros", "Wilds", "Landings", "Consumers", "NLO", "TotalGP&Landings")          #names for groups of Gains
N.GainGroups <- length(Grouped.Gain.names)
Grouped.Gains <- array(data=0, dim=c(N.GainGroups, TS.length), dimnames = list(paste0("GainsBy.", Grouped.Gain.names), names(SimTime)))      #total GP for each group of gainers
GHF.names <- Grouped.Gain.names
GHF.names[9] <- "Wilds&NLOs"
Grouped.HarvestsFrom <- array(data=0, dim=c(N.GainGroups, TS.length), dimnames = list(paste0("HarvestsFrom.", GHF.names), names(SimTime)))   #total harvests from each group (some zero)
GHB.names <- Grouped.Gain.names
GHB.names[6] <- "Fisheries"
GHB.names[9] <- "Consumers"
Grouped.HarvestsBy <- array(data=0, dim=c(N.GainGroups, TS.length), dimnames = list(paste0("HarvestsBy.", GHB.names), names(SimTime)))       #total harvests by consumers again

#7.3.3 Projected time series
#These arrays hold only Taylor-series projections
proj.SelfLosses<- array(data=0, dim=c(N.Components, TS.length), dimnames=list(paste0("projSLof.", Components), names(SimTime)))              #self losses for components by projection; only WildOrNLOs have SLs
TotalLossesByProj <- array(data=0, dim=c(N.Components, TS.length), dimnames=list(paste0("projTotLossOf.", Components), names(SimTime)))      #total (self + Harvest) losses for components; only WildOrNLOs have losses
D1.BmassByProj <- array(data=0, dim=c(N.Components, TS.length), dimnames = list(paste("projD1.Bmass", Components, sep="."), names(SimTime))) #D1.biomass time-series projected from GLV 
D2.BmassByProj <- array(data=0, dim=c(N.Components, TS.length), dimnames = list(paste("projD2.Bmass", Components, sep="."), names(SimTime))) #D2.biomass time-series projected from GLV
D3.BmassByProj <- array(data=0, dim=c(N.Components, TS.length), dimnames = list(paste("projD3.Bmass", Components, sep="."), names(SimTime))) #D3.biomass time-series projected from GLV

#7.3.4 Organic-biomass time series for wild and NLO components
#Initial values are put in the last col of the time-series arrays; they will be moved to the 1st column at Main menu Option8 below
Bmass <- array(data=NA, dim=c(N.Components, TS.length), dimnames = list(paste("Bmass", Components, sep="."), names(SimTime)))                #biomass for each component over time; some are always zero (for simpler indexing)
Bmass[ , Last.Cts] <- Bmass.init[1, ]                                                                                                        #initial biomasses are stored in the last col of the TS
#Biomass series grouped for outputs
Grouped.Bmass.names <- paste0("Bmass.", Grouped.Gain.names)                                                                                  #most grouped bmass names are the same as grouped gain names
Grouped.Bmass.names[6] <- "Bmass.AllCatch"                                                                                                   #change Landings to Caught in grouped biomass matrix
Grouped.Bmass.names[9] <- "Bmass.Wilds+Catch+NLO"                                                                                            #change total name
TrophicGroups.Bmass <- array(data=NA, dim=c(N.GainGroups, TS.length), dimnames = list(Grouped.Bmass.names, names(SimTime)))                  #grouped Bmass matrix with 1 row for each group
TrophicGroups.Bmass[1, Last.Cts] <- sum(Bmass.init[is.Auto])                                                                                 #total initial biomass of autotrophs put in the 1st row of the last col
TrophicGroups.Bmass[2, Last.Cts] <- sum(Bmass.init[is.Predator])                                                                             #of predators, put in the second row
if(N.Scavengers > 0) TrophicGroups.Bmass[3, Last.Cts] <- sum(Bmass.init[is.Scavenger]) else TrophicGroups.Bmass[3, Last.Cts] <- NA           #of scavengers in 3rd row
TrophicGroups.Bmass[4, Last.Cts] <- sum(Bmass.init[is.Hetero])                                                                               #of heterotrophs in 4th row
TrophicGroups.Bmass[5, Last.Cts] <- sum(Bmass.init[is.Wild])                                                                                 #of Wilds in 5th row
if(N.Fisheries > 0) TrophicGroups.Bmass[6, Last.Cts] <- 0                                                                                    #of Catch components in 6th row, initially always 0 because no fishing done yet
TrophicGroups.Bmass[7, Last.Cts] <- sum(Bmass.init[is.Consumer])                                                                             #of consumers in 7th row
if(N.NLOs > 0) TrophicGroups.Bmass[8, Last.Cts] <- sum(Bmass.init[is.NLO])                                                                   #of NLO
TrophicGroups.Bmass[9, Last.Cts] <- sum(TrophicGroups.Bmass[c(5, 8), Last.Cts], na.rm=TRUE)                                                  #total ecosystem organic mass=Wilds + NLOs ; Caughts not included because removed

#7.3.5 Nutrient time series
Nmasses <- array(data=NA, dim=c(6, TS.length), dimnames = list(c("Nmass.H2O", "Nmass.Wilds", "Nmass.NLO", "Nmass.TotSystem", "Nmass.Landed", "Nmass.CumLanded"), names(SimTime))) #default Nmass time-series for H2O, NLO and Wilds
NconcInNLO <- array(data=NA, dim=c(1, TS.length), dimnames = list("Nconc.NLO.", names(SimTime))) #default Nconc in NLO.  Only NLO conc is variable so it needs a time series array; dummy if nutrients are not in the model
if(N.Nutr > 0)
   {NmassInH2O.names <- paste(row.names(Nmasses)[1], Nutrients, sep=".")                                     #nutr.mass in water names  ; new row names are needed to identify the given nutrient(s)
   NmassInWilds.names <- paste(row.names(Nmasses)[2], Nutrients, sep=".")                                    #total nutr.mass in all wild components 
   NmassInNLO.names <- paste(row.names(Nmasses)[3], Nutrients, sep=".")                                      #nutr.mass in NLO names
   NmassInSys.names <- paste(row.names(Nmasses)[4], Nutrients, sep=".")                                      #total nutr.mass in the whole ecosystem
   NmassLanded.names <- paste(row.names(Nmasses)[5], Nutrients, sep=".")                                     #total nutr.mass landed, ie. removed from the ecosystem
   NmassCumLanded.names <- paste(row.names(Nmasses)[6], Nutrients, sep=".")                                  #total nutr.mass landed, ie. cumulative total removed from the ecosystem
   Nmasses <- array(data=0, dim=c(N.Nutr * 6, TS.length), dimnames = list(c(NmassInH2O.names, NmassInWilds.names, NmassInNLO.names, NmassInSys.names, NmassLanded.names, NmassCumLanded.names), names(SimTime))) #Nmass array
   NinH2O.R <- 1:N.Nutr                                                                                      #rows in Nmasses for water, assuming 1 water component
   NinWilds.R <- (N.Nutr + 1) : (2 * N.Nutr)                                                                 #rows in Nmasses for all wilds
   NinNLO.R <- (2 * N.Nutr + 1) : (3 * N.Nutr)                                                               #rows in Nmasses for any NLO present
   NinSys.R <- (3 * N.Nutr + 1) : (4 * N.Nutr)                                                               #rows in Nmasses for total Nmass in ecosystem
   NinLandings.R <- (4 * N.Nutr + 1) : (5 * N.Nutr)                                                          #rows in Nmasses for Nmass landed by any fisheries present
   NinCumLandings.R <- (5 * N.Nutr + 1) : (6 * N.Nutr)                                                       #rows in Nmasses for cumulative Nmass landed by any fisheries present
   Nmasses[NinH2O.R, Last.Cts] <- Nmass.init[ , is.H2O]                                                      #transfer nutr masses from Nmass.init array to last col of NmassInH2O time series; moved to 1st col at Main menu 8
   Nmasses[NinWilds.R, Last.Cts] <- Nconc[ , is.Wild] %*% Bmass[is.Wild , Last.Cts]                          #Nmasses held in living components by matrix multiplication
   Nmasses[NinSys.R, Last.Cts] <- Nmasses[NinH2O.R, Last.Cts] + Nmasses[NinWilds.R, Last.Cts]                #calculate system Nmass in case no NLO is in the model
   #initial Nmasses in fishery landings are inserted when the first LP step has been carried out; see #8.8.3.7 code:- if(N.Fisheries > 0) etc
   Nmasses[NinCumLandings.R, Last.Cts] <- 0                                                                  #zero the cumulative landings for first step
   if(N.NLOs == 1)                                                                                           #If NLO is in the model:-
      {Nmasses[NinNLO.R, Last.Cts] <- Nmass.init[ , is.NLO]                                                  #transfer NLO nutr masses from Nmass.init array to last col; moved to 1st col at Main menu 8 below
      NconcInNLO <- array(data=0, dim=c(max(N.Nutr, 1), TS.length), dimnames = list(paste0("Nconc.NLO.", Nutrients), names(SimTime)))  #Nconc in NLO.  Only NLO concs are variable; dummy if nutrients are not in the model
      NconcInNLO[ , Last.Cts] <- Nconc[ , is.NLO]                                                            #initial concs for NLO were calculated in #4.2; the columns of Nconc match the rows of Bmass matrices
      Nmasses[NinSys.R, Last.Cts] <- Nmasses[NinSys.R, Last.Cts] + Nmasses[NinNLO.R, Last.Cts]               #add Nmasses in NLO to total system Nmass
      }
   }   

#7.3.6 Fisheries' time series
F.effort <- array(data=NA, dim=c(1, TS.length), dimnames = list("FishingEffort", names(SimTime)))                                #dummy fishing effort T.S.
QuotaLeft <- array(data=NA, dim=c(1, TS.length), dimnames=list("QuotaLeft", names(SimTime)))                                     #dummy quota remaining
if(N.Fisheries > 0)                                                                                                              #If fisheries are in the model,
   {F.effort <- array(data=NA, dim=c(N.Fisheries, TS.length), dimnames = list(paste0("Effort.", Fisheries), names(SimTime)))     #fishing effort T.S.
   F.effort[ , Last.Cts] <- Effort.init[1, is.Fishery]                                                                           #store f.effort in last col of time-series array; moved to 1st at Main menu Option8 below
   if(N.QuotumFishery.pairs > 0)                                                                                                 #If quota are in the model:-
      {QuotaLeft <- array(data=NA, dim=c(N.QuotumFishery.pairs, TS.length), dimnames=list(paste0("QuotaLeft.", QuotumFishery.pairs), names(SimTime)))#Quota remaining
      QuotaLeft[ , Last.Cts] <- t(Quota.init)[t(is.ActiveQuotum)]                                                                #remove infinity values and put quota into fishery order; t() allows for R working by cols
      }
   }                                                          

#7.3.7 Habitat usage
HabLeft <- array(data=NA, dim=c(1, TS.length), dimnames=list("HabSizeLeft", names(SimTime)))                                     #dummy for habitat left to fill
if(N.Habitats > 0)                                                                                                               #If essential habitats are in the model,
   {HabLeft1 <- array(data=HabSize.init[is.Habitat]  - HabOccup %*% Bmass.init[1, ] , dim=c(N.Habitats, 1), dimnames=list(paste0("SizeOf.", Habitats), "SizeUnits")) #empty habitat size - initial biomass using habitats
   if(any(HabLeft1[ , 1] < 0))                                                                                                   #If any habitat is over-filled (HabLeft1 < 0) at initialisation :-
      {Advise(WithRefTo=paste0("Initial biomasses of components requiring essential habitats as supplied in Parameters file:- \n\t", Params.file, " ."), ErrText="One or more essential habitats contain more initial biomass of components requiring those habitats than the habitat size allows.  \n\tEither increase HabSize.init, decrease occupation coefficients, or decrease initial biomasses occupying the habitat.", AdvText="Living components may only occupy one essential habitat at a time.  Their initial biomasses must be small enough to fit into \n\ttheir required habitat.  The size of habitat required by each component is its biomass x its occupation coefficient. \n\tBelow: Initial biomasses, occupancy coefficients, habitat sizes, and habitat remaining, as read from the file:-", ShowData=list(Bmass.init, HabOccup, HabSize.init, HabLeft1), CallSection="#7.3.7", Fatal=TRUE, MinAdviceLevel2Show=0)
      }
   HabLeft <- array(data=NA, dim=c(N.Habitats, TS.length), dimnames=list(paste0("HabSizeLeft.", Habitats), names(SimTime)))      #array for habitat-left to fill
   HabLeft[ , Last.Cts] <- HabLeft1                                                                                              #insert HabLeft values in last col; moved to 1st at Main menu Option8 below
   }  

#7.3.8 Physical time series for Temperature and Light
T.index <- array(data=NA, dim=c(1, TS.length), dimnames = list("Temperature.index", names(SimTime)))                             #define time series array for T.index varying 0 to 1; serves as dummy if no T.index supplied
T.index[1, Last.Cts] <- 1                                                                                                        #insert default=1 Temperature index in last column
if(!is.na(T.index0.Rpar))                                                                                                        #If a T.index row was found in Params:-  Note: T.index0.Rpar always exists, Rpar.T.index only if T.i
   {if(length(Rpar.T.index) == 1 )                                                                                               #and If there is exactly one Temperature row:-
      {T.index[1, Last.Cts] <- T.index.init                                                                                      #store T.index.init supplied in H2O col of Params.file into last column of time series
      } 
   }     
L.index <- array(data=NA, dim=c(1, TS.length), dimnames = list("Light.index", names(SimTime)))                                   #define the time series array for light; serves as dummy if no T.index supplied
L.index[1, Last.Cts] <- 1                                                                                                        #insert default 1 in last column; moved to 1st at Main menu Option8 below
if(!is.na(L.index0.Rpar))                                                                                                        #If a L.index row has been found in Params:-
   {if(length(Rpar.L.index) == 1)                                                                                                #and If there is exactly one L.index row in Params, necessarily in is.H2O:-
      {L.index[1, Last.Cts] <- L.index.init                                                                                      #store initial L.index for water into last element; moved to 1st at Main menu Option8 below
      }
   } 
   
#7.3.9 Time series of varying parameters
#7.3.10.1 Varying SelfGrow coeffs of autotrophs; only the adjusted ones are stored
#SelfGrow coeffs may have been adjusted for L.index.init and/or T.index.init in #4.8
SG.LTadj <- array(data=0, dim=c(1, TS.length), dimnames = list("SGadj4LT.index", names(SimTime)))                                #default time-series array for adjusted SelfGrow with light and/or temperature indexes
N.SGadj <- 0                                                                                                                     #default number of adjusted SelfGrow values
is.SGadj <- is.LoBoundSG.T | is.HiBoundSG.T | is.alphaSG.I                                                                       #TRUE for components having SG adjusted for alphaSG.I, LoBoundSG.T or HiBoundSG.T
if(sum(is.SGadj) > 0)                                                                                                            #If any SelfGrow is adjusted for alphaSG.I, LoBoundSG.T or HiBoundSG.T,
   {N.SGadj <- sum(is.SGadj)                                                                                                     #adjust the number of adjusted SelfGrow values
   SG.LTadj <- array(data=0, dim=c(N.SGadj, TS.length), dimnames = list(paste0("SGadjust4LT.index.", Components[is.SGadj]), names(SimTime))) #expand SG.LTadj, 1 TS per row, and name rows
   SG.LTadj[ , Last.Cts] <- SelfGrow[1, is.SGadj]                                                                                #put initial, previously adjusted values of SG into last col of Time Series
   }

#7.3.9.2 Varying Harvest coeffs of heterotrophs; only the adjusted ones are stored
#If any cols of Harv coeffs appear to be missing or unadjusted, check that they have alphaHvst.T > 0 for that column; if not, they are not stored as a varying coeff
#Harvest coeffs may have been adjusted for T.index.init in #4.9
Hvst.Tadj <- array(data=0, dim=c(1, TS.length), dimnames = list("HarvAdj4T", names(SimTime)))                                    #default time-series array for Harv coeffs that might be adjusted with T.index, including UAHs
N.HvstColsAdj <- 0                                                                                                               #default number of adjusted Harv columns
is.HvstColsAdj <- is.LoBoundHvst.T | is.HiBoundHvst.T | is.alphaHvst.T                                                           #TRUE for components having Harv adjusted for alphaHvst.T, LoBoundSG.T or HiBoundSG.T
is.HvstAdjust4T.matrix <- array(data=FALSE, dim=c(N.Components, N.Components), dimnames=list(Components, Components))            #stores locations in Harv matrix of Harv coeffs adjusted for T.index
if(sum(is.HvstColsAdj) > 0)                                                                                                      #If any Harv cols are adjusted for alphaHvst.T, LoBoundHvst.T or HiBoundHvst.T,
   {N.HvstColsAdj <- sum(is.HvstColsAdj)                                                                                         #adjust the number of adjusted Harv cols
   FoodHetero.pairs.Adj4T <- NULL                                                                                                #create a pair.name matrix to be added to
   for(Ch in which(is.HvstColsAdj == TRUE))                                                                                      #For each harvester having Harv adjusted for T.index,
      {cons <- Components[Ch]                                                                                                    #the harvester's name
      prey.Rh <- which(Params[Rpar.Harv, Ch] > 0)                                                                                #row Ns in Harv of prey; orig Harvs from Params are used in case Harv values have been zeroed
      prey <- Components[prey.Rh]                                                                                                #vector of names of prey of harvester
      if(length(prey) > 0)                                                                                                       #If this harvester does have prey:-
         {FoodHetero.pairs.Adj4T <- c(FoodHetero.pairs.Adj4T, paste0("HarvAdj4T.", prey, ">", cons))                             #FoodHetero.pairs of its prey components
         is.HvstAdjust4T.matrix[prey.Rh, Ch] <- TRUE                                                                             #update this matrix location to TRUE
         }       
      }
   N.HarvCoefsAdj4T <- sum(is.HvstAdjust4T.matrix)                                                                               #N of Harv coeffs adjusted for T.index; TRUE=1, FALSE=0
   Hvst.Tadj <- array(data=0, dim=c(N.HarvCoefsAdj4T, TS.length), dimnames = list(FoodHetero.pairs.Adj4T, names(SimTime)))       #time-series array for Harv coeffs adjusted with T.index
   Hvst.Tadj[ , Last.Cts] <- Harv[which(is.HvstAdjust4T.matrix==TRUE, arr.ind=TRUE)]                                             #put initial, previously adjusted values of Harv into last col of Time Series
   RCHvst.Tadj <- which(is.HvstAdjust4T.matrix==TRUE, arr.ind=TRUE)                                                              #Rch holds row and col positions for each location in Harv holding adjustable Harv rates 
   }   
#----------------------------------------------------------------


#7.4 Left-side matrices of constraints
#The orderings of ObjFun elements, and constraint matrix cols and constraint limits, must be exactly the same
#Constraints are named with (i) a letter for the cause, if appropriate; (ii) a short name in upper case; (iii) a letter for the component (l.c) or component set (u.c) constrained
#Constraint matrices (or K in SigmaGP equations) contain the left-side coefficients of GP and pGP variables to be optimised by linear programming. They are prepared now
#Constraint limits (right-side maxima) must be adjusted with new Bmasses for each projection, so they are prepared within the #8.8 loop, not here.
#Non-negativity constraints, used for LP, are not needed because the R simplex() function automatically applies them
   #SUBSUBSECTIONS
      #7.4.1 Autotrophic GP constraints, GP.a
      #7.4.2 Partial GP constraints, m.pGP.g
      #7.4.3 Food constraints, m.FOOD.G and d.FOOD.S
      #7.4.4 NLO food constraints, d.FOOD.S
      #7.4.5 Heterotrophic GP constraints, M.SATE.h
      #7.4.6 Heterotrophic harvest constraints, M.HARV.h
      #7.4.7 Nutrient constraints, n.NUTR.A and n.EXCR.h
      #7.4.8 Fishery quota constraints, m.QUOTA.f
      #7.4.9 Habitat constraints, z.HAB.W

#7.4.1 Autotrophic GP constraints, GP.a
#GP.a constrains GP of autotrophs; required for all simulations
#All.C is the constraint coefficient matrix K in equation (2) of the theory paper describing this simulator; it is created here
GP.a.names <- paste0("GP.", Autotrophs)                                                                                #row names for GP.a constraints
All.C <- array(data=0, dim=c(N.Autos, N.LPvars), dimnames=list(GP.a.names, ObjFun.names))                              #create matrix for the Left sides of minimal constraints
GP.a.R <- 1:N.Autos                                                                                                    #set of rows (.R) in All.C in which GP.a is stored 
for(AllC.R in GP.a.R)                                                                                                  #for each autotroph,
   {All.C[AllC.R, Auto.Obj[AllC.R]] <- 1                                                                               #insert 1 in its row and col; this will include GP of each auto during LP
   }
N.C <- AllC.R                                                                                                          #total number of constraints so far

#7.4.2 Partial GP constraints, m.pGP.g
#m.pGP.g constrains pGP of every consumer g (including fisheries) from each of its foods m; required for all simulations
m.pGP.g.names <- paste0(sub(">", ".pGP.", FoodConsumer.pairs))                                                         #row names for m.pGP.g constraints
m.pGP.g.R <- (N.C + 1):(N.C + N.FoodConsumer.pairs)                                                                    #rows for the m.pGP.g constraints
All.C <- rbind(All.C, array(data=0, dim=c(N.FoodConsumer.pairs, N.LPvars), dimnames=list(m.pGP.g.names, ObjFun.names)))#bind rows for m.pGP.g onto All.C
for(AllC.R in m.pGP.g.R)                                                                                               #For each row of m.pGP.g:-
   {All.C[AllC.R, Consumer.Obj[AllC.R - N.C]] <- 1                                                                     #put 1 in col for this food-cons pair; Consumer.Obj holds cols of ObjFun for food>cons pairs, see #7.2
   }
N.C <- AllC.R                                                                                                          #total number of constraints so far

#7.4.3 Food constraints, m.FOOD.G
#m.FOOD.G constrains GPs of consumers of m and allows calculation of new bmasses of m from slack variables; required for all simulations
#Note: net removals of food during Delta_t must be < or = to food present at t, as adjusted for self losses and growth of the food. Thus LS coeffs are -ive where B_food is added to (eg. by GP) and +ive where harvested
#m.FOOD.G rows for m not a food, eg m is an unfished top predator, are present though redundant for LP; they are needed because their slack vars -> starting biomasses for the next projection
m.FOOD.G.names <- paste0(Wilds, ".FOOD.G")                                                                             #names for each wild m.FOOD.G row; suffix is .G (for consumers) if eaten
m.FOOD.G.names[!is.WildFood[1, Wilds]] <- paste0(Wilds[!is.WildFood[1, Wilds]], ".FOOD.")                              #no G suffix if m is not eaten, ie. it is an unfished top predator
m.FOOD.G.R <- (N.C + 1):(N.C + N.Wilds)                                                                                #rows for m.FOOD.G constraints in All.C; each row is a potential food but may be uneaten
mFGR.1 <- N.C                                                                                                          #store current number of constraints for indexing
All.C <- rbind(All.C, array(data=0, dim=c(N.Wilds, N.LPvars), dimnames=list(m.FOOD.G.names, ObjFun.names)))            #row-bind rows for m.FOOD.G constraints
N.C <- N.C + N.Wilds                                                                                                   #update number of constraint rows in All.C
#Find the harvest coefficient of GP to put into the left side of m.FOOD.G; there is one for each consumer of the food in the row so that matrix multiplication during LP supplies the sum of all harvests of this food
for(AllC.R in m.FOOD.G.R)                                                                                              #For each potential wild food m of any consumer:-
   {Param.C <- which(is.Wild)[AllC.R - mFGR.1]                                                                         #find the col#s in Params representing the food referred to by AllC.R
   AssimOfAllCrow.C <- which(Assim[Param.C, ] > 0)                                                                     #find col#s in Assim of this food's assimilators; named integer(0) of length=0 if there are none
   if(length(AssimOfAllCrow.C) > 0)                                                                                    #If food referred to by AllC.r is assimilated (ie. used for GP) by 1 or more components:-
      {ACReatenBy.Obj <- which(ObjFun.Foods.Loc == Components[Param.C])                                                #find col#s in objfunc where AllC.r food is eaten or fished productively
      All.C[AllC.R, ACReatenBy.Obj] <- InverseAssim[Param.C, AssimOfAllCrow.C]                                         #insert 1/Assim into row AllC.r for each col where AllC.r is eaten; estimates harvs of ACR by cons during LP
      }                                                                                                 
   #Correct the LS of m.FOOD.G for the GP of the food m itself
   Co <- which(ObjFun.GProducers.Loc == Components[Param.C])                                                           #cols of ObjFun for this food as producer (autos <- 1 col, prey <- 1 col for each of their foods, NLO <- 0)
   All.C[AllC.R, Co] <- All.C[AllC.R, Co] - 1                                                                          #insert -1s in AllC.r in each col for AllC.r as a GProducer.  Compensates for pGPs of the food itself
   }

#7.4.4 NLO food constraints, d.FOOD.S
#d.FOOD.S constrains scavengers eating NLO; required if NLO is present
d.FOOD.S.R <- NULL                                                                                                     #default row location in All.C if NLO is not in the model
if(N.NLOs > 0)                                                                                                         #In this version of ECOLPS, N.NLOs = 0 or 1 only; code below may allow for more for poss. developments
   {d.FOOD.S.names <- paste0(NLOs, ".FOOD.S")                                                                          #names for each NLO d.FOOD.S row; suffix is .S (for saprotrophs) if eaten
   d.FOOD.S.R <- (N.C + 1):(N.C + N.NLOs)                                                                              #rows for d.FOOD.S in All.C; each row is an NLO food for scavengers; here just one row because N.NLOs=1
   dFGR.1 <- N.C                                                                                                       #store for indexing
   All.C <- rbind(All.C, array(data=0, dim=c(N.NLOs, N.LPvars), dimnames=list(d.FOOD.S.names, ObjFun.names)))          #row-bind rows of d.FOOD.S constraint(s) onto the complete matrix of LS constraints
   N.C <- N.C + N.NLOs                                                                                                 #update the number of constraint rows in All.C
   for(AllC.R in d.FOOD.S.R)                                                                                           #For each type of NLO d (only 1 in this version of ECOLPS):-
      {Param.C  <- which(is.NLO)[AllC.R - dFGR.1]                                                                      #find col#s in Params for the NLO type referred to by AllC.R
      AssimOfAllCrow.C <- which(Assim[Param.C , ] > 0)                                                                 #find col#s in Assim of this NLO's assimilators, ie scavengers; named integer(0) of length=0 if none
      if(length(AssimOfAllCrow.C) > 0)                                                                                 #If NLO referred to by AllC.R is assimilated (ie. used for GP) by 1 or more components:-
         {ACReatenBy.Obj <- which(ObjFun.Foods.Loc == Components[Param.C])                                             #find col#s in obj.fun where AllC.R food is eaten productively; 1 col for each different scavenger
         All.C[AllC.R, ACReatenBy.Obj] <- 1                                                                            #insert 1s in AllC.R in cols where this NLO is eaten; sums harvs of food AllC.R=NLO (because Assim=1 for NLO)
         }
      }   
   #Correct the LS of d.FOOD.S for dropped harvests by wild consumers that do not assimilate 100% of the foods they harvest; code below assumes 1 NLO component only
   #Each food.consumer col in the LS matrix multiplies the pGP found by LP for that f.c.pair.  Entries below are pGainsToDropped.factor to convert pGP to unassimilated harvest dropped to NLO (assumed at bottom of sea)
   for(Gcol in which(is.Consumer))                                                                                     #find each consumer's column in Params and therefore in Growth, Assim, Harv, etc
      {for(Grow in which(is.WildFood))                                                                                 #find each wild food's column ditto; excludes NLO which, if not 100% assimilated, is dropped back to NLO
         {Of.name <- paste0(Components[Grow], ">", Components[Gcol])                                                   #fabricate name "food>consumer", as used in the O.f, to find column in Obj.fun having the same header
         ColInObjFun <- which(ObjFun.names == Of.name)                                                                 #find the col# in O.f that matches this food>consumer
         ColIn.pGTD.f <- which(pGains.Loc == Of.name)                                                                  #find the col# in pGainsToDropped.factor that matches this food>consumer
         All.C[d.FOOD.S.R, ColInObjFun] <- -pGainsToDropped.factor[1, ColIn.pGTD.f]                                    #put pGTD.factor in LS of the d.FOOD.S constraint (which here has 1 row only)
         }
      }   
   }

#7.4.5 Heterotrophic GP constraints, M.SATE.h
#M.SATE.h constrains total GP of h from all of h's foods because of max physiological growth capabilities
#Note: In R, a NULL vector has testable length=0; an NA vector has length >or= 1, and is.na(vec) -> error if length(vec) > 1
M.SATE.h.R <- NULL                                                                                                     #default if Gmax coeffs were not read
if(!is.na(Gmax0.Rpar))                                                                                                 #If a row of Gmax coeffs was read from Params.file:-
   {All.C <- rbind(All.C, array(data=0, dim=c(N.Heteros, N.LPvars), dimnames=list(paste("M.SATE.", Heterotrophs, sep=""), ObjFun.names))) #1 row put on All.C for every heterotroph
   M.SATE.h.R <- (N.C + 1):(N.C + N.Heteros)                                                                           #rows for the M.SATE.h constraints; each row is a heterotroph
   for(AllC.R in M.SATE.h.R)                                                                                           #For each row of M.SATE.h,
      {All.C[AllC.R, which(ObjFun.Heteros.Loc == Heterotrophs[AllC.R - N.C])] <- 1                                     #insert 1 in each food col of hetero ACR; this sums pGains across all foods of ACR during LP
      }      
   N.C <- AllC.R                                                                                                       #total number of constraints so far
   }

#7.4.6 Heterotrophic harvest constraints, M.HARV.h
#M.HARV.h constrains total harvest of h from all of its foods because of max physiological harvesting capabilities
M.HARV.h.R <- NULL                                                                                                     #default if Hmax coeffs were not read
if(!is.na(Hmax0.Rpar))                                                                                                 #If a row of Hmax coeffs was read from Params.file:-
   {All.C <- rbind(All.C, array(data=0, dim=c(N.Heteros, N.LPvars), dimnames=list(paste("M.HARV.", Heterotrophs, sep=""), ObjFun.names))) #1 row for every heterotroph
   M.HARV.h.R <- (N.C + 1):(N.C + N.Heteros)                                                                           #rows for the M.HARV.h constraints; each row is a heterotroph
   for(AllC.R in M.HARV.h.R)                                                                                           #For each row of M.HARV.h,
      {FoodsOfACR.Obj <- which(ObjFun.Heteros.Loc == Heterotrophs[AllC.R - N.C])                                       #find food cols in ObjFun for hetero ACR
      Thisis.Hetero <- which(Components == Heterotrophs[AllC.R - N.C])                                                 #find col number in Params of hetero ACR
      FoodsOfACR.Rpar <- which(Assim[ , Thisis.Hetero] > 0)                                                            #find food rows in Assim that are assimilated by hetero ACR
      All.C[AllC.R, which(ObjFun.Heteros.Loc == Heterotrophs[AllC.R - N.C])] <- InverseAssim[FoodsOfACR.Rpar, Thisis.Hetero] #1/Assim in food row this hetero; this sums pHarvs across all foods of ACR during LP
      }      
   N.C <- AllC.R                                                                                                       #total number of constraints so far
   } 

#7.4.7 Nutrient constraints, n.NUTR.A, n.EXCR.h
#7.4.7.1 n.NUTR.A constraints
#n.NUTR.A constrains growth of autotrophs when nutrients essential for photosynthesis are limiting in the water.  
 #Note.1: not all nutrient constraints are relevant, eg. when Nconc is zero or Bmass is zero.  Removing irrelevant rows caused coding complications later, so irrelevant rows have been left in.  Users should be aware that false Activity of an irrelevant constraint can occur because biomass_t=0 or Nconc=0 => a limit that is zero => the slack variable is zero => false activity is signalled.
#Note.2: These constraints, as coded, are not applicable to phytoheterotrophs, e.g. corals, taking up nutrients from both water and foods.  Untested suggestions for them: 1) set Nconc for tissues of phytoheterotrophs low to prevent the n.NUTR.A constraint limiting photosynthetic growth; 2) increase n.EXCR.h right-side limit (see below) to Inf for the phytoheterotrophs to prevent the n.EXCR.h constraint limiting their heterotrophic growth; or 3) input the auto and hetero parts as separate components with the hetero feeding on the auto and selfloss of both inversely dependent on biomasses of the other.
n.NUTR.A <- NULL
n.NUTR.A.R <- NULL                                                                                                     #defaults if no nutrients in the model
if(N.Nutr > 0)                                                                                                         #If nutrients are in the model (because Nmass rows were supplied):-
   {#create rows for n.NUTR.A
   All.C <- rbind(All.C, array(data=0, dim=c(N.Nutr, N.LPvars), dimnames=list(paste0(Nutrients, ".NUTR.A"), ObjFun.names))) #add a blank N.Nutr x N.LPvars array for the n.NUTR.A rows of All.C 
   N.C1 <- N.C + N.Nutr                                                                                                #row number in All.C of last n.NUTR.A row; N.C1 is used to count last row
   n.NUTR.A.R <- (N.C + 1):N.C1                                                                                        #row numbers in All.C for the n.NUTR.A constraints; each row is for one nutrient
   N.C <- N.C1                                                                                                         #total number of constraints so far
   All.C[n.NUTR.A.R, Auto.Obj] <- Nconc[1:N.Nutr, is.Auto]          #insert tissue concs of nutrients as coeffs for autotrophs; Nconc has the same col ordering as Components; Auto.Obj=>autotrophic column elements of ObjFun
   #subtract n excreted into water by heterotrophs
   for(Het in 1:N.Heteros)                                                                                             #For each heterotroph,
      {HetAsCons.Obj <- which(ObjFun.Heteros.Loc == Heterotrophs[Het])                                                 #col(s) in the ObjFun row vector holding this hetero as a consumer
      FoodsOfThisHet <- ObjFun.Foods.Loc[HetAsCons.Obj]                                                                #names of foods of this hetero; the foods may be autos, heteros or NLO
      FoodsOfThisHet.pairs <- paste(FoodsOfThisHet, Heterotrophs[Het], sep=">")                                        #make names to match format of ObjFun names with these foods and this hetero
      FoodsOfThisHetero.Obj <- which(ObjFun.names %in% FoodsOfThisHet.pairs)                                           #cols of ObjFun matched by these food-hetero pairs
      FoodsOfThisHet.RC <- which(Components %in% FoodsOfThisHet)                                                       #RC=rows of biomass, rows of Assim, and cols of Nconc for foods of this hetero
      HetAsCons.RC <- which(Components == Heterotrophs[Het])                                                           #RC=row of biomass, col of Assim, and col of Nconc for this hetero as a consumer 
      for(foth in 1:length(FoodsOfThisHet))                                                                            #For each Food of this Hetero, 
         {netConc <- Nconc[ , FoodsOfThisHet.RC[foth]] - Nconc[ , HetAsCons.RC]                                        #conc difference: Nconc in food of this hetero - Nconc in hetero. There is one element for each nutrient
         All.C[n.NUTR.A.R, FoodsOfThisHetero.Obj[foth]] <- -netConc                                                    #insert -conc_n,m + conc_n,h at col for m eaten by h and row for nutrient n 
         }
      }   
   }
#7.4.7.2 n.EXCR.h constraints
#n.EXCR.h limits heterotrophic GP if they cannot take in sufficient nutrient from their food. 
n.EXCR.h <- NULL                                                                                                       #dummy in case called
n.EXCR.h.R <- NULL                                                                                                     #dummy for the rows in constraint coeff matrix
if(N.Nutr > 0)                                                                                                         #If nutrients are in the model,
   {#create rows for n.EXCR.h ; only include rows with conc_n,h > 0 because only those can constrain GP of h           #this is to cut out ineffective constraints
   n.EXCR.h <- array(data=0, dim=c(1, N.LPvars), dimnames=list(NULL, ObjFun.names))                                    #initiate a coeff matrix for n.EXCR.h
   N.C1 <- N.C + 1                                                                                                     #mark the 1st row of n.EXCR.h constraints in All.C
   excr <- 1                                                                                                           #counter for each n.EXCR.h constraints, 1 per hetero per nutrient
   for(Het in 1:N.Heteros)                                                                                             #For each heterotroph,
      {FoodsOfHet.Obj <- which(ObjFun.Heteros.Loc == Heterotrophs[Het])                                                #locate the positions of its foods in the objective function
      FoodsOfHet.RC <- which(Components %in% ObjFun.Foods.Loc[FoodsOfHet.Obj])                                         #locate the positions of its foods in Nconc (with cols for each component)
      for(Nut in 1:N.Nutr)                                                                                             #For each nutrient in the model,
         {if(Nconc[Nut, N.Autos + Het] > 0)                                                                            #Only If Nconc of hetero is > 0,
            {for(food in 1:length(FoodsOfHet.Obj))                                                                     #For each food of Het,
               {n.EXCR.h[excr, FoodsOfHet.Obj] <- Nconc[Nut, N.Autos + Het] - Nconc[Nut, FoodsOfHet.RC]                #add hetero's Nconc and subtract food's Nconc
               }                                                                                                       
            row.names(n.EXCR.h)[excr] <- paste0(Nutrients[Nut], ".EXCR.", Heterotrophs[Het])                           #label this row
            n.EXCR.h <- rbind(n.EXCR.h, array(data=0, dim=c(1, N.LPvars)))                                             #bind on the next row, empty
            excr <- excr + 1                                                                                           #update the row counter
            }
         }
      }
   n.EXCR.h <- n.EXCR.h[-excr,  ]                                                                                      #remove the last empty row
   N.C <- N.C + excr -1                                                                                                #update the number of constraints in All.C
   n.EXCR.h.R <- N.C1 : N.C                                                                                            #store the rows in All.C holding potentially active n.EXCR.h constraints
   All.C <- rbind(All.C, n.EXCR.h)                                                                                     #bind rows of n.EXCR.h onto existing All.C coeff matrix
   }

#7.4.8 Fishery quota constraints, m.QUOTA.f
#Note: some supplied quota may not be fished against, eg. when Growth[food, fishery] = 0
m.QUOTA.f.R <- NULL                                                                                                    #default row location in All.C if quota components are not in the model
if(N.ActiveQuotaFisheries > 0)                                                                                         #If any actively fished quota are in the model:-  
   {All.C <- rbind(All.C, array(data=0, dim=c(N.ActiveQuota, N.LPvars), dimnames=list(paste0(ObjFun.Foods.Loc[Quota.Obj], ".QUOTA.",  ObjFun.Fisheries.Loc[Quota.Obj]), ObjFun.names))) #rbind quota rows; Quota.Obj defined in #7.2
   m.QUOTA.f.R <- (N.C + 1):(N.C + N.ActiveQuota)                                                                      #number of rows for the m.QUOTA.f constraints; each row is a quotum-fishery pair. 
   for(AllC.R in m.QUOTA.f.R)                                                                                          #for each quotum row,
      {All.C[AllC.R, Quota.Obj[AllC.R - N.C]] <- 1                                                                     #put 1 into cols of fished quotum-fishery pairs => summation of GPs during LP for this l-f pair
      }
   N.C <- nrow(All.C)                                                                                                  #renew count of all constraints
   }
   
#7.4.9 Habitat constraints, z.HAB.W
#z.HAB.W constrains retained GP of any W requiring finite habitat z
#Note: HabOccup is N.Hab x N.Components. It pre-multiplies Temp.z.HAB.W to give an N.Hab x N.LPvars matrix; this are habitat rows in the K matrix which pre-multiplies vector j (see eq 2 in Part 1) so that constraint limits y are not exceeded.
z.HAB.W.R <- NULL                                                                                                      #default row location in All.C if habitat components are not in the model
if(N.Habitats > 0)                                                                                                     #If essential habitats are in the model:-
   {#the negative m.FOOD.G matrix is needed for habitat constraints; the NLO food row is excluded from Temp.z.HAB.W because NLO is not limited by habitat; m.FOOD.G from #7.4.3
   if(N.NLOs == 0) Temp.z.HAB.W <- -All.C[m.FOOD.G.R, ] else Temp.z.HAB.W <- -All.C[m.FOOD.G.R, ][-which(is.NLO), ]    #store temporarily omitting an NLO row if present
   All.C <- rbind(All.C, array(data=0, dim=c(N.Habitats, N.LPvars), dimnames=list(paste0(Habitats, ".HAB.W"), ObjFun.names))) #initialise habitat constraint rows at the end of the current All.C matrix
   z.HAB.W.R <- (N.C + 1):(N.C + N.Habitats)                                                                           #z.HAB.W rows in All.C; each is an essential habitat for one or more wild components
   All.C[z.HAB.W.R, ] <- HabOccup[ , is.Wild] %*% Temp.z.HAB.W                                                         #coeffs in z.HAB.W are found by matrix multiplication
   N.C <- nrow(All.C)                                                                                                  #renew count of all constraints
   }
#----------------------------------------------------------------


#7.5 Constraint-related time series
#These are needed for interpreting time series of simulated variables
Slacks <- array(data=NA, dim=c(N.C, TS.length), dimnames = list(paste0("Slack.", substr(row.names(All.C), 1, 15)), names(SimTime)))   #slack-variable time series; there are N.C constraints; slack=0 if constraining
Zero.Slacks <- array(data=NA, dim=c(1, TS.length), dimnames = list("N.ZeroSlacks", names(SimTime)))                                   #number of zero slack variables to show active constraints as a time series
Limits <- array(data=NA, dim=c(N.C, TS.length), dimnames = list(paste0("Lim.", substr(row.names(All.C), 1, 15)), names(SimTime)))     #constraint-limit time series; limit -> max GP possible if no other constraints are active
#----------------------------------------------------------------


#7.6 Non-time-series structures for simulations.
#Elements of non-time-series structures are re-written for every Delta but the matrix structures stay the same so they are created once here
#Clarifications: Gains for fisheries are landings; losses of wilds to fisheries are catches. Gains for NLO are dropped harvests and dropped particulate self losses; self losses from NLO are only dissolved self losses going to water; particulate self losses of NLO to water are zero (they fall back to NLO); harvests of NLO by scavengers are biomass of NLO
   #SUBSUBSECTIONS
      #7.6.1 Non-time-series structures for 1st-order Taylor series projections
      #7.6.2 Non-time-series structures for 2nd-order Taylor series projections
      #7.6.3 Non-time-series structures for 3rd-order Taylor series projections
      #7.6.4 Other matrices re-used for each projection

#7.6.1 Non-time-series structures for 1st-order Taylor series projections
#In this version of ECOLPS, matrix dimensions not involving .pairs include all N.Components, not just the relevant N.variables; this reduces the number of row and col indices needed -> easier maintenance of code.  The memory wasted with rows entirely filled by zeros is less important.
GainR.names <- paste("from.", Components, sep="")                                                                                #row names for proj Gain matrices
GainC.names <- paste("Gains.", Components, sep="")                                                                               #col names for proj Gain matrices
LossR.names <- paste("Loss.", Components, sep="")                                                                                #row names for proj Loss matrix
LossC.names <- paste("to.", Components, sep="")                                                                                  #col names for proj Loss matrix
BmassR.names <- paste(Components, ".Bmass", sep="")                                                                              #row names for BmassEffort.prods
BmassC.names <- paste(Components, ".Bmass", sep="")                                                                              #col names for BmassEffort.prods, multiplication symbol "x " not yet added
if(N.Fisheries > 0)                                                                                                              #If fisheries are in the model:-
   {GainC.names[is.Fishery] <- paste0("Landed.", Fisheries)                                                                      #col names for fishery-landing cols in Gain matrix
   LossC.names[is.Fishery] <- paste0("Caught.", Fisheries)                                                                       #col names for fishery-catch cols in Loss matrix
   BmassC.names[is.Fishery] <- paste0(Fisheries, ".Effort")                                                                      #col names for any fishery components in biomass x biomassOrEffort product matrices
   BmassR.names[is.Fishery] <- paste0(Fisheries, ".Effort")                                                                      #row names for any fishery components in biomass x biomassOrEffort product matrices
   }
projD1.pGains <- array(data=0, dim=c(N.Components, N.Components), dimnames=list(GainR.names, paste0("projD1.p", GainC.names)))   #projected 1st deriv Gain matrix (this is G dot in SI Annex A)
projD1.pLosses <- array(data=0, dim=c(N.Components, N.Components), dimnames=list(paste0("projD1.", LossR.names), LossC.names))   #projected 1st deriv Loss matrix (this is L dot in SI Annex A)
BmassEffort.prods <- array(data=0, dim=c(N.Components, N.Components), dimnames=list(BmassR.names, paste0("x ", BmassC.names)))   #current biomass x (biomass and fishing effort) products
proj.UnAssimHarvs <- array(data=0, dim=c(max(1, N.UnAssimHarv.pairs), 1), dimnames = list(paste0("proj.UnAssimHvst.", UnAssimHarv.pairs), "UnAssimHarvest")) #proj.UnAssimilated harvests with +ve Harv and 0 Assim; may be none

#7.6.2 Non-time-series structures for 2nd-order Taylor series projections
projD2.pGains <- array(data=0, dim=c(N.Components, N.Components), dimnames=list(GainR.names, paste0("projD2.", GainC.names)))    #projected 2nd deriv gain matrix (this is G dot dot in SI Annex A)
projD2.pLosses <- array(data=0, dim=c(N.Components, N.Components), dimnames=list(paste0("projD2.", LossR.names), LossC.names))   #projected 2nd deriv Loss matrix (this is L dot dot in SI Annex A)
D1.Bmass.BorE <- array(data=0, dim = c(N.Components, N.Components), dimnames = list(paste0("D1.", BmassR.names), paste0("x", BmassC.names))) #D1.B x (B or E)
Bmass.D1BorE <- array(data=0, dim = c(N.Components, N.Components), dimnames = list(BmassR.names, paste0("xD1.", BmassC.names)))  #Bmass x D1.(Bmass, Effort) 
D1.BmassEffort.prods <- array(data=0, dim = c(N.Components, N.Components), dimnames = list(paste0("D1.(",BmassR.names), paste0("x", BmassC.names, ")"))) #D1.(B x (B, E))      

#7.6.3 Non-time-series structures for 3rd-order Taylor series projections
projD3.pGains <- array(data=0, dim=c(N.Components, N.Components), dimnames=list(GainR.names, paste0("projD3.", GainC.names)))    #projected 3rd deriv Gain matrix (this is G dot dot dot in SI Annex A)
projD3.pLosses <- array(data=0, dim=c(N.Components, N.Components), dimnames=list(paste0("projD3.", LossR.names), LossC.names))   #projected 3rd deriv Loss matrix (this is L dot dot dot in SI Annex A)
D1.Bmass.D1BorE <- array(data=0, dim = c(N.Components, N.Components), dimnames = list(paste0("D1.", BmassR.names), paste("x D1.", BmassC.names, sep=""))) #D1.B x D1.(B, E) 
Bmass.D2BorE <- array(data=0, dim = c(N.Components, N.Components), dimnames = list(BmassR.names, paste0("x D2.", BmassC.names))) #Bmass x D2.(B, Eff)
D2.Bmass.BorE <- array(data=0, dim = c(N.Components, N.Components), dimnames = list(paste0("D2.", BmassR.names), paste0("x ", BmassC.names))) #D2.B x (B, E)
D2.BmassEffort.prods <- array(data=0, dim = c(N.Components, N.Components), dimnames = list(paste0("D2.(",BmassR.names), paste0("x ", BmassC.names, ")"))) #D2.(B x (B, E))

#7.6.4 Other matrices re-used for each projection
All.Cmax <- array(data=NA, dim=c(N.C, 1), dimnames=list(row.names(All.C), "MAXIMUM"))                                            #col array for constraint maxima; N.C is the total number of constraints
proj.pGains <- array(data=0, dim=c(N.Components, N.Components), dimnames = list(paste0("from.", Components), paste0("proj.", GainC.names)))#projected pGains by Taylor series; partial Gain <- row & col position
#----------------------------------------------------------------


#7.7 Other variables and constants needed for simulation steps
#Alphabetic order is presented so far as code needs allow
EndOut <- N.LPproj + 1                                           #default end of outputs
NumericConstrOptions <- c("0":"15")                              #fixed number of numeric options in constraints menu as a string; this MUST be changed if number of numeric options are changed
NumericVarsOptions <- c("1":"5","11":"15","21":"25","31":"37","38","41":"43","51":"55","61":"66","71","72") #fixed number of numeric options in variables menu as a string; this MUST be changed if numeric options are changed
PrintOut.interval <- 1                                           #default time increment (Deltas) between printed and graphed time points; 1 -> every point is shown
StartOut <- 1                                                    #graph plotting AND printing of output starts from this element of SimTime by default
OutputDeltas <- seq(from=StartOut, to=EndOut, by=PrintOut.interval) #default Delta values used for printing/graphing time series

AllowedConstrResponses <- c("", NumericConstrOptions,"A","a","B","b","C","c","E","e","F","f","G","g","I","i","J","j","K","k","L","l","M","m","N","n","O","o","P","p","Q","q","R","r","S","s","T","t","U","u","w","W","X","x","Y","y","z","Z")                                                           #allowed responses to Constraints menu; MUST be changed if allowed responses are added with new code, or removed
AllowedMainResponses <- c("","1":"9","C","c", "R", "r","U","u")  #allowed responses to Main menu; MUST be changed if allowed responses are added with new code, or removed
AllowedVarsResponses <- c("", NumericVarsOptions,"A","a","B","b","C","c","E","e","F","f","G","g","I","i","J","j","K","k","L","l","M","m","N","n","O","o","P","p","Q","q","R","r","S","s","T","t","U","u","V","v","w","W","X","x","Y","y","z","Z")                                                       #allowed responses to Variables menu; ; MUST be changed if allowed responses are added with new code, or removed
AxLabelFactor <- 1.5                                             #default script size of X and Y axes labels for graphs; change with S option in submenus
Block <- 0                                                       #number of this block of simulations. 1 is added in the projection loop below. MainMenu, below, allows further blocks of simulations
Coef1 <- 0.5                                                     #coefficient for 1st-order terms of a projected integral (constant = 1/2); see Annex A of paper
Coef2 <- 0.1666666667                                            #coefficient for 2nd-order integral terms (constant = 1/6)
Coef3 <- 0.0416666667                                            #coefficient for 3rd-order integral terms (constant = 1/24)
Col.plots <- 1                                                   #set default = 1 = graph symbols in colour; change with Q option in submenus
ConstrCount <- 0                                                 #ConstrCount allows 'while()' looping below until ESC is pressed
DefaultMainMenuChoice <- 2                                       #default main-menu choice when [Enter] is pressed without any other typed input
DefaultTitleText <- Params.file                                  #default title for printed and graphic output; see ProcessOutput() where date & time is added; change with T option in menus
DeleteEmptySeries <- TRUE                                        #default: automatically delete empty series (holding nothing but 0s or NAs) from the Output.set sent for graphing or printing
EndOutYear <- Years[EndOut]                                      #default end-of-outputs in years; EndOut is above
FirstMenu <- TRUE                                                #if TRUE, Main Menu is not shown to user 
GraphLabel <- "*"                                                #default text applied to graph when menu option L=label is chosen
Graph.plotted <- FALSE                                           #if FALSE, prevents addition of text with the L option in submenus if a graph does not yet exist
is.Output.set <- FALSE                                           #if FALSE, an Output.set must be created before adding rows; if TRUE, new rows are appended to the set already existing
is.slacks.out <- TRUE                                            #if TRUE, slack variables are selected for output by the constraints menu
ItemName <- NA                                                   #name of the type of variable selected for the Output.set.  NA needed for the 1st main menu 
Jitter <- 0                                                      #default is no jitter applied to plotted points
KeyAuto <- TRUE                                                  #number of lines in graph keys is set automatically until over-ruled by the K option in Vars and Constraint's menus
LengthOut <- EndOut - StartOut + 1                               #length (number of deltas) of the complete Output.set in Deltas
MainCount <- 0                                                   #MainCount allows 'while()' looping below until ESC is pressed
MainMessage0<- c("See information read from inputs listed above depending on ADVICE LEVEL:- AL =", AL, ".") #this message opens the main menu
MaxOrdinate <- Inf                                               #set maximum ordinate value > 0; Inf is unrestrictive; fixing the max allows matching ordinate scales in related graphs
N.ActiveConstraintsSelected <- 0                                 #set the number of active constraints so far selected; used to create activity markers up the ordinate when active constraints selected from constraints menu
N.extraYears <- 0                                                #the number of years appended to the simulation by use of MainMenuOption=9; 0 here because option 9 not yet available
N.keyLines <- 4                                                  #number of lines allowed for key; a value is needed for function calls even if KeyAuto is TRUE; too many lines cramp the graph; too few crops the key
N.OutputDeltas <- length(OutputDeltas)                           #default number of time points to print; adjustable with B and F options in submenus
N.progressReports <- 0                                           #number of progress reports screened
N.requestedRows.OS <- 0                                          #number of rows (=variables) requested for Output.set, including some that may be empty
N.rows.OS <- 0                                                   #number of rows (=variables) in the Output.set for printing, graphing or analysis, currently none; added to in submenus; clear with C option
nextStep <- 0                                                    #default interval = nextStep+1 between active constraints going up ordinate
Ordinate.factor <- 1                                             #default spacing of active constraint markers up the graphic ordinate; 1 gives 1, 2, 3; 2 gives 2 4 6 etc
pc <- 0                                                          #current CPU time; used for timing LP steps
pc.total <- 0                                                    #total CPU time used for LP steps
PlotSymbolSize <- 0.8                                            #set size of plot symbol for R's cex parameter; typical value = 0.8, allowed = 0.5 to 2.0 by 0.01
Premature.end <- FALSE                                           #if TRUE, the requested N.LPproj loops were/not completed      
ShowMainMenu <- FALSE                                            #if TRUE, the Main Menu is not screened; it is omitted for first block simulations
Simulation <- TRUE                                               #if TRUE, allow the 1st LP step
SlacksMessage <- ""                                              #this stub message is printed with the slack variables menu to alert user to output or the lack of it
StartOutYear <- Years[StartOut+1]                                #graph plotting AND printing of output starts from this year
Thin <- 0                                                        #thinning of output points: = PrintOut.interval -1;  therefore 0 -> show every point; adjustable with I option in submenus
TickMarkNumberSize <- 1                                          #default size of tick mark numbers on both X and Y axes; change with S option in submenus
TimeLabel <- TimeLabel.def <- "Years" #"Time, \U0394s"           #default label for X axis of graphs. Use "\U0394s" for 'Deltas'; change with X option in submenus
TitleText <- DefaultTitleText                                    #set title to default text
TitleTime <- TRUE                                                #include date and time in the title
VarsCount <- 0                                                   #VarsCount allows 'while()' looping below until ESC is pressed
Vars.loc <- 0                                                    #start row for searching Environ.file if present
VarsMessage1 <- ""                                               #supplementary Vars message
#----------------------------------------------------------------
#################################################################End of section 7. INITIALISE DATA STRUCTURES FOR SIMULATIONS

                                                                                                      

#8. MENUS AND SIMULATION LOOPS
   #SUBSECTIONS
      #8.0 Main menu
      #8.1 Select CONSTRAINTS for output (MainMenuChoice == "1")
      #8.2 Select SIMULATED VARIABLES for printed or graphic output (MainMenuChoice == "2")
      #8.3 Select NUMBER OF ZERO SLACK VARIABLES for printed or graphic output (MainMenuChoice == "3")
      #8.4 Show LP VARIABLES CALCULATED for the last projection step (MainMenuChoice == 4)
      #8.5 Reset default response to this menu (MainMenuChoice == "5")
      #8.6 Toggle DELETION of empty rows from Output.set AUTOMATICALLY (Default=Delete)(MainMenuChoice == "6")
      #8.7 Show INPUT data set
      #8.8-8.9 Simulate N.LPproj times, OVER-WRITING (MainMenuChoice =8) or APPENDING TO (MainMenuChoice =9) any pre-existing results
      #8.10 Clear or Unclear Output.set
      #8.11 End of Main Menu loop, and end of main program
#Note: To add another menu item, you must program it below and add the keyboard response to AllowedMainResponses in #7.7

#8.0 Main menu
#Logic: when processing first arrives here, ShowMainMenu=FALSE signals that the main menu is not screened, and a MainMenuChoice of 8 is automatically supplied.  Processing then passes directly through to the linear programming stage; Block number is incremented from 0 to 1 -> main menu is shown after N.LPproj projections for the 1st block -> any further processing is controlled by the user.  This logic is used so that the LP code need only be in one place, and not be written as a function (which hindered de-bugging).
#In this section, a while() loop is started to bring the user back to the main menu (up to 100 times if needed).  Keyboard input is dealt with after the menu has been screened      
while(MainCount < 100)                                                                                       #allows this many repeat visits to main menu; the while() loop extends to the end of this file
   {if(Block > 0)
      {cat("\n\nMAIN MENU Options for ", M, " simulation steps stored as time series:-")                     #M, the LP projection counter, is used here, not N.LPproj, in case projections terminated prematurely
      cat("
1 -> Select CONSTRAINTS for printed or graphic output
2 -> Select SIMULATED VARIABLES for printed or graphic output  
3 -> Print NUMBERS of ZERO SLACK VARIABLES
4 -> Show PARAMETERS and VARIABLES CALCULATED for the LAST simulation step
5 -> New DEFAULT RESPONSE for this menu 
6 -> Toggle DELETION of empty rows from Output.set AUTOMATICALLY (currently:-", ifelse(DeleteEmptySeries==TRUE, "Deleting", "NOT deleting"), ")
7 -> Show original INPUT table for parameters 
8 -> CONTINUE current simulation, OVER-WRITING existing results     
9 -> CONTINUE current simulation, APPENDING new results
C -> CLEAR current Output.set      R -> REMOVE the active graphics window      
[Enter] -> DEFAULT response = ", DefaultMainMenuChoice, "
[ESC] -> Terminate simulation (variables remain in memory)                                                                                
")
      if(MainCount == 1) cat("\a\nADVICE:", MainMessage0)                                                    #main message0 (written in #7.7) is only shown in loop 1. \a rings bell to notify end of processing and MAIN menu
      if(MainCount > 0)  cat("\a\nADVICE:", MainMessage, "\n")                                               #main message (without the 0, written below,) is always shown after block 0 is completed
      RepeatedMainMessage <- paste0("OUTPUT.SET now contains ", ifelse(is.Output.set == FALSE, 0, N.rows.OS), " time series and spans ", Years[1], " to ", Years[EndOut], " years thinned by ", Thin, " points. \nENTER 1 number, 1 letter, or nothing-> default choice:-\n")                                                                   #tells what is already in Output.set; if these series are not needed, use C=clear output.set, or E   =edit output.set
      cat(RepeatedMainMessage)                                                                               #screen repeated message (see above) updates on the size of the Output.set, and tells how to enter a response
#Deal with keyboard input
      MainMenuChoice <- as.character(readLines(n=1))                                                         #read one line of keystrokes followed by [ENTER] 
      MainMenuChoice <- gsub(" ", "", MainMenuChoice)                                                        #remove a leading blank from keyboard input
      MainMenuChoice <- substr(MainMenuChoice,1, 1)                                                          #only the first character is read with this menu
      if(MainMenuChoice %in% AllowedMainResponses == FALSE | length(MainMenuChoice) > 1)                     #If input character is not assigned, [see AMR in #7.7]; or more than 1 number or letter was typed:-
         {MainMessage <- c("Input unrecognised.  Please enter exactly 1 number or letter from ", AllowedMainResponses, "or [Enter] or [ESC].") #advise
         next()                                                                                              #and return to Main Menu
         }
      if(MainMenuChoice %in% c("8","9") & Premature.end == TRUE)                                             #If continuing simulations are requested when previous block was not completed:-
         {MainMessage <- "The last block of simulations could not be completed.  Further simulations are not possible."     #advise
         next()                                                                                              #and return to Main menu
         }
      if(MainMenuChoice == "" )                                                                              #If [ENTER] by itself is pressed, it is read as ""    
         {MainMenuChoice <- DefaultMainMenuChoice                                                            #provide the default response
         cat(DefaultMainMenuChoice, "\n")                                                                    #show choice on screen
         }
      } else                                                                                                 #Else, if this is simulation block number 0, ie. the first,
      {ShowMainMenu <- TRUE                                                                                  #show MM for blocks 1 or higher, not for block 0
      MainMenuChoice <- 8                                                                                    #set choice=8 to go directly to simulation code for block 0
      }
#----------------------------------------------------------------


#8.1 Select CONSTRAINTS for printed or graphic output (MainMenuChoice == "1")
   #SUBSUBSECTIONS
      #8.1.0 Constraints menu
      #8.1.1 Deal with keyboard input to Constraint menu
      #8.1.2 Selected option from Constraints' menu was numerical
      #8.1.3 Selected option from Constraints' menu was alphabetic
#Note: if Constraints-menu options are changed, AllowedConstrResponses in #7.7 must also be changed   

#8.1.0 Constraints menu      
   if(MainMenuChoice == "1")                                                                                 #If output of active constraints is requested:-
      {ItemName <- "Active Constraints"                                                                      #name the default output data set for heading printouts 
      ItemName2 <- "Activity markers"                                                                        #name the constraint property to be selected for output
      ItemN <- 1                                                                                             #item number = 1 denotes Activity markers
      ConstrMessage <- paste0("\nADVICE: You have moved to CONSTRAINTS MENU from Main Menu (via option ", MainMenuChoice,  "): ") #this message opens constraints menu at first showing     
      if(ConstrCount == 0) Ordinate.label <- ItemName                                                        #O.l is set for 1st time; alter with ConstrMenu option "Y"; CC was 0'd in #7.7
      ConstrCount <- 0                                                                                       #zero constraint-menu loop counter
      while(ConstrCount < 100)                                                                               #allows this many repeat visits to constraints menu; end of while loop is at end of #8.1.2
         {RepeatedConstrMessage <- paste0("\bADVICE: OUTPUT.SET now contains ", N.rows.OS, " time series and spans ", Years[StartOut], " to ", Years[EndOut], " years thinned by ", Thin, " points")
         cat("\n\nCONSTRAINTS MENU. Options for ", M, " simulation steps stored as time series.") 
         cat("
0 -> TOGGLES which constraint variable for OUTPUT: Activity markers at t+1 OR Slack variables at t OR Constraint limits at t 
Note: Activity may be shown falsely if B_w,t or Q_n,w,t = 0 and, as a result, the constraint limit = 0
1 -> ALL constraints        2 -> AUTOTROPHIC GP (GP.a)       3 -> CONSUMER pGP (m.pGP.g)     4 -> HETEROTROPHIC GPs (M.SATE.h)
5 -> HETEROTROPHIC HARVEST (M.HARV.h)        6 -> LIVING FOOD (m.FOOD.G)                     7 -> NLO FOOD (d.FOOD.S)
8 -> AUTOTROPHIC NUTRIENT (n.NUTR.A)         9 -> HETEROTROPHIC NUTRIENT (n.EXCR.h)                
10-> PARTIAL LANDINGS (m.pGP.g subset)       11-> FISHERIES QUOTA (m.QUOTA.f)                12-> ESSENTIAL HABITAT (z.HAB.W)
13-> Constraints with AUTOTROPHS             14-> Constraints with HETEROTROPHS              15-> Constraints with FISHERIES
A -> ACTIVITY-marker ordinates               B -> BEGINNING time of Output                   C -> CLEAR current Output.set
E -> EDIT rows of Output.set                 F -> FINISHING time of Output                   G -> GRAPH current Output.set
I -> Time INTERVAL between points            K -> Number of KEY lines                        L -> Add text LABEL to ACTIVE graph
M -> Column TOTALS                           N -> NUMBER of positives                        O -> max ORDINATE        
P -> PRINT current Output.set to screen      Q -> COLOUR / B&W graphic symbols               R -> REMOVE the active GRAPHICS window       
S -> Size of axes' labelling                 T -> TITLE of output                            W -> Graph WINDOW width and height           
X or Y -> Label for X or Y axis              Z -> Size of plot SYMBOL                       [Enter] -> Return to MAIN MENU
[ESC] -> Terminate simulation (variables remain in memory)\n ")            
         cat(ConstrMessage)                                                                                  #screen the new message
         cat("\n", RepeatedConstrMessage)                                                                    #screen the regular message
         cat("\nADVICE:  Now selecting", toupper(ItemName2), "\b. ENTER numbers and/or 1 optional letter (1 space between items):-\n") 
         ConstrMessage <- NULL                                                                               #remove previous constrMessage, ready for the next announcement
                                                        
#8.1.1 Deal with keyboard input to Constraint menu
         ConstrMenuChoice.In <- as.character(readLines(n=1))                                                 #read response as integers and characters 
         ConstrMenuChoice.In <- gsub("  ", " ", ConstrMenuChoice.In)                                         #correct any double spaces to single spaces (does not work for 3 spaces)
         ConstrMenuChoice.In <- unlist(strsplit(ConstrMenuChoice.In, split=" "))                             #split the string into separate choices as characters using the single blank
         N.ConstrMenuChoices <- length(ConstrMenuChoice.In)                                                  #the number of menu items chosen
         N.requestedRows.OS <- N.requestedRows.OS + N.ConstrMenuChoices                                      #accumulated number of menu items chosen, including empties
         MenuOption <- 1                                                                                     #now dealing with first option
         if(N.ConstrMenuChoices == 0)                                                                        #[ENTER] => zero length string meaning return to Main menu
            {MainMessage <- "You have moved back to MAIN MENU from Constraints Menu.  "
            break()                                                                                          #break out of while(ConstrCount < 100) loop back to main menu
            }
         for(ConstrMenuChoice in ConstrMenuChoice.In)                                                        #For each item in the response,
            {if(ConstrMenuChoice %in% AllowedConstrResponses == FALSE)                                       #If character is not assigned, [see #7.7]
               {ConstrMessage1 <- c("Input unrecognised.  Please enter 1 or more numbers and spaces, 0 or 1 letter to process, from\n ", AllowedConstrResponses, "\nor [Enter] or [ESC].") #advise
               next()                                                                                        #advise and return to Constraint menu
               }
            if(ConstrMenuChoice == "0")                                                                      #0 => toggle through output variable types: Active / Slacks / Limits
               {if(ItemN == 1) {ItemN <- 2; ItemName2 <- "Slack variables"} else                             #If ItemN is currently 1, corresponding to active constraints, change to 2, else
                  {if(ItemN == 2) {ItemN <- 3; ItemName2 <- "Constraint limits"} else                        #If ItemN is currently 2, corresponding to slack variables; change to 3, else
                     {if(ItemN == 3) {ItemN <- 1; ItemName2 <- "Activity markers"}                           #If ItemN is currently 3, corresponding with constraint limits; change back to 1
                     }
                  }
               ConstrMessage1 <- "The type of constraint variable being selected has changed."
               ConstrMessage <- paste(ConstrMessage, ConstrMessage1, sep="\nADVICE: ")                       #concatenate the new message with any already existing, with line break between
               ConstrMessage1 <- NULL                                                                        #clear CM1
               MenuOption <- MenuOption + 1                                                                  #increment count of options chosen in one line of response to menu
               next()                                                                                        #break out of while(ConstrCount < 100) loop back to main menu
               }

#8.1.2 Selected option from Constraints' menu was numerical 
#Note: logic for selecting constraints for output differs from that for selecting variables (see next menu).  Here, required row numbers in All.C, the constraint matrix, are collected in OutputRowVec
            if(ConstrMenuChoice %in% NumericConstrOptions)                                                   #IF A NUMERICAL OPTION IS CHOSEN:-
               {OutputRowVec <- NULL                                                                         #clear OutputRowVec with NULL (elements can be concatenated); ORV is a vector for constraint row numbers
               if(ConstrMenuChoice == "1") OutputRowVec <- 1:N.C                                             #select all constraint row numbers
               if(ConstrMenuChoice == "2") OutputRowVec <- GP.a.R                                            #select autotrophic GP row numbers
               if(ConstrMenuChoice == "3") OutputRowVec <- m.pGP.g.R                                         #consumer pGP row numbers
               if(ConstrMenuChoice == "4" & !(is.null(M.SATE.h.R))) OutputRowVec <- M.SATE.h.R               #If Gmax supplied, select hetero GP (sated) row numbers; if Gmax not supplied, M.SATE.h.R=NA
               if(ConstrMenuChoice == "5" & !(is.null(M.HARV.h.R))) OutputRowVec <- M.HARV.h.R               #If Hmax supplied, select hetero harvest row numbers, 
               if(ConstrMenuChoice == "6") OutputRowVec <- m.FOOD.G.R                                        #select FOOD row numbers
               if(ConstrMenuChoice == "7" & !(is.null(d.FOOD.S.R))) OutputRowVec <- d.FOOD.S.R               #select NLO FOOD row number(s)
               if(ConstrMenuChoice == "8" & !(is.null(n.NUTR.A.R))) OutputRowVec <- n.NUTR.A.R               #If nutrients are in the model, select nutrient rows for autotrophs 
               if(ConstrMenuChoice == "9" & !(is.null(n.EXCR.h.R))) OutputRowVec <- n.EXCR.h.R               #If nutrients are in the modelselect nutrient rows for heterotrophs
               if(ConstrMenuChoice == "10" & !(is.null(m.pGP.g.R)) & N.Fisheries > 0) OutputRowVec <- m.pGP.g.R[(N.FoodHetero.pairs + 1):(N.FoodConsumer.pairs)]#If fisheries are in the model, select landings' rows 
               if(ConstrMenuChoice == "11" & !(is.null(m.QUOTA.f.R))) OutputRowVec <- m.QUOTA.f.R            #If quota are set, select quota-constraint row numbers of fisheries constrained 
               if(ConstrMenuChoice == "12" & !(is.null(z.HAB.W.R))) OutputRowVec <- z.HAB.W.R                #If habitats are in the model, select habitat-size row numbers, 
               if(ConstrMenuChoice == "13")                                                                  #If ALL-constraints-on-autotrophs is requested:-
                  {OutputRowVec <- GP.a.R                                                                    #select auto GP constraints; they always exist because they are part of the minimal model
                  for(aa in 1:N.Autos)                                                                       #For each autotroph,
                     {OutputRowVec <- c(OutputRowVec, grep(Autotrophs[aa], row.names(All.C)[m.FOOD.G.R]) + m.FOOD.G.R[1] - 1) #find all food constraints with an autotroph in the code name using grep(); concats with ORVec
                     }
                  if(!(is.null(n.NUTR.A.R))) OutputRowVec <- c(OutputRowVec, n.NUTR.A.R)                     #If nutr constr on autos exist (probably the is.null() is not needed), put into selec vec
                  if(!(is.null(z.HAB.W.R)))                                                                  #If a z.HAB.W constraint is present:-
                     {for(hab in 1:N.Habitats)                                                               #For each one,
                        {if(any(z.HAB.W[hab, Auto.Obj] > 0)) OutputRowVec <- c(OutputRowVec, z.HAB.W.R[hab]) #If growth of any autotroph is constrained by this habitat, concat the row number of this constraint
                        }     
                     }
                  }
               if(ConstrMenuChoice == "14")                                                                  #If ALL-constraints-on-heterotrophs is selected; they always exist because they are part of the minimal model:-
                  {OutputRowVec <- m.pGP.g.R                                                                 #add rows of all m.pGP.g constraints applying to consumers
                  Fery.R <- NULL                                                                             #prepare empty list of fishery rows; they are not heterotrophs and should be removed here
                  if(N.Fisheries > 0)                                                                        #If fisheries are in the model:-
                     {for(hh in 1:N.Fisheries)                                                               #for each fishery,
                        {Fery.R <- c(Fery.R, grep(Fisheries[hh], row.names(All.C)[m.pGP.g.R]) + m.pGP.g.R[1] - 1) #calculate row #s having this fishery's name using grep() and concatenate new rows in Fery.R
                        }
                     OutputRowVec <- OutputRowVec[!is.na(ifelse(OutputRowVec %in% Fery.R, NA, OutputRowVec))]#remove the m.pGP.Fishery rows
                     }  
                     
                  OutputRowVec <- c(OutputRowVec, m.FOOD.G.R)                                                #select m.FOOD.G constraints on GPs of heteros; they always exist
                  if(!is.null(n.EXCR.h.R)) OutputRowVec <- c(OutputRowVec, n.EXCR.h.R)                       #If nutrient constr on heteros exist, select them even if redundant
                  if(!is.null(z.HAB.W.R))                                                                    #If habitat constraints on heteros exist, select them even if redundant
                     {for(hab in 1:N.Habitats)                                                               #For each one,
                        {if(any(z.HAB.W[hab, Hetero.Obj] > 0)) OutputRowVec <- c(OutputRowVec, z.HAB.W.R[hab])#if growth of any heterotroph is constrained by this habitat, concat the row number of this constraint
                        }     
                     }
                  }
               if(ConstrMenuChoice == "15")                                                                  #If all-constraints-on-fisheries is selected:-
                  {if(N.Fisheries > 0)                                                                       #If fisheries are in the model:-
                     {for(ff in 1:N.Fisheries)                                                               #For each fishery,
                         {OutputRowVec <- c(OutputRowVec, grep(Fisheries[ff], row.names(All.C)))             #select row.names including this fishery (always as consumer) with grep() and concatenate
                         }
                     }
                  }
               #PROCESS THE NUMERICAL OPTION SELECTED
               if(is.null(OutputRowVec))                                                                     #IF OUTPUTROWVEC IS STILL NULL, ie. selected constraints are unfound and no row numbers have been added:- 
                  {ConstrMessage1 <- paste0("Option ", paste0(ConstrMenuChoice, ": "), "Requested constraints are all missing, have no information, or were never active. No time series were added.\n\t" ) #start new ConstrMessage
                  } else                                                                                     #Else IF OUTPUTROWVEC IS NOT NULL, ie. selected constraints have been found and row numbers have been added:- 
                  {N.rows.TS <- length(OutputRowVec)                                                         #the number of rows of constraints now selected, now stored in Temp.set
                  if(ItemN == 1)                                                                             #IF ACTIVITY MARKERS ARE REQUIRED (see #8.1.1), convert slack variables to activity markers:-
                     {Temp.set <- array(data=Slacks[OutputRowVec, 1:(M+1)], dim=c(N.rows.TS, M+1), dimnames=list(paste0("Activ.", row.names(All.C)[OutputRowVec]), names(SimTime)[1:(M+1)])) #Temp.set holds slacks
                     for (iii in 1:N.rows.TS)                                                                #For each row of Temp.set, 
                        {Temp.set[iii, ] <- ifelse(Temp.set[iii, ] == 0, 1, NA)                              #If Slack=0 insert 1, else insert NA (meaning that constraint is not binding, not plotted)
                        }
                     }
                  
                  if(ItemN == 2)                                                                             #IF OUTPUT OF SLACK VARIABLES IS REQUIRED (see #8.1.1)
                     {Temp.set <- array(data=Slacks[OutputRowVec, 1:(M+1)], dim=c(N.rows.TS, M+1), dimnames=list(row.names(Slacks)[OutputRowVec], names(SimTime)[1:(M+1)])) #array to hold slacks
                     }

                  if(ItemN == 3)                                                                             #IF OUTPUT OF CONSTRAINT LIMITS IS REQUIRED (see #8.1.1)
                     {Temp.set <- array(data=Limits[OutputRowVec, 1:(M+1)], dim=c(N.rows.TS, M+1), dimnames=list(row.names(Limits)[OutputRowVec], names(SimTime)[1:(M+1)])) #array to hold limits
                     }
                  Temp.set[ , M+1] <- NA                                                                     #set last value to NA so as not plotted as a dip to 0 (Slacks & Limits are stored at M, not at M+1)
                  is.rowEmpty <- rep(TRUE, N.rows.TS)                                                        #create marker of empty rows, ie. holding 0s and NAs only, in Temp.set
                  for(TS.R in 1:N.rows.TS)                                                                   #For each row of Temp.set,
                     {if(ItemN == 1) {if(sum(Temp.set[TS.R, ], na.rm=TRUE) != 0) is.rowEmpty[TS.R] <- FALSE} #activity marks are valid if not zero or NA
                     if(ItemN > 1) {if(any(!is.na(Temp.set[TS.R, ]))) is.rowEmpty[TS.R] <- FALSE}            #slack vars and limits are valid if not NA
                     }
                  if(sum(is.rowEmpty) == N.rows.TS)                                                          #IF THE NUMBER OF EMPTY ROWS = THE NUMBER OF ROWS IN TEMP.SET, ie. all are empty:-
                     {ConstrMessage1 <- paste0("Option ", ConstrMenuChoice, "; Requested constraints are ALL missing, have no information, or were never active. No time series were added.")
                     }   

                  if(sum(is.rowEmpty) < N.rows.TS)                                                           #IF THE NUMBER OF EMPTY ROWS < THE NUMBER OF ROWS IN TEMP.SET, ie. some useful rows are present:-
                     {if(DeleteEmptySeries == FALSE)                                                         #IF EMPTY ROWS ARE TO BE RETAINED IN THE OUTPUT.SET (MainMenu option 6):-
                        {if(ItemN == 1)                                                                      #If Activity markers are required, change the 1s marking activity to integers to space constr up the ordinate
                           {for (iii in 1:N.rows.TS)                                                         #For each row of Temp.set,
                              {nextMark <- iii * Ordinate.factor                                             #step active constr markers up ordinate by O.f (set in #7.7); O.f=0 -> nS is unchanged -> markers at same ordin. 
                              Temp.set[iii, ] <- ifelse(Temp.set[iii, ] == 1, nextStep + nextMark, NA)       #If mark=1 insert iii*Ord.fac instead.  If mark is NA or 0, insert NA (not binding, not plotted)  
                              }
                           nextStep <- nextStep + nextMark                                                   #increment nextStep in case other activity marks are selected 
                           }   
                        if(is.Output.set == FALSE)                                                           #IF OUTPUT.SET DOES NOT YET EXIST:-
                           {Output.set <- array(data=Temp.set, dim=c(N.rows.TS, M+1), dimnames=list(row.names(Temp.set), names(SimTime)[1:(M+1)])) #create Output.set with all rows of Temp.set
                           is.Output.set <- TRUE 
                           } else                                                                            #else IF OUTPUT.SET DOES EXIST ALREADY:-
                           {Output.set <- array(data=rbind(Output.set, Temp.set), dim=c(N.rows.OS + N.rows.TS, M+1), dimnames=list(c(row.names(Output.set), row.names(Temp.set)), names(SimTime)[1:(M+1)]))#bind on Temp.set
#                           if(length(grep(ItemShortName, unlist(strsplit(Ordinate.label, ", ")))) == 0) Ordinate.label <- paste0(Ordinate.label, ", ", ItemShortName) #concat. short name if not already present in Ordinate.label
                           }
                        ConstrMessage1 <- paste0(paste0("Option ", ConstrMenuChoice, "; "), paste0(N.rows.TS, " time series added to Output.set, including " ), paste0(sum(is.rowEmpty), " empty time series. [E=Edit removes; MainMenu 6 auto-deletes]"))
                        } else                                                                               #else IF EMPTY ROWS ARE TO BE DELETED FROM THE OUTPUT.SET:-
                        {Temp.set <- array(data=Temp.set[!is.rowEmpty, ], dim=c(N.rows.TS - sum(is.rowEmpty), M+1), dimnames=list(row.names(Temp.set)[!is.rowEmpty], names(SimTime)[1:(M+1)])) #create T.set with non-empty rows
                        N.rows.TS <- N.rows.TS - sum(is.rowEmpty)                                            #update N rows in Temp.set to exclude empties
                       if(ItemN == 1)                                                                        #change the 1s marking constraint activity to integers that will space constr up the ordinate
                           {for (iii in 1:N.rows.TS)                                                         #For each row of Temp.set,
                              {nextMark <- iii * Ordinate.factor                                             #step active constr markers up ordinate by O.f; O.f  can be changed to 0 -> nS is unchanged -> markers at same ordin. 
                              Temp.set[iii, ] <- ifelse(Temp.set[iii, ] == 1, nextStep + nextMark, NA)       #If mark=1 insert iii*Ord.fac instead.  If mark is NA or 0, insert NA (not binding, not plotted)  
                              }
                           nextStep <- nextStep + nextMark                                                   #increment nextStep in case other activity marks are selected
                           }
                        if(is.Output.set == FALSE)                                                           #IF OUTPUT.SET DOES NOT YET EXIST:-
                           {Output.set <- array(data=Temp.set, dim=c(N.rows.TS, M+1), dimnames=list(row.names(Temp.set), names(SimTime)[1:(M+1)])) #create Output.set with non-empty rows
                           is.Output.set <- TRUE
#                           Ordinate.label <- ItemShortName                                                  #the ordinate label for graphs is created with the short name                    
                           } else                                                                            #else IF OUTPUT.SET DOES EXIST:-
                           {Output.set <- array(data=rbind(Output.set, Temp.set), dim=c(N.rows.OS + N.rows.TS, M+1), dimnames=list(c(row.names(Output.set), row.names(Temp.set)), names(SimTime)[1:(M+1)])) #bind on non-empty rows
#                           if(length(grep(ItemShortName, unlist(strsplit(Ordinate.label, ", ")))) == 0) Ordinate.label <- paste0(Ordinate.label, ", ", ItemShortName) #concat. short name if not already present in Ordinate.label
                           }
                        ConstrMessage1 <- paste0(paste0("Option ", ConstrMenuChoice, ": "), paste0(sum(!is.rowEmpty), " time series added to Output.set; ", sum(is.rowEmpty), " empty series deleted. [MainMenu 6 stops auto-deletion.]" ))                       
                        }
                     N.rows.OS <- N.rows.OS + N.rows.TS                                                      #current number of rows in Output.set
                     }                                                                                       #end of if(sum(is.rowEmpty) < N.rows.TS) block
                  }                                                                                          #end of if(!is.null(OutputRowVec)) block
               }                                                                                             #end of If numerical option was chosen from Constraints menu                  

#8.1.3 Selected option from Constraints' menu was alphabetic
            if(ConstrMenuChoice %in% c("P","p","G","g"))                                                     #If printing or graphing is requested:-
               {if(is.Output.set == TRUE)                                                                    #and If the Output.set exists:-
                  {ConstrMessage1 <- ProcessOutput(ConstrMenuChoice, ItemName, Ordinate.label, N.rows.OS, TitleText, TitleTime, KeyAuto, N.keyLines, MaxOrdinate, MinYearOut, MaxYearOut) #print or graph output; return message to say so [ProcessOutput() is in ECOLPSfunctions.R
                  if(ConstrMenuChoice == "G" | ConstrMenuChoice == "g") Graph.plotted <- TRUE                #remember that a graph has been plotted (in case a text label will be applied)
                  } else
                  {ConstrMessage1 <- "No Output.set exists to print or graph:"
                  Graph.plotted <- FALSE                                                                     #remember that a graph has not been plotted (in case a text label will be requested)
                  }
               }
            if(ConstrMenuChoice %in% c("A","a"))                                                             #If adjustmetn of the ordinate spacing of Active constraints up the ordinate is requested:-
               {#Note: analysis of peaks/valleys ("A") is only available for Variables, not for activity markers. For them, "A" adjusts the ordinate spacing of Active constraints on a graph
               if(ItemN == 1)                                                                                #If active constraints are being selected:-
                  {cat("\nEnter the factor used to space the next constraint-activity bars up the graph ordinate, currently '", Ordinate.factor, "'.  Default=1 :- \nEnter a non-negative integer=>\n")
                  InputNumber <- readLines(n=1)                                                              #read keystrokes of user as characters
                  if(all(unlist(strsplit(InputNumber,split=character(0))) %in% c("","0","1","2","3","4","5","6","7","8","9"))) #check that all characters are +ve integers. No easier way found!
                     {if(InputNumber=="")                                                                    #If typed input was just [Enter]:-
                        {Ordinate.factor <- 1                                                                #default value = 1
                        } else                                                                               #else If typed input exists:-
                        {Ordinate.factor <- as.numeric(InputNumber) #read keyboard input                     #read the keyboard input
                        }
                     ConstrMessage1 <- paste0("\nOrdinate.factor has been changed to ", Ordinate.factor, ":  ") #notify changed ord.factor             
                     } else
                     {ConstrMessage1 <- "\nUnexpected input. Ordinate.factor is unchanged:  "                #notify of unacceptable typed input
                     }
                  } else ConstrMessage1 <- "\nThe 'A' option only has effect when ACTIVE constraints are being selected:  "  #notify that A does not work for slacks or limites
               }
            if(ConstrMenuChoice %in% c("M", "m"))                                                            #If totalling of the columns of the Output.set is required,
               {ColSums <- array(data=Csums(Output.set, N.rows.OS, M+1), dim=c(1, M+1), dimnames=list("Col.totals", rep(NA, M+1)))
               Output.set <- rbind(Output.set, ColSums)                                                      #bind on totals as final row of existing Output.set (use E to remove)
               N.rows.OS <- N.rows.OS + 1                                                                    #correct the number of rows
               ConstrMessage1 <- "Final row of Output.set now has totals of existing columns"
               }
            if(ConstrMenuChoice %in% c("N", "n"))                                                            #If If counts of positive values in the Output.set is required,
               {ColNumbs <- Csums(Output.set > 0, N.rows.OS, M+1)
               Output.set <- rbind(Output.set, array(data=ColNumbs, dim=c(1, M+1), dimnames=list("Col.counts of +ves", rep(NA, M+1))))  #bind on counts as final row of existing Output.set (use E to remove)
               N.rows.OS <- N.rows.OS + 1                                                                    #correct the number of rows
               ConstrMessage1 <- "Final row of Output.set now has COUNTS of positive values in existing columns"
               }
            if(ConstrMenuChoice %in% c("E","e","T","t","R","r","C","c","B","b","F","f","I","i","L","l","J","j","K","k","O","o","W","w", "X","x","Y","y","S","s","Q","q", "z", "Z", "?"))#If adjustment of the Output.set is requested:-
               {tempList <- EditOutput(ConstrMenuChoice, ItemName, Ordinate.label, N.rows.OS, KeyAuto, MinOrdinate, MaxOrdinate)    #adjust Output.set with (E), (T), etc options
               #EditOutput returns (list(1 Message, 2 Temp.set, 3 is.Output.set, 4 TitleText, 5 StartOut, 6 EndOut, 7 OutputDeltas, 8 Thin, 9 WinWidth, 10 WinHeight, 11 N.keyLines, 12 ItemName, 13 Ordinate.label, 14 TimeLabel, 15 AxLabelFactor, 16 Col.plots, 17 Graph.plotted, 18 MaxOrdinate, 19 TitleTime, 20 PlotSymbolSize, 21 KeyAuto, 22 GraphLabel, 23 Jitter, 24 MaxYearOut, 25 MinYearOut))        #25 items are returned in an R list read with [[ ]]
               ConstrMessage1 <- tempList[[1]]                                                                         #feedback message for next menu
               if(ConstrMenuChoice %in% c("E","e")) Output.set <- as.array(tempList[[2]])                              #revise Output.set, only changed by the edit options
               if(ConstrMenuChoice %in% c("C","c")) {is.Output.set <- tempList[[3]]; Ordinate.label <- ItemName <- tempList[[12]]; N.ActiveConstraintsSelected <- 0; Ordinate.factor <- 1; nextStep <- 0; N.requestedRows.OS <- 0}                                                                                                                            #revise is.Output.set
               if(ConstrMenuChoice %in% c("T", "t")) {TitleText <- tempList[[4]]; TitleTime <- tempList[[19]]}         #revise TitleText; TitleTime (switch for adding time to graph titles)
               if(ConstrMenuChoice %in% c("B", "b")) {OutputDeltas <- tempList[[7]]; MinYearOut=tempList[[25]]} #revise StartOut, OutputDeltas, etc 
               if(ConstrMenuChoice %in% c("F", "f")) {EndOut <- tempList[[6]]; OutputDeltas <- tempList[[7]]; MaxYearOut=tempList[[24]]} #revise EndOut, OutputDeltas, etc
               if(ConstrMenuChoice %in% c("I", "i")) {TitleText <- tempList[[4]]; OutputDeltas <- tempList[[7]]; Thin <- tempList[[8]]} #revise Title text to show thinning, OutputDeltas, Thin
               if(ConstrMenuChoice %in% c("W", "w")) {WinWidth <- tempList[[9]]; WinHeight <- tempList[[10]]}          #revise graphic window width, height
               if(ConstrMenuChoice %in% c("K", "k")) {N.keyLines <- tempList[[11]]; KeyAuto <- tempList[[21]]}         #revise N key lines on graphs; autokey switch
               if(ConstrMenuChoice %in% c("Y", "y")) {Ordinate.label <- tempList[[13]]}                                #revise label for the vertical axis of graphs
               if(ConstrMenuChoice %in% c("X", "x")) {TimeLabel <- tempList[[14]]}                                     #revise label for the time axis of graphs
               if(ConstrMenuChoice %in% c("S", "s")) {AxLabelFactor <- tempList[[15]]}                                 #revise axis label and numbering size factor 
               if(ConstrMenuChoice %in% c("Q", "q")) {Col.plots <- tempList[[16]]}                                     #revise Col.plots switch, 0=B&W, 1=Coloured symbols  
               if(ConstrMenuChoice %in% c("R", "r")) {Graph.plotted <- FALSE}                                          #revise Graph.plotted switch for Label in EditOutput()
               if(ConstrMenuChoice %in% c("O", "o")) {MaxOrdinate <- tempList[[18]]}                                   #revise maximum graph ordinate -> common axis for multiple plots, or v.high values omitted -> more detail
               if(ConstrMenuChoice %in% c("Z", "z")) {PlotSymbolSize <- tempList[[20]]}                                #revise plot-symbol size with cex parameter (0.5 to 1.5)
               if(ConstrMenuChoice %in% c("L", "l")) {GraphLabel <- tempList[[22]]}                                    #update GraphLabel for use with labelling option, default="Time"
               if(ConstrMenuChoice %in% c("J", "j")) {Jitter <- tempList[[23]]; TitleText <- tempList[[4]]}            #adjust jitter factor for plotted points, default=0 ; adjust title text to show jittering              
               N.OutputDeltas <- length(OutputDeltas)                                                                  #revise N elements in OutputDeltas for output given begin, finish, and intervals between points
               if(is.Output.set == TRUE) {N.rows.OS <- nrow2(Output.set, M + 1)} else {N.rows.OS <- 0}                 #update the number of rows now in Output.set, incl. if only 1          
               }
            ConstrMessage1 <- paste0("\nADVICE: ", ConstrMessage1)
            ConstrMessage <- paste0(ConstrMessage, ConstrMessage1)                                                     #concatenate new message with any already existing, line break between
            ConstrMessage1 <- NULL                                           
            MenuOption <- MenuOption + 1                                                                               #increment count of options chosen in one line of response to menu
            }                                                                                                          #end of for(ConstrMenuChoice in ConstrMenuChoice.In) loop
         ConstrCount <- ConstrCount + 1                                                                                #increment count of visits to the constraints menu
         }                                                                                                             #end of while(ConstrCount < 100) loop
      }                                                                                                                #end of if(MainMenuChoice == 1) block   
#----------------------------------------------------------------


#8.2 Select SIMULATED VARIABLES for output (MainMenuChoice == "2")
   #SUBSUBSECTIONS
      #8.2.0 Variables menu
      #8.2.1 Deal with keyboard input to Variables menu
      #8.2.2 Selection from Variables' menu was numerical
      #8.2.3 Selection from Variables' menu was alphabetical
#Note: if Variables-menu options are changed, AllowedVarsResponses in #6.7 must also be changed   

#8.2.0 Variables menu
   if(MainMenuChoice == "2")
      {ItemName <- "Sim.Vars"                                                                                          #default name for heading printouts etc
      ItemName2 <- "Simulated variables"                                                                               #another name
      ItemN <- 4                                                                                                       #used by EditOutput() in ECOLPSfunctions.R, choice Y (change ordinate label)
      VarsMessage <- paste0("\nADVICE: You have moved to VARIABLES MENU from Main Menu (via option ", MainMenuChoice, ").") #this message opens the variables menu for the first time
      if(VarsCount == 0) Ordinate.label <- ItemName                                                                    #label for TS graph ordinate
      VarsCount <- 0                                                                                                   #zero the variables menu loop counter for this approach to the menu
      while(VarsCount < 100)                                                                                           #count of repeat visits to variables menu; end of this while-loop is in #8.2.3
         {RepeatedVarsMessage <- paste0("\bADVICE: OUTPUT.SET now holds ", ifelse(is.Output.set == FALSE, 0, N.rows.OS), " time series from ", Years[StartOut], " to ", Years[EndOut], " years thinned by ", Thin, " points")
         cat("\n\nVARIABLES MENU. Options for ", M, " simulation steps stored as time series.") 
         cat("
ORGANIC MASSES:  1 -> Wilds, NLO        2 -> Autotrophs          3 -> Grazers, Predators 4 -> Scavengers & NLO    5 -> Trophic groups   
GAINs by LP:     11-> Components        12-> Autotrophs          13-> partial Consumers  14-> Scavengers & NLO    15-> Grouped       
HARVESTS by LP:  21-> from Wilds, NLO   22-> by Consumers        23-> partial            24-> Dropped harvests    25-> Grouped harvests
NUTRIENTS:       31-> Nmasses in H2O  32-> Nmasses in Wilds  33-> Nmasses in NLO  34-> Nmasses in Ecosystem  35-> Nmasses Landed  36-> NutrConc in NLO  
PHYSICAL:        41-> TEMPERATURE index 42-> LIGHT index         43-> HABITAT unfilled **** 37-> CumNmasses Landed    38 -> 34 + 37
FISHERIES by LP: 51-> partial Landings  52-> partial Catches     53-> partial Discarded  54-> Quota remaining     55-> Fishing effort
by PROJECTION:   61-> Self losses       62-> Total losses(21+61) 63-> Harvests(unassim.) 64-> D1.Bmass  65-> D2.Bmass   66-> D3.Bmass (not used)
PARAMETERS ETC   71-> SelfGrow adj. by Light(&Temp.)   72-> Harvest adjusted by Temp.    
A -> ANALYSE peaks, dips, Output.set      B -> BEGIN time of Output      C -> CLEAR current Output.set     E -> EDIT rows of Output.set        
F -> FINISH time of Output                G -> GRAPH current Output.set  I -> Time INTERVAL between points J -> JITTER points of next plot
K -> Number of KEY lines on graphs        L -> LABEL the ACTIVE graph    M -> Column TOTALS                N -> NUMBER of column posiives       
O -> max ORDINATE                         P -> PRINT current Output.set  Q -> COLOUR/B&W graph symbols     R -> REMOVE active GRAPH          
S -> Size of axes' labelling              T -> TITLE of output           W -> GraphWINDOW width,height     X or Y -> Label X or Y axis    
Z -> plot SYMBOL size                     [ESC] -> TERMINATE simulation (variables remain in memory)       [Enter] -> Return to MAIN MENU \n")
         cat(VarsMessage)                                                 #screen the new message
         cat("\n", RepeatedVarsMessage)                                   #screen the regular message
         cat("\nADVICE:  Now selecting", toupper(ItemName2), "\b. ENTER numbers and/or 1 optional letter (1 space between items):-\n") 
#print(VarsMessage); print(RepeatedVarsMessage)
         VarsMessage <- NULL                                              #remove previous VarsMessage, ready for the next announcement

#8.2.1 Deal with keyboard input to Variables menu
         VarsMenuChoice.In <- as.character(readLines(n=1))                                                             #read response as a sequence of integers and characters 
         VarsMenuChoice.In <- gsub("  ", " ", VarsMenuChoice.In)                                                       #correct any double spaces to single spaces (does not work for 3 spaces)
         VarsMenuChoice.In <- unlist(strsplit(VarsMenuChoice.In, split=" "))                                           #split the string into separate choices as characters using a single space as the separator
         N.VarsMenuChoices <- length(VarsMenuChoice.In)                                                                #number of menu items chosen
         N.requestedRows.OS <- N.requestedRows.OS + N.VarsMenuChoices                                                  #accumulated number of rows chosen for output, including previous choices and any empty rows
         MenuOption <- 1                                                                                               #now dealing with first menu option typed in
         if(length(VarsMenuChoice.In) == 0)                                                                            #[ENTER] => zero length string meaning return to Main menu
            {MainMessage <- "You have moved back to MAIN MENU from Variables Menu"
            break()                                                                                                    #break out of while(VarsCount < 100) loop back to main menu
            }
         for(VarsMenuChoice in VarsMenuChoice.In)                                                                      #For each item in the response,
            {if(VarsMenuChoice %in% AllowedVarsResponses == FALSE)                                                     #If character is not assigned, [see #7.7]:-
               {VarsMessage1 <- c("Input unrecognised.  Please enter 1 or more numbers with space, 0 or 1 letter to process, from\n ", AllowedVarsResponses, "\nor [Enter] or [ESC].") #advise
               next()                                                                                                  #return to Variables menu
               }
            
#8.2.2 Selection from Variables' menu was numerical
#Note: array() statements below permit row names to exist; R removes them if there is only 1 row in Temp.set
#NOTE: if you add a new VarsMenuChoice to those below, you MUST also add it to the list called NumericVarsOptions in #7.7
#Note: logic for selecting variables for output differs from that for selecting constraints (previous menu).  Here, the required time series of simulated variables are stacked in Temp.set

            if(VarsMenuChoice %in% NumericVarsOptions)                                                                 #If a numerical option was chosen:-
               {Temp.set <- NULL                                                                                       #setting NULL permits the first element to be filled; empty rows are filled with with NAs or 0s
               if(VarsMenuChoice == "1") Temp.set <- Bmass[is.WildOrNLO , 1:(M+1)]                                     #all wild and NLO biomasses are selected
               if(VarsMenuChoice == "2") Temp.set <- array(Bmass[is.Auto, 1:(M+1)], dim=c(N.Autos, M+1), dimnames= list(row.names(Bmass)[is.Auto], names(SimTime)[1:(M+1)])) #auto biomasses only
               if(VarsMenuChoice == "3") Temp.set <- array(Bmass[is.Grazer | is.Predator, 1:(M+1)], dim=c(N.Grazers + N.Predators, M+1), dimnames= list(row.names(Bmass)[is.Grazer | is.Predator], names(SimTime)[1:(M+1)])) #grazer and predator biomasses only; this includes scavengers that graze or predate as well as consuming NLO
               if(VarsMenuChoice == "4") if(N.NLOs > 0) Temp.set <- Bmass[is.Scavenger | is.NLO, 1:(M+1)] else Temp.set <- array(data=NA, dim=c(max(N.Scavengers + N.NLOs, 2), M+1), dimnames=list(c("Scavengers", "NLO"), names(SimTime)[1:(M+1)]))                                                                                                   #scavengers and NLO biomasses only
               if(VarsMenuChoice == "5") Temp.set <- TrophicGroups.Bmass[ , 1:(M+1)]                                   #biomasses totalled by trophic category of component
               if(VarsMenuChoice == "11") Temp.set <- Gains[ , 1:(M+1)]                                                #GPs totalled for individual wild and fishery components
               if(VarsMenuChoice == "12") Temp.set <- array(pGains[Autos.RpG, 1:(M+1)], dim=c(N.Autos, M+1), dimnames= list(row.names(pGains)[Autos.RpG], names(SimTime)[1:(M+1)]))  #pGains for individual autos
               if(VarsMenuChoice == "13") Temp.set <- array(pGains[Consumers.RpG, 1:(M+1)], dim=c(N.FoodConsumer.pairs + N.UnAssimHarv.pairs, M+1), dimnames= list(row.names(pGains)[Consumers.RpG], names(SimTime)[1:(M+1)])) #pGains for consumers x foods 
               if(VarsMenuChoice == "14") Temp.set <- array(pGains[c(Scavengers.RpG, NLO.RpG), 1:(M+1)], dim=c(length(c(Scavengers.RpG, NLO.RpG)), M+1), dimnames= list(row.names(pGains)[c(Scavengers.RpG, NLO.RpG)], names(SimTime)[1:(M+1)]))                                                                                                              #pGains for scavengers and NLO
               if(VarsMenuChoice == "15") Temp.set <- Grouped.Gains[ , 1:(M+1)]                                        #gains totalled by grouped components
               if(VarsMenuChoice == "21") Temp.set <- HarvestsFrom[ , 1:(M+1)]                                         #harvests taken from wild and NLO components
               if(VarsMenuChoice == "22") Temp.set <- HarvestsBy[ , 1:(M+1)]                                           #Harvs taken by wild and fishery components       
               if(VarsMenuChoice == "23") Temp.set <- pHarvs[ , 1:(M+1)]                                               #pHarvs for all pairs
               if(VarsMenuChoice == "24") Temp.set <- Dropped.pHarvs[ , 1:(M+1)]                                       #wild dropped harvs
               if(VarsMenuChoice == "25") Temp.set <- rbind(Grouped.HarvestsFrom, Grouped.HarvestsBy)[ , 1:(M+1)]      #grouped harvests
               if(VarsMenuChoice == "31") if(N.Nutr > 0) Temp.set <- array(Nmasses[NinH2O.R , 1:(M+1)], dim=c(N.Nutr, M+1), dimnames= list(row.names(Nmasses)[NinH2O.R], names(SimTime)[1:(M+1)])) else Temp.set <- array(data=NA, dim=c(max(N.H2Os, 1), M+1), dimnames=list("NmassesInH2O", names(SimTime)[1:(M+1)]))                                       #nutrients in water
               if(VarsMenuChoice == "32") if(N.Nutr > 0) Temp.set <- array(Nmasses[NinWilds.R , 1:(M+1)], dim=c(N.Nutr, M+1), dimnames= list(row.names(Nmasses)[NinWilds.R], names(SimTime)[1:(M+1)])) else Temp.set <- array(data=NA, dim=c(1, M+1), dimnames=list("NmassesInWilds", names(SimTime)[1:(M+1)]))                                              #nutrients in Wilds
               if(VarsMenuChoice == "33") if(N.Nutr > 0 & N.NLOs > 0) Temp.set <- array(Nmasses[NinNLO.R , 1:(M+1)], dim=c(N.Nutr, M+1), dimnames= list(row.names(Nmasses)[NinNLO.R], names(SimTime)[1:(M+1)])) else Temp.set <- array(data=NA, dim=c(max(N.NLOs, N.Nutr, 1), M+1), dimnames=list("NmassesInNLO", names(SimTime)[1:(M+1)]))                  #nutrients in NLO 
               if(VarsMenuChoice == "34") if(N.Nutr > 0) Temp.set <- array(Nmasses[NinSys.R, 1:(M+1)], dim=c(N.Nutr, M+1), dimnames= list(row.names(Nmasses)[NinSys.R], names(SimTime)[1:(M+1)])) else Temp.set <- array(data=NA, dim=c(1, M+1), dimnames=list("NmassesInEcoSys", names(SimTime)[1:(M+1)]))                                                   #nutrients in ecosystem
               if(VarsMenuChoice == "35") if(N.Nutr > 0 & N.Fisheries > 0) Temp.set <- array(Nmasses[NinLandings.R, 1:(M+1)], dim=c(N.Nutr, M+1), dimnames= list(row.names(Nmasses)[NinLandings.R], names(SimTime)[1:(M+1)])) else Temp.set <- array(data=NA, dim=c(max(N.Nutr, N.Fisheries, 1), M+1), dimnames=list("NmassesInLandings", names(SimTime)[1:(M+1)])) #nutrients in fishery landings
               if(VarsMenuChoice == "36") if(N.Nutr > 0 & N.NLOs > 0) Temp.set <- array(NconcInNLO[ , 1:(M+1)], dim=c(max(1, N.Nutr), M+1), dimnames= list(row.names(NconcInNLO), names(SimTime)[1:(M+1)])) else Temp.set <- array(data=NA, dim=c(max(N.Nutr, N.NLOs, 1), M+1), dimnames=list("NconcInNLO", names(SimTime)[1:(M+1)]))                         #nutrient concs in NLO
               if(VarsMenuChoice == "37") if(N.Nutr > 0 & N.Fisheries > 0) Temp.set <- array(Nmasses[NinCumLandings.R, 1:(M+1)], dim=c(N.Nutr, M+1), dimnames= list(row.names(Nmasses)[NinCumLandings.R] , names(SimTime)[1:(M+1)])) else Temp.set <- array(data=NA, dim=c(max(N.Nutr, N.Fisheries, 1), M+1), dimnames=list("NmassesInCumLandgs", names(SimTime)[1:(M+1)])) #cumulative nutrs in fishery landings
               if(VarsMenuChoice == "38") 
                  {if(N.Nutr > 0 & N.Fisheries > 0) 
                      {Temp.set <- array(Nmasses[NinSys.R, 1:(M+1)] + Nmasses[NinCumLandings.R, 1:(M+1)], dim=c(N.Nutr, M+1), dimnames= list(paste0(row.names(Nmasses)[NinSys.R], "+CumLand") , names(SimTime)[1:(M+1)]))
                      Temp.set[ , M+1] <- NA                                                                           #set the last value to NA to prevent last point dropping on graph because cumLanded is 0
                      } else 
                      {Temp.set <- array(data=NA, dim=c(max(N.Nutr, N.Fisheries, 1), M+1), dimnames=list("NmassesInEcoSys&CumLandgs", names(SimTime)[1:(M+1)])) #nutrs in ecosystem + fishery landings
                      }
                  }
               if(VarsMenuChoice == "41") Temp.set <- array(T.index[ , 1:(M+1)], dim=c(1, M+1), dimnames= list(row.names(T.index), names(SimTime)[1:(M+1)])) #temperature index
               if(VarsMenuChoice == "42") Temp.set <- array(L.index[ , 1:(M+1)], dim=c(1, M+1), dimnames= list(row.names(L.index), names(SimTime)[1:(M+1)])) #light index
               if(VarsMenuChoice == "43") Temp.set <- array(HabLeft[ , 1:(M+1)], dim=c(max(1, N.Habitats), M+1), dimnames= list(row.names(HabLeft), names(SimTime)[1:(M+1)])) #habitat left
               if(VarsMenuChoice == "51") if(N.Fisheries > 0) Temp.set <- array(pGains[Fisheries.RpG, 1:(M+1)], dim=c(length(Fisheries.RpG), M+1), dimnames= list(row.names(pGains)[Fisheries.RpG], names(SimTime)[1:(M+1)])) else  Temp.set <- array(data=NA, dim=c(1, M+1), dimnames=list("pGain.Fishery", names(SimTime)[1:(M+1)]))                    #pGains for fisheries
               if(VarsMenuChoice == "52") if(N.Fisheries > 0) Temp.set <- array(pHarvs[Fisheries.RpG, 1:(M+1)], dim=c(length(Fisheries.RpG), M+1), dimnames= list(row.names(pHarvs)[Fisheries.RpG], names(SimTime)[1:(M+1)])) else  Temp.set <- array(data=NA, dim=c(1, M+1), dimnames=list("pHarv.Fishery", names(SimTime)[1:(M+1)]))                    #pHarvs for fisheries  
               if(VarsMenuChoice == "53") if(N.Fisheries > 0) Temp.set <- array(Dropped.pHarvs[Fisheries.RpG, 1:(M+1)], dim=c(length(Fisheries.RpG), M+1), dimnames= list(row.names(Dropped.pHarvs)[Fisheries.RpG], names(SimTime)[1:(M+1)]))  else  Temp.set <- array(data=NA, dim=c(1, M+1), dimnames=list("Dropped.pHarvs.Fishery", names(SimTime)[1:(M+1)]))  #dropped pHarvs for fisheries
               if(VarsMenuChoice == "54") if(N.QuotumFishery.pairs > 0) Temp.set <- array(QuotaLeft[ , 1:(M+1)], dim=c(N.QuotumFishery.pairs, M+1), dimnames= list(row.names(QuotaLeft), names(SimTime)[1:(M+1)]))  else  Temp.set <- array(data=NA, dim=c(1, M+1), dimnames=list("QuotaLeft", names(SimTime)[1:(M+1)]))                                    #quota remaining for fisheries
               if(VarsMenuChoice == "55") if(N.Fisheries > 0) Temp.set <- array(F.effort[ , 1:(M+1)], dim=c(max(1, N.Fisheries), M+1), dimnames=list(row.names(F.effort), names(SimTime)[1:(M+1)]))  else  Temp.set <- array(data=NA, dim=c(1, M+1), dimnames=list("Fishing.Effort", names(SimTime)[1:(M+1)]))                                              #fishing effort
               if(VarsMenuChoice == "61") Temp.set <- array(proj.SelfLosses[is.WildOrNLO, 1:(M+1)], dim=c(N.WildOrNLOs, M+1), dimnames=list(row.names(proj.SelfLosses)[is.WildOrNLO], names(SimTime)[1:(M+1)])) #proj. self losses
               if(VarsMenuChoice == "62") Temp.set <- array(TotalLossesByProj[is.WildOrNLO, 1:(M+1)], dim=c(N.WildOrNLOs, M+1), dimnames=list(row.names(TotalLossesByProj)[is.WildOrNLO], names(SimTime)[1:(M+1)]))#total SLosses
               if(VarsMenuChoice == "63") if(N.UnAssimHarv.pairs > 0) Temp.set <- array(pHarvs[projPairs, 1:(M+1)], dim=c(max(1, N.UnAssimHarv.pairs), M+1), dimnames=list(row.names(proj.UnAssimHarvs), names(SimTime)[1:(M+1)]))  else  Temp.set <- array(data=NA, dim=c(max(N.UnAssimHarv.pairs, 1), M+1), dimnames=list("UnAssimHarv.pairs", names(SimTime)[1:(M+1)])) #unassimilated harvest pairs

               if(VarsMenuChoice == "64") Temp.set <- D1.BmassByProj[is.WildOrNLO, 1:(M+1)]                            #projected 1st derivatives of biomasses
               if(VarsMenuChoice == "65") Temp.set <- D2.BmassByProj[is.WildOrNLO, 1:(M+1)]                            #projected 2nd . . .   
               if(VarsMenuChoice == "66") Temp.set <- D3.BmassByProj[is.WildOrNLO, 1:(M+1)]                            #projected 3rd . . .
               if(VarsMenuChoice == "71") Temp.set <- array(SG.LTadj[ , 1:(M+1)], dim=c(nrow(SG.LTadj), M+1), dimnames=list(row.names(SG.LTadj), names(SimTime)[1:(M+1)])) #Varying SelfGrow with L.index
               if(VarsMenuChoice == "72") Temp.set <- array(Hvst.Tadj, dim=c(nrow(Hvst.Tadj), M+1), dimnames=list(row.names(Hvst.Tadj), names(SimTime)[1:(M+1)])) #Varying Harv with T.index

#Note: Code below for Variables menu is simpler than that for Constraints menu because there is only 1 type of variable, whereas constraints have 3 types (Active, Slacks, Limits) that may be output
               N.rows.TS <- nrow2(Temp.set, M+1)                                                                       #N rows in Temp.set, includ if Temp.set is still only NULL. nrow2 works with 1 or more rows [unlike nrow() ]
               is.rowEmpty <- rep(TRUE, N.rows.TS)                                                                     #create marker of empty rows in Temp.set
               if(N.rows.TS > 0)                                                                                       #IF THERE IS AT LEAST 1 ROW IN TEMP.SET:-
                  {for(TS.R in 1:N.rows.TS)                                                                            #For each row of Temp.set,
                     {if(any(!is.na(Temp.set[TS.R, ]) & Temp.set[TS.R, ] != 0)) {is.rowEmpty[TS.R] <- FALSE}           #empty is false for this row
                     }
                  }    
               if(sum(is.rowEmpty) >= N.rows.TS) VarsMessage1 <- paste0("Option ", VarsMenuChoice, "; Requested variables are ALL missing or have no information" ) #message for N empty rows = N rows in TEMP.SET,
              if(DeleteEmptySeries == FALSE)                                                                           #IF EMPTY ROWS ARE TO BE RETAINED IN THE OUTPUT.SET:-
                  {if(is.Output.set == FALSE)                                                                          #and IF OUTPUT.SET DOES NOT YET EXIST:-
                     {Output.set <- array(data=Temp.set, dim=c(N.rows.TS, M+1), dimnames=list(row.names(Temp.set), names(SimTime)[1:(M+1)])) #create Output.set with all rows of Temp.set 
                     is.Output.set <- TRUE                                                                             #O.s now exists
                     #Ordinate.label <- ItemShortName                                                                  #the ordinate label for graphs is created with the short name          
                     } else                                                                                            #else IF OUTPUT.SET DOES EXIST ALREADY:- 
                     {Output.set <- array(data=rbind(Output.set, Temp.set), dim=c(N.rows.OS + N.rows.TS, M+1), dimnames=list(c(row.names(Output.set), row.names(Temp.set)), names(SimTime)[1:(M+1)]))#bind on all rows of Temp.set
                     }
                  VarsMessage1 <- paste0(paste0("Option ", VarsMenuChoice, ": "), paste0(N.rows.TS, " time series added to Output.set, including " ), paste0(sum(is.rowEmpty), " empty time series. [E=Edit removes; MainMenu 6 auto-deletes.]"))                                                                                                          #message for rows added
                  } else                                                                                               #else IF EMPTY ROWS ARE TO BE DELETED FROM THE OUTPUT.SET:-
                  {Temp.set <- array(data=Temp.set[!is.rowEmpty, ], dim=c(N.rows.TS - sum(is.rowEmpty), M+1), dimnames=list(row.names(Temp.set)[!is.rowEmpty], names(SimTime)[1:(M+1)])) #create T.set with non-empty rows
                  N.rows.TS <- N.rows.TS - sum(is.rowEmpty)                                                            #update N rows in Temp.set to exclude empties
#                  Temp.set[ , M+1] <- ifelse(Temp.set[ , M+1] == 0, NA, Temp.set[ , M+1])                             #change last value to NA if 0 to prevent plotting a final drop
                  if(is.Output.set == FALSE)                                                                           #IF OUTPUT.SET DOES NOT YET EXIST:-
                     {Output.set <- Temp.set                                                                           #create Out.set with non-empty rows
                     is.Output.set <- TRUE                                                                             #mark output set as existent
                     } else                                                                                            #else IF OUTPUT.SET DOES EXIST:-  
                     {Output.set <- rbind(Output.set, Temp.set)                                                        #bind on non-empty rows of Temp.set
                     }
                  VarsMessage1 <- paste0(paste0("Option ", VarsMenuChoice, ": "), paste0(sum(!is.rowEmpty), " time series added to Output.set; ", sum(is.rowEmpty), " empty series deleted. [MainMenu 6 stops auto-deletion.]" ))
                  }
               N.rows.OS <- nrow2(Output.set, M+1)                                                                     #N.rows.OS was initially set to 0 in #7.7. Here it is adjusted for Temp.set
            }                                                                                                          #end of If numerical option was chosen from Vars menu                  
 
#8.2.3 Selection from Variables' menu was alphabetical (VarsMenuChoice = "P","p","G","g","A" or "a" . . . )
            if(VarsMenuChoice %in% c("A","a","G","g","P","p"))                                                         #If printing, graphing or analysis of peaks and valleys is requested:-
               {if(is.Output.set == TRUE)                                                                              #and If an Output.set already exists:-
                  {VarsMessage1 <- ProcessOutput(VarsMenuChoice, ItemName, Ordinate.label, N.rows.OS, TitleText, TitleTime, KeyAuto, N.keyLines, MaxOrdinate, MinYearOut, MaxYearOut) #print/graph output; return message to say so
                  if(VarsMenuChoice == "G" | VarsMenuChoice == "g") Graph.plotted <- TRUE                              #remember that a graph has been plotted (in case a text label will be applied)
                  } else                                                                                               #else If an Output.set does not already exist:-
                  {VarsMessage1 <- "No Output.set exists to process.  "          
                  Graph.plotted <- FALSE                                                                               #remember that a graph has not been plotted (in case a text label will be requested)
                  }
               }
            if(VarsMenuChoice %in% c("M", "m"))                                                                        #If totalling of the columns of the Output.set is required,
               {ColSums <- array(data=Csums(Output.set, N.rows.OS, M+1), dim=c(1, M+1), dimnames=list("Col.totals", rep(NA, M+1))) #form the col sums using Csums() (which also works with 1 row)
               Output.set <- rbind(Output.set, ColSums)                                                                #bind on totals as final row of existing Output.set (use E to remove)
               N.rows.OS <- N.rows.OS + 1                                                                              #correct the number of rows
               VarsMessage1 <- "Final row of Output.set now has totals of existing columns"
               }                                                                                                 
            if(VarsMenuChoice %in% c("N", "n"))                                                                        #If counts of positive values in the columns of the Output.set is required,
               {ColNumbs <- Csums(Output.set > 0, N.rows.OS, M+1)                                                      #form the col counts using Csums() (which also works with 1 row)
               Output.set <- rbind(Output.set, array(data=ColNumbs, dim=c(1, M+1), dimnames=list("Col.counts of +ves", rep(NA, M+1))))  #bind on counts as final row of existing Output.set (use E to remove)
               N.rows.OS <- N.rows.OS + 1                                                                              #correct the number of rows
               VarsMessage1 <- "Final row of Output.set now has COUNTS of positive values in existing columns"
               }
            if(VarsMenuChoice %in% c("B","b","C","c","E","e","F","f","I","i","J","j","K","k","L","l","O","o","Q","q","R","r","S","s","T","t","W","w","X","x","Y","y","z","Z","?"))#If adjustment of the Output.set is requested:-
               {tempList <- EditOutput(VarsMenuChoice, ItemName, Ordinate.label, N.rows.OS, KeyAuto, MinOrdinate, MaxOrdinate)      #adjust Output.set with (E), (T), etc options; EditOutput() is in ECOLPSfunctions.R
               #EditOutput returns (list(1 Message, 2 Temp.set, 3 is.Output.set, 4 TitleText, 5 StartOut, 6 EndOut, 7 OutputDeltas, 8 Thin, 9 WinWidth, 10 WinHeight, 11 N.keyLines, 12 ItemName, 13 Ordinate.label, 14 TimeLabel, 15 AxLabelFactor, 16 Col.plots, 17 Graph.plotted, 18 MaxOrdinate, 19 TitleTime, 20 PlotSymbolSize, 21 KeyAuto, 22 GraphLabel, 23 Jitter, 24 MaxYearOut, 25 MinYearOut))        #25 items are returned in an R list read with [[ ]]
               VarsMessage1 <- tempList[[1]]                                                                           #feedback message
               if(VarsMenuChoice %in% c("E","e"))  {Output.set <- as.array(tempList[[2]])}                             #replace complete Output.set
               if(VarsMenuChoice %in% c("C","c"))  {is.Output.set <- tempList[[3]]; Ordinate.label <- ItemName <- tempList[[12]]; Ordinate.factor <- 1; nextStep <- 0; N.requestedRows.OS <- 0} #is.Output.set (TRUE/FALSE)->clear
               if(VarsMenuChoice %in% c("T", "t")) {TitleText <- tempList[[4]]; TitleTime <- tempList[[19]]}           #title text; TitleTime (=toggle data & time in graph titles)
               if(VarsMenuChoice %in% c("B", "b")) {OutputDeltas <- tempList[[7]]; MinYearOut <- tempList[[25]]}       #adjust start of output; adjust OutputDeltas to implement this
               if(VarsMenuChoice %in% c("F", "f")) {EndOut <- tempList[[6]]; OutputDeltas <- tempList[[7]]; MaxYearOut <- tempList[[24]]}   #adjust end point of output;  adjust OutputDeltas to implement this
```
