## Supplementary material for "Virtual aquatic ecology: stepped simulations of gross production within constraints explain biomass changes": ECOLPS simulator in R: ECOLPSfunctions.R

```
#ECOLPSfunctions.R:- Copyright CC BY-SA 2025 by A.John R.Cotter. See https://creativecommons.org/ This program is distributed WITHOUT ANY WARRANTY WHATSOEVER.
#Version.AJRC: 32, last edited 1 December 2025, Version.Zenodo: 1.0.0,   Contact
#This code must be stored in the same folder as ECOLPS.R and Controls.txt for operational use; it is kept seperate and called with source() to avoid cluttering the start of the main program
#Note: these functions can read variables in the main program but only overwrite them if the re-calculated variable (or array, list, etc) is alone as the last code line, or if a return() statement is used. 
#Only 1 variable (or array, list, etc) can be returned. So, if more than 1 is needed, the variable names have to be passed with a list[[var1, var2, , , , ]].  eg. EditOutput() has 20-something returns 
   #CONTENTS - functions called by ECOLPS
      #Advise() <- function(WithRefTo, ErrText, AdvText, ShowData, CallSection, Fatal, MinAdviceLevel) 
      #EditOutput() <- function(Choice, ItemName, Output.label, N.rows, KeyAuto, MaxOrdinate)      
      #findOneLine() <- function(SearchTxt, Headers, call.section, file.txt, Var.name, domain, ZeroOK)
      #is.DatumOK() <- function(Datum, VarName, NA.OK, CharOK, NumbOK, LogicalOK, MustBeFile=NA, MustBeInSet=NA, MustNotBeInSet=NA, ZeroOK=NA, InfinOK=NA, nonIntegerOK=NA, MaxOK=NA, MinOK=NA, MaxNotOK=NA, MinNotOK=NA)
      #nrow2() <- function(VecOrMat, N.cols)
      #ncol2() <- function(VecOrMat, N.rows)
      #PickActiveConstraints() <- function(PickSet, ConstraintBindings)
      #Plot.results() <- function(Plot.matrix, AxLabel, TitleText, N.plotVars, TitleTime, KeyAuto, N.keyLines, MaxOrdinate)
      #ProcessOutput() <- function(Choice, ItemName, N.rows, TitleText, TitleTime, KeyAuto, N.keyLines, MaxOrdinate, MaxYearOut)
      #ReOrder() <- function(Rows, Source, Stem, Suffixes, Primary, PrimaryName, ErrLine, Test)  
      #Restart.TS() <- function(TS.matrix, Last.col, Years)
      #ParamAdj() <- function(L.Index, SlopeSG.L, MidSG.L, SG)
      #ParaAdj.T() <- function(T.Index, Mu.T, Sigma.T, Coeff)
      #ShowCalcs() <- function(MM)
      #Csums() <- function(A, nR, nC)
      #Rsums() <- function(A, nR, nC)
                                  
Advise <- function(WithRefTo, ErrText, AdvText, ShowData, CallSection, Fatal, MinAdviceLevel2Show) {
#Screens error and/or most advisory messages, uniformly, with/without data and locations
if(!is.na(ErrText[1]))cat("\n************************************************************************************************************************************************\n")
#suppressWarnings(
if(!is.na(WithRefTo[1])) cat("\nWITH REFERENCE TO: ", WithRefTo, "\n\n")                                           #cat() writes to screen
if(!is.na(ErrText[1])) cat("ERROR:- ", ErrText, "\n\n")
if(!is.na(AdvText[1]) & AL >= MinAdviceLevel2Show) cat("ADVICE:- ", AdvText, "\n")                                 #AL is the advice level set by Controls.txt
if(!is.null(ShowData))                                                                                             #If any element is present,
   {if(class(ShowData[1]) == "character") {cat(ShowData, "\n")} else {cat("\nRELEVANT DATA:\n") ; print(ShowData)} #show data on screen
   }
if(!is.na(CallSection)) cat("\nAdvice sent from code section ", CallSection)                                       #If source of message was provided, advise the code section number
if(Fatal) stop("\b\b\b\b\b\b\b\nFATAL ERROR; program terminated.", call.=FALSE)                                    #If Fatal=True, stop the program
}
################################################################# End of function Advise


EditOutput <- function(Choice, ItemName, Ordinate.label, N.rows, KeyAuto, MinOrdinate, MaxOrdinate){
#EditOutput() implements tasks requested by alphabetic letters entered in response to Constraints' and Variables' menus.  
#If Output.set exists, it is duplicated as Temp.set and edited here as required, then returned to the calling code
#Choice is the letter entered by user in response to the Constraint or Variables menus; ItemName is used to identify recent data in Output.set; N.rows refers to the number of rows in the Output.set
if(is.Output.set == TRUE) {Temp.set <- Output.set} else {Temp.set <- NA}                                           #If an Output.set exists, make a copy.  Else, O.set <- NA
   #SECTIONS
      #EO1 Edit the current Output.set of active constraints or slack variables
      #EO2 Check or change the title of output
      #EO3 Close all graphics windows
      #EO4 Clear existing Output.set
      #EO5 Remove clearance of existing Output.set
      #EO6 Adjust the beginning time of printed and graphic outputs 
      #EO7 Adjust the finishing time of printed and graphic outputs      
      #EO8 Adjust interval between time-series points in printed and graphic output; this is 'thinning'
      #EO9 Label the active plot with text additional to the title
      #EO10 Change the size of graphic windows
      #EO11 Toggle automatic/manual setting of the number of key lines on graphs
      #EO12 Change the Time label in graphic output    
      #EO13 Change the label for Variables in output
      #EO14 Change the size of axes' labels relative to the plot character
      #EO15 Switch for plotting graphic symbols in black and white only, or colour
      #EO16 Fix the maximum ordinate
      #EO17 Alter the size of plot symbols
      #EO18 Change Jitter for plotted points
#----------------------------------------------------------------


#EO1 Edit the current Output.set of active constraints or slack variables
if(Choice == "E" | Choice == "e")                                                                                  #If editing of Output.set is selected:-
   {if(is.Output.set == TRUE)                                                                                      #and If the Output.set exists,
      {for(ii in 1:N.rows)                                                                                         #For each row of Output.set,
         {cat("\n", format(ii, width=3), format(row.names(Temp.set)[ii], width=15), Temp.set[ii, 1:min(10, M)])    #display 1st cols of the row with some formatting
         }
      cat("\n\nFirst 10 columns of the current Output.set of", ItemName, "are shown above to assist identification of variables.  \nSelect ROW NUMBERS (one at a time) to be omitted from graphic or printed output.  [Enter] (by itself) terminates.\n\n")                                                                                         #advise
      Delete.rows <- suppressWarnings(as.numeric(scan(nmax=(N.rows - 1), what=character())))                       #read keyboard input with scan(), leaving min 1 row; should stop a fatal error if a nonnumeric is typed
      Delete.rows <- Delete.rows[which(Delete.rows >= 1 & Delete.rows <= N.rows)]                                  #limit to existing rows
      if(length(Delete.rows) > 0)                                                                                  #If the number of rows to be deleted is positive:-
         {cat("\nRows to be omitted are:\n")                                                                       #advise
         for(i in 1:length(Delete.rows))                                                                           #For each row, 
            {cat("\n", format(Delete.rows[i], width=3), format(row.names(Temp.set)[Delete.rows[i]], width=25), Temp.set[Delete.rows[i], 1:min(10, M)]) #display all rows to be deleted
            }
         cat("\nAre these right?  [Enter] = YES, they are all right; OR enter the displayed row numbers (one at a time) to be undeleted.\n") #allow user a chance to amend mistakes
         Undelete.rows <- suppressWarnings(as.numeric(scan(nmax=length(Delete.rows), what=character())))           #read rows to be undeleted
         if(length(Undelete.rows) > 0)                                                                             #If the number is positive,
            {Delete.rows <- Delete.rows[which(Delete.rows != Undelete.rows)]                                       #remove from the rows to be deleted
            }
         if(length(Delete.rows) > 0)                                                                               #If a positive number of rows is to be deleted:-
            {if(DeleteEmptySeries == TRUE)                                                                         #and If option 6 of Main Menu says that empty series should be deleted:-
               {Temp.set <- array(data=Temp.set[-Delete.rows, ], dim=c(N.rows - length(Delete.rows), M + 1), dimnames=list(row.names(Temp.set)[-Delete.rows], names(Years)[Start.M:(Start.M + M)])) #remove them from Temp.set; redefining an array to do this is intended to prevent R discarding the row and column structure of Output.set
               } else                                                                                              #else If option 6 says no delete:- 
               {Temp.set[Delete.rows, ] <- NA
               }                                                                                                   #fill with NAs, eg. as row place holders for matching TS symbols across graphs
            }
         }
      Message <- paste0(length(Delete.rows), " rows have been deleted from the Output.set")  
      } else                                                                                                       #else If is.Output.set==FALSE:-
      {Message <- "No editing is possible"                                                                         # message to be shown under next Menu
      }                    
   }                                                                                                               #end of if(ConstrMenuChoice == E) block                          
#----------------------------------------------------------------


#EO2 Check or change the title of output
if(Choice == "T" | Choice == "t")                                                                                  #If the title of output is to be adjusted:-
   {cat("\nEnter the title for printed and graphic output, currently '", TitleText, "':- \nText => REPLACE current title with text; @ => toggle inclusion of date & time; * => default title; 0 => NO main title; NO text => KEEP current title\nENTER one option =>\n")
   InputText <- as.character(readLines(n=1))                                                                       #read keystrokes of user as characters
   if(!is.null(InputText) & !is.na(InputText) & InputText != "")                                                   #If input text is not null or invalid:-
      {if(InputText == "*")                                                                                        #If # was typed:-
         {TitleText <- DefaultTitleText; Message <- paste("Graphs and tables will have the default title '", DefaultTitleText, "'", sep="")  #restore the default title; see MainProgram #7.7
         } else
         {if(InputText == "0")                                                                                     #If 0 was typed:-
            {TitleText <- ""; TitleTime <- FALSE; Message <- "Graphs and tables will have no title.  "  
            } else 
            {if(InputText == "@")                                                                                  #If * was typed:-
               {TitleTime <- !TitleTime                                                                            #toggle the TitleTime switch (adds time to title if true)
               if(TitleTime == TRUE) 
                  {Message <- "The title of graphs and tables WILL include the date and time of printing"
                  } else 
                  {Message <- "The title of graphs and tables will NOT include the date and time of printing"
                  }
               } else
               {TitleText <- InputText                                                                             #else If text was typed in:-
               Message <- paste0("The title of outputs has been changed to '", TitleText, "'")                     #input text => new title
               }
            }
         }
      } else   
      {Message <- paste0("Unexpected or null input. The title of outputs '", TitleText, "' is unchanged")          #message to be shown under next Menu
      } 
   }
#----------------------------------------------------------------


#EO3 Close the active graphics window
if(Choice == "R" | Choice == "r")                                                                  #If the active graph window is to be closed:-
   {act.dev <- dev.cur()                                                                           #get the graphic-device i.d. of the active device
   if(act.dev > 1)                                                                                 #If the active device is higher than 1:-
      {dev.off()                                                                                   #close it; if this was the last, R sets the first g.d as active; otherwise R sets the first in the circular list of g.ds
      if(act.dev > 2) dev.set(dev.prev())                                                          #adjusts R's choice to the g.d one before the closed device but stops a blank g.d being screened when g.d is closed
      Message <- paste0("Graphic window ", act.dev, " has been closed")                            #message to be shown under next Menu
      } else
      {Message <- "Request to close graphic window 1 (the 'null device')   cannot be implemented"
      }
   }
#----------------------------------------------------------------


#EO4 Clear existing Output.set
if(Choice == "C" | Choice == "c")                                                                  #If clearance of the current Output.set is selected:-
   {is.Output.set <- FALSE                                                                         #reset the existence switch to signal that Output.set is ready to be overwritten
   Message <- "The Output.set has been cleared"                                                    #advice for screening under the next menu
   }
#----------------------------------------------------------------

            
#EO5 Remove clearance of existing Output.set
if(Choice == "U" | Choice == "u")                                                                  #If Output.set was cleared (option above) in error:-
   {if(nrow2(Output.set, N.OutputDeltas) > 0)                                                      #and If there is at least 1 row in Output.set:-
      {is.Output.set <- TRUE                                                                       #reverse the existence switch
      Message <- "The previous Output.set has been undeleted"                                      #prepare advice
      } else
      {Message <- "No Output.set has been found to undelete"                                       #message to be shown under next Menu
      }   
   }
#----------------------------------------------------------------

   
#EO6 Adjust the beginning time of printed and graphic outputs 
if(Choice == "B" | Choice == "b")                                                                  #If the start time of output is to be reset (does not adjust Output.set itself):-        
   {cat("\nENTER a value in years at which to BEGIN all printed and graphic output of time series, currently ", MinYearOut, " years.\n")
   StartOutYear.tmp <- as.numeric(readLines(n=1))                                                  #read from keyboard input
   if(StartOutYear.tmp < Years[1]) StartOutYear.tmp <- Years[1]                                    #Default correction of user: MinYearOut cannot be less than the year of the first data point 
   ERR <- is.DatumOK(StartOutYear.tmp, "New startOutput year", NA.OK=FALSE, CharOK=FALSE, NumbOK=TRUE, LogicalOK=FALSE, MinOK=0, MaxNotOK=Years[EndOut])#check supplied values
   if(ERR == TRUE)                                                                                 #If an invalid value is found:-
      {Message <- paste0("\nADVICE:- Unaccepted or null input. First time of printed and graphic output is unchanged from: ", MinYearOut) #message to be shown under next Menu
#                    1        2          3              4          5          6            7       8      9        10        11          12        13               14            15            16         17           18      
      return(list(Message, Temp.set, is.Output.set, TitleText, StartOut, EndOut, OutputDeltas, Thin, WinWidth, WinHeight, N.keyLines, ItemName, Ordinate.label, TimeLabel, AxLabelFactor, Col.plots, Graph.plotted, MaxOrdinate, TitleTime, PlotSymbolSize, KeyAuto, GraphLabel, Jitter, MaxYearOut, MinYearOut))                  #Return after erroneous input; 25 items must be returned by EditOutput()
#   19           20          21          22       23         24         25
      }
   MinYearOut <- StartOutYear.tmp                                                                  #store the new MinYearOut
   MinDeltaOut <- (MinYearOut - Years[1]) * YearLength +1                                          #store the equivalent number of Deltas
   OutputDeltas <- seq(from=MinDeltaOut, to=OutputDeltas[MaxYearOut * YearLength + 1], by=PrintOut.interval) #vector to select time cols from result matrices to print. OutputDeltas holds row numbers not years (differs by +1)
   Message <- paste0("Printed and graphic outputs have been adjusted to begin from ", MinYearOut, " years")  #message to be shown under next Menu
   }
#----------------------------------------------------------------


#EO7 Adjust the finishing time of printed and graphic outputs      
if(Choice == "F" | Choice == "f")                                                                  #If the end time of output is to be reset (does not adjust Output.set itself):-        
   {cat("\nENTER a value in years at which to FINISH all printed and graphic output of time series, currently ", MaxYearOut, " years.\n")
   EndOutYear.tmp <- as.numeric(readLines(n=1))                                                    #read from keyboard input
   if(EndOutYear.tmp > Years[EndOut]) EndOutYear.tmp <- Years[EndOut]                              #Default correction of user: MaxYearOut cannot be more than the year of the last data point 
   ERR <- is.DatumOK(EndOutYear.tmp, "New endOutput year", NA.OK=FALSE, CharOK=FALSE, NumbOK=TRUE, LogicalOK=FALSE, MinNotOK=0, MaxOK=Years[EndOut])#check supplied values
   if(ERR == TRUE)                                                                                 #If an invalid value is found:-
      {Message <- paste0("\nADVICE:- Unaccepted or null input. Last time of printed and graphic output is unchanged from: ", MaxYearOut) #message to be shown under next Menu
#                         1        2          3              4          5          6            7       8      9        10        11          12        13               14            15            16         17           18      
      return(list(Message, Temp.set, is.Output.set, TitleText, StartOut, EndOut, OutputDeltas, Thin, WinWidth, WinHeight, N.keyLines, ItemName, Ordinate.label, TimeLabel, AxLabelFactor, Col.plots, Graph.plotted, MaxOrdinate, TitleTime, PlotSymbolSize, KeyAuto, GraphLabel, Jitter, MaxYearOut, MinYearOut))                  #Return after erroneous input; 25 items must be returned by EditOutput()
   #  19           20          21          22       23         24         25
      }
   MaxYearOut <- EndOutYear.tmp                                                                    #store the new MaxYearOut
   MaxDeltaOut <- (MaxYearOut - Years[1]) * YearLength +1                                          #store the equivalent number of Deltas
   OutputDeltas <- seq(from=OutputDeltas[1], to=MaxDeltaOut, by=PrintOut.interval)                 #vector to select time cols from time-related result matrices to print. OutputDeltas holds row numbers not years (differs by +1)
   Message <- paste0("Printed and graphic outputs have been adjusted to finish at: ", MaxYearOut, " years")
   }                                                                         
#----------------------------------------------------------------


#EO8 Adjust interval between time-series points in printed and graphic output; this is 'thinning'
#Thinning of output points may be useful for identifying the plot symbol or otherwise for clarity.  Note that peaks may become lower because the top point is omitted; vice versa for dips
#OutputDeltas is a vector created to mark the columns from time-related output matrices destined for printing or graphing
if(Choice == "I" | Choice == "i")                                                                  #If the interval between output points is to be reset (does not adjust Output.set itself)
   {cat("\nENTER the number of simulated results to be removed between each printed or graphed result in order to improve clarity, currently ", Thin, " .  [0 shows all].\n")
   Thin.tmp <- as.numeric(readLines(n=1))                                                          #read keyboard input of how many points to omit between shown points
   if(is.numeric(Thin.tmp) & !is.null(Thin.tmp) & Thin.tmp >= 0 & !is.na(Thin.tmp) & Thin.tmp <= (EndOut - StartOut - 1) & Thin.tmp != "")  #If input is valid:-
      {Thin <- min(Thin.tmp, EndOut - StartOut)                                                    #store Thin.tmp
      PrintOut.interval <- Thin + 1                                                                #Thin=0 gives sequence 1,2,3,... as expected
      OutputDeltas <- seq(from=StartOut, to=EndOut, by=PrintOut.interval)                          #vector to select cols from time-related matrices of results to print
      Message <- paste("Printed and graphic outputs will have ", Thin, " point(s) removed between consecutive displayed points")  
      firstBracket <- regexpr("[[]", TitleText)                                                    #adjust graph title text to indicate thinning: find any open left square bracket already present in TitleText
      if(firstBracket > 0)                                                                         #If a bracket is present:-
         {TitleText <- substr(TitleText, 1, firstBracket - 1)                                      #remove the last characters including the bracket
         }
      if(Thin > 0) TitleText <- paste(TitleText, "  [thin ", Thin, " pts]", sep="")                #append the thinning text to the title (removable using the T menu response)
      } else                                                                                       #If input is not valid:-
      {Message <- paste0("Unexpected or null input. Current thinning of time series is unchanged from: ", Thin, "points") #message to be shown under next Menu
      }
   }
#----------------------------------------------------------------
   
#EO9 Label the active plot with text additional to the title
#This can be useful for graphs to go into reports, or just to record what you set for the run.  Label only works on R's "active" plots, ie. the last one plotted
if(Choice == "L" | Choice == "l")                                                                                      #If a label is requested:-
   {if(Graph.plotted == FALSE)                                                                                         #and If no active graph exists:-
      {Message <- "ADVICE: No graph yet plotted for this simulation run -> no labelling"                               #explain why the labelling command has been ignored
      } else                                                                                                           #else If a graph does exist to accept a label:-
      {x.coord <- Years[EndOut] * 0.9 ; y.coord <- 0.9 * min(max(Temp.set[  , OutputDeltas], na.rm=TRUE), MaxOrdinate) #default coordinates put label at top right of graph, allowing for reduced MaxOrd by "O" option
      font.size <- 1.5                                                                                                 #default font size
      cat("\nEnter a LABEL to apply on this graph, currently '", GraphLabel, "':- \nText => REPLACE current label with text; * or time => time of plot (default); date => date of plot; d&t => date and time; file => input parameters file; 0 => abandon labelling; NO text => KEEP current label\nENTER one options =>\n")
      keystrokes <- ""                                                                                                 #prime with no character
      keystrokes <- readLines(n=1)                                                                                     #read keyboard input for new text in place of default label
#copy<-keystrokes
#      
      if(keystrokes == "*" | tolower(keystrokes) == "time")                                                            #If graph label is time of plotting,
         {GraphLabel <- paste0("@ ", substr(as.character(Sys.time()), 12, 19))                                         
         } else                                                                                                        #else if not,
         {if(tolower(keystrokes) == "date") 
             {GraphLabel <- paste0("on ", paste0(substr(date(), 5, 10), " ", substr(date(), 21, 24)))                  #If label text is the date of plotting
             } else                                                                                                    #else if not,
             {if(tolower(keystrokes) == "d&t" ) 
                 {GraphLabel <- date()                                                                                 #If d&t was typed, label is date and time of plotting
                 } else                                                                                                #else if not,
                 {if(tolower(keystrokes) == "file") 
                     {GraphLabel <- Params.file                                                                        #If file was typed, label is the input parameters file name
                     }  else
                     {if(keystrokes != "")                                                                             #If keystrokes is not blank, ie. not the default label,
                         {GraphLabel <- keystrokes                                                                     #accept the provided text
                         }
                     }
                 }
             }
         }        
      char1 <- strsplit(keystrokes, split=character(0))[[1]][1]                                                        #isolate 1st character of keystrokes; needed for identifying 'no action required'
      char1 <- ifelse(is.na(char1), "", char1)                                                                         #adjust an NA
      cat("\nEnter x and y CO-ORDINATES (may be negative), and font size for the label on 3 lines, 1 item per line OR just [Enter] => default value:- ", x.coord, ", ", y.coord, ", ", font.size, " >\n" ) 
      keystrokes <- suppressWarnings(as.numeric(readLines(n=3)))                                                       #read keystrokes of user as vector of 3 numeric vars over 3 lines; suppress warnings if not
      keystrokes <- ifelse(is.na(keystrokes), "", keystrokes)                                                          #NA or blank input leaves keystrokes unchanged
      if(keystrokes[1] != "") x.coord <- as.numeric(keystrokes[1])                                                     #If a number key was pressed, characters entered replace default coordinates
      if(keystrokes[2] != "") y.coord <- as.numeric(keystrokes[2])
      if(keystrokes[3] != "") font.size <- as.numeric(keystrokes[3])                                                   #adjust font size
      if(char1 != "0")                                                                                                 #If the 1st character of keystrokes is not zero:-
         {text(x=x.coord, y=y.coord, labels=GraphLabel, cex=font.size, xpd=TRUE, pos=2)                                #label graph; xpd=T => text outside plot region; pos is code for justification
         Message <- paste0("Graph has been labelled with ", GraphLabel)                                                #advice
         } else                                                                                                        #or If unexpected text was typed:-
         {Message <- "Unexpected or null input. The graph has not been labelled"                                       # message to be shown under next Menu
         }
      }                                                                                                                #end of if(Graph.plotted == TRUE)            
   }   
#----------------------------------------------------------------


#EO10 Change the size of graphic windows
#Graphic windows may be adjusted for your screen size and current requirements
if(Choice == "W" | Choice == "w")                                                                  #If adjustments to the screen window for simulated results are selced:-
   {cat("\nENTER the WIDTH to be set for graphic windows, currently, ", WinWidth, ". Maximum is ", MaxWinWidth, "\n") 
   WinWidth.tmp <- as.numeric(readLines(n=1))                                                      #read from from keyboard input
   if(is.numeric(WinWidth.tmp) & !is.null(WinWidth.tmp) & WinWidth.tmp >= 5 & !is.na(WinWidth.tmp) & WinWidth.tmp != "")  #If input is not null or invalid,
      {WinWidth <- min(WinWidth.tmp, MaxWinWidth)                                                  #store the new WinWidth
      Message1 <- paste0("Graphic window width is now set to ", WinWidth)                          #advise
      } else                                                                                       #or If input is invalid:-
      {Message1 <- "Unexpected or null input for width of graphic window => no change"             #advise
      }
   cat("\nENTER the HEIGHT to be set for graphic windows, currently, ", WinHeight, ". Maximum is ", MaxWinHeight, "\n")  
   WinHeight.tmp <- as.numeric(readLines(n=1))                                                     #read characters from keyboard input 
   print(WinHeight.tmp)     
   if(is.numeric(WinHeight.tmp) & !is.null(WinHeight.tmp) & WinHeight.tmp >= 2 & !is.na(WinHeight.tmp) & WinHeight.tmp <= MaxWinHeight & WinHeight.tmp != "")  #If input is not null or invalid,
      {WinHeight <- min(WinHeight.tmp, MaxWinHeight)                                               #store the new WinHeight
      Message2 <- paste0("Graphic window height is now set to ", WinHeight)                        #advixe
      } else                                                                                       #else if input is invalid,
      {Message2 <- "Unexpected or null input for height of graphic window => no change"  #advise   #advice
      }
   Message <- c(Message1, Message2)                                                                #return 2 messages as a vector (R only returns one item from a function); message to be shown under next Menu
   }            
#----------------------------------------------------------------

   
#EO11 Toggle automatic/manual setting of the number of key lines on graphs
#For general use, the automatic setting is probably OK. Otherwise, adjust keylines to give the best balance between the graph height and the readability of key items, given the width of the graph 
if(Choice == "K" | Choice == "k")                                                                  #If the number of keylines shown at the base of graphs is to be adjusted:-
   {maxKeyLines <- 20                                                                              #this is a sensible maximum; a different value may be appropriate for your screen
   cat("Set the number of KEY lines (including 1 for 'Symbols') at the foot of graphs manually or automatically.  Currently auto setting is ", KeyAuto, ".  \nWhen automatic, min is 0, max is ", maxKeyLines, "lines. [Enter] -> switch on automatic setting OR:  N [Enter] -> fix at N lines => \n")
   N.keyLines.tmp <- as.numeric(readLines(n=1))                                                    #read from keyboard input
   if(!is.na(N.keyLines.tmp) & is.numeric(N.keyLines.tmp) & N.keyLines.tmp >= 0 & N.keyLines.tmp <= maxKeyLines)  #If input is not null or invalid, ie. a valid number:-
      {N.keyLines <- N.keyLines.tmp                                                                #transfer keyboard input to N.keyLines
      KeyAuto <- FALSE                                                                             #mark manual setting
      Message <- paste("The number of key lines in graphic windows is now fixed at ", N.keyLines)  #advise
      } else
      {KeyAuto <- TRUE                                                                             #mark automatic setting
      N.keyLines <- 4                                                                              #value is only used for returning; the value used is calculated when the graph is plotted         
      Message <- "The number of key lines in graphic windows is now chosen automatically"          #message to be shown under next Menu
      }
   } 
#----------------------------------------------------------------

   
#EO12 Change the Time label in graphic output
#The abscissa or horizontal axis of graphic output is always time. Default axis label is TimeLabel.def set in #7.7 of the main ECOLPS program, eg. "Deltas"
if(Choice == "X" | Choice == "x")
   {cat("\nChange the TIME label for graphic output, currently '", TimeLabel, "'. \n[Enter] => KEEP current label; Text => replace current label with TEXT; * => replace with default; 0 => NO label. \nENTER response:-\n")
   TimeLabel.tmp <- as.character(readLines(n=1))                                                   #read new axis label from keyboard input
   if(!is.null(TimeLabel.tmp) & !is.na(TimeLabel.tmp) & TimeLabel.tmp != "")                       #If input is not null or invalid:-
      {if(TimeLabel.tmp == "0")                                                                    #If 0 was typed:-
         {TimeLabel <- ""                                                                          #time label is blank
         Message <- "The Time axis in graphic output will have no label"                                     
         } else                                                                                    #else If something else:-
         {if(TimeLabel.tmp == "*")                                                                 #If * was typed:-
            {TimeLabel <- TimeLabel.def; Message <- paste0("The Time axis in graphic output will be labelled with the default:")  #use \U0394 in quotes for the Delta symbol; TimeLabel.def is from #7.7 in main program
            } else                                                                                 #If text was input:-  
            {TimeLabel <- TimeLabel.tmp; Message <- paste("The label for the Time axis in graphic output has been changed to: '", TimeLabel, "'", sep="")  #input text => new label
            }
         }
      } else                                                                                       #else If input was invalid:-
      {Message <- paste0("Unexpected or null input. Existing Time label for graphic output is unchanged from: ", TimeLabel) #advise message to be shown under next Menu
      } 
   } 
#----------------------------------------------------------------


#EO13 Change the label for Variables in output
#The ordinate or vertical axis of graphic output is always used to scale the variables plotted. The name supplied automatically may be inappropriate, eg. "Simulated variables"
if(Choice == "Y" | Choice == "y")                                                                  #If the ordinate axis is to be re-labelled:-
   {cat("\nChange the label for Variables in output, currently '", Ordinate.label, "'. \n[Enter] => KEEP current label; Text => replace current label with TEXT; * => replace with the default label; 0 => NO label. \nENTER response:-\n")
   Ordinate.label.tmp <- as.character(readLines(n=1))                                              #read new axis label from keyboard input
   if(!is.null(Ordinate.label.tmp) & !is.na(Ordinate.label.tmp) & Ordinate.label.tmp != "")        #If input is valid:-
      {if(Ordinate.label.tmp == "0")                                                               #If 0 was typed:-
         {Ordinate.label <- ""; Message <- "Variables in output will have no label."  
         } else 
         {if(Ordinate.label.tmp == "*")                                                            #If * was typed:-
            {if(ItemN == 1) Ordinate.label <- "Active constraints"                                 #ItemN is set in #8.1 of main ECOLPS program
            if(ItemN == 2) Ordinate.label <- "Slack variables"
            if(ItemN == 3) Ordinate.label <- "Constraint limits"
            if(ItemN == 4) Ordinate.label <- "Simulated variables"
            Message <- paste0("Variables in output will be labelled with the default '", Ordinate.label, "'")
            } else
            {Ordinate.label <- Ordinate.label.tmp; Message <- paste("The label for variables in output has been changed to: '", Ordinate.label, "'", sep="")  #if none of the previous options apply, input text => new label
            } 
         }
      } else                                                                                       #else If input is invalid:-
      {Message <- paste0("Unexpected or null input. Existing label for variables in output is unchanged from: ", ItemName) #advise message to be shown under next Menu
      } 
   } 
#----------------------------------------------------------------
   
   
#EO14 Change the size of axes' labels relative to the plot characters
#Useful for formatting graphs for publications
if(Choice == "S" | Choice == "s")
   {Default.AxLabel.Factor <- 1.5                                                                  #default size of axis label characters is 1.5 x plot character size
   cat("\nChange the size of axes labels relative to the plot characters using a factor, currently ", AxLabelFactor, "\n[Enter] => KEEP current factor; 0.0 to 3.0 => NEW factor; * => Default=1.5. \nENTER response:-\n") 
   AxLabelFactor.tmp <- readLines(n=1)                                                             #read factor from keyboard input; non-numeric => NA
   if(!is.null(AxLabelFactor.tmp) & !is.na(AxLabelFactor.tmp) & AxLabelFactor.tmp != "")           #If input is not null, NA or blank:-
      {if(AxLabelFactor.tmp == "*")                                                                #If input is *  :-
         {AxLabelFactor <- Default.AxLabel.Factor                                                  #use default, in code above
         Message <- paste0("Axis-label factor for graphic axes will be the default size: ", AxLabelFactor)
         }
      if(AxLabelFactor.tmp %in% seq(0, 3, by=0.1))                                                 #if input is from 0.1, 0.2, 0.3 . . . 3.0 :-
         {AxLabelFactor <- as.numeric(AxLabelFactor.tmp)                                           #use the given value
         Message <- paste0("Axis-label factor for graphic axes has been changed to: ", AxLabelFactor)
         }
      } else                                                                                       #else If input is invalid:-
      {Message <- paste0("Unexpected or null input. Existing Axis-label factor is unchanged from: ", AxLabelFactor) #advise message to be shown under next Menu 
      }
   }   
#----------------------------------------------------------------


#EO15 Switch for plotting graphic symbols in black and white only, or colour
#Useful for printing graphs when colour reproduction is not available
if(Choice == "Q" | Choice == "q")
   {cat("\nColour or black-and-white symbols on graphs?  Currently ", Col.plots, "\n[Enter] => KEEP current switch value, 0 => B&W, 1 => Coloured symbols. \nENTER response:-\n")
   Col.plots.tmp <- as.numeric(readLines(n=1))                                                     #read switch from keyboard input; non-numeric => NA      
   if(!is.null(Col.plots.tmp) | !is.na(Col.plots.tmp) | Col.plots.tmp != "")                       #If input is not null or invalid:-
      {Col.plots <- Col.plots.tmp                                                                  #transfer keyboard input to Col.plots
      Message <- paste0("Col.plots switch for graphic axes has been changed to ", Col.plots, " (0=B&W, 1=Colour)")
      } else                                                                                       #else If input is invalid:-
      {Message <- paste0("Unexpected or null input. Existing Col.plots switch for symbol colour is unchanged from: ", Col.plots) #advise message to be shown under next Menu
      }
   }
#----------------------------------------------------------------


#EO16 Fix the maximum ordinate
#useful for side-by-side comparisons of graphs with the same scaling, and to remove high ordinates that shrink the scale for smaller points of more interest
if(Choice == "O" | Choice == "o")
   {cat("\nSet the maximum ordinate for time series, currently ", MaxOrdinate, "\n[Enter] => current maximum; a number > 0 => fixed maximum; Inf => no maximum. \nENTER response:-\n")
   MaxOrdinate.tmp <- as.numeric(readLines(n=1))                                                   #read value from keyboard input; non-numeric -> NA
   if(length(MaxOrdinate.tmp) > 0 & !is.na(MaxOrdinate.tmp) & MaxOrdinate.tmp > 0)                 #If input is not null or invalid:-
      {MaxOrdinate <- MaxOrdinate.tmp                                                              #transfer keyboard input to MaxOrdinate
      Message <- paste0("Maximum ordinate value for graphs has been changed to ", MaxOrdinate)
      } else                                                                                       #else If input is invalid:-
      {Message <- paste0("Unexpected or null input. Existing maximum ordinate value is unchanged from: ", MaxOrdinate) #advise message to be shown under next Menu 
      }
   }   


#EO17 Alter the size of plot symbols
#Bigger plot symbols, possibly with thinning of plotted points (option "I"), may improve presentation of graphs
if(Choice == "Z" | Choice == "z")
   {Default.PlotSymbolSize <- 0.8                                                                  #default plot symbol size; change the value if required
   ValidResponse <- FALSE
   cat("\nChange the size of plot characters, currently set to ", PlotSymbolSize, "\n[Enter] => KEEP current factor;  0.5 to 2.0 => NEW plot-symbol size; * => Default=0.8. \nENTER response:-\n") 
   PlotSymbolSize.tmp <- readLines(n=1)                                                            #read size from keyboard input; non-numeric => NA
   if(!is.null(PlotSymbolSize.tmp) & !is.na(PlotSymbolSize.tmp) & PlotSymbolSize.tmp != "")        #If keyboard input is valid:-
      {if(PlotSymbolSize.tmp == "*")                                                               #If input is *
         {PlotSymbolSize <- Default.PlotSymbolSize                                                 #use the default size, set in code above
         Message <- paste0("Plot-symbol size will be the default: ", Default.PlotSymbolSize)  #
         ValidResponse <- TRUE                                                                     #remember that response was valid
         }
      if(PlotSymbolSize.tmp %in% seq(0.5, 2, by=0.01))                                             #If input is a value in 0.50, 0.51, 0.52, . . . 2.00
         {PlotSymbolSize <- as.numeric(PlotSymbolSize.tmp)                                         #store the value
         Message <- paste0("Plot-symbol size for graphic axes has been changed to: ", PlotSymbolSize)
         ValidResponse <- TRUE                                                                     #remember that response was valid
         }
      }
   if(ValidResponse == FALSE)                                                                      #else If keyboard input is invalid:-
      {Message <- paste0("Unexpected or null input - existing Plot-symbol size is unchanged from: ", PlotSymbolSize) #advise message to be shown under next Menu
      }
   } 
   
#EO18 Change Jitter for plotted points
#Jittering of plotted points assists visibility of overlapping series.  The jittered points are not stored with the Output.set so there is no loss of accuracy in subsequent plots or printouts
#Jittering is effected in ProcessOutput
if(Choice == "J" | Choice == "j")
   {ValidResponse <- FALSE
   cat("\nChange the random jitter factor for plotted points, currently set to ", Jitter, "\n[Enter] => KEEP current factor;  0.005, 0.010,... < 0.2 => NEW jitter value; * or 0 => Default=0. \nENTER response:-\n") 
   Jitter.tmp <- readLines(n=1)                                                                    #read value from keyboard input; non-numeric => NA
   if(!is.null(Jitter.tmp) & !is.na(Jitter.tmp) & Jitter.tmp != "")                                #If keyboard input is valid,
      {if(Jitter.tmp == "*" | Jitter.tmp == "0")                                                   #If input is * or 0 as character
         {Jitter <- 0                                                                              #use the default size, set in code above
         Message <- "Jitter factor will be 0: "                                                    #message to be shown on screen
         ValidResponse <- TRUE                                                                     #remember that response was valid
         if(length(grep(", Jittered", TitleText)) > 0) TitleText <-  gsub(", Jittered", "", TitleText) #overwrite mention of jitter with blank in graph title
         }
      valids <- seq(0.005, 0.2, by=0.005)   
      if(as.numeric(Jitter.tmp) %in% valids)                                                       #If input converted to a numeric is a value in 0.005, 0.01, 0.015, . . . 0.2
         {Jitter <- as.numeric(Jitter.tmp)                                                         #store the value as a numeric
         Message <- paste0("Jitter factor for plotted points has been changed to: ", Jitter)
         ValidResponse <- TRUE                                                                     #remember that response was valid
         if(length(grep(", Jittered", TitleText)) == 0) TitleText <- paste0(TitleText, ", Jittered")
         }
      }
   if(ValidResponse == FALSE)                                                                      #else If keyboard input is invalid:-
      {Message <- paste0("Unexpected or null input - existing Jitter factor is unchanged from: ", Jitter) #advise message to be shown under next Menu
      }
   }  
#----------------------------------------------------------------

#             1        2          3              4          5          6            7       8      9        10        11          12        13               14            15            16         17           18           19
return(list(Message, Temp.set, is.Output.set, TitleText, StartOut, EndOut, OutputDeltas, Thin, WinWidth, WinHeight, N.keyLines, ItemName, Ordinate.label, TimeLabel, AxLabelFactor, Col.plots, Graph.plotted, MaxOrdinate, TitleTime, PlotSymbolSize, KeyAuto, GraphLabel, Jitter, MaxYearOut, MinYearOut))                                                        #24 items are returned by EditOutput()
   #20             21          22      23       24           25
}#################################################################End of EditOutput() function


findOneLine <- function(SearchTxt, Headers, call.section, file.txt, Var.name, domain, ZeroOK){                   
#Finds elements of row- or column-vectors (=Headers) that contain SearchTxt.  An error is displayed and the program stopped if > 1 header contains SearchTxt (ie. an ambiguous result). 
#ZeroOK tells whether not finding the search text is an error or not.  Program stops if it is; otherwise returns NA
#domain can be "row" or "column" for use in the error message
LineN <- grep(SearchTxt, ignore.case=T, Headers)                                                   #grep() looks for SearchTxt in the Headers vector.  LineN is a vector of element numbers containing the search text.
AdjLineN <- LineN - 1
if(length(LineN) < 1 & ZeroOK == FALSE | length(LineN) > 1)                                        #If grep() finds no element and ZeroOK is false, or more than one element with SearchTxt is present:-
   {print(Advise(WithRefTo=paste0("Location of '", Var.name, "' in ", domain, " headings of \n\t", file.txt), ErrText=paste0("Zero, or more than 1 ", domain, " header contains the search text: '", SearchTxt, "' .  There should be one, and only one occurrence of this text."), AdvText=paste0("ECOLPS cannot decide which ", domain, " to use. The file must be adjusted. Check the search domain below and SearchTxt found: \n\t"), ShowData=c(Headers, paste0("\n\tLocation:  ", AdjLineN)), CallSection=paste0("findOneLine() called from ", call.section), Fatal=TRUE, MinAdviceLevel=0))                                                                                #either error stops the program
   } 
if(length(LineN) < 1)                                                                              #If grep() finds no element with SearchTxt present, and ZeroOK is TRUE:-
   {LineN <- NA                                                                                    #standardise the result as NA
   }  
LineN                                                                                              #return LineN
}
##################################################################  End of function findOneLine()


is.DatumOK <- function(Datum, VarName, NA.OK, CharOK, NumbOK, LogicalOK, MustBeFile=NA, MustBeInSet=NA, MustNotBeInSet=NA, ZeroOK=NA, InfinOK=NA, nonIntegerOK=NA, MaxOK=NA, MinOK=NA, MaxNotOK=NA, MinNotOK=NA){
#Used for validating data input in files or by keyboard; values that do not match the requirements of the storage variable are flagged as ERR=TRUE 
#Detects classes and values of Datum not complying with rules given as arguments to this function.  ERR is returned for further actions if needed.  The first 6 arguments are REQUIRED.  Args 3 onwards are logical variables.  
#This function responds to the combination of arguments; eg. to test for an invalid integer, you must set NA.OK=F, CharOK=F, NumbOK=T, LogicalOK=F, nonIntegerOK=F.  This function only handles NULL, zero-length values, NA, character, numeric or logical values. One NA may be in a MustBeInSet but Datum=NA will not be marked as ERR=FALSE unless NA.OK=TRUE. NULL values and zero-length data always create an error.  
ERR <- FALSE; Msg <- NULL                                                                          #Default value of ERR is false, ie. no problem.
if(is.null(Datum) | length(Datum)==0)                                                              #If Datum is NULL or of length zero:-
   {ERR <- TRUE                                                                                    #set ERR and message
#   cat("\n\nis.datumOK?  NO: A supplied value, ", VarName, "= ", Datum, ", breaks a rule for this variable:-")  #these statements may be required but may in the way somewhere.  If needed, remove #
#   cat("\nRule: A NULL or zero-length value is not allowed.\n")
   return(ERR)                                                                                     #leave this function now(because tests below may fail with Datum=NULL);
   }                                                                                               #At this point, Datum is not NULL and has positive length
if(is.na(Datum))                                                                                   #If Datum is NA:-
   {if(NA.OK == FALSE)                                                                             #and, if NA is not allowed:-
      {ERR <- TRUE                                                                                 #set error flag
#      cat("\n\nis.datumOK?  NO: A supplied value, ", VarName, "= ", Datum, ", breaks a rule for this variable:-")
#      cat("\nRule: A value of NA (not available) is not allowed.\n")
      return(ERR)                                                                                  #leave this function now(because tests below may fail with Datum=NA)
      } else                                                                                       #else If NA is allowed:-
      {return(ERR)                                                                                 #return ERR=FALSE; leave this function now(because tests below may fail with Datum=NA even if it is allowed) 
      }  
   }                                                                                               #at this point, Datum is not NULL, has positive length, and is not NA
if(is.character(Datum))                                                                            #If Datum is a character:-
   {if(CharOK==FALSE)                                                                              #and If CharOK (required argument) is false:-
      {ERR <- TRUE; Msg <- "A character value is not allowed."                                     #set ERR and message
      } else                                                                                       #else if Datum is a character and is allowed to be:- 
      {if(!is.na(MustBeFile))                                                                      #If MustBeFile is supplied as an argument:-
         {if(MustBeFile==TRUE & !file.exists(Datum))                                               #and if MustBeFile is true but the file does not exist:-
            {ERR <- TRUE; Msg <- "The value must be the address of an existing file."              #set ERR and message
            }                                                             
         }                                                                
      if(!is.na(MustBeInSet[1]))                                                                   #If MustBeInSet is supplied as an argument:-
         {if(!is.element(Datum, MustBeInSet))                                                      #and If Datum is not an element of the supplied set:-
            {ERR <- TRUE; Msg <- c("The value must be a member of the set:-   {", MustBeInSet, "}")#set ERR and message
            }                                                             
         }                                                                
      if(!is.na(MustNotBeInSet[1]))                                                                #If MustNotBeInSet is supplied as an argument:-
         {if(is.element(Datum, MustNotBeInSet))                                                    #and If Datum is an element of the supplied set:-
            {ERR <- TRUE; Msg <- c("The value must NOT match any member of the set:-   {", MustNotBeInSet, "}") #set ERR and message
            }
         }                                                             
      if(LogicalOK==FALSE & (Datum=="TRUE" | Datum=="FALSE"))                                      #If a logical variable is found but is not allowed:-
         {ERR <- TRUE; Msg <- "The value must not be one of the logical variables:'TRUE' or 'FALSE'"  #set and message
         }
      }                                                                
   }                                                                
if(is.numeric(Datum))                                                                              #If Datum is numeric:-
   {if(NumbOK==FALSE)                                                                              #and If NumbOK  (required arg) is false,
      {ERR <- TRUE; Msg <- "A numeric value is not allowed."                                       #set ERR and message
      } else                                                                                       #else If Datum is numeric and is allowed to be:-
      {if(!is.na(ZeroOK))                                                                          #If ZeroOK is supplied as an argument:-
         {if(ZeroOK==FALSE & Datum == 0)                                                           #and If ZeroOK is false but Datum is zero:-
            {ERR <- TRUE; Msg <- "A value of zero is not allowed."                                 #set ERR and message
            }
         }                                                          
      if(!is.na(InfinOK))                                                                          #If InfinOK is supplied as an argument:-
         {if(InfinOK==FALSE & abs(Datum) == Inf)                                                   #and If InfinOK is false but absolute(Datum) is infinite:-
            {ERR <- TRUE; Msg <- "A value of +/-Inf (infinity) is not allowed."                    #set ERR and message
            }
         }   
      if(!is.na(MustBeInSet[1]))                                                                   #If MustBeInSet is supplied as an argument:-
         {if(!is.element(Datum, MustBeInSet))                                                      #and If Datum is not an element of the supplied set:-
            {ERR <- TRUE; Msg <- c("The value must be in the set:-   {", MustBeInSet, "}")         #set ERR and message
            }
         }
      if(!is.na(MustNotBeInSet[1]))                                                                #If MustNotBeInSet is supplied as an argument:-
         {if(is.element(Datum, MustNotBeInSet))                                                    #and if Datum is an element of the supplied set:-
            {ERR <- TRUE; Msg <- c("The value must NOT match any member of the set:-   {", MustNotBeInSet, "}") #set ERR and message
            }
         }            
      if(!is.na(nonIntegerOK))                                                                     #If nonIntegerOK is supplied as an argument:- 
         {if(nonIntegerOK==FALSE & floor(abs(Datum)) < abs(Datum))                                 #and If Datum is neither a positive nor a negative integer:-
            {ERR <- TRUE; Msg <- "The value must be an integer."                                   #set ERR and message 
            }
         }
      if(!is.na(MaxOK))                                                                            #If MaxOK is supplied as an argument:-
         {if(Datum > MaxOK)                                                                        #and if MaxOK is exceeded:-
            {ERR <- TRUE; Msg <- paste("The value must be less than or equal to ", as.character(MaxOK), sep="") #set ERR and message
            }
         }   
      if(!is.na(MinOK))                                                                            #If MinOK is supplied as an argument:-
         {if(Datum < MinOK)                                                                        #and if MinOK is greater than Datum:-
            {ERR <- TRUE; Msg <- paste("The value must be greater than or equal to ", as.character(MinOK), sep="") #set ERR and message
            }
         }         
      if(!is.na(MaxNotOK))                                                                         #If MaxNotOK is supplied as an argument:-
         {if(Datum >= MaxNotOK)                                                                    #and If MaxNotOK is not greater than Datum:-
            {ERR <- TRUE; Msg <- paste("The value must be less than ", as.character(MaxNotOK), sep="") #set ERR and message
            }
         }   
      if(!is.na(MinNotOK))                                                                         #If MinNotOK is supplied as an argument:-
         {if(Datum <= MinNotOK)                                                                    #and If Datum is not greater than MinNotOK:-
            {ERR <- TRUE; Msg <- paste("The value must be greater than ", as.character(MinNotOK), sep="")  #set ERR and message
            }                                                                         
         }                                                                
      }
   }
if(is.logical(Datum) & !ERR)                                                                       #If Datum is logical:-
   {if(LogicalOK==FALSE)                                                                           #and if Datum may not be logical:-
      {ERR <- TRUE; Msg <- "A logical value is not allowed."                                       #set ERR and message
      } else
      {if(!is.na(MustBeInSet[1]))                                                                  #If MustBeInSet is supplied as an argument:-
          {if(!is.element(Datum, MustBeInSet))                                                     #and if Datum is not an element of the supplied set:-
             {ERR <- TRUE; Msg <- c("The value must be in the set:-   {", MustBeInSet, "}")        #set ERR and message
             }
          }
      if(!is.na(MustNotBeInSet[1]))                                                                #If MustNotBeInSet is supplied as an argument:-
         {if(is.element(Datum, MustNotBeInSet))                                                    #and If Datum is an element of the supplied set:-
            {ERR <- TRUE; Msg <- c("The value must NOT match any member of the set:-   {", MustNotBeInSet, "}")
            }
         }            
      }
   }      
if(ERR)                                                                                            #If there is an error, advise, giving rule broken:- [Following code is replaced by special code in Main Program]
   {#cat("\n\nis.datumOK?  NO: A supplied value, ", VarName, "= ", Datum, ", breaks a rule for this variable:-")  #these statements could be used (remove #) if calling program does not supply appropriate advice
   #cat("\nRule:", Msg, "\n")
   #cat("**NOTE: The class of a value may be changed by the classes of other values in the same data structure.\n")  #eg. a numeric can be changed to character by R if a single value in the vector is character
   }    
return(ERR)                                                                                        #Return ERR to allow program to stop or use default values.
}
################################################################# End of function is.DatumOK

   
nrow2 <- function(VecOrMat, N.cols){                             
#Function to find the number of rows in a row vector, a col vector, or a matrix, given N.cols known. R's nrow() does not work with 1 row or 1 col only. They can occur frequently during operations of ECOLPS.  nrow() does work for 1-row arrays but the array attribute may be lost when subsetting to yield a single row.
nrow2 <- floor(length(VecOrMat)/N.cols)                                                            #Matrices and vectors are stored as vectors. Dividing N elements by N.cols -> N.rows ; floor rounds down non-integers and may give 0
}
#################################################################End of function nrow2()


ncol2 <- function(VecOrMat, N.rows){                             
#Function to find the number of cols in a row vector, a col vector, or a matrix, given N.rows known. R's ncol() does not work with 1 row or 1 col only. They can occur frequently during operations of ECOLPS.  ncol() does work for 1-col arrays but the array attribute may be lost when subsetting to yield a single row.
nrow2 <- floor(length(VecOrMat)/N.rows)                                                            #Matrices and vectors are stored as vectors. Dividing N elements by N.rows -> N.cols ; floor rounds down non-integers and may give 0
}
#################################################################End of function nrow2()


PickActiveConstraints <- function(PickSet, ConstraintBindings){
#Function to find row numbers of sometime-active constraints in a subset of all constraints (ConstraintBindings from Slacks) specified by row numbers in Pickset
#ConstraintBindings has all constraint rows with 0=unbinding, 1=binding (=active) for each Delta (time).  Pickset is its vector of row numbers
PickSet.active <- rep(FALSE, N.C)                                                                  #vector to mark row numbers of constraints within PickSet active at least once during simulation. N.C is N cols in the set
for(PSa in PickSet)                                                                                #For each row of the selected constraints,
   {if(sum(ConstraintBindings[PSa, ], na.rm=TRUE) > 0) PickSet.active[PSa] <- TRUE                 #If there was at least one time when the constraint was active, mark as TRUE
   }
N.ActiveConstr <- sum(PickSet.active) > 0                                                          #number of active constraints in PickSet
if(N.ActiveConstr > 0)                                                                             #If any constraints in PickSet are active at any time:-
   {ActiveConstr <- which(PickSet.active == TRUE)                                                  #AC holds a vector of numbers of constraints active at least once
   return(ActiveConstr)                                                                            #return row numbers of PickSet that are active
   } else return(NA)                                                                               #else If none, return NA
}
#################################################################End of function PickActiveConstraints()


Plot.results <- function(Plot.matrix, AxLabel, TitleText, N.plotVars, TitleTime, KeyAuto, N.keyLines, MaxOrdinate, MinYearOut, MaxYearOut){
#Plot.results() is called by ProcessOutput() below. It implements the time-series plotting option for selected output variables
#Simulation results are plotted as time series using R graphics functions.  Plot.matrix is times x variables, as needed for plot().  Argument AxLabel labels the y-axis.  Argument  OutputDeltas [used by plot() and lines() statements below] allows restriction of plots to a subset of available times. It is a list of row numbers selected from all times for plotting with menu options B, F or I.  Time in Deltas is converted to time in Years for plotting purposes using YearLength (=Deltas per year) in Controls.txt.
   #SECTIONS
      #PR1. Prepare plot dimensions, symbols, and colours 
      #PR2. Prepare graphic parameters
      #PR3. Plot first variable to create axes and labels, then plot all other variables with lines() 
      #PR4. Add key to plot
Plot.matrix <- t(Plot.matrix)                                                   #first, transpose the plot matrix for plot() so that time runs vertically, variables horizontally
#Plot.matrix[nrow(Plot.matrix), ] <- ifelse(!is.na(Plot.matrix[nrow(Plot.matrix), ]) & Plot.matrix[nrow(Plot.matrix), ] != 0, Plot.matrix[nrow(Plot.matrix), ], NA) #replace 0s <- NA in last row to avoid stepdown; NAs don't plot 
#print(Plot.matrix)
#PR1. Prepare key lines, symbols, and colours 
if(N.plotVars > 62)                                                             #If there are more variables to plot than there are available symbols:-  [In practice, 62 is probably way too many variables on one graph]
   {cat("\nADVICE:- Too many variables [", N.plotVars, "] requested for one time-series graph.  Only the first 62 symbols will be plotted.  Use other Menu options to see all results by groups.\n")  #advise                      
   N.plotVars <- 62                                                             #reduce the number of variables to the maximum available
   }
if(KeyAuto == TRUE)                                                             #If automatic setting of the number of keylines is TRUE:- [changed with K option of Constraints and Variables menus]
   {N.keyItemsPerLine <- KeyLineLength %/% KeyItemWidth                         #set the number of key items that will fit in each line; KLL and KIW are supplied in Controls.txt
   N.keyLines <- 3 + N.plotVars %/% N.keyItemsPerLine                           #required number of keylines including 1 for key title, 1 for blank footer, and 1 for any part of a line.  N.keyLines is applied in par(oma=...) below.
   } 

Plot.symbols <- noquote(c(as.character(c(1:9,0)), letters, LETTERS))[1:N.plotVars]  #Full list of available plot symbols for variables.  Max of 10+26*2=62 on one graph
Pal.BnW <- rep("black", 72)                                                     #palette for black and white graphs; all colours are black. Below: remove all but one palette by placing # at start of line
Pal.Col <- rep(c("#000000", "#D55E00", "#117733", "#88CCEE", "#CC6677", "#DDCC77", "#44AA99", "#E69F00", "#F0E442", "#661100", "#332288", "#888888"), 6)  #palette for coloured graphs, codes 1 - 12 recycled 6 times gives 72codes 
#Pal.Col are by approx name: black, orange, green, sky, pink, straw, turquoise, gold, yellow, brown, mauve, grey. To view these colors, at R prompt, submit Pal.Col <- . . .; windows() ; pie(rep(1,12), col=Pal.Col) 
                                                                                #Or, for a different sequence of colours:-  
#Pal.Col <- rep(c("#000000", "#D55E00", "#117733", "#CC6677", "#88CCEE", "#DDCC77", "#44AA99", "#E69F00", "#F0E442", "#661100", "#332288", "#888888"), 6)  #palette for coloured graphs, codes 1 to 12 recycled 6 times gives 72codes 
#Pal.Col are by approx name: black, orange, green, pink, sky, straw, turquoise, gold, yellow, brown, mauve, grey. To view these colors, at R prompt, submit Pal.Col <- . . .; windows() ; pie(rep(1,12), col=Pal.Col)   

#----------------------------------------------------------------

#PR2. Prepare graphic parameters
Y.Axmax <- ifelse(MaxOrdinate < Inf, MaxOrdinate, max(Plot.matrix, na.rm=TRUE)) #maximum y value is MaxOrdinate If finite [& not Active constr being plotted], Else = max value in the plot matrix
Plot.matrix.adj <- ifelse(Plot.matrix > Y.Axmax, NA, Plot.matrix)               #this line omits elements of Plot.matrix higher than Y.Axmax; NAs are not plotted
Y.Axmin <- suppressWarnings(min(min(Plot.matrix, na.rm=TRUE), 0))               #the minimum y value; it is zero for +ve variables and -ve if negatives are present
if(Jitter > 0) Plot.matrix.adj[ , ] <- pmax(Plot.matrix.adj[ , ] +  runif(length(Plot.matrix), -1, +1) * (Y.Axmax - Y.Axmin) * Jitter, 0) #jitter plotted points by factor * random-uniform value in -1 and +1  * vert range, or 0
Plot.type <- "b"                                                                #Plot symbols as points with no lines between them; usually there is not room for lines between
y.ticks <- 10                                                                   #number of tick marks on Y axis for standard variables
windows(width=WinWidth, height=WinHeight, restoreConsole=TRUE)                  #prevents over-writing of previous graphs; must be before par(); returns to keyboard input after plot
#if(AxLabelFactor <= 0) par(oma=c(N.keyLines, 0,0,0), mar=c(0.5,0.5,2,1), lheight=1.2, mgp=c(2.6,1,0)) else  par(oma=c(N.keyLines, 0,0,0), mar=c(3.8,4.1,2,1), lheight=1.2, mgp=c(2.6,1,0))
#marVec <- ifelse(AxLabelFactor > 0, c(3.8,4.1,2,1), c(0.5,0.5,2,1))            #these 2 lines are probably not needed but kept just in case they are found necessary!
par(oma=c(N.keyLines, 0,0,0), mar=c(3.8,4.1,2,1), lheight=1.2, mgp=c(2.6,1,0))  #oma=>set lower outer margin size in lines (1 needed for 'Symbols:-' heading); mar=>set margin widths for annotation of axes; lheight=>set line height of key in lower outer margin; mgp=>set axis title, labels and position; recommended default is mar=c(3.8,4.1,2,1)       
AxLim.y <- c(Y.Axmin, Y.Axmax)                                                  #limits of y axis for quantitative variables
AxLim.x <- c(MinYearOut, MaxYearOut)                                            #limits of x axis for time. Note: you can change MaxYearOut with the F option, and Years[StartOut] with the B option
if(Col.plots == 0)                                                              #If Col.plots switch read from Controls.txt is zero:-
   {palette (Pal.BnW)                                                           #all plotted lines and symbols will be black; only works for simple graphs
   LineType <- rep(c(1:6), 12)                                                  #a different line type (from 6 available) is used for every variable
   }
if(Col.plots == 1)                                                              #If Col.plots switch is 1:-   
   {palette (Pal.Col)                                                           #load Pal.Col with 12 colours repeated 6 times, constructed above
   LineType <-  c(rep(1,12),rep(2,12),rep(3,12),rep(4,12),rep(5,12),rep(6,12))  #a different line type (from 6 available) is used for each set of 12 colours, supposing you can distinguish them on the graph
   }
#----------------------------------------------------------------

#PR3. Plot first variable to create axes and labels, then plot all other variables with lines()
PSS <- PlotSymbolSize                                                           #size of number or letter marking time series points; typically 0.8 
plot( Plot.matrix.adj[ , 1] ~ Years[OutputDeltas], data=Plot.matrix.adj, type=Plot.type, ylab=AxLabel, ylim=AxLim.y, xlab=TimeLabel, xlim=AxLim.x, lab=c(5, y.ticks, 7), cex.lab=1.5 * AxLabelFactor, pch=Plot.symbols[1], cex=PSS, cex.axis=AxLabelFactor, lwd=3, col=palette()[1], lty=LineType[1], xpd=TRUE)        #cex controls symbol size, lwd does width of symbol border. [Insert extra plot par to extend x-axis, eg ", xlim=c(0, 1000)"]
grid(nx=NULL)                                                                   #puts a faint grid over the plot region aligned with tick marks
if(N.plotVars > 1)                                                              #If more than one variable is to be plotted on the same axes using lines():-
   {for (j in 2:N.plotVars)                                                     #For each of the remaining variables to be plotted,   
      {OutTimesVec <- c(OutputDeltas[1], OutputDeltas[2:(length(OutputDeltas)-1)], OutputDeltas[length(OutputDeltas)]) #staggers intermediate points depending on Interval & j for better visibility
      lines(Plot.matrix.adj[ , j] ~ Years[OutTimesVec], data=Plot.matrix, type=Plot.type,  pch=Plot.symbols[j], cex=PSS, lwd=3, col=palette()[j], lty=LineType[j]) #plot points with calculated gaps between
      }                                                                  
   }
#----------------------------------------------------------------

#PR4. Add key to plot
LocFromLeft <- 0                                                                #left justify the key; this variable ranges up to 1, see below
KeyLineN <- 0                                                                   #current line number of key, starting from 0 in the area beneath the graph
mtext("  Symbols:-  ", side=1, line=KeyLineN, outer=TRUE, adj=0, cex=1, lheight=2, family="mono", col=palette()[1], at=LocFromLeft) #palette()[1] is black for the heading "Symbols"
KeyLineN <- 1                                                                   #put the key below the heading
for (x in 1:ncol(Plot.matrix))                                                  #For each variable column in Plot.matrix,
   {if(LocFromLeft > 1 - KeyItemWidth/ KeyLineLength)                           #If there is not room for another plot symbol:- [KeyItemWidth and KeyLineLength were set in Controls.txt]
      {LocFromLeft <- 0                                                         #start a new line from 0
      KeyLineN <- KeyLineN + 1                                                  #add one to KeyLineN
      }
   mtext(paste(Plot.symbols[x], "=", sep=""),  side=1, line=KeyLineN, outer=TRUE, adj=0, cex=1, lheight=2, family="mono", col=palette()[x], at=LocFromLeft)#write the plot symbol in the plot colour with an "="
   LocFromLeft <- LocFromLeft + 2 / KeyLineLength                               #insert a small space between = and var name
   mtext(substr(colnames(Plot.matrix)[x], 1, (KeyItemWidth - 2)), side=1, line=KeyLineN, outer=TRUE, adj=0, cex=1, lheight=2, family="mono", col=palette()[1], at=LocFromLeft) #write variable name in black
   LocFromLeft <- LocFromLeft + (KeyItemWidth - 2) / KeyLineLength              #move LocFromLeft to the right as measured by width needed for each key item
   }
if(TitleTime == FALSE) {DatedTitleText <- TitleText} else {DatedTitleText <- paste0(TitleText, " - ", substr(as.character(date()),5,16))} #if time switch is true, add date and time string to title
title(main=DatedTitleText, line=1)                                              #the graph can be titled with this statement but par(mar= ) above may need adjustments
}
################################################################# End of function Plot.results()


ProcessOutput <- function(Choice, ItemName, Ordinate.label, N.rows, TitleText, TitleTime, KeyAuto, N.keyLines, MaxOrdinate, MinYearOut, MaxYearOut){
#Performs printing on screen, graphing, or analysis of peaks and valleys in Output.set (stored at program body level); returns a message as a report of the action
#Choice is the letter entered by user in response to the constraint or variable menus; ItemName is the name for the types of variable in Output.set; N.rows is the length of the Output.set
#MaxYearOut is the length of the X-axis for time-series graphs
   #SECTIONS
      #PO0 Preliminary dimensioning and subsetting as Temp.set
      #PO1 Printing = screening
      #PO2 Plotting
      #PO3 Analysis of peaks and valleys

#PO0 Preliminary dimensioning and subsetting as Process.set
#Process.set is a temporary set so that the original Output.set is unchanged and could be extended etc if required later
#OutputDeltas subsets by time in columns; see Thinning in EO8 of EditOutput(); this code should handle Output.set from simulations that terminated prematurely
if(N.rows == 0) {Message <- "The Output.set has no rows to process: "; return(Message)}            #message goes under next menu
if(N.rows == 1) Process.set <- array(Output.set[OutputDeltas], dim=c(1, length(OutputDeltas)), dimnames=list(row.names(Output.set), names(SimTime)[OutputDeltas])) #OutputDeltas subsets by time in columns; see Thinning in EO8 of EditOutput(). Note: EndOut=last M if Output.set is from a simulation that could not be completed (See #7.10.3.7.2 in main code)      
if(N.rows > 1) Process.set <- array(Output.set[ , OutputDeltas], dim=c(N.rows, length(OutputDeltas)), dimnames=list(row.names(Output.set), names(SimTime)[OutputDeltas])) #OutputDeltas subsets by time in columns; see Thinning in EO8 of EditOutput(). Note: EndOut=last M if Output.set is from a simulation that could not be completed (See #7.10.3.7.2 in main code)
#----------------------------------------------------------------                                                                                     

#PO1 Printing = screening
if(Choice == "P" | Choice == "p")                                               #If screening of Output.set is selected:-
   {Title.tmp <- TitleText                                                      #store the existing title
   if(length(grep(", Jittered", Title.tmp)) > 0) Title.tmp <-  gsub(", Jittered", "", Title.tmp) #remove mention of Jittering if present in temporary title because printouts are never jittered
   cat("\n\n", Title.tmp, "\n")                                                 #print a general heading on screen
   cat("\n  ", gsub("(\\w)(\\w*)", "\\U\\1\\L\\2", Ordinate.label, perl=TRUE), "versus Years\n")#print a specific heading; perl code, from Crawley MJ, capitalises 1st letter of ItemName
   if(N.rows > 1) print(Process.set, digits=4)                                  #If more than 1 row, use print(Process.set[ , OutputDeltas[OutputDeltas <= (M+1)]])
   if(N.rows == 1) print(noquote(format(Process.set, digits=3)))                #R does not print a row name with 1-row vectors; this code is a fix
   Message <- "See the DATA printed above"                                      #advise that a table is printed above (because the menu may obscure the table)
   }
#----------------------------------------------------------------


#PO2 Plotting
if(Choice == "G" | Choice == "g")                                               #If plotting of Output.set is selected:-
   {MaxYvalue <- 10e+200                                                        #arbitrary 'excessive value' for sensible plots; prevents R errors when ylim is too high
   if(sum(Process.set, na.rm=TRUE) == 0) MaxOrdinate <- 1                       #If Process.set only has NAs in it, set MaxOrdinate to 1:- [This value is not passed back to main program]
   if(suppressWarnings(max(Process.set, na.rm=TRUE)) < MaxYvalue | MaxOrdinate < MaxYvalue) #If the maximum value for plotting is not excessive, or values > MaxOrdinate will be censored:-
      {Plot.results(Process.set, Ordinate.label, TitleText, N.rows, TitleTime, KeyAuto, N.keyLines, MaxOrdinate, MinYearOut, MaxYearOut) #graph the transposed results with Plot.results().  Ordinate.label labels the y-axis
      Message <- "See the PLOT in the active graphics window"                   #advice for screening below the next menu output
      } else                                                                    #else If the max value for plotting is too high:-
      {Message <- paste0("Some values to be plotted exceed ", MaxYvalue, " . Plot ABORTED. Use F to crop end of series, \nor O to crop the higher values")
      }
   }


#PO3 Analysis of peaks and dips 
#Lists peaks and dips in time series, their time-locations and values, plus frequencies and wavelengths.  Input for this function is the TRANSPOSE of Process.set. There must be enough time points to enable peaks and dips to exist.
#Comparison of early and late waves allows a check on stability of cycles if there is interest in steady-state conditions
#Code below assumes that peaks and dips mean something in the Output.set.  eg. There is no point in looking for them in a set of Active constraint markers.  Also, numerical wobbles in time series close to steady state, ie. tracing a flat path over time, may be mistaken for model-dependent oscillations -> number of peaks, wavelength, frequencies etc will be misleading/inaccurate.  
#NB. This code is not suitable for a Process.set that has been thinned!! Thinning would alter visible peaks, dips, wavelengths and amplitudes.
if(Choice == "A" | Choice == "a")                                                                            #If analysis of peaks and dips is selected from the Variables menu [only]:-
   {Process.set <- t(Process.set)                                                                            #transpose the data set to give TIMES x VARIABLES
   N.t <- nrow(Process.set)                                                                                  #N of rows corresponding to times; row names are 'years'=Deltas.
   if(N.t < 6) {Message <- "Too few time points; analysis of peaks and dips cannot be usefully done: "; return(Message)}  #the message goes to the screen after the next menu
   N.v <- ncol(Process.set)                                                                                  #N of cols corresponding to variables
   TS.names <- colnames(Process.set)                                                                         #names of variables
   #array declarations
   Peak.pts  <- array(0, dim=c(N.t, N.v), dimnames = list(row.names(Process.set), TS.names))                 #initialise array with peak points initially marked with 1; a peak is the highest point locally
   Dip.pts  <- array(0, dim=c(N.t, N.v), dimnames = list(row.names(Process.set), TS.names))                  #initialise array with dip points marked with -1; a dip is the lowest point locally
   Peaks <- array(0, dim=c(nrow(Peak.pts), N.v), dimnames = list(row.names(Peak.pts), TS.names))             #values at peaks will be stored in this array
   Dips <- array(0, dim=c(nrow(Dip.pts), N.v), dimnames = list(row.names(Dip.pts), TS.names))                #values at dips will be stored in Dips
   Peak.times <- array(0, dim=c(nrow(Peak.pts), N.v), dimnames = list(row.names(Peak.pts), TS.names))        #times of peaks 
   Dip.times <- array(0, dim=c(nrow(Dip.pts), N.v), dimnames = list(row.names(Dip.pts), TS.names))           #times of dips
   N.peaks <- array(0, dim=c(1, N.v), dimnames = list("N.peaks", TS.names))                                  #number of peaks in entire Process.set
   N.dips <- array(0, dim=c(1, N.v), dimnames = list("N.dips", TS.names))                                    #number of dips in entire Process.set
   AnnualDrift.peaks <- array(NA, dim=c(1, N.v), dimnames = list("PeakDrift/Delta", TS.names))               #difference in height between first and last peaks; slope is averaged per Delta ('year')  
   AnnualDrift.dips <- array(NA, dim=c(1, N.v), dimnames = list("DipDrift/Delta", TS.names))                 #difference in height between first and last dips averaged per Delta ('year') 
   FirstWaveL.peaks <- array(NA, dim=c(1, N.v), dimnames = list("Peaks.1stWaveLength", TS.names))            #time interval = wavelength between first 2 peaks
   FirstWaveL.dips <- array(NA, dim=c(1, N.v), dimnames = list("Dips.1stWaveLength", TS.names))              #time interval = wavelength between first 2 dips
   LastWaveL.peaks <- array(NA, dim=c(1, N.v), dimnames = list("Peaks.LastWaveLength", TS.names))            #time interval = wavelength between last 2 peaks
   LastWaveL.dips <- array(NA, dim=c(1, N.v), dimnames = list("Dips.LastWaveLength", TS.names))              #time interval = wavelength between last 2 dips
   AnnualWaveLDrift.peaks <- array(NA, dim=c(1, N.v), dimnames = list("Peaks.WaveLDrift/Delta", TS.names))   #drift of wavelength per delta for peaks
   AnnualWaveLDrift.dips <- array(NA, dim=c(1, N.v), dimnames = list("Dip.WaveLengthDrift/Delta", TS.names)) #drift of wavelength per delta for dips
   FirstAmplitude <- array(NA, dim=c(1, N.v), dimnames = list("FirstAmplitude", TS.names))                   #amplitude = difference between 1st peak and 1st subsequent dip
   LastAmplitude <- array(NA, dim=c(1, N.v), dimnames = list("LastAmplitude", TS.names))                     #amplitude of last wave
   AnnualAmplitudeDrift <- array(NA, dim=c(1, N.v), dimnames = list("AmplitudeDrift/Delta", TS.names))       #drift of amplitude per delta
   #Points (=time locations) where peaks and dips occurred, numbers of them, and their biomass values

   for(Var.C in 1:N.v)                                                                                       #For each variable supplied in Process.set,
      {Peak.count <- 1; Dip.count <- 1                                                                       #initialise counters
      for(ii in 2:(N.t - 1))                                                                                 #For peaks or dips from year 2 (row) to 1 before last because 3 consec values are needed to decide a peak or dip
         {if(is.na(Process.set[ii-1, Var.C]) | is.na(Process.set[ii, Var.C]) | is.na(Process.set[ii+1, Var.C])) next() #if any value needed to identify a peak at ii is NA, move to the next ii loop 
         if((Process.set[ii, Var.C] > Process.set[ii-1, Var.C]) & (Process.set[ii, Var.C] > Process.set[ii+1, Var.C])) #finds a peak if TRUE
            {Peaks[ii, ] <- Process.set[ii, ]                                                                #transfer peak value to Peaks in the same row (other rows are zero)
            Peak.pts[ii, Var.C] <- Peak.count                                                                #transfer count of peaks to Peak.pts in the same row; this overwrites the 1 there previously
            Peak.count <- Peak.count + 1                                                                     #increment the counter
            }
         if((Process.set[ii, Var.C] < Process.set[ii-1, Var.C]) & (Process.set[ii, Var.C] < Process.set[ii+1, Var.C]))  #finds a dip if TRUE
            {Dips[ii, ] <- Process.set[ii, ]                                                                 #transfer the dip value to Dips in the same row (other rows are zero)
            Dip.pts[ii, Var.C] <- Dip.count                                                                  #transfer the count of dips to Dip.pts in the same row
            Dip.count <- Dip.count + 1                                                                       #increment the counter
            }
         } 
      Peak.pts[ , Var.C] <- ifelse(is.na(Peak.pts[ , Var.C]), 0, Peak.pts[ , Var.C])                         #NAs might be present; convert to 0s, ie. not a peak                                                                 
      Dip.pts[ , Var.C] <- ifelse(is.na(Dip.pts[ , Var.C]), 0, Dip.pts[ , Var.C])                            #NAs might be present; convert to 0s, ie. not a  dip                                                                  
      Peak.times[ , Var.C] <- ifelse(Peak.pts[ , Var.C] > 0,  as.numeric(gsub("y", "", row.names(Process.set))[which(row.names(Process.set) %in% row.names(Peak.pts))]), 0) #assign a Deltas column, and insert the row times
      Dip.times[ , Var.C] <- ifelse(Dip.pts[ , Var.C] < 0,  as.numeric(gsub("y", "", row.names(Process.set))[which(row.names(Process.set) %in% row.names(Dip.pts))]), 0) #assign a Deltas column, and insert the row times
      N.peaks[1, Var.C] <- max(Peak.pts[ , Var.C])                                                           #N of peaks is the peak count stored for the last peak of the series
      N.dips[1, Var.C] <- max(Dip.pts[ , Var.C])                                                             #N of dips is the dip count stored for the last dip of the series
#For peaks
      if(N.peaks[1, Var.C] < 2)                                                                              #If there is only 1 or 0 peaks,
         {FirstWaveL.peaks[1, Var.C] <- NA                                                                   #first wavelength for peaks cannot be found
         LastWaveL.peaks[1, Var.C] <- NA                                                                     #nor the last
         AnnualWaveLDrift.peaks[1, Var.C] <- NA                                                              #nor the drift of wavelengths
         } else                                                                                              #else, If there are 2 or more peaks,
         {Peak1.R <- which(Peak.pts[ , Var.C] == 1)                                                          #the row of the first peak in Process.set etc
         Peak2.R <- which(Peak.pts[ , Var.C] == 2)                                                           #the row of the second peak
         PeakPenultimate.R <-  which(Peak.pts[ , Var.C] == N.peaks[1, Var.C] - 1)                            #the row of the penultimate peak
         PeakLast.R <-  which(Peak.pts[ , Var.C] == N.peaks[1, Var.C])                                       #the row of the last peak
         FirstPeak.Time <- as.numeric(gsub("y", "", row.names(Process.set)[Peak1.R]))                        #the time of first peak . . .
         SecondPeak.Time <- as.numeric(gsub("y", "", row.names(Process.set)[Peak2.R]))
         PenultimatePeak.Time <- as.numeric(gsub("y", "", row.names(Process.set)[PeakPenultimate.R]))
         LastPeak.Time <- as.numeric(gsub("y", "", row.names(Process.set)[PeakLast.R]))
         FirstWaveL.peaks[1, Var.C] <- SecondPeak.Time - FirstPeak.Time                                      #the wavelength =period between the 1st 2 peaks
         LastWaveL.peaks[1, Var.C] <-  LastPeak.Time - PenultimatePeak.Time                                  #same for last 2 peaks
         if(N.peaks[1, Var.C] < 3)                                                                           #If there are fewer than 3 peaks:-
            {AnnualDrift.peaks <- NA                                                                         #the drift of peak heights isn't found
             AnnualWaveLDrift.peaks[1, Var.C] <- NA                                                          #the drift of wavelength cannot be found
            } else                                                                                           #else there are 3 or more peaks,
            {AnnualDrift.peaks[1, Var.C] <- (Peaks[PeakLast.R, Var.C] - Peaks[Peak1.R, Var.C]) / (LastPeak.Time - FirstPeak.Time) #drift of peak heights per delta
             AnnualWaveLDrift.peaks[1, Var.C] <- (LastWaveL.peaks[1, Var.C] - FirstWaveL.peaks[1, Var.C]) / (PenultimatePeak.Time - FirstPeak.Time) #drift of wavelength between peaks per delta
            }
         }
#For dips         
      if(N.dips[1, Var.C] < 2)                                                                               #If there is only 1 or 0 dips:-
         {FirstWaveL.dips[1, Var.C] <- NA                                                                    #first wavelength dips cannot be found
         LastWaveL.dips[1, Var.C] <- NA                                                                      #nor the last
         AnnualWaveLDrift.dips[1, Var.C] <- NA                                                               #nor the drift of wavelengths
         } else                                                                                              #else, If there are 2 or more dips:-
         {Dip1.R <- which(Dip.pts[ , Var.C] == 1)                                                            #the row of the first dip in transposed Process.set etc
         Dip2.R <- which(Dip.pts[ , Var.C] == 2)                                                             #the row of the second dip
         DipPenultimate.R <-  which(Dip.pts[ , Var.C] == N.dips[1, Var.C] - 1)                               #the row of the penultimate dip
         DipLast.R <-  which(Dip.pts[ , Var.C] == N.dips[1, Var.C])                                          #the row of the last dip
         FirstDip.Time <- as.numeric(gsub("y", "", row.names(Process.set)[Dip1.R]))                          #the time of the first dip . . . 
         SecondDip.Time <- as.numeric(gsub("y", "", row.names(Process.set)[Dip2.R]))
         PenultimateDip.Time <- as.numeric(gsub("y", "", row.names(Process.set)[DipPenultimate.R]))
         LastDip.Time <- as.numeric(gsub("y", "", row.names(Process.set)[DipLast.R]))
         FirstWaveL.dips[1, Var.C] <- SecondDip.Time - FirstDip.Time                                         #the wavelength =period between the 1st 2 dips
         LastWaveL.dips[1, Var.C] <-  LastDip.Time - PenultimateDip.Time                                     #same for last 2 dips
         if(N.dips[1, Var.C] < 3)                                                                            #If there are fewer than 3 dips:-
            {AnnualDrift.dips <- NA                                                                          #the drift of dip heights isn't found
            AnnualWaveLDrift.dips[1, Var.C] <- NA                                                            #the drift of wavelength cannot be found
            } else                                                                                           #else If there are 3 or more dips:-
            {AnnualDrift.dips[1, Var.C] <- (Dips[DipLast.R, Var.C] - Dips[Dip1.R, Var.C]) / (LastDip.Time - FirstDip.Time) #drift of dip heights per delta
            AnnualWaveLDrift.dips[1, Var.C] <- (LastWaveL.dips[1, Var.C] - FirstWaveL.dips[1, Var.C]) / (PenultimateDip.Time - FirstDip.Time) #drift of wavelength between dips per delta
            }
         }
#For peaks and dips
      if(N.peaks[1, Var.C] < 3 | N.dips[1, Var.C] < 3)                                                       #If there are fewer than 3 peaks or dips:-
         {FirstAmplitude[1, Var.C] <- NA                                                                     #first amplitude cannot be found
         LastAmplitude[1, Var.C] <- NA                                                                       #last amplitude cannot be found
         AnnualAmplitudeDrift[1, Var.C] <- NA                                                                #nor the drift of amplitudes
         } else                                                                                              #else, If there are 3 or more peaks and dips:-
         {DipAfterFirstPeak.R <- ifelse(FirstDip.Time > FirstPeak.Time, Dip1.R, Dip2.R)                      #row in transposed Process.set of the 1st dip after the first peak 
         FirstAmplitude[1, Var.C] <- Process.set[Peak1.R, Var.C] - Process.set[DipAfterFirstPeak.R, Var.C]   #1st amplitude is 1st peak - 1st subsequent dip
         PeakBeforeLastDip.R <-  ifelse(LastPeak.Time < LastDip.Time, PeakLast.R, PeakPenultimate.R)         #row of the last peak before the last dip
         LastAmplitude[1, Var.C] <- Process.set[PeakBeforeLastDip.R, Var.C] - Process.set[DipLast.R, Var.C]  #last amplitude is difference between last peak before last dip, and the last dip 
         AnnualAmplitudeDrift[1, Var.C] <- (LastAmplitude[1, Var.C] - FirstAmplitude[1, Var.C]) / (LastDip.Time - FirstDip.Time) #drift of amplitudes per unit time.
         }
      }

#Output         
   cat("\n\nANALYSIS OF PEAKS AND DIPS\n")
   cat("\nTIMINGS (Deltas), PEAKS, DIPS, AND VALUES AT PEAKS OR DIPS OF OTHER COMPONENT(S):-\n")             #print out peaks to screen
   Peaks <- Peaks[which(rowSums(Peaks) > 0), ]
if(length(Peaks) < 4) {Message <- "Too few peaks and dips were found in the time series for analysis" ; return(Message)}
   N.ActivityMarkers <- length(grep("Active", row.names(Output.set)))                                        #activity markers are rows of 1s, 2s, 3s, etc.. array() statements fail with these in Process.set (=t(Output.set))
   if(N.ActivityMarkers > 0)                                                                                 #length(grep()) counts how many activity markers are in Process.set
      {Message <- "Constraint activity markers are included in Output.set. Analysis of peaks and dips failed:"  #if more than 0, an error message is returned
      return(Message)
      }
#print(Peaks); stop("843")
   Peaks <- array(cbind(as.numeric(gsub("y", "", row.names(Peaks))), Peaks), dim=c(nrow(Peaks), ncol(Peaks) + 1), dimnames= list(paste0("Peak.", 1:nrow(Peaks)), c("Deltas", colnames(Peaks)))) #put in a time=Deltas column
   Dips <- Dips[which(rowSums(Dips) > 0), ]                                                                  #Dips only holds rows with 1 or more dips
   Dips <- array(cbind(as.numeric(gsub("y", "", row.names(Dips))), Dips), dim=c(nrow(Dips), ncol(Dips) + 1), dimnames= list(paste0("Dip.",1:nrow(Dips)), c("Deltas", colnames(Dips))))  #put in a time=Deltas column
   PeaksAndDips <- data.frame(rbind(Peaks, Dips))                                                            #bind rows into data.frame for neat printing of peak and dip values, together with other variables at those times
   PeaksAndDips <- PeaksAndDips[order(PeaksAndDips$Deltas), ]                                                #sort all rows by time 
   print(noquote(format(PeaksAndDips, width=12, digits=4)))                                                  #print on screen                      
   cat("\nNUMBERS OF PEAKS AND DIPS:-\n")                                                                    #title on screen
   N.PeaksAndDips <- data.frame(rbind(N.peaks, N.dips))                                                      #number of peaks and dips in each variable column
   print(noquote(format(N.PeaksAndDips, width=12)))                                                          #print on screen N peaks and dips
   cat("\nDRIFT OF PEAKS AND DIPS FROM FIRST TO LAST:-\n")                                                   #title on screen
   DriftPerDelta <- data.frame(rbind(AnnualDrift.peaks, AnnualDrift.dips))                                   #drift of peak and drift heights from first to last per delta
   print(noquote(format(DriftPerDelta, width=12, digits=4)))                                                 #print on screen drift of peaks per Delta
   cat("\nWAVELENGTHS (Deltas) OF PEAKS AND DIPS:-\n")                                                       #title on screen
   WaveLengths <- data.frame(rbind(FirstWaveL.peaks, LastWaveL.peaks, AnnualWaveLDrift.peaks, FirstWaveL.dips, LastWaveL.dips, AnnualWaveLDrift.dips))  #bind rows into data.frame for neat printing
   print(noquote(format(WaveLengths, width=12, digits=4)))                                                   #print wave lengths on screen
   cat("\nAMPLITUDES (Bmass) BETWEEN PEAK and FIRST SUBSEQUENT DIP:-\n")                                     #print amplitudes on screen
   Amplitudes <- data.frame(rbind(FirstAmplitude, LastAmplitude, AnnualAmplitudeDrift))                      #amplitudes and their drift per delta
   print(noquote(format(Amplitudes, width=12, digits=4)))                                                    #print amplitudes and their drift per delta
   cat("\n* * * End of Analysis of Peaks and Dips * * *\n\n\n")                                              #credits on screen
   Message <- "See the ANALYSIS of peaks and dips printed above"                                             #message to go on screen reminding user to scroll upwards
   }                                                                                                         #End of if(Choice == "A") block
return(Message)                                                                                              #return the message
}
#################################################################End of function ProcessOutput()


ReOrder <- function(Rows, Source, Stem, Suffixes, Primary, PrimaryName, ErrLine, Test){ 
#Sorts rows of data for the stem variable, read from the source file (Params or Vars), to match the ordering of names in Primary. This is used to put all rows and cols into the same order for easier processing.  Outputs re-ordered, rows and suffixes as a list of 2 items.
#Rows are those to be re-ordered, Source is the file.name whence they came, Stem is the pre-dot part, and Suffixes are the post-dot part of the row.names, Primary is the primary list whose ordering should be followed by Suffixes (from the column headers of Params), ErrLine is the calling line number in case an error is found, Test is shown below.
if(Test == "=")                                                                                              #If Suffixes should equal Primary names, ie. not be a proper subset of them:-
   {if(!setequal(Suffixes, Primary))                                                                         #If the two sets of names do not have exactly the same members:-
      {cat("\nERROR: The set of suffixes of", Stem, "variables in '", Source, "' are not exactly the same as the set of", PrimaryName, ".\nCheck the sorted lists below and adjust names in the input file:-\n\n")
      N.diff <- length(Primary) - length(Suffixes)                                                           #If Primary has more members than Suffixes, or vice versa
      if(N.diff > 0) {Suffixes <- c(Suffixes, rep(NA, N.diff))}                                              #if N.diff is +ve, supplement Suffixes with NAs to make up the length
      if(N.diff < 0) {Primary <- c(Primary, rep(NA, N.diff))}                                                #if N.diff is -ve, supplement Primary with NAs to make up the length
      Temp.matrix <- matrix(data=c(Suffixes, Primary), nrow=2, byrow=TRUE, dimnames=list(c(paste(Stem, "suffixes", sep="."), PrimaryName), NULL))
      print(Temp.matrix, quote=FALSE)                                                                        #print Suffixes and Primary with matching values one above the other
      cat("\nThe set of", Stem, "suffixes that are not members of the", PrimaryName, "set is {", setdiff(Suffixes, Primary), "}")  #advise further
      cat("\nThe set of", PrimaryName, "that are not members of the set of", Stem, "suffixes is {", setdiff(Primary, Suffixes), "}\n")
      ErrMessage <- paste("\b\b\b\b\b\b\b\nProgram terminated at ReOrder.a called from ", ErrLine, sep="")   #advise error and
      stop(ErrMessage, call.=FALSE)                                                                          #stop the program
      }                                                                                                      #the two sets of names do have exactly the same members; skip to "#Given no errors or stops:"
   }
if(Test == "<")                                                                                              #If Suffixes should be a subset of Primary names:-
   {if(length(setdiff(Suffixes, Primary)) > 0)                                                               #If Suffixes has members that Primary does not have:-
      {cat("\nERROR: The set of suffixes of", Stem, "variables in '", Source, "' has members not in the primary set,", PrimaryName, ".\nThese additional members cannot be used.  Check the sorted lists below and adjust names in the input file.\n\n")         
      cat(paste(Stem, "suffixes", sep="."), sort(Suffixes), "\n")                                            #screen names
      cat(PrimaryName, sort(Primary), "\n")                                                                  #primary name
      cat("\nThe surplus members found in", paste(Stem, "suffixes", sep="."), "are:-\n", setdiff(Suffixes, Primary), "\n") #assistance to find unknown suffixes      
      ErrMessage <- paste("\b\b\b\b\b\b\b\nProgram terminated at ReOrder.b called from ", ErrLine, sep="")   #advise error and
      stop(ErrMessage, call.=FALSE)                                                                          #stop the program
      }
   #Given no errors or stops:- 
   Ordering <- match(Primary, Suffixes)                                                                      #Get the ordering of the suffixes so that they match that of the Primary set
   Ordering <- Ordering[!is.na(Ordering)]                                                                    #remove NAs that occur when Test == "<"
   ReOrderedSuffs <- Suffixes[Ordering]                                                                      #Use the ordering to re-order the suffixes
   if(length(Ordering) > 1) ReOrderedRows <- Rows[Ordering, ] else ReOrderedRows <- Rows                     #Use it to re-order the rows of data (if() avoids dimensioning error)
   list(ReOrderedRows, ReOrderedSuffs)                                                                       #This final statement passes these two objects back to the program body
   }
} 
##################################################################  End of function ReOrder()


Restart.TS <- function(TS.matrix, Last.col, Years, NewValue){                                                #define function name and calling params
#Re-start starts time series for main-menu option 9 (over-writing); the last col of data will be moved to the first col of the new TS matrix; NewValue is the value supplied to put into New.matrix array definition
New.matrix <- array(data=NewValue, dim= c(nrow(TS.matrix), N.LPproj + 1), dimnames = list(row.names(TS.matrix), paste0(Years, "y"))) #define a local matrix; Years is a vector of dates for the time series
New.matrix[ , 1] <- TS.matrix[ , Last.col]                                                                   #move the last col of time-series matrix to the first col
return(New.matrix)                                                                                           #return the restarted matrix
}
#################################################################End of function Restart.TS()


#ShowCalcs shows internal calculations for the MM'th simulated projection
#Useful for working out how results were calculated, given the formulas in the papers
ShowCalcs <- function(MM){                                                                                   #calculate from step MM
names(LP.out$soln) <- ObjFun.names                                                                           #objective-function names
names(LP.out$slack) <- row.names(All.C)                                                                      #slack variable row names
cat("\n                        PARAMETERS AND VARIABLES FOR SIMULATION STEP [M] = ", MM, " at Years[M] = ", Years[MM], ":\n")  #screen title
cat("*****Biomasses and parameters used for simulation *****\n")
print(Bmass[ , MM])                                                                                          #screen biomass
TempDataFrame <- data.frame(t(SelfGrow), t(SelfLoss), t(ParticulateSL))                                      #store these cols in temporary dataframe for easy printing
print(t(TempDataFrame[is.WildOrNLO, ]))                                                                      #print on screen temp dataframe
print(Harv[is.WildOrNLO, !is.Auto ])                                                                         #print selected rows and cols of Harv
print(Growth[is.WildOrNLO, !is.Auto])                                                                        #ditto Growth
cat("\n*****Projected-derivative matrices *****\n")
print(projD1.pGains[!is.H2O, !is.H2O])                                                                       #print projected D1 gains
print(projD1.pLosses[!is.H2O, !is.H2O])                                                                      #print projected D1 losses
print(D1.BmassByProj[!is.H2O, MM])                                                                           #print projected D1 biomasses
cat("\n")
print(projD2.pGains[!is.H2O, !is.H2O])
if(proj.SLorder > 1)                                                                                         #If the Taylor-series order of approximation was > 1 [as recommended for LV series only] :-
   {print(projD2.pLosses[!is.H2O, !is.H2O])
   print(D2.BmassByProj[!is.H2O, MM])
   } else {cat("\n*****No projD2.pLosses or D2.BmassByProj matrices were required for proj.SLorder= ", proj.SLorder, "\n")}
cat("\n")
print(projD3.pGains[!is.H2O, !is.H2O])
if(proj.SLorder > 2)                                                                                         #If the Taylor-series order of approximation was > 2 [as recommended for LV series only] :-
   {print(projD3.pLosses)
   print(D3.BmassByProj[!is.H2O, MM])
   } else {cat("\n*****No projD3.pLosses or D3.BmassByProj matrices matrices were required for proj.SLorder= ", proj.SLorder, "\n")}
cat("\n*****Projected partial gains to Taylor-series order 3, and losses to T-s order ", proj.SLorder, ", Delta = ", Delta, " *****\n")  #title on screen
print(proj.pGains[!is.H2O, !is.H2O])                                                                         #print all outputs
cat("\n")
print(proj.SelfLosses[!is.H2O, MM])
cat("\n*****Constraints and right-side maximum limits *****\n")
Constr.used <- All.Cmax - LP.out$slack
colnames(Constr.used) <- "Used"
Constr.dataframe <- data.frame(All.Cmax, LP.out$slack, Constr.used)                                          #combine these variables as columns
print(t(Constr.dataframe))                                                                                   #print as rows by transposing the data.frame
if(N.Nutr > 0)
   {cat("\n*****Nutrients at the start of step", MM, " *****\n")
   Nutr.dataframe <- data.frame(Nmasses[ , MM])
   print(Nutr.dataframe)
   if(N.NLOs > 0)   
      {Nutr.dataframe <- data.frame(NconcInNLO[ , MM])
      print(Nutr.dataframe)
      }
   }
if(N.Habitats > 0)
   {cat("\n*****Essential-habitat sizes available for occupation *****\n")
   InitSize <- HabSize.init[which(is.Habitat)]
   OccupCoef <- array(data=HabOccup[ , is.Wild], dim=c(N.Habitats, N.Wilds), dimnames=list(Habitats, paste0("Occup.", Wilds)))
   Biomass <- array(data=rep(Bmass[which(is.Wild), MM], N.Habitats), dim=c(N.Wilds, N.Habitats))
   Biomass <- t(Biomass); dimnames(Biomass) <- list(Habitats, paste0("Bmass.", Wilds))
   SizeUsed <- array(data=HabOccup[ , is.Wild] %*% Bmass[which(is.Wild), MM], dim=c(N.Habitats, 1), dimnames=list(Habitats, "SizeUsed"))
   HabSizeLeft <- array(data=HabLeft[ , MM], dim=c(N.Habitats, 1), dimnames=list(Habitats, "HabitatLeft"))
   Hab.dataframe <- data.frame(InitSize, OccupCoef, Biomass, SizeUsed, HabSizeLeft)
   print(Hab.dataframe)
   }
cat("\n*****Linear Programming results *****\n")
cat(paste0("TotalGP=", LP.out$value, ", and individual pGPs:- \n"))
print(round(LP.out$soln, 5))
cat("\n***************************************************************************************************\n")   
}                                                          #end of function ShowCalcs
   

#Csums and Rsums are intended for summing cols or rows in 1 or 2d vectors, matrices, arrays or sub-arrays without distinction between them
#They may have 1 or more rows, 1 or more columns, or be dimensionless; the dim() function did not always work for all of these structures, hence dims are best provided as arguments nR and nC
Csums <- function(A, nR, nC){
if(nR == 0 | nC == 0) {Cs <- 0 ; return(Cs)}
#if(!is.numeric(nR)) nR <- 1                                                    #default number of rows
#if(!is.numeric(nC)) nC <- length(A) / nR                                       #default number of cols
A <- array(data=A, dim=c(nR, nC))                                               #fixing the type of structure is necessary for uniform performance of this function
if(nR == 1)                                                                     #If A has one row only:-
   {Cs <- A                                                                     #colsums are just the elements of A
   } else                                                                       #else
   {if(nR > 1)                                                                  #If A has more than 1 row:-
      {Cs <- colSums(A, na.rm=TRUE)                                             #colSums() works OK; NAs are removed
      }
   }   
return(Cs)                                                                      #return the calculated col sums
}

Rsums <- function(A, nR, nC){
if(nR == 0 | nC == 0) {Cs <- 0 ; return(Rs)}
#if(!is.numeric(nC)) nC <- 1                                                    #default number of cols
#if(!is.numeric(nR)) nR <- length(A) / nC                                       #default number of rows
A <- array(data=A, dim=c(nR, nC))                                               #fixing the type of structure is necessary for uniform performance of this function
if(nC == 1)                                                                     #if A has 1 col only:- 
   {Rs <- A                                                                     #rowsums are just the elements of A
   } else                                                                       #else
   {if(nC > 1)                                                                  #If A has more than 1 col:-
      {Rs <- rowSums(A, na.rm=TRUE)                                             #rowSums() works OK; NAs are removed
      }
   }    
return(Rs)                                                                      #return the calculated row sums
}   
```
