## Supplementary material for "Virtual aquatic ecology: stepped simulations of gross production within constraints explain biomass changes": ECOLPS simulator in R: Trial1.CompareLVsim2Theor.R

```
#This code was used to prepare the fig. 4 in Part I of "Constrained simulation of relative organic biomasses . . ."
r.a <- c(0.4, 0.5, 0.62, 0.75, 0.87, 1, 1.2)                     #values of self-growth coefficient
#Sim <- c( 0.375,  0.625,  0.925,  1.25,  1.55,  1.875, 2.375)   #simulated steady-state bmasses of heterotroph h using 1st order self losses in food constr
Sim <- c( 0.375,  0.626,  0.925,  1.25,  1.55,  1.875, 2.375)    #steady-state bmasses of h using 3rd order self losses in food constr
Theo <- c(0.375, 0.625, 0.925, 1.25, 1.55, 1.875, 2.375)         #theoretical biomasses calculated with eq. 9 for B_h
B.h <- data.frame(r.a, Sim, Theo, row.names=1:7)                 #make a data frame for the plot
#Simulated <- c(0.0167, 0.0213, 0.0256, 0.0303, 0.0339, 0.0385, 0.0435) #simulated frequencies with each r.a using 1st order self losses in food constr; obtained with A option of Variables menu in LINSIM
Simulated <- c(0.0154, 0.020, 0.0244, 0.0278, 0.0313, 0.0345, 0.0385) #frequencies using 3rd order self losses in food constr
Theoretical <- c(0.194, 0.25, 0.304, 0.35, 0.394, 0.433, 0.487)  #theoretical values calculated for frequencies, phi, with eq. 11 of Part I
phi <- data.frame(r.a, Simulated, Theoretical, row.names=1:7)    #make a data frame for the plot
graphics.off()                                                   #clear all graphs on screen
windows(width=6, height=4, restoreConsole=TRUE)                  #open a new window for the plot (required here)
op <- par()                                                      #store current graphic parameters of R
par(mfrow=c(1, 2), omi=c(.3,.3,0,0), mar=c(3,2.5,2,2))   #oma=c(1,1,0,0), #change some of them
plot(Sim ~ Theo, data=B.h, pch=3, cex=1.0, las=1, cex.lab=1.4,   #plot biomass points
    cex.axis=1, xlab="", ylab="", lwd=3, xlim=c(0.35,2.5), ylim=c(0.35, 2.5))
abline(lm(Sim ~ Theo), lwd=2)                                    #fit a line
title(main="Biomass")
plot(Simulated ~ Theoretical, data=phi, pch=3, cex=1.0, las=1, cex.lab=1.4, #plot frequency points
    cex.axis=1. , ylab="", xlab="", lwd=3, xlim=c(0.15,0.5), ylim=c(0.015, 0.045))
abline(lm(Simulated ~ Theoretical, data=phi), lwd=2)             #fit a line
title(main="Frequency")
mtext("Simulated", side=2, outer=T, cex=1.5)                     #add marginal text headings
mtext("Theoretical", side=1, outer=T, cex=1.5)
par <- op                                                        #restore original graphic parameters
```
